# Effects of antibiotics on the abundance of antibiotic resistance determinants during and after antibiotic administration to beef cattle: A systematic review and meta-analysis of longitudinal studies

**DOI:** 10.64898/2026.05.31.729110

**Authors:** Matt Lloyd Jones, Alfredo Sánchez-Tójar, Alison Bethel, Anne Frances Clare Leonard, Emma Lamb, Natalia Casanova, Johana Dominguez, María Paula Quiroga, Daniela Centron, Adriana Peralta Alonso, Mariano Fernández-Miyakawa, William Gaze, Alejandro Petroni, Ruth Garside

## Abstract

**Background:** Beef feedlots are increasing concerns that antibiotic use in beef cattle selects for antibiotic resistance, but limitations of primary studies and previous syntheses make it difficult to confirm a consistent effect. We conducted a rigorous systematic review and meta-analysis to summarise: 1) the effect during and after antibiotic administration; 2) its moderation by time since administration started/ended.

**Methods:** Eligible studies longitudinally compared beef cattle administered antibiotics to those that were not, measuring resistance determinants in faeces and/or environments. Information sources included Web of Science, CAB Abstracts, and Medline (last searches: 05/03/25). Risk of bias was assessed using RoB 2 and ROBINS-I. Meta-analysis was conducted where feasible, using individual participant data where necessary.

**Results:** The 33 included studies were mostly small trials of North American feedlot cattle, all with high risks of bias. Meta-analysis of 11 studies of tylosin, ceftiofur, and chlortetracycline indicated positive effects on absolute abundance of resistance both during (SMDH = 0.4; 95% CI = 0.11 to 0.69, p = <0.01) and after (SMDH = 0.52; 95% CI = 0.33 to 0.71, p = <0.01) antibiotic administration. Log-transformed time was positively associated with effect size during (Slope = 0.63; 95% CI = 0.1 to 1.16, p = 0.02), and negatively associated after (Slope = -0.65; 95% CI = -1.24 to -0.06, p = 0.03)

**Discussion:** Available evidence indicates time-dependent selection for antibiotic resistance in beef cattle, warranting further regulation to limit human health risks. Simultaneously, uncertainty about precise effect sizes warrants further research.

**Funding:** BBSRC

**Registration:** https://doi.org/10.17605/OSF.IO/RXQHT

## 1 Introduction

### 1.1 Rationale

Whilst beef farming consumes relatively low quantities of antibiotics compared to other livestock sectors such as dairy, poultry and pig farming^1,2^, the global spread of intensive beef cattle production systems may be changing this. In particular, North American beef feedlot systems associated with higher antibiotic use are steadily becoming the system of choice in other parts of the world such as Australia, Latin America and Asia^3^. In these systems, antibiotics are fed and injected for prophylactic, growth promotion, and therapeutic purposes – sometimes with daily and mass administration — presenting an obvious potential threat to the ongoing efficacy of antibiotics in both veterinary and human medicine^4^. This is because such widespread antibiotic use risks selecting for the spread of antibiotic resistance both on-farm (e.g. through faecal transmission of antibiotic-resistant bacteria and genes to animals and humans) and off-farm (e.g. through faecal contamination of food and the environment)^5^.

A 2017 WHO-commissioned systematic review of mainly cross-sectional studies concluded that broad interventions to restrict antibiotic use in food-producing animals are associated with a reduced proportion of antibiotic-resistant bacteria and genes being isolated from them, directly informing WHO guidelines on reducing antibiotic use in livestock generally^6,7^. However, direct evidence for a causative relationship between antibiotic use in beef cattle and increased antibiotic resistance has eluded comprehensive synthesis, with two recent systematic reviews encountering several challenges. A 2020 systematic review of tylosin administration^8^ tentatively concluded that administering tylosin to cattle is associated with an increased proportion of macrolide-resistant bacteria isolated from them (compared to no tylosin administration), based on a semi-quantitative analysis of 4 studies due to the low number of studies identified and their inconsistent reporting. A more recent systematic review^9^ on the effect of antibiotic use on antibiotic resistance in Salmonella encountered similar difficulties, concluding that the certainty of the evidence base was too low to be able to draw meaningful conclusions (e.g. to inform quantitative microbial risk assessment). Thus, whilst these reviews make an important contribution by highlighting the difficulty of drawing firm conclusions from the current evidence base, we are still lacking a comprehensive review conclusively answering the question of whether antibiotic use in beef cattle production selects for antibiotic resistance.

This review aims to comprehensively assess the evidence for antibiotic use in beef cattle production systems selecting for antibiotic resistance, improving upon previous efforts in several ways. Firstly, our systematic review builds on the two previous reviews of the causative evidence^8,9^ by adopting a broader scope including all antibiotics tested as interventions, and all microorganisms and resistance genes tested as outcomes in its eligibility criteria. This allows us to answer the question of whether antibiotic use in beef cattle production in general drives antibiotic resistance, capitalising upon the broader evidence base and dealing with any additional heterogeneity through synthesis methods. Secondly, our meta-analysis directly engages with the potential moderation of this effect by the timing of its assessment during and after antibiotic administration. Previous reviews only used this longitudinal information was only in a semi-quantitative way^8^ or only used the last sampling point from each longitudinal transect^9^. Finally, our approach is underpinned by a systematic review methodology that more closely adheres to contemporary guidelines. In particular, we sought to balance our methodological innovation with standardised and transparent methods for conducting and reporting searches, screening, risk of bias, certainty assessments (Supplementary Table 1), and any deviations from protocol (Supplementary Table 2).

### 1.2 Objectives

Our review objective is articulated in the following PICO-structured research question^10^:

*In beef cattle (P), what is the effect of administering an antibiotic (I) versus not administering that antibiotic (C) on the abundance of resistance determinants (e.g. resistant bacteria, genes) for that antibiotic’s class in their faeces and environments impacted by it (O)?*

We tested two primary hypotheses to answer this question:

*H1. When an antibiotic is administered to beef cattle, there will be a higher abundance of resistance to that antibiotic in the faeces and environments of the cattle to which it is administered, compared to that of non-administered cattle*.

*H2. The strength of this effect (H1) will increase over time during the period of antibiotic administration, and decrease over time after the cessation of antibiotic administration*.

## 2 Methods

### 2.1 Registration and reporting

A protocol for this systematic review and meta-analysis was pre-registered at Open Science Framework on November 3rd, 2020 under the DOI 10.17605/OSF.IO/RXQHT^11^, with the final systematic review and meta-analysis reported following PRISMA2020 guidelines^12^ and a checklist included in Supplementary Table 1. During the process of conducting the review, some deviations that we considered necessary were made. The most significant of these was re-focusing the quantitative synthesis on the effect on absolute rather than relative abundance of resistance determinants for interpretability and statistical reasons (Supplementary Information 1). All deviations are fully reported and justified in Supplementary Table 2 using a standardised reporting tool^13^.

### 2.2 Eligibility criteria

We considered studies to be eligible for answering our research question according to our pre-defined PICO(S) criteria (Supplementary Information 2), which are summarised as:

- **Population:** Cattle being raised for beef production and consuming only solid food (i.e. weaned).
- **Intervention(s):** Administration of an antibiotic to beef cattle either through feed or injection. An intervention was considered eligible provided there was a measure of resistance to the antibiotic class to which it belonged (see Outcome). Indirect administration of antibiotic contaminants of other media (e.g. the common practice of feeding cattle spent distiller’s grains from the fuel ethanol industry, which can contain virginiamycin as a contaminant) is eligible provided the dose of the contaminant antibiotic is quantified.
- **Control(s):** No administration of the antibiotic that defines the intervention (though other antibiotics may be given to both intervention and control/comparator groups).
- **Outcome:** An estimate of either: 1) the absolute abundance of both resistant and all bacteria in the targeted population; or 2) the relative abundance of resistance in the targeted population. Estimates of absolute abundance refer to either culture-dependent methods to quantify the density of resistant and all bacteria in the target population (e.g. resistant cells per gram of faeces, all cells per gram of faeces measured via direct plating), or culture-independent methods to quantify the density of resistance and bacterial marker genes (e.g. genes per gram of faeces, 16S genes per gram of faeces measured via qPCR or metagenomics). Estimates of relative abundance can include direct plating and qPCR methods which may be presented or reformulated as relative abundance (e.g. resistance genes per 16S copies), as well as antimicrobial susceptibility testing methods which can only estimate relative abundance (e.g. proportion of 10 tested isolates that were resistant). The targeted resistance phenotype or genotype should relate specifically to the class of the antibiotic administered in the intervention (i.e. direct selection). The targeted bacterial populations should be from samples from fresh beef cattle faeces (direct from the rectum or deposited <24 hours before) or beef cattle associated environments (i.e. pen soil, water or feed). The bacterial populations in which resistance is assessed should be established bacteria rather than bacteria with which the animal has been experimentally colonised or infected (though it is fine for studies to use experimental colonisation/infection as a cointervention alongside an antibiotic intervention).
- **Study designs:** Longitudinal studies, minimally defined as those with a baseline measurement before the start of the intervention and at least one measured after the start of the intervention.

### 2.3 Information sources & search strategy

The information sources and search strategy were chosen and executed primarily by an Information Specialist (AB), working with a Latin American librarian (APA) and the wider multi-disciplinary team. The main set of searches comprised English-language searching of key bibliographic databases: Clarivate Web of Science core collection (SCI-EXPANDED, SSCI, A&HCI, CPCI-S, CPCI-SSH, ESCI; last searched 4 April 2025), Ovid CAB Abstracts (last searched 4 April 2025), Ovid MEDLINE (last searched 4 April 2025), Scopus (last searched 7 July 2020), Ovid Global Health (last searched 7 July 2020), CABI VetMed Resource (last searched 24 September 2020), Google Scholar (last searched 14 July 2020), EBSCO Environment Complete (last searched 30 September 2020), NAL Agricola (last searched 7 July 2020), LILACS (last searched 7 July 2020), PQDT Global (last searched 7 July 2020), British Library Explore (last searched July 2020). AB and APA also conducted English and/or Spanish-language searches of databases and grey literature sources more focused on Latin America. This was to find relevant evidence from Latin America, which contains several of the world’s largest beef producers and is experiencing a shift toward intensive production systems. These sources were LILACS (last searched in Spanish 12 February 2021) Google Académico (last searched in Spanish 6 April 2021), LA Referencia (last searched in English & Spanish 19 February 2021), SIDALC (last searched in English & Spanish 24 February 2021), REDIB (last searched in English & Spanish 18 February 2021), SciELO (last searched in Spanish 7 April 2021), SNRD (last searched in Spanish 17 February 2021), and Redalyc (last searched in English & Spanish 25 March 2021). All these bibliographic databases were initially searched in 2020 to obtain an initial set of included studies and identify that minimal set of databases that needed to be searched to obtain them. This minimal set of Clarivate Web of Science, Ovid CAB Abstracts, and OVID MEDLINE was then included in search updates in 2022, 2024, and 2025 (hence why their last search dates are April 2025). Citation searching on the final set of included studies was not performed because a test performed in 2021 revealed that citation searching using four different methods did not capture any includable studies could not be found with a simpler search update.

The full search strategies for each information source and search strategies can be found in Supplementary Information 3 but briefly, were broad searches for records relating to cattle and antibiotic resistance, with faecal-related outcome measures.

### 2.4 Selection process

Records (mostly titles and abstracts of scientific publications) obtained from searches were first deduplicated in EndNote^14^, with further duplicates removed in the screening platform Rayyan (see below). Records obtained from the search updates were deduplicated against both themselves and previously-screened records. However, as this was not always effective because of re-indexing by the bibliographic database providers, we also removed records published in the period covered by the last search (which should have been screened already). The remaining records were then double-blind screened according to our eligibility criteria via the Rayyan platform^15^. MLJ double screened all records, whilst a second reviewer (AL, AST, EL, DC, JD, MPQ, NC or RG) screened a random but unique subset of these records. Conflicts between reviewers were resolved through discussion, with a third reviewer (AP) brought in to resolve any conflicts remaining after discussion. After reaching a consensus on inclusions at record stage, MLJ then attempted to retrieve reports (i.e. full texts — usually in the format of a PDF of scientific publication) for each of them. The review team then repeated the screening process implemented at record stage for the retrievable reports.

The study selection process is documented in a PRISMA2020 flow diagram, produced using the raw bibliographic files for each set and subset of articles, MLJ’s own in-development package (RayyanR v0.22) for interacting with the Rayyan API^16^, and the ‘PRISMA2020’ R package version 1.1.2^17^.

### 2.5 Data collection process and data items

For all studies, we extracted data relating to population, intervention(s), control(s), outcome(s) and study for each study for each publication in a study characteristics table (Supplementary Information 4). For population, we sought information on the type (e.g. feedlot steers). For interventions, we sought information on the antibiotic administered, its dose, and cointerventions for each intervention arm. For controls, we sought information on any cointerventions (defined as antibiotics that were given to both the intervention and control arm). For outcome(s) we sought information on the method and matrix (e.g. faeces) used to measure the abundance of resistance/all bacteria (e.g. direct enumeration), and the reported effects on absolute and/or relative abundance in the publication. For study, we matched each publication to the unique study on which it reported (assigning each a name in the format ‘Institution - Interventions’), and sought information on the study design and number of animals/pens in the intervention/control arms. For practical reasons, these data were extracted by one reviewer (MLJ) and checked by a second reviewer (discussing and resolving any discrepancies with MLJ). Where necessary, MLJ emailed the authors to clarify details.

### 2.6 Study risk of bias assessment

Risk of bias was assessed for all included using Cochrane’s Risk of Bias 2 (RoB 2) tool for randomised controlled trials, and Cochrane’s ROBINS-I tool for non-randomised/observational studies. Risk of bias was assessed by a single reviewer (MLJ) but checked by a second using these tools at the publication, rather than individual outcome level. This made assessing these studies — which frequently had multiple study arms, outcome measures and sampling points — more manageable, taking advantage of the fact that there was typically some methodological unity between effect sizes reported in the same publication (e.g. one publication relating to a study might report culture-based outcomes whilst another might report culture-independent outcomes from the same samples). Hence, groups of outcomes reported in the same publication — for example, different resistance genes or relative and absolute measures of resistance (see next section) — were therefore considered simultaneously when assessing risk of bias (e.g. baseline differences in the outcome, measurement of the outcome).

### 2.7 Effect measures

To compare effects of antibiotic administration on antibiotic resistance across studies in a standardised way, we aimed to estimate the standardised mean difference accounting for heteroscedasticity (SMDH). Specifically, we sought to use the SMDH to estimate the difference between the mean absolute abundance of resistant bacteria/resistance genes between bacterial populations associated with beef cattle that had been administered antibiotics (the intervention group), and the mean absolute abundance of resistant bacteria/resistance genes in bacterial populations associated with beef cattle that had not been administered antibiotics, in terms of standard deviations (SDs). The SMDH^18,19^ was specifically chosen as the effect measure because — given the comprehensive scope of our synthesis — our eligibility criteria resulted in a set of included studies that used different methods with very different scales of measurement and errors (e.g. culture-based versus qPCR based measures, different target bacterial species, different target resistance genes). SMD(H)s are designed to cope with such differences in method and scale of measurement by using the distance in terms of SD as the measure of the effect. SMDHs were never reported in publications themselves, so we sought to calculate them ourselves. The minimal level of data required for this were the means, standard deviations (SDs), and sample sizes of the absolute and relative abundance of resistant bacteria/resistance genes in the bacterial populations sampled, for both the intervention and control/comparator groups in each study for every time point measured in the study.

### 2.8 Synthesis methods

#### 2.8.1 Selection process

All studies selected for inclusion in the systematic review were included in an initial narrative synthesis, regardless of whether we could obtain sufficient data to calculate the SMDH. To further permit their inclusion in the quantitative synthesis MLJ requested the raw data necessary to calculate SMDHs — given outcomes were frequently reported in scientific publications in ways not immediately amenable to this (e.g. selected outcomes, marginal means, difficult-to-extract figures). This was done by sending at least two email requests to the first, corresponding, or last author(s) of each study, with follow-up emails to clarify details where necessary. Where study data was only available in the format of published figures, MLJ extracted it from published reports and their figures manually and using WebPlotDigitizer version 5.2^20^.

Studies for which we could obtain the data necessary to calculate SMDHs in this way were included in the quantitative synthesis — provided they assessed the antibiotic resistance outcome using direct plating and/or PCR methods. Outcomes assessed by antimicrobial susceptibility testing were excluded due to statistical issues with using this data to quantitatively estimate the abundance of resistance that only became clear after obtaining study data (Supplementary Table 2). Outcomes assessed by metagenomics were excluded as planned in our pre-registered protocol^11^, because by definition they target many genes (i.e. resistance outcomes) simultaneously and hence are better suited to broad-scale assessing the effect on aggregate outcomes like richness and composition of the resistome.

#### 2.8.2 Preparation of data

Data for the narrative synthesis required no preparation other than converting all antibiotic doses to mg/kg in the same way as we did for the quantitative synthesis (see below). Data for the quantitative synthesis required more extensive processing and combined with other data from reports before meta-analysis, which MLJ did using custom scripts written in the R programming language v4.3.1^21^. This process and the decisions made during it are detailed and justified in Supplementary Information 5, with the full list of extracted items detailed in Supplementary Information 6. As part of processing, measurements of the absolute abundance of antibiotic resistance determinants in the intervention and control arms across each study’s longitudinal sampling regimen were translated into SMDHs using the ‘escalc’ function of the metafor R package v4.8.0^22^.

#### 2.8.3 Results of individual studies

Before formally testing our hypotheses with the subset of studies included in the meta-analysis, the results of all studies included in the systematic review were first included in a narrative synthesis. This synthesis sought to understand the range of effects or associations reported in the relevant literature as a whole, regardless of whether these studies were suitable for inclusion in a meta-analysis. For this, we drew on the whole evidence base including studies using methods that can assess effects on the absolute and relative abundance of resistance (direct plating, qPCR and metagenomics), as well as studies using methods that can only assess effects on the relative abundance of resistance (antimicrobial susceptibility testing). This synthesis consisted of descriptive statistics of study characteristics underpinned by the data extracted into the study characteristics table, though we avoided explicit vote counting given its severe limitations as a synthesis method^23,24^.

To complement the narrative synthesis of the results of individual studies, we also quantitatively synthesised the results of individual studies included in the meta-analysis. For this, we meta-analytically aggregated the effect sizes (SMDHs) at the study level to produce a forest plot using metafor methods^25^. We produced separate forest plots for effects measured in faeces and environmental samples, and within these two groupings outcomes measures pre-antibiotic administration (i.e. baseline measurements) or post-antibiotic administration (Supplementary Information 7).

#### 2.8.4 Meta-analysis

##### 2.8.4.1 Meta-analysis of the overall effect (hypothesis 1)

All quantitative synthesis was performed in v4.3.1 of the R programming language^21^, with meta-analysis primarily performed using the *metafor* R package v4.8.0^22^. Given our hypotheses centred around the effect of antibiotic administration on antibiotic resistance during and after antibiotic administration (see ‘Introduction: Objectives’), we first split post-antibiotic administration data for faecal outcomes into sub-datasets relating to 1) effects measured during antibiotic administration and 2) effects measured after antibiotic administration (defined as at least 1 day since the last antibiotic administration event). Pre-antibiotic administration data were excluded from the synthesis because including them would result in inaccurate estimates of the effect due to antibiotic administration (since antibiotics had not yet been administered for these samples). Data from environmental samples were excluded from the synthesis because their scarcity precluded meta-analysis (see Results).

To test our first hypothesis, we fit intercept-only random effects models fit to ‘during’ and ‘after’ data subsets, implemented as multilevel meta-analytical models using the ‘rma.mv’ function in *metafor*. These models included the following three random effects:

1. **Study ID** to account for non-independence among effect sizes from the same study. We used the actual study (e.g. trial) rather than publication (i.e. scientific paper) as the study ID, since a few publications reported alternative outcomes from the same study.
2. **Timing of sampling nested within study arm** to account for temporal autocorrelation between repeated measures taken within the same study arm (i.e. an intervention or control/comparator group). This was modelled with a continuous-time autoregressive structure (“struct = ‘CAR’” option in rma.mv). More information about this approach can be found in metafor documentation^26^ and an example implementation^27^.
3. **Unit-level observation ID** to account for residual within-study variance. This is to account for any residual variance between multiple effect estimates from the same study, after accounting for the effects of study and repeated sampling within the same study arm (including alternative assessments of the outcome at the same time point within the same study arm). More information about this approach can be found in metafor documentation^29^ and example implementations^31^.

The response variable in all models was the SMDHs representing the effect on the absolute abundance of resistance determinants, weighted by their associated sampling variances. We also fit equivalent models for the absolute abundance of all determinants and their associated sampling variances. This allowed us to contextualise the effect on the resistant portion of the bacterial population in terms of changes in the wider bacterial population (i.e. the relative effect), without introducing interpretively and statistically problematic relative abundance measures into our primary synthesis (Supplementary Information 1).

The presence/absence of heterogeneity in each of the meta-analytical models was primarily assessed using both the Q test^32^. We also quantified heterogeneity using the I^2^ statistic using the Jackson approach^33^ for multilevel meta-analytic models, which compares the variance-covariance matrix of the fixed effects of the model with and without random effects. I^2^ estimates were interpreted sensu Higgins^34^, using the same terminology to describe them when reporting results. Confidence and prediction intervals were also computed within *metafor*, meaning that prediction intervals were identical to the confidence interval when τ^2^ = 035.

##### 2.8.4.2 Meta-analysis of the moderation of the effect by timing of sampling (hypothesis 2)

To test our second hypothesis, we tested the extent to which timing of sampling could explain any heterogeneity identified in the intercept-only models. This was done by adding timing of sampling as a fixed effect/moderator to the previously intercept-only (random effects) models, making them uni-moderator multilevel (mixed effects) meta-regressions. We tested both linear (‘H2 During Linear’ model & ‘H2 After Linear’ model) and non-linear relationships of timing of sampling (‘H2 During Nonlinear’ model & ‘H2 After Nonlinear’ model), with the latter fit as a restricted cubic spline^36,37^. Given the relatively small number of studies available for meta-analysis, the latter was fit with the minimum of 3 ‘knots’ (the points in the time series where different cubic polynomials are joined), resulting in 2 polynomials/coefficients fit to the effect of time. The importance of the moderator was assessed through a combination of whether the model converged, omnibus tests of moderators (Q_m_ tests), and model coefficients — with convergence considered the first evidence of fit, a statistically significant p-value for the Q_m_ test considered the second piece of evidence of fit, a statistically significant p-value for the timing of sampling coefficient (linear model) or at least one of the splines (restricted cubic spline) considered the third piece of evidence of fit. Again, the above process was repeated for effects on the absolute abundance of all determinants.

#### 2.8.5 Sensitivity analyses

Given variation in effect sizes can be driven by (meta)analytical decisions^38^, we subjected our candidate models for hypotheses 1 and 2 to several sensitivity analyses (Supplementary Information 8). Briefly, these were: 1) a leave-one-out analysis; 2) removing/correcting potentially problematic studies; 3) assuming different degrees of correlation between effect sizes from the same study^39,40^.

#### 2.8.6 Subgroup and other alternative analyses

We conducted various subgroup analyses (i.e. of categorical moderators) and other alternative analyses (i.e. of non-categorical moderators) to explore alternative hypotheses for heterogeneity among effect sizes. These were conducted on the whole post-antibiotic administration dataset without splitting into ‘during’ and ‘after’ periods to maximise the power of subgroup analyses (representation in each subgroup) and because these analyses were considered as alternative moderators of the effect to the period and timing of sampling. The alternative moderators considered fell into PICO(S) groupings and were:

- **Population:** Reproductive class; breed; weight; age; antibiotic treatment history of the cattle studied.
- **Intervention(s) and control(s)/comparator(s):** Antibiotic class administered in the intervention; antibiotic dose administered in the intervention (expressed as daily dose; log10-transformed daily dose, cumulative daily dose before each effect estimate, and log10-transformed cumulative daily dose before each effect estimate); whether or not the intervention and control/comparator groups were also administered other (cointervention) antibiotics; whether or not the intervention and control/comparator cattle were housed in the same enclosure.
- **Outcome:** Microorganisms targeted; method used to measure resistance.
- **Study design:** The overall design of the study (simplified to observational, completely randomised, or randomised complete block designs)

### 2.9 Reporting bias assessment

To further understand the reliability of our quantitative synthesis, we tested for reporting bias (aka publication bias) in the SMDHs underpinning our tests of hypotheses 1 and 2. The overall effect during (“During H1” model), the linear moderating effect of time during (“During H2” model), the overall effect after (“After H1” model), and the linear moderating effect of time after (“After H2” model) were considered as distinct review outcomes for publication bias testing and certainty assessment. Following Nakagawa^41^, both small sample size and time-lag publication bias were tested for each of these four Review Outcomes (Supplementary Information 9).

### 2.10 Certainty assessment

We assessed certainty in the size of the overall effect during (“During H1” model), the linear moderating effect of time during (“During H2” model), the overall effect after (“After H1” model), and the linear moderating effect of time after (“After H2” model). We used the standard GRADE approach^42,43^ to assess certainty separately for each of these four review outcomes from the meta-analysis (Supplementary Information 10).

## 3 Results

### 3.1 Study selection

Searches of bibliographic databases returned 18154 records (i.e. titles and abstracts), with 9035 after pre-screening de-duplication (Supplementary Figure 1). After screening, 217 records were identified as eligible for report retrieval, of which 182 reports were retrievable. The relatively high percentage of unretrievable reports (16.13%) was investigated and found to be mainly related to apparently undigitised publications from the late 1960s to early 1990s (median = 1984) and/or records that were difficult to link to reports because bibliographic metadata was very limited or had been translated from another language. The 182 retrieved reports were assessed for eligibility, resulting in the exclusion of 152 reports. This resulted in the initial inclusion of 30 reports in the review, with an additional 8 reports identified via search updates (Supplementary Figures 2-4). No new studies were identified from Latin American searches (Supplementary Figure 5). Exclusion reasons are detailed in Supplementary Table 3.

This process resulted in the inclusion of a total of 38 reports in the systematic review reporting on the results of 33 unique studies (Figure 1). All reports were journal papers^44–81^.

**Figure 1:**
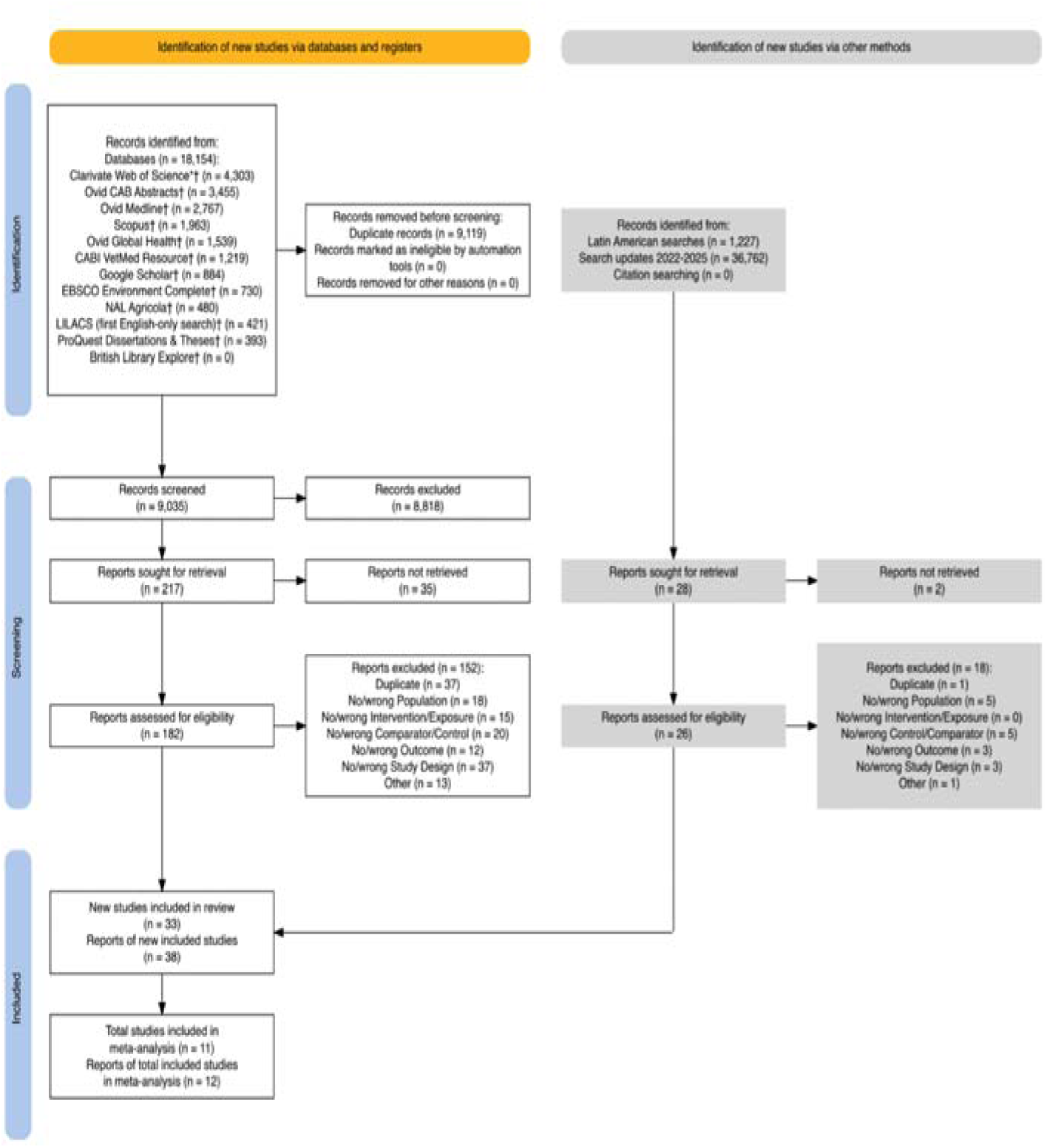
PRISMA2020 flow diagram of the systematic review process for the results of all search methods. *Specific databases searched via the Web of Science platform were SCI-EXPANDED, SSCI, A&HCI, CPCI-S, CPCI-SSH, ESCI. †Searched in English. ‡Searched in Spanish. N.B. LILACS English & Spanish language searches are listed separately because they were searched at different times

### 3.2 Study characteristics

#### 3.2.1 Population

All 33 study populations were beef cattle, with 27 being raised in feedlot-type facilities or actual feedlots (reflecting the predominance of North American studies, 4 in indoor facilities (e.g. winter and/or veal housing), and 2 on pasture. 23 of these 33 study cattle populations were in the United States, 7 in Canada, 1 in Uruguay, 1 in France, and 1 in the United Kingdom. Publications did not always report the year in which the populations were studied, but the studies were published between 1977 and 2025 (Median = 2016).

#### 3.2.2 Interventions

Interventions belonging to an antibiotic class to which resistance was assessed were included. 19 of the studies contained a single intervention arm, whilst 14 contained multiple intervention arms. Given the latter, there were a total of 55 interventions arms across the studies.

The antibiotics defining these 55 intervention arms (i.e. not administered in the control/comparator group) were the tetracyclines chlortetracycline and oxytetracycline (18 intervention arms), the macrolides tylosin, tulathromycin and tilmicosin (17 intervention arms), the extended-spectrum cephalosporin ceftiofur (7 intervention arms); the fluoroquinolones enrofloxacin and marbofloxacin (3 intervention arms); the tetracycline/sulfonamide complex chlortetracycline and sulfamethazine (4 intervention arms); the streptogramin virginiamycin (3 intervention arms); the phenicol florfenicol (2 intervention arms); the moenomycin complex bambermycin/flavomycin (1 intervention). Whilst the ionophore monensin was theoretically eligible for inclusion as an intervention, none of the studies included monensin intervention arms (though several included it as a cointervention alongside the intervention antibiotics; see below).

Reported doses for the 55 intervention arms (which we standardised into daily doses in milligram per kilogram of body weight) ranged from 0.001 mg/kg of body weight to 40 mg/kg of body weight.

30 of the 55 antibiotic intervention arms were administered orally, whilst 25 were administered via injection. Of the 30 administered orally, 26 were included in-feed and 4 in a free-choice mineral supplement (i.e salt-lick type medicated block). Of the 25 administered via injection, 15 were administered subcutaneously and 10 were administered intramuscularly.

#### 3.2.3 Controls

17 of the studies had a control arm for each intervention arm in the study, whilst 16 contained control arms that were shared between multiple intervention arms for the purposes of analysis. Given the latter, there were a total of 39 control arms across the 55 intervention arms.

13 of these control arms were administered a cointervention whose interaction with the intervention was also being studied (and hence was also given to the paired intervention arm). Most of these cointerventions were monensin, with 5 consisting of in-feed monensin alone, 3 of in-feed monensin and tylosin, and 1 of in-feed monensin and trenbolone acetate. 1 cointervention was in-feed chlortetracycline. Of the non-antibiotic cointerventions, 1 was trenbolone acetate, 1 was copper, 1 was a direct-fed antimicrobial.

#### 3.2.4 Outcomes

Outcomes assessing resistance to an antibiotic class administered in an intervention were included. 20 of the studies assessed a single outcome, whilst 13 assessed multiple outcomes. Given the latter, there were a total of 75 outcomes assessed across the studies.

The matrices in which the 75 outcomes were assessed were fresh cattle faeces (70 outcomes), pen surface soil (3 outcomes), animal feed (1 outcome) and trough water (1 outcome).

Target organisms for the 75 outcomes were *Escherichia coli* (27 outcomes), *Enterococcus spp.* (16 outcomes), *Salmonella* (6 outcomes), *Campylobacter spp.* (4 outcomes), and Enterobacteriaceae in general 1 outcomes). 21 outcomes targeted bacteria in general (by nature of using culture-independent methods).

Methods of assessment for the 75 outcomes were direct plating of samples onto antibiotic and antibiotic-free agar (29 outcomes), antimicrobial susceptibility testing of isolated bacteria (25 outcomes), qPCR quantification of resistance and 16S genes (18 outcomes), and metagenomics (3 outcomes).

#### 3.2.5 Study design

Among the 33 included studies (Table 1), 26 were randomised controlled trials. 14 of these had completely-randomised designs (CRD) and 12 had randomised block designs (RBD) where cattle were blocked by weight, age and/or other factors before randomisation - though accounting for blocking in the studies’ analysis (e.g. by including a random effect of block) was rare. The remaining 7 studies had non-randomised designs, of which 1 had case-control and 3 prospective cohort designs. The remaining 3 were described as randomised in published reports, but were better regarded as prospective cohort designs^54,55,80^. For two studies^80^ this was because even after attempted randomisation, each study arm was confounded by cattle source and experimental pasture, respectively. The other study^54^ was described as a completely randomised study but the primary intervention (tylosin with monensin) was assigned non-randomly based on previous antimicrobial treatment exposure (natural and conventionally-raised), with randomisation only used to assign conventionally-raised animals to a more secondary intervention (metaphylactic antimicrobials).

**Table 1:**
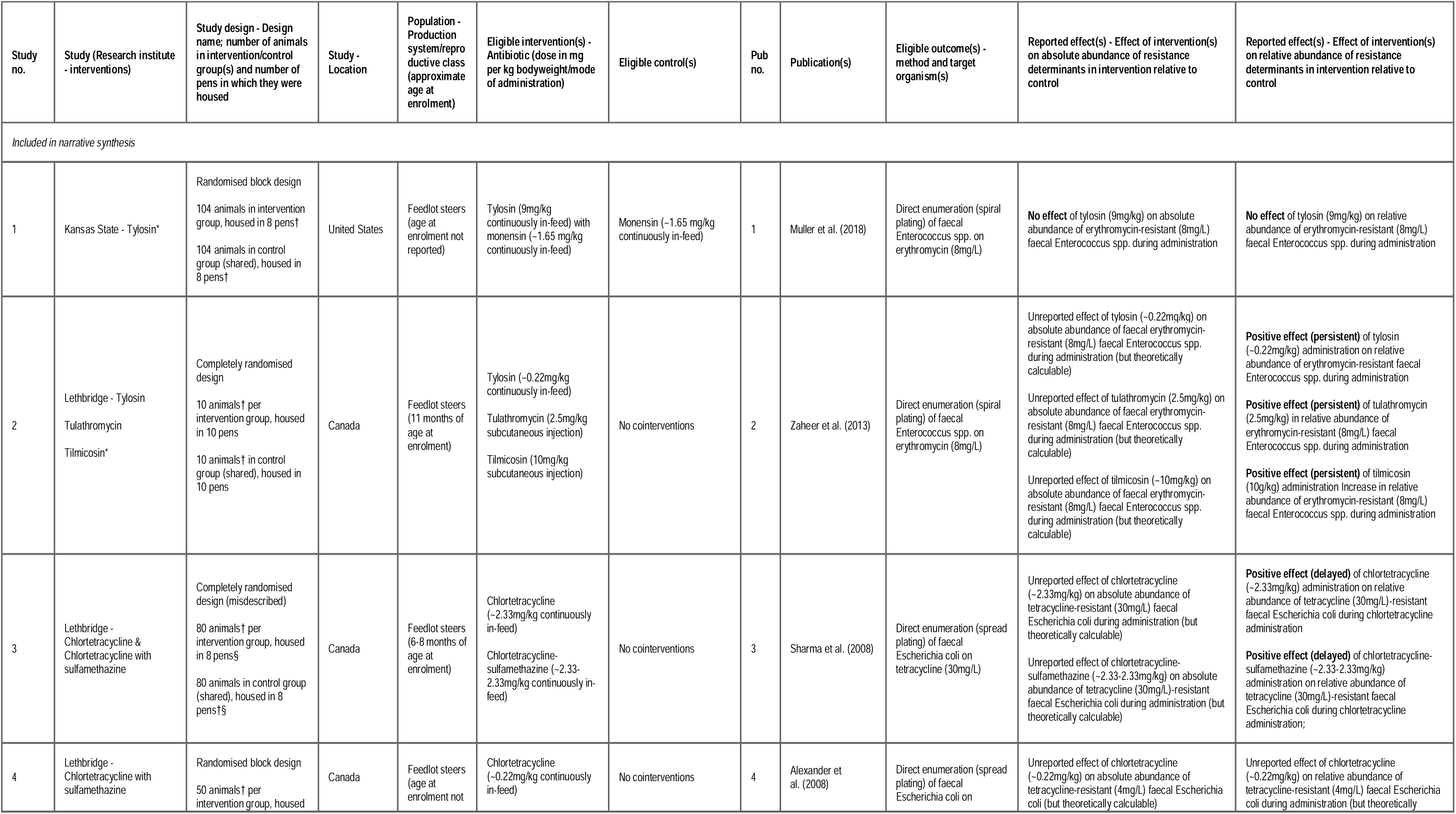
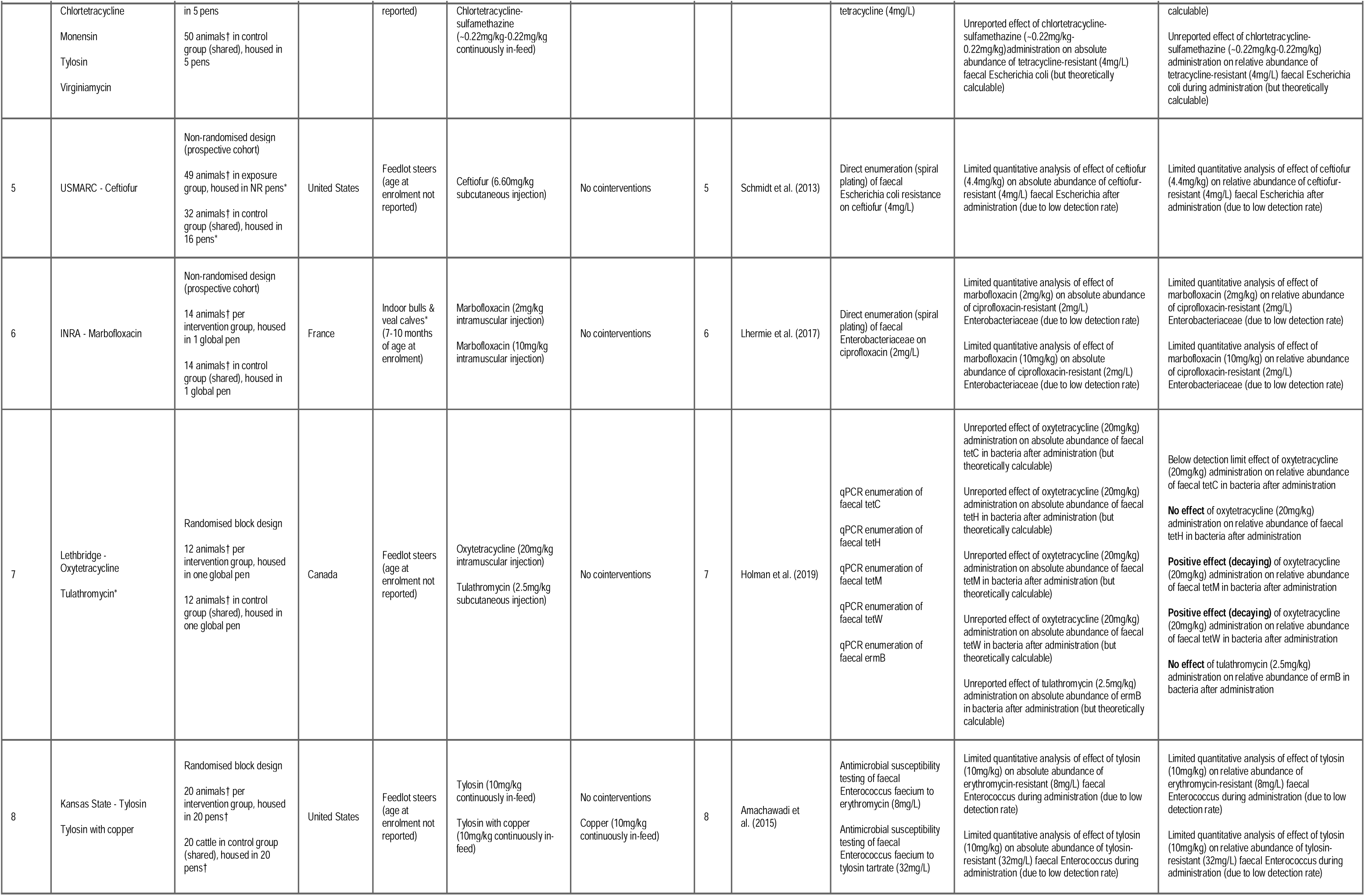
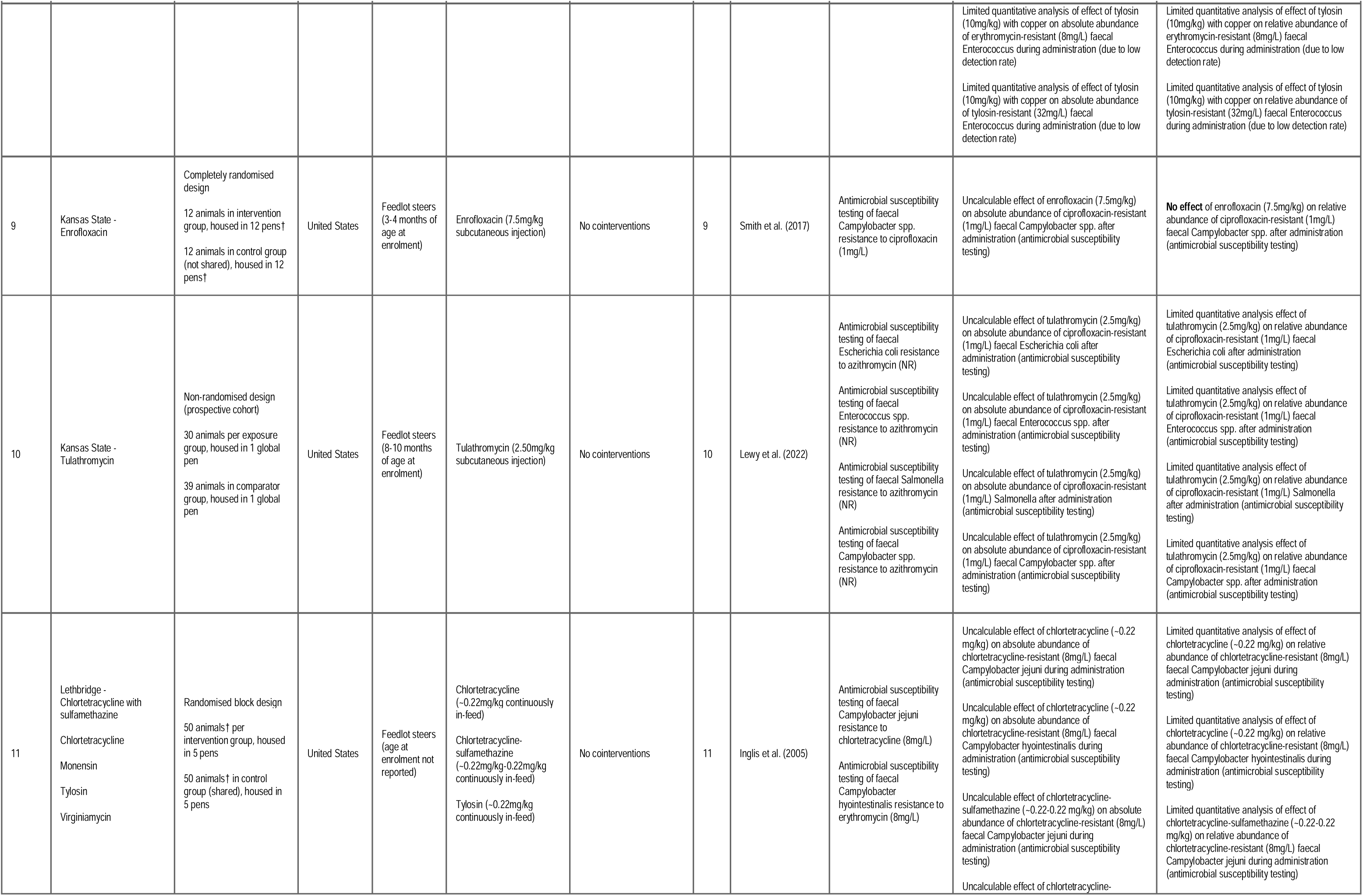
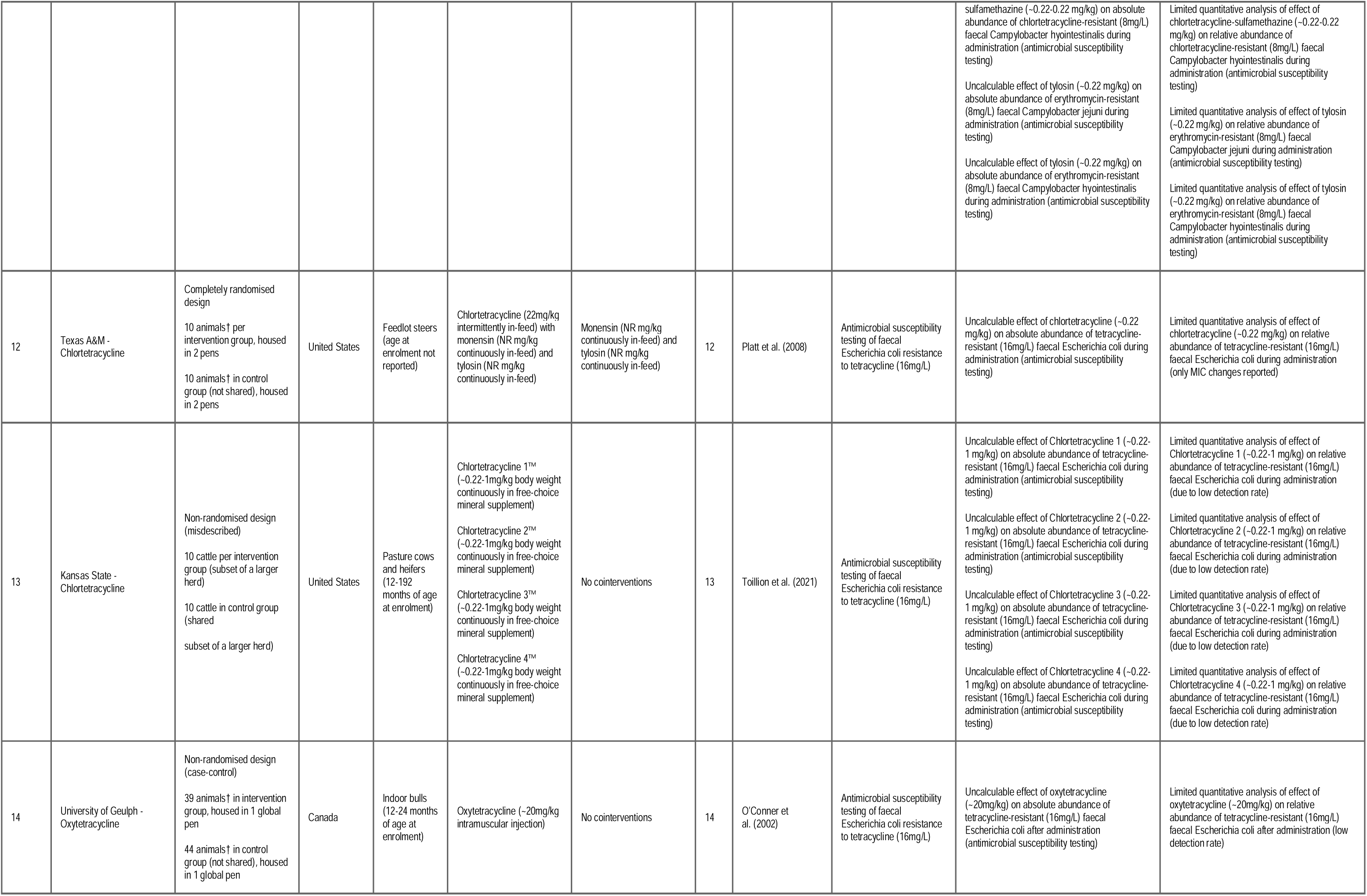
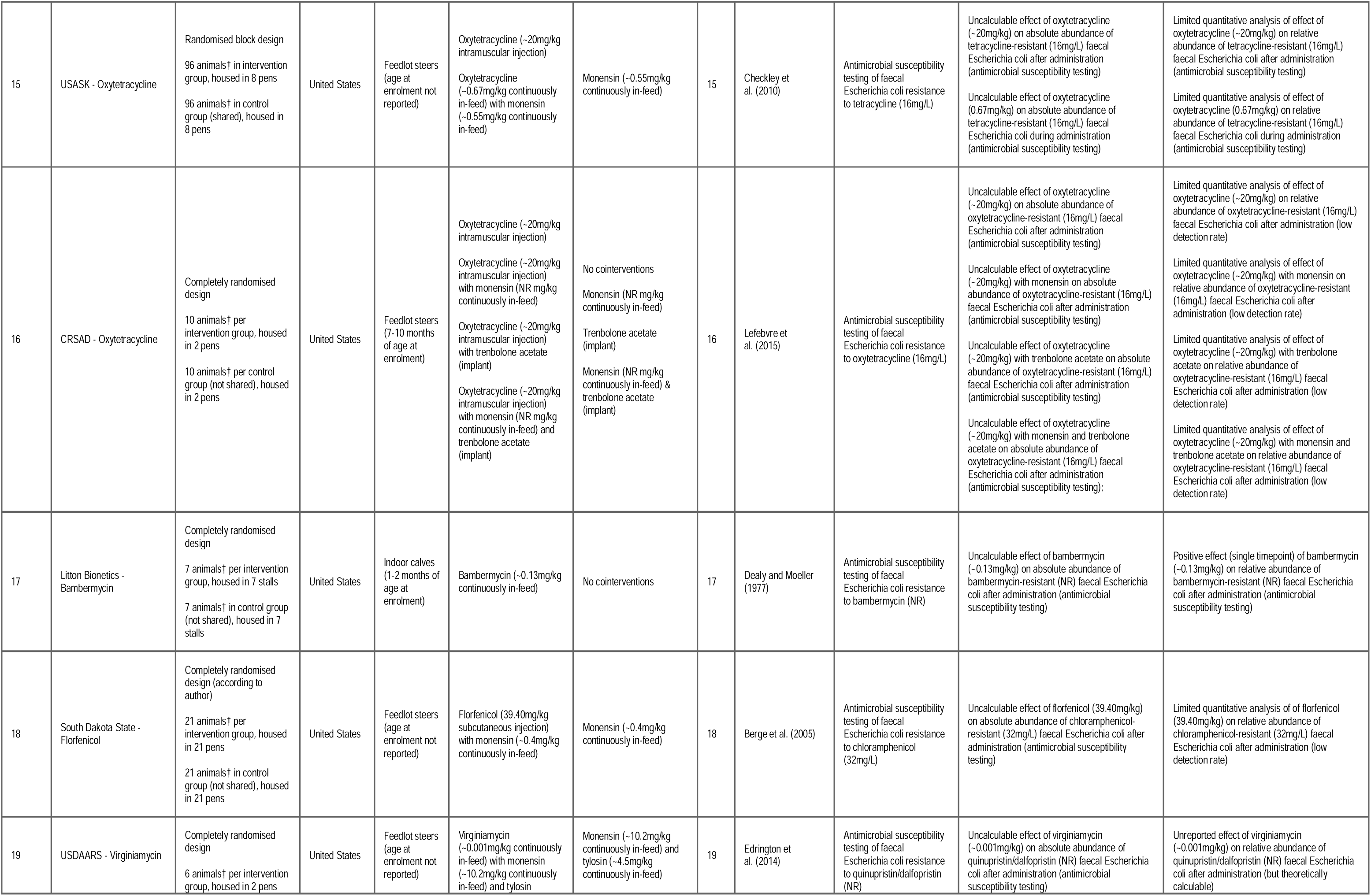
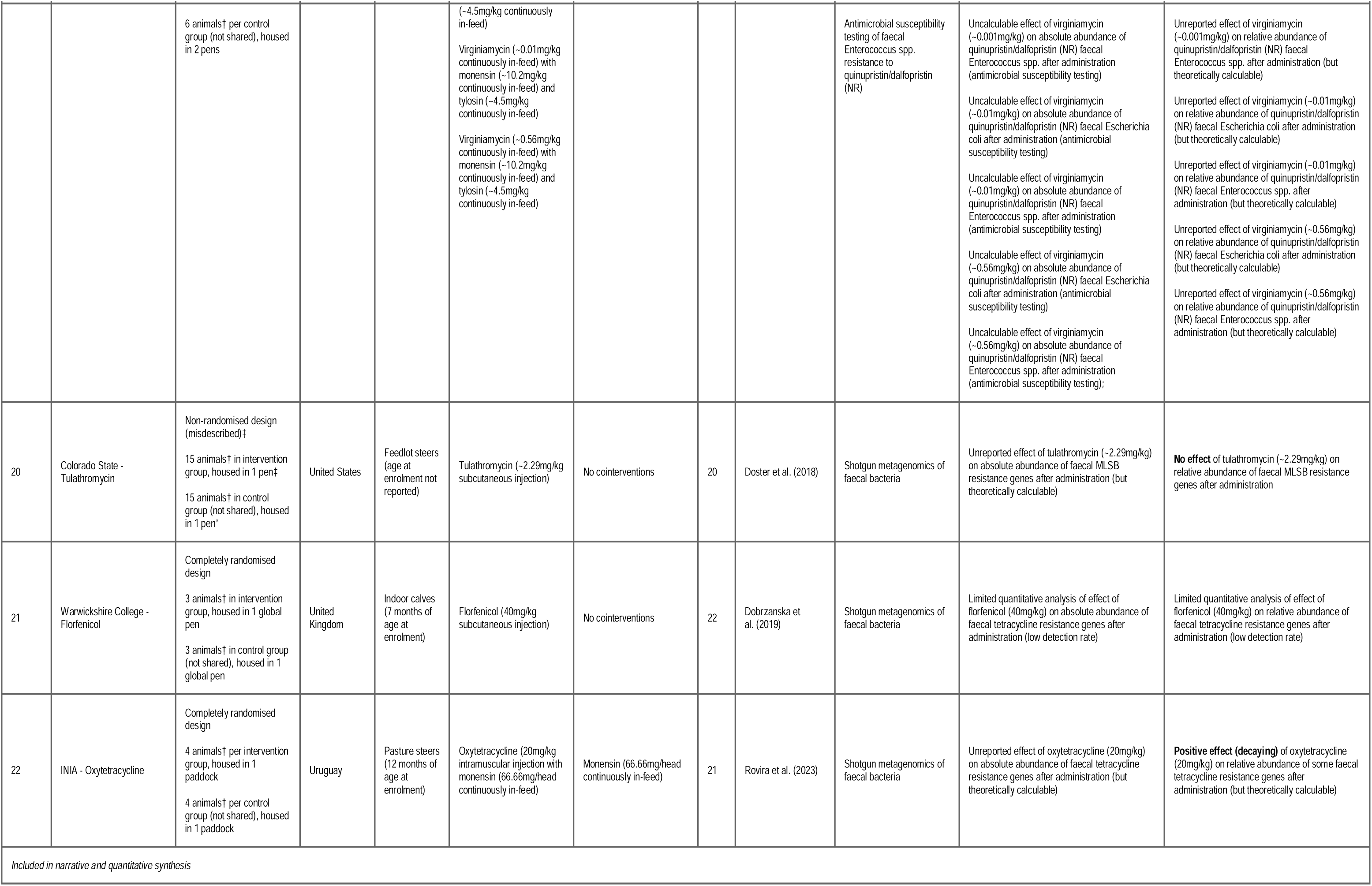
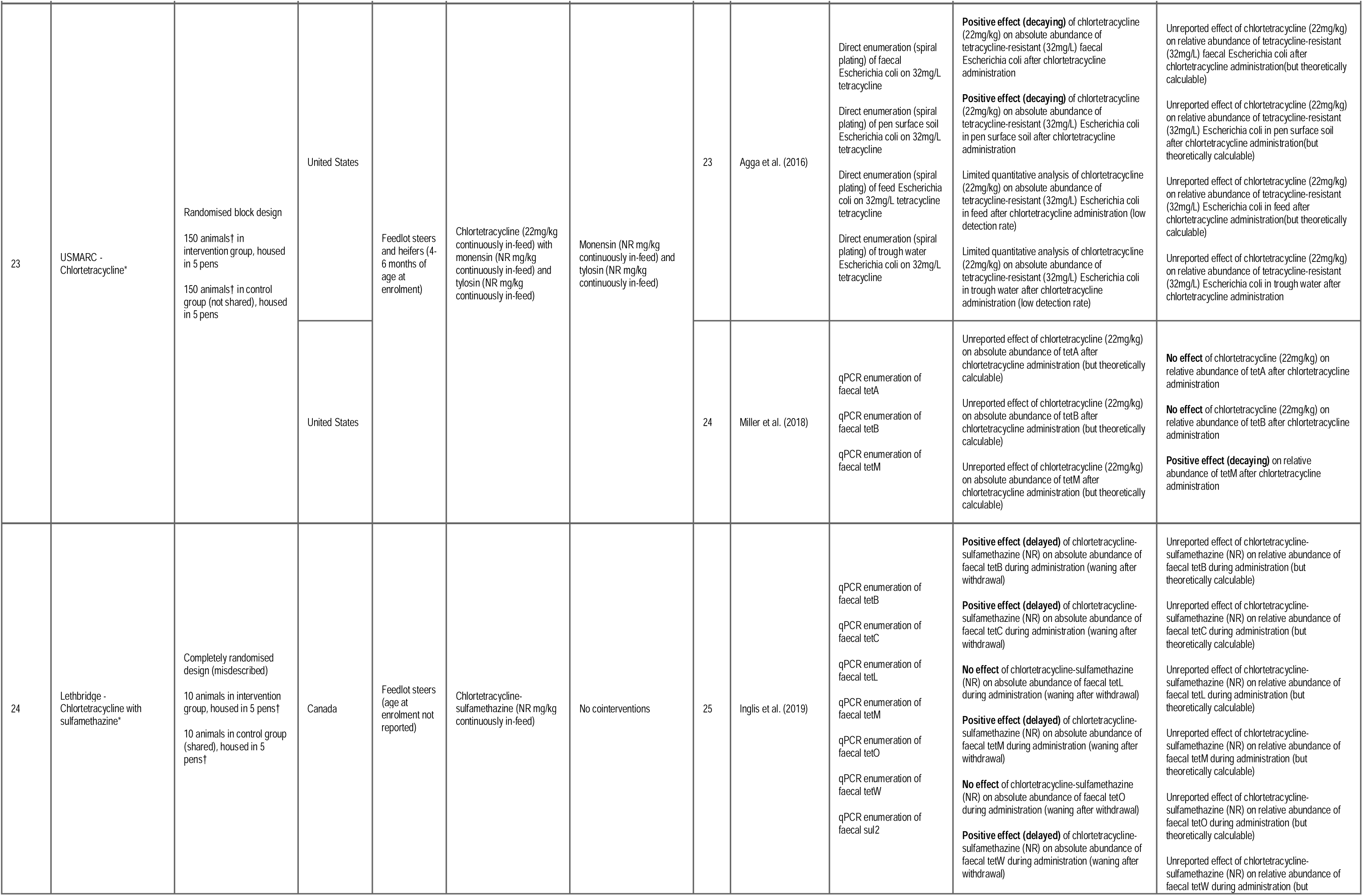
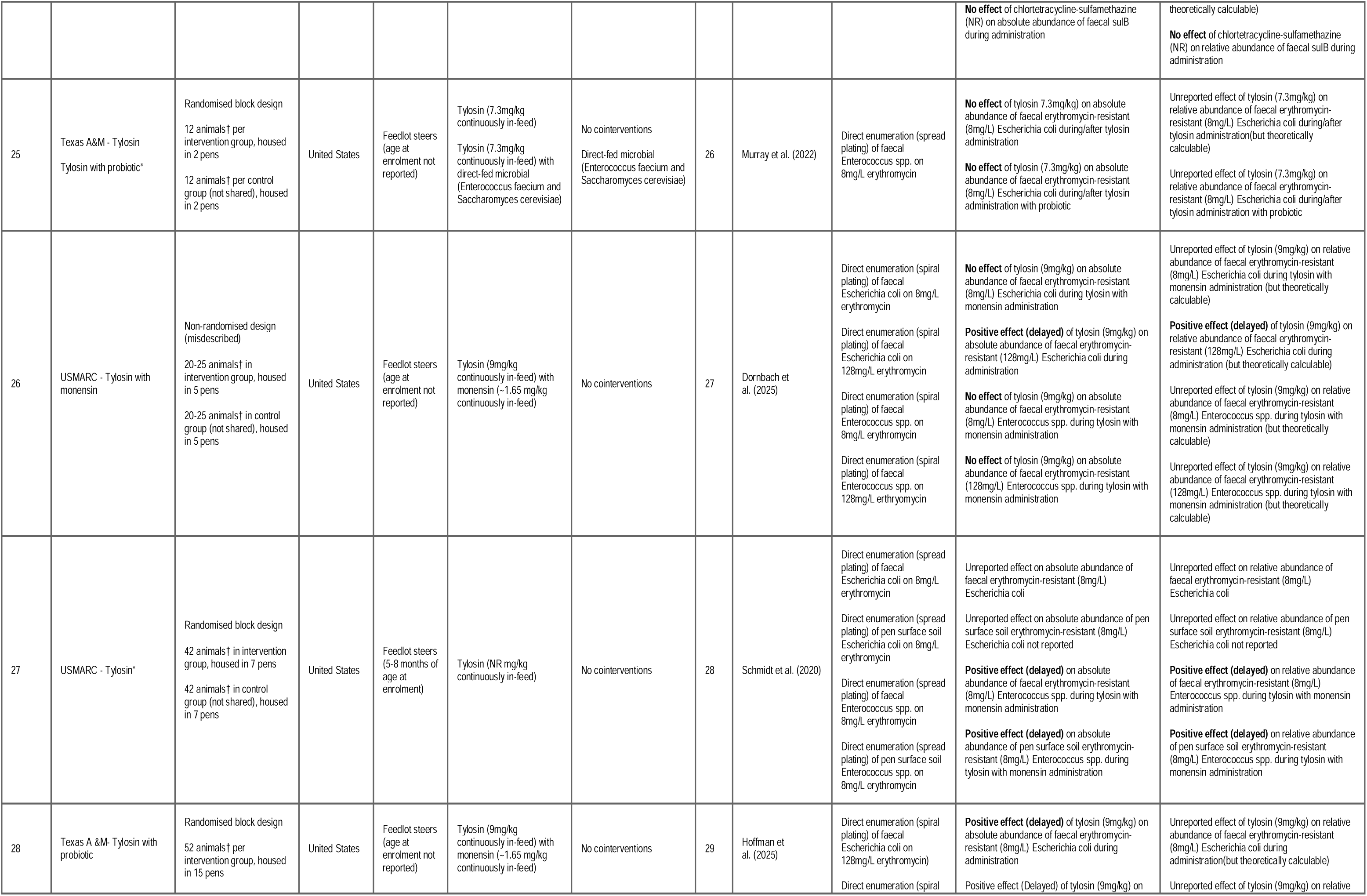
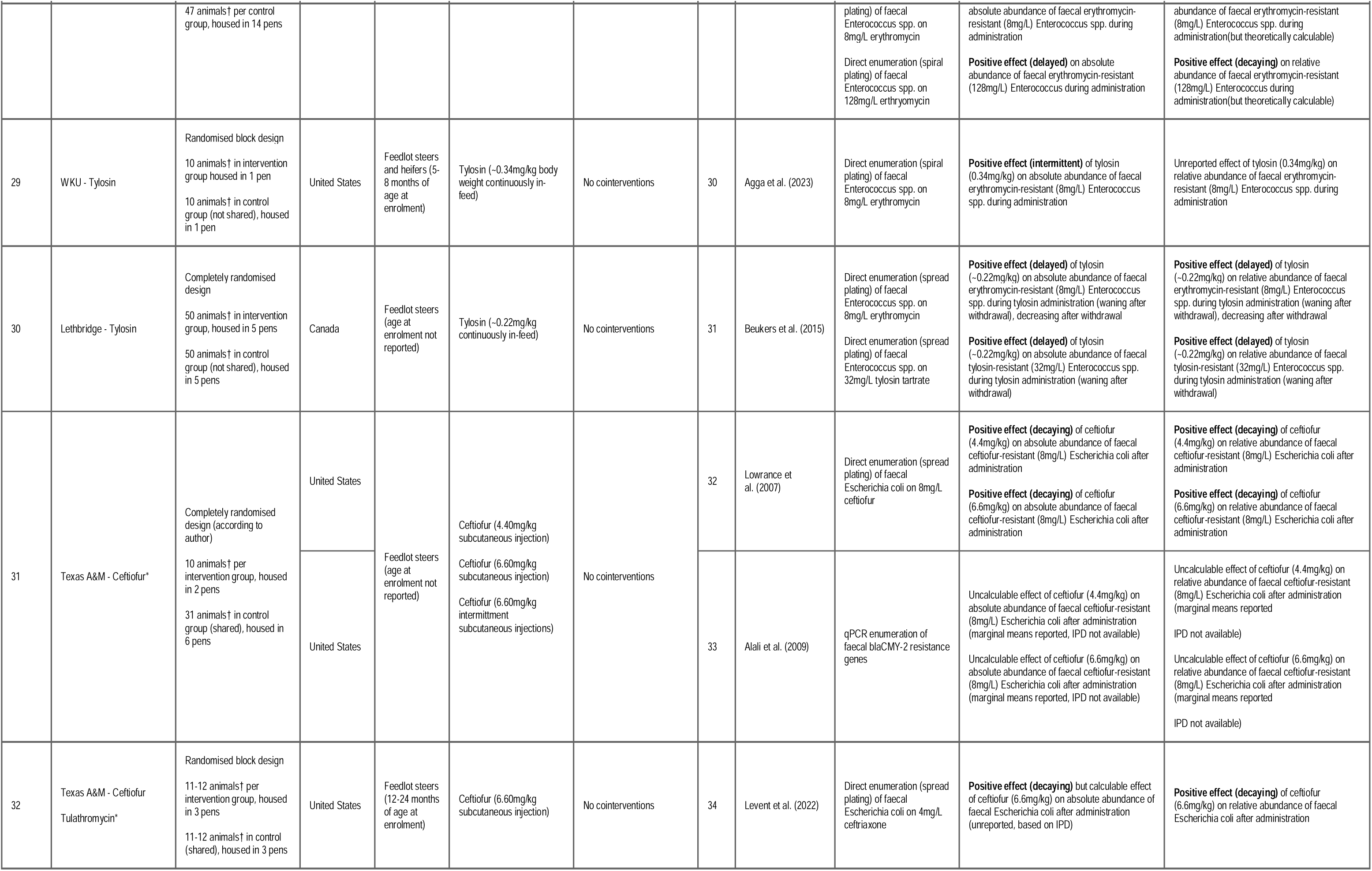
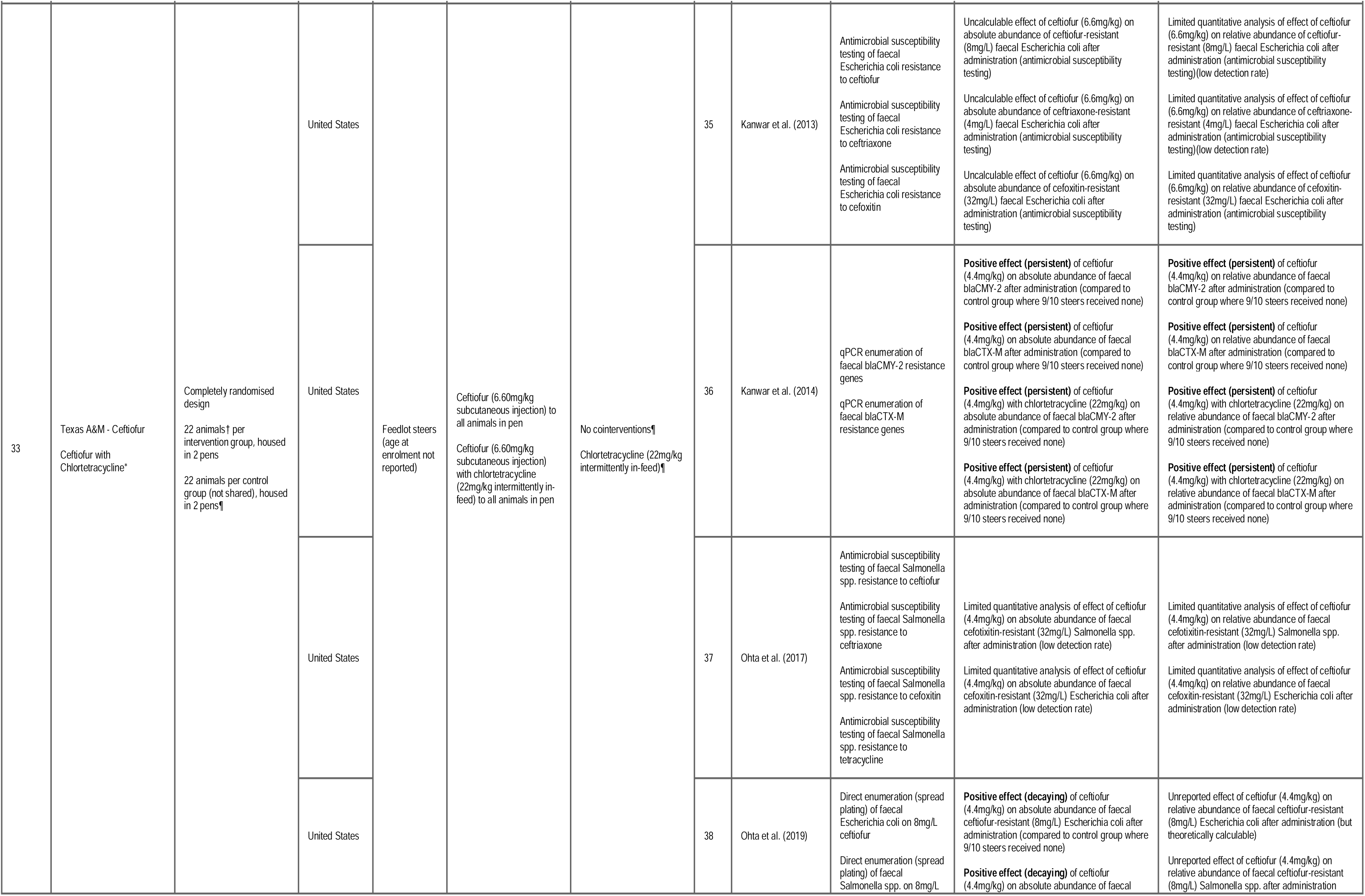
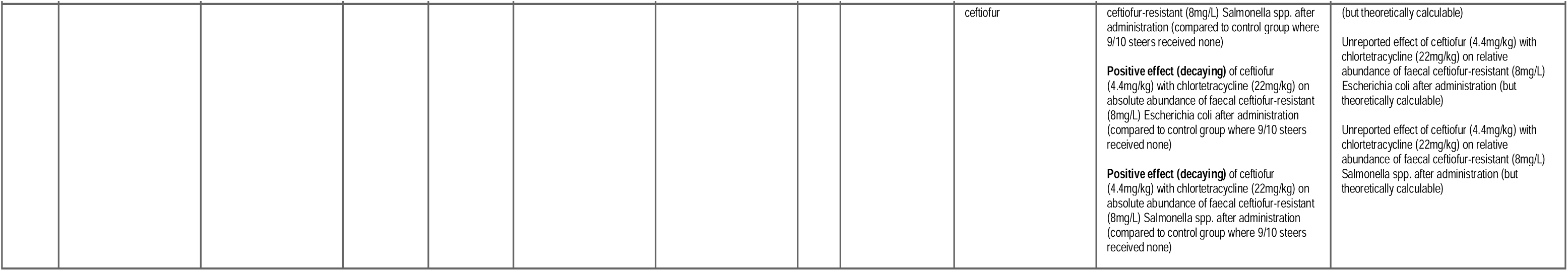
Study characteristics of the 33 included studies including study design, population, intervention(s), control(s)/comparator(s), and outcome(s). These studies occasionally reported multiple alternative outcomes from the same study (e.g. culture-dependent versus culture-independent measures of resistance) which were typically reported in separate publications. Hence there are 39 publications corresponding to the 33 studies, which are tabulated alongside the outcomes on which they reported within a study; *Data for this effect were included in the meta-analysis (reasons for excluding studies and effects in Supplementary Information 11 & 12, respectively); † unit of analysis as reported in publication; ‡ animals each group were a subset of the total number of cattle in each group; ¶ 1/11 animals in the control group were treated with ceftiofur (6.60mg/kg subcutaneous injection) and analysed as such in the original papers. For meta-analysis, we processed the individual participant data so as to remove this animal, thus ’decontaminating’ the control group and bringing it into line with other studies’ control groups.

### 3.3 Risk of bias in studies

We assessed risk of bias for the 33 included studies using the information provided in the publications in which they were reported (Supplementary Information 13 - 15; Supplementary Figure 6). Among the 26 randomised controlled trials, lack of reported blinding of those delivering the interventions and/or analysing study data was a key issue across all studies — leading to a risk that the randomisation process had been undermined (Domain 1) and deviations from the intended interventions had been made (Domain 2). The other key issue was the universal lack of pre-registration of studies, which led to a risk of selection of the reported results (Domain 5). For several studies, the use of antimicrobial susceptibility testing on a small number of isolates was also deemed an imprecise and inaccurate method of measuring the (relative) abundance of resistance (Domain 4), and hence these outcomes were excluded from the quantitative synthesis (see Methods: Synthesis methods: Selection process). Among the 7 studies with non-randomised designs, the key issues were that disease and treatment status were confounded (i.e. only sick animals were treated) and there was time-varying confounding (animals were recruited into treatment groups at different times).

Although not explicitly assessed as part of the risk of bias tools used, we also encountered unit-of-analysis issues in some of the studies when assessing risk of bias. In particular, some randomised block designs did not account for blocks in their analysis. Less commonly, several studies did not account for pen effects when housing animals across multiple pens. These issues further added to risk of bias in the results as reported in the literature — though for the subset of studies included in the meta-analysis, we corrected these issues where possible (see below).

### 3.4 Results of individual studies

#### 3.4.1 As reported in publications

Although ostensibly measured in the included studies, eligible effects were rarely reported in sufficient detail in publications establish their direction (Table 1). Often, this was because the low detection rate of resistance determinants (particularly for antimicrobial susceptibility testing) had resulted in a limited quantitative synthesis in the final publication (i.e. the effect could not be estimated). For other publications, it was less clear why the effects were not reported even though apparently measured.

Where effects were reported, publications reported either no effect or a positive effect of antibiotic administration on the absolute abundance of associated resistant bacteria/resistance genes in faeces (Table 1). Negative effects (whereby antibiotic interventions decrease the abundance of resistance determinants relative the control) were never reported. Where a positive effect was reported, it was often reported to be time-dependent (e.g. a delayed positive effect during antibiotic administration, decaying or persistent after antibiotic administration). This time-dependency appeared to be related to certain antibiotic interventions e.g. delayed effects when feeding tylosin^50,58^, decaying effects after injection of tetracyclines^57,60^, more persistent the effects after injection of MLSBs/cephalosporins^62,81^. For relative abundance outcomes, no effect or a positive effect was reported. The two metagenomic studies that reported effects only reported them in relative terms, reporting no effect and a positive effect on the relative abundance of relevant resistance genes.

Two studies assessed the effect in faeces-impacted environmental samples (e.g. pen surface soil, feed, trough water) as well as fresh faeces^45,77^. Only pen surface soil was assessed in both studies, in which the effect was reported to mirror that in faeces (i.e. resistance levels increasing and decreasing at similar times in both sample types).

#### 3.4.2 As calculated from available data (initial quantitative assessment)

11 of the 33 studies had data that was available and eligible for meta-analysis (Supplementary Information 11; Supplementary Information 12). These 11 studies were reported across 12 reports^44–46,50,60,62,64,67,69,70,73,77^.

In faeces, there was either no effect or a positive effect of antibiotic administration on the absolute abundance of resistance at the study-level (i.e. aggregating multiple effects from the same study) for each of the 11 studies. The overall effect across studies was a small-to-medium positive one (SMDH = 0.47; 95% CI 0.3 to 0.63, p = <0.01, Figure 2), which did not exist pre-antibiotic administration (SMDH = 0.06; 95% CI -0.18 to 0.31, p = 0.6; Supplementary Figure 7). When using SMDHs based on relative rather than absolute abundance, the results were very similar (Supplementary Figures 8 - 11).

**Figure 2:**
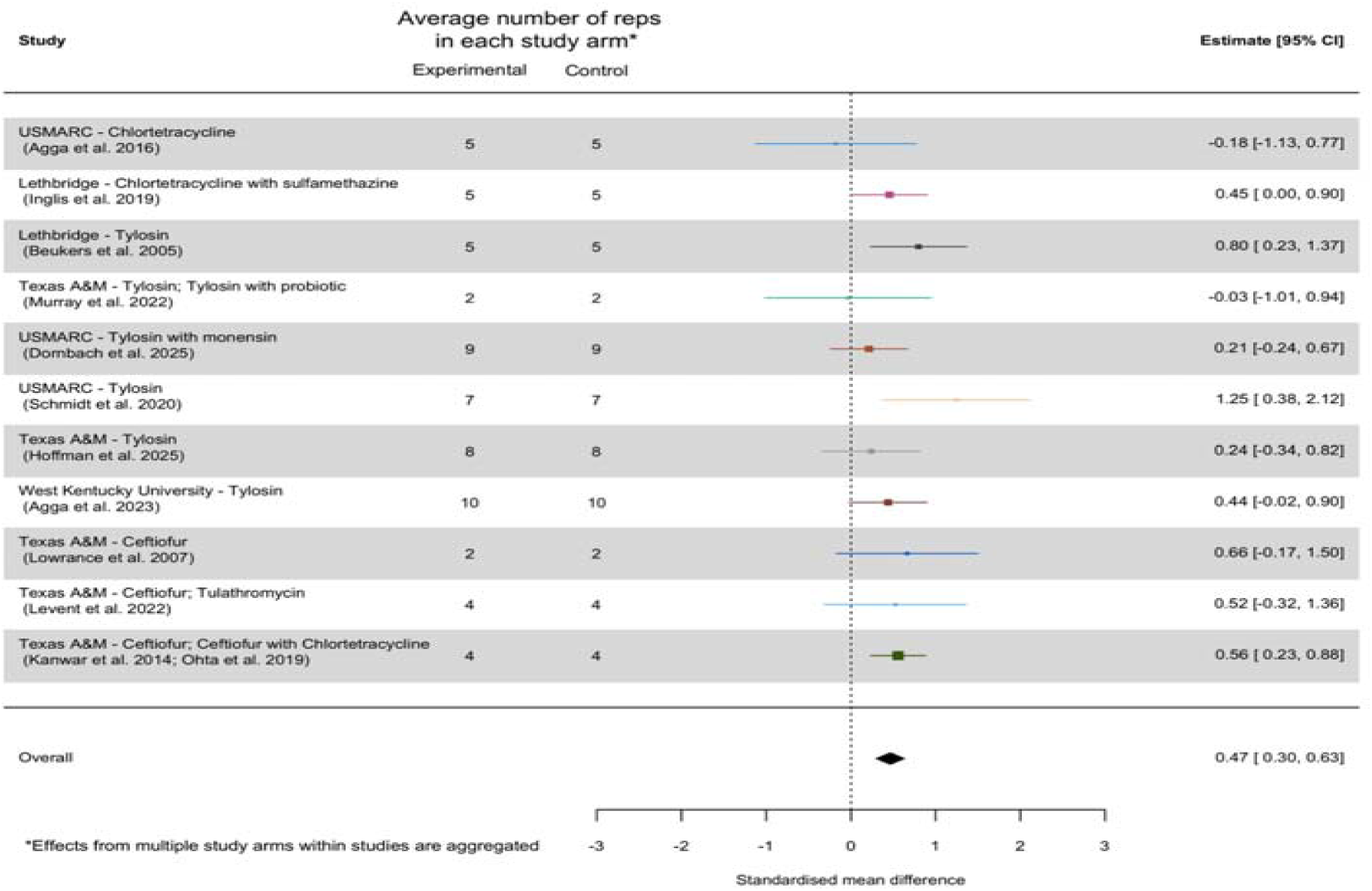
Forest plot of effects on the absolute abundance of antibiotic resistant bacteria in faeces post-antibiotic administration. Effects were aggregated at the study level using an intercept-only random effects model accounting for non-independence between effects from the same study, between temporally-autocorrelated effects from the same study arm, and residual variance as random effects. These were then pooled into an overall effect using an equal-effects model sensu Viechtbauer (2024b).

In the 2 studies of pen surface soil, the direction of the effect of antibiotic administration on the absolute abundance of resistance was positive at the study level and overall (Supplementary Figure 12), which was not the case before antibiotic administration (Supplementary Figure 13). However, no overall effect was detected (SMDH = 0.55; 95% CI -0.53 to 1.62, p = 0.3). When using SMDHs based on the relative abundance of resistance, we detected small-to-large positive effects in pen surface soil at the study-level and overall (Supplementary Figures 14 - 17).

### 3.5 Results of syntheses (hypotheses 1 and 2)

#### 3.5.1.1 Effects measured during antibiotic administration

##### 3.5.1.1.1 Primary analysis

7 of the 11 studies included in the quantitative synthesis measured effects during the administration of antibiotics. 6 of these measured the effect during tylosin administration, and 1 during chlortetracycline-sulfamethazine administration. 6 of these studies assessed the effect on antibiotic resistance bacteria (via direct plating), and 1 on antibiotic resistance genes (via qPCR). All but one of these studies^54^ included in the meta-analysis were randomised, though there was a high risk of bias for all studies.

Across effects measured during the administration of antibiotics, there was a small-to-medium positive overall effect of antibiotic administration on the absolute abundance of resistance (SMDH = 0.4; 95% CI 0.11 to 0.69, 95% PI -0.52 to 1.32, p = <0.01; Figure 3A), with considerable heterogeneity (I^2^ = 82.25%, Q_e(df=107)_ = 145.5, p = <0.01). When log10-transformed timing of sampling was added to the model as a linear moderator, there was a significant positive relationship between days of antibiotic administration and effect size (Q_m(df=1)_ = 5.36, p = 0.02; Intercept = -0.78, p = 0.1; Slope = 0.63, p = 0.02; Figure 3B), with estimated heterogeneity reduced (I^2^ = 61.05%, Q_e(df=106)_ = 134.75, p = 0.03). Profile likelihood analyses indicated that the random effects for the two models were well estimated (Supplementary Information 16; Supplementary Figures 18 - 19). A more complicated meta-regression model fitting time as a restricted cubic spline with 3 knots did not detect a non-linear effect of time (Q_m(df=2)_ = 5.18, p = 0.07; Intercept and Slope not relevant because model is non-linear; Supplementary Figure 20B).

**Figure 3:**
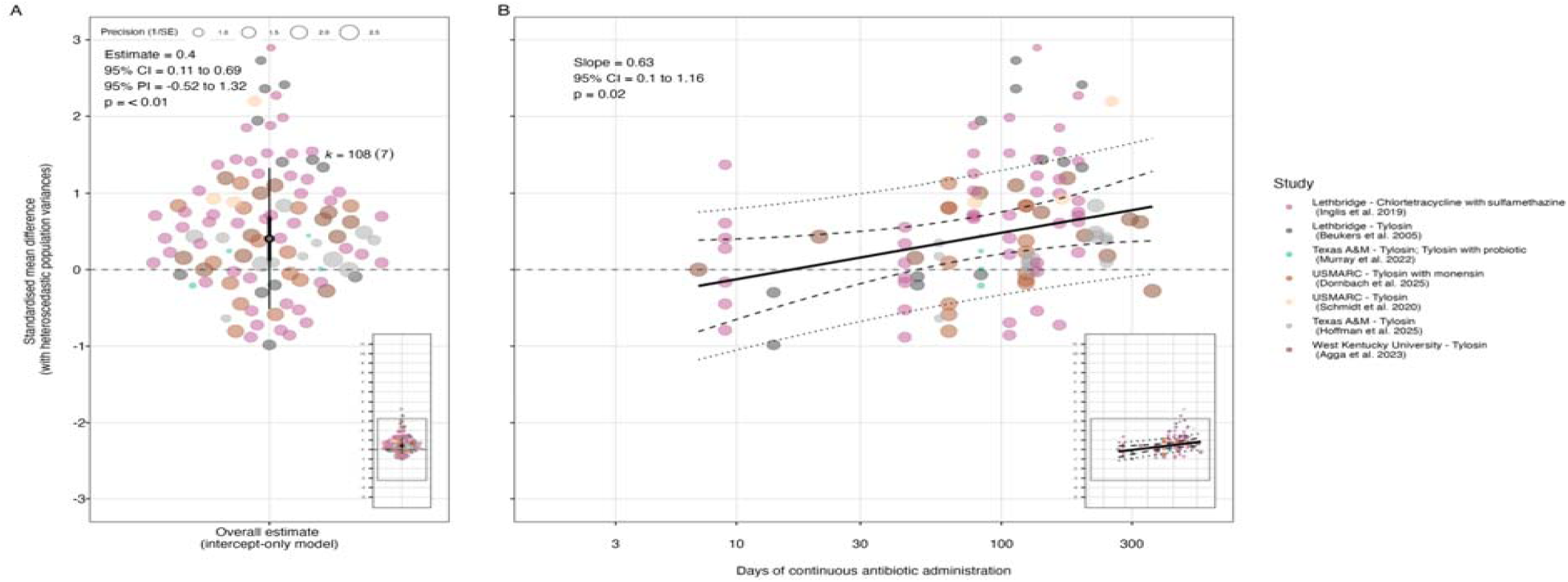
Plots representing results of meta-analytical models testing our two hypotheses for effects measured during the administration of antibiotics. In these plots, colours of the points represent the studies from which they are derived, and their size is proportional to their inverse standard error (1/SE; small points are less precise estimates, larger points are more precise estimates). A) Orchard plot representing the modelled overall effect of antibiotic administration on the absolute abundance of resistant bacteria/resistance genes relating to the intervention antibiotic class in the targeted bacterial population. This overall effect alongside its heterogeneity estimate, are derived from the intercept-only model. In this plot, the overall effect (model intercept after accounting for random effects) is represented by the ‘trunk’ of the tree (black circle), with its confidence interval represented by its ‘branch’ (shorter, thicker line through it), and its prediction interval represented by its ‘twig’ (longer, thinner line through it). B) Bubble plot representing the modelled relationship between timing of sampling (days of continuous antibiotic administration) and effect size (SMDH) from the meta-regression model. In this plot, the modelled effect of time is represented by the bold line, with the dashed line and dotted lines representing the 95% confidence and prediction intervals, respectively.

Effects of antibiotic administration on the absolute abundance of all determinants (susceptible and resistant; the denominator in relative abundance measures) provided further context regarding the nature of selection during antibiotic administration. In contrast to the effect on resistance determinants only, there was a small-to-medium negative overall effect on the absolute abundance of all determinants (SMDH = -0.37; 95% CI -0.59 to -0.15, 95% PI -1.14 to 0.4, p = <0.01; Supplementary Figure 21A). This indicated that there was a consistent suppression of the susceptible portion of the targeted bacterial population during antibiotic administration, likely contributing to the enrichment of the resistant portion. However, there was no detectable effect of either linear timing of sampling (Q_m(df=1)_ = 2.23, p = 0.1; Intercept = -1.08, p = 0.03; Slope = 0.37, p = 0.1; Supplementary Figure 21B) or non-linear timing of sampling (Q_m(df=2)_ = 2.21, p = 0.3; Intercept and Slope not relevant because model is non-linear; Supplementary Figure 22B).

##### 3.5.1.1.2 Sensitivity testing

Both the overall effect on the absolute abundance of resistance determinants during antibiotic administration (hypothesis 1) and its moderation by timing of sampling (hypothesis 2) were relatively robust to sensitivity analyses focused on removing or correcting studies/effect sizes. A statistically-significant small-to-medium overall positive effect (hypothesis 1) persisted regardless of which study was removed from the meta-analysis in the leave-one-out analysis (Supplementary Figure 23), and when removing groups of potentially problematic studies, except when removing 3 studies for which were not able to account for blocking (Supplementary Figure 24). The moderation of this effect by time (hypothesis 2) was more sensitive — becoming not statistically significant when removing two of the seven studies individually (Supplementary Figure 25), and when removing studies for which we had to extract data from published figures and when removing studies for which we were not able to account for pen-level effects (Supplementary Figure 26).

The overall effect and its moderation by timing of sampling were more sensitive to assumed correlation between effects from the same study; as we increased assumed correlation, the point estimate decreased and confidence and prediction intervals for both fits widened sharply (Supplementary Figures 27 & 28). However, this was to be expected given our analysis was based on a relatively small number of studies which often contained multiple arms and/or outcomes. Furthermore, whilst the overall effect quickly became non-statistically significant (even when fixing correlation at 0.2; Supplementary Figure 27), the time effect remained significant regardless of the amount of assumed correlation (suggesting it accounted for much of the correlation between effect sizes from different studies; Supplementary Figure 28).

##### 3.5.1.2 Effects measured after antibiotic administration

###### 3.5.1.2.1 Primary analysis

7 of the 11 studies included in the quantitative synthesis measured effects after the administration of antibiotics. 3 of these measured the effect after ceftiofur administration, 2 after tylosin administration, and 2 after chlortetracycline/chlortetracycline-sulfamethazine administration. 5 of these studies assessed the effect on antibiotic resistance bacteria (via direct plating), 1 on antibiotic resistance genes (via qPCR), and 1 on both. All interventions included in this meta-analysis were randomised, though there was high risk of bias for all studies.

Across effects after the cessation of antibiotic administration, there was a small-to-medium positive overall effect of antibiotic administration on the absolute abundance of resistance determinants which was similar in size to the effect during antibiotic administration (SMDH = 0.52; 95% CI 0.33 to 0.71, 95% PI 0.33 to 0.71, p = <0.01; Figure 4A). No significant heterogeneity between studies was identified in this model even before accounting for potential moderators (pseudo I^2^ = 0%, Q_e(df=86)_ = 48.3, p = 1). When we added log10-transformed timing of sampling to the model as a moderator, we detected a significant linear effect of time (Q_m(df=1)_ = 4.63, p = 0.03; Intercept = 1.32, p = <0.01; Slope = -0.65, p = 0.03; Figure 4B). Again, no significant heterogeneity was detected (I^2^ = 0%, Q_e(df=85)_ = 43.68, p = 1). Profile likelihood analyses indicated that the random effects for the two models were well estimated (Supplementary Information 16; Supplementary Figures 29 - 30). We also did not detect a significant non-linear effect of timing of sampling fitted as a restricted cubic spline with 3 knots in the meta-regression (Q_m(df=2)_ = 5.95, p = 0.05; Intercept and Slope not relevant because model is non-linear; Supplementary Figure 31B).

**Figure 4:**
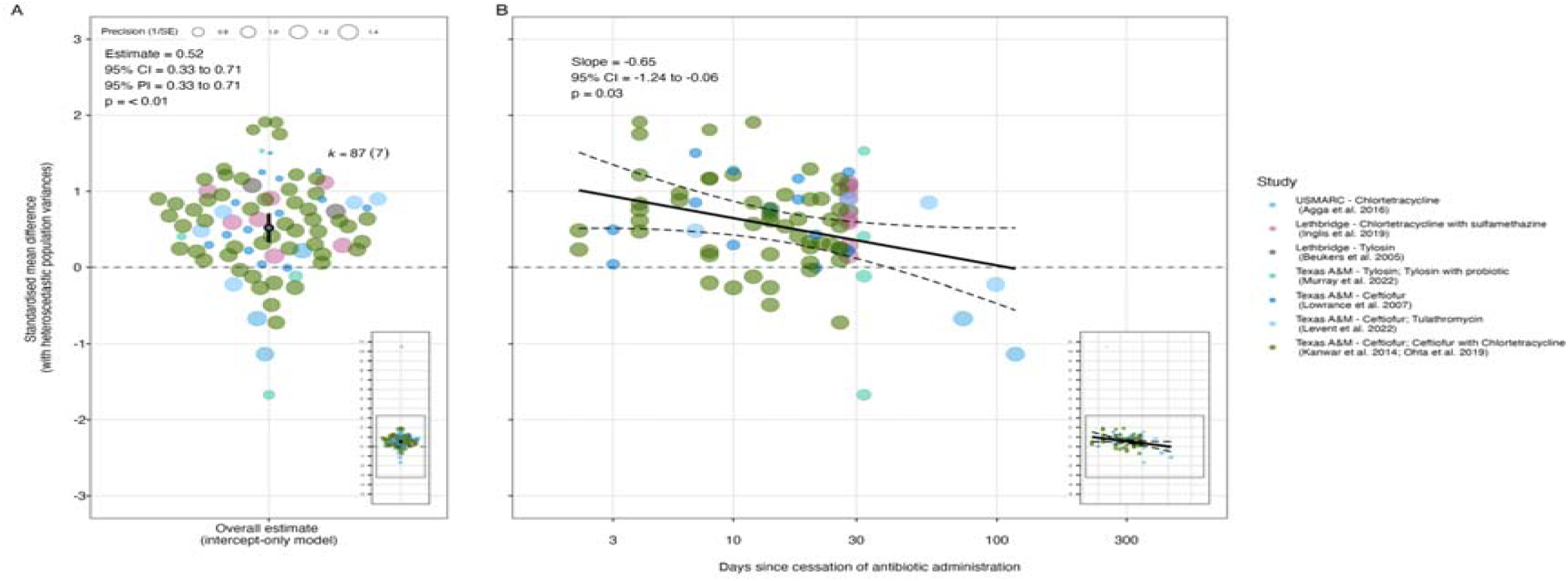
Plots representing results of meta-analytical models testing our two hypotheses for effects measured after the administration of antibiotics. In these plots, colours of the points represent the studies from which they are derived, and their size is proportional to their inverse standard error (1/SE; small points are less precise estimates, larger points are more precise estimates). A) Orchard plot representing the modelled overall effect of antibiotic administration on the absolute abundance of resistant bacteria/resistance genes relating to the intervention antibiotic class in the targeted bacterial population. This overall effect alongside its heterogeneity estimate, are derived from the intercept-only model. In this plot, the overall effect (model intercept after accounting for random effects) is represented by the ‘trunk’ of the tree (black circle), with its confidence interval represented by its ‘branch’ (shorter, thicker line through it), and its prediction interval represented by its ‘twig’ (longer, thinner line through it - not visible here as is identical to the confidence interval when τ2=0; Viechtbauer, 2025). B) Bubble plot representing the modelled relationship between timing of sampling (days of continuous antibiotic administration) and effect size (SMDH) from the meta-regression model. In this plot, the modelled effect of time is represented by the bold line, with the dashed line and dotted lines representing the 95% confidence and prediction intervals, respectively.

The positive overall effect on resistance determinants after antibiotic administration was mirrored by a small-to-medium negative overall effect on all determinants, though this was not significantly different from no effect (SMDH = -0.3; 95% CI -0.6 to 0, 95% PI -1.19 to 0.59, p = 0.05; Supplementary Figure 32A). There were no significant effects of either linear timing of sampling (Q_m(df=1)_ = 1.85, p = 0.2; Intercept = -1.01, p = 0.06; Slope = 0.55, p = 0.2; Supplementary Figure 32B), or non-linear timing of sampling (Q_m(df=2)_ = 3.79, p = 0.2; Intercept and Slope not relevant because model is non-linear; Supplementary Figure 33B).

###### 3.5.1.2.2 Sensitivity testing

Our estimates of the effect after antibiotic administration (hypothesis 1) and its moderation by timing of sampling (hypothesis 2) were relatively robust to sensitivity analyses focused on removing or correcting studies/effect sizes. The statistically-significant small-to-medium overall positive effect (hypothesis 1) persisted regardless of which study was removed from the meta-analysis in the leave-one-out analysis (Supplementary Figure 34), and only became non-significant when removing the 3 studies for which we had to extract data from published figures (Supplementary Figure 35). The moderation of this effect by time (hypothesis 2) was similarly robust, only losing its significance when removing the ‘USMARC - Chlortetracycline’ study in the leave-one-out analysis (Supplementary Figure 36). Removing this study (which was also the only one for which we were not able to account for blocking) also had the strongest effect in other sensitivity analyses (Supplementary Figure 37).

The overall effect after antibiotic administration and its moderation by timing of sampling were more sensitive to assuming increasing amounts of correlation between effect sizes from the same study, with confidence and prediction intervals for both fits widening sharply with increasing assumed correlation. Accordingly, the overall effect again became not statistically significant by an assumed correlation of 0.2 (Supplementary Figure 38), and again the time effect remained significant regardless of the amount of assumed correlation (Supplementary Figure 39).

##### 3.5.1.3 Subgroup and other alternative analyses

Subgroup and other alternative analyses were conducted on the full dataset of 195 post-antibiotic administration longitudinal measurements from all 11 studies eligible for quantitative synthesis, as an alternative to splitting the dataset into ‘during’ and ‘after’ periods and testing for effects of timing of sampling.

Across many subgroup analyses of various PICO(S) characteristics of studies (Supplementary Information 18; Supplementary Figures 40 - 52), characteristics of the intervention stood out as the clearest alternative explanation to timing of sampling for differences in effect sizes. In particular, the class of the antibiotic administered in the intervention detected a significant positive effect for the two most well-studied classes of antibiotics (Figure 5). In macrolides, lincosamides and streptogramin B/MLSB (which made up the majority of measurements taken during antibiotic administration) the size of the effect sensu Cohen^82^ was small-to-medium (Estimate = 0.38; 95% CI 0.12 to 0.65; 95% PI -0.33 to 1.1; p = <0.01). In extended-spectrum cephalosporins (which made up the majority of measurements taken after antibiotic administration) the size of the effect was medium (Estimate = 0.57; 95% CI 0.27 to 0.87; 95% PI -0.16 to 1.29; p = <0.01). Tetracyclines and sulfonamides were represented by two studies and one study, respectively, and no significant effect was detected in either class (Figure 5). Subgroup analysis by mode of administration produced similar results, given mode of administration was confounded by class of antibiotic administered (Supplementary Figure 40).

**Figure 5:**
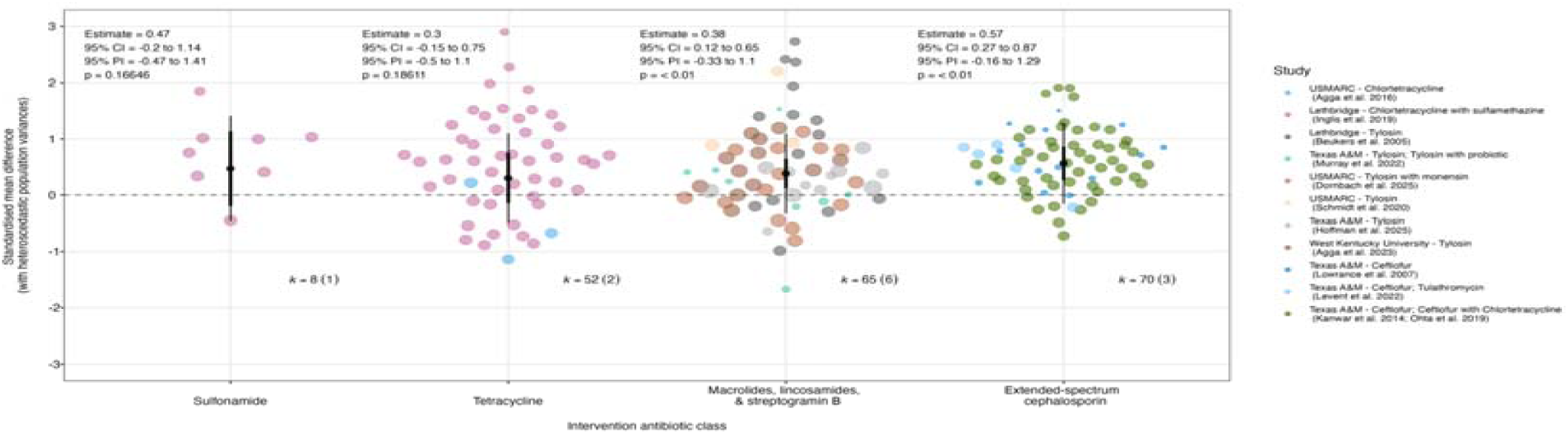
Orchard plot representing results of the meta-analytical model testing for a subgroup effect of the class of antibiotic administered in the intervention. Points are effect sizes, with their colours representing the studies from which they are derived and their size being proportional to their inverse standard error (1/SE; small points are less precise estimates, larger points are more precise estimates). The pooled effect for each subgroup (coefficient after accounting for random effects) is represented by the ‘trunk’ of the tree (black circle), with its confidence interval represented by its ‘branch’ (shorter, thicker line through it), and its prediction interval represented by its ‘twig’ (longer, thinner line through it). For the numbers above each subgroup, ‘k’ represents the number of effect sizes, whilst the number in brackets represents the number of studies from which they were derived.

#### 3.5.2 Reporting bias

We found minimal evidence of publication bias in effects measured during or after antibiotic administration. Exploratory funnel plots of the residuals of the hypothesis 1 models (overall effect during and after) and selected models for hypothesis 2 (linear time effect during and non-linear time effect after) showed no clear relationship (Supplementary Figures 53 & 54). Formal tests found no relationship between the square root of the inverse of the effective sample size and effect size (small study publication bias) during (Intercept = 0.24, Slope = 0.29; p = 0.8) and marginal evidence of a positive relationship between effect size and sample size antibiotic administration (Intercept = 0.39; Slope = 0.18; p = 0.9). We did not detect any relationship between mean-centered publication year (time-lag publication bias) and effect size during (Intercept = 0.61, Slope = -0.06; p = 0.1) or after (Intercept = 0.61, Slope = -0.06; p = 0.1) antibiotic administration.

#### 3.5.3 Certainty of evidence

Whilst evidence consistently pointed to the presence of a positive effect of antibiotic administration on antibiotic resistance, we judged there to be ‘low’ certainty in the exact size of this effect following a GRADE approach (Table 2; Supplementary Table 4).

**Table 2:** Summary of Findings: Summary of findings of the meta-analytical results for the review outcomes — the absolute abundance of resistance determinants for a particular antibiotic class during and after its administration, with and without accounting for the moderating effect of time. Briefly, the population was weaned beef cattle, the intervention was antibiotic administration, and the comparator was no administration of the intervention antibiotic. Illustrative comparative differences are derived from similar effect sizes (in a similar time window). Certainty of evidence was assessed using the GRADE approach (Schünemann et al., 2013; Schünemann et al., 2023)

| Outcome | Illustrative comparative difference |  |  | Relative effect (95% CI) | Number of studies | Certainty of the evidence (GRADE) in the size of the effect |
| --- | --- | --- | --- | --- | --- | --- |
|  | Example | Without antibiotic | With antibiotic |  |  |  |
| Faecal density of resistance determinants for a particular antibiotic class during its administration to beef cattle | Abundance of erythromycin-resistant (128µg/ml) E.coli in faeces during in-feed tylosin administration (~75mg/head/day; ~0.28mg/kg body weight/day) in the intervention group (Hoffman et al. 2025; SMDH = 0.42) | 10<sup>2.04</sup> resistant bacteria per g of faeces at 242 days since start of antibiotic administration | 10<sup>2.16</sup> resistant bacteria per g of faeces at 242 days since start of antibiotic administration | SMDH = 0.4 (95% CI 0.11 to 0.69) | 7 (3 completely randomised; 3 randomised complete block; 1 non-randomised) | -----<br>**low** |
| Faecal density of resistance determinants for a particular antibiotic class during its administration to beef cattle, accounting for the moderating effect of time | Increase in abundance of tetM in faeces during in-feed chlortetracycline-sulfamethazine administration (350-350mg/head/day; ~1.5mg/kg body weight/day) in the intervention group (Ingilis et al. 2019; Slope of SMDH = 0.52) | Increase from 10<sup>4.77</sup> resistance genes per g of faeces at 9 days since start of antibiotic administration to intervention group, to 10<sup>6.71</sup> resistant bacteria per g of faeces at 191 days since start of antibiotic administration to intervention group | Increase from 10<sup>6.08</sup> resistance genes bacteria per g of faeces at 9 days since start of antibiotic administration to intervention group, to 10<sup>8.07</sup> resistant bacteria per g of faeces at 191 days since start of antibiotic administration to intervention group | Slope of SMDH = 0.63 (95% CI 0.1 to 1.16) | 7 (3 completely randomised; 3 randomised complete block; 1 non-randomised) | -----<br>**low** |
| Faecal density of resistance determinants for a particular antibiotic class after its administration to beef cattle | Abundance of ceftiofur-resistant (8µg/ml) E. coli in faeces after ceftiofur administration (4.4mg/kg body weight/day) in the intervention group (Lowrance et al. 2007; SMDH = 0.5) | 10<sup>3.56</sup> resistant bacteria per g of faeces 3 days after antibiotic administration | 10<sup>4.18</sup> resistant bacteria per g of faeces 3 days after antibiotic administration | SMDH = 0.52 (95% CI 0.33 to 0.71) | 7 (4 completely randomised; 3 randomised complete block) | -----<br>**low** |
| Faecal density of resistance determinants for a particular antibiotic class after its administration to beef cattle, accounting for the moderating effect of time | Decrease in abundance of ceftiofur-resistant (8µg/ml) E. coli in faeces between 4 and 26 days since end of ceftiofur administration (6.6mg/kg body weight/day) in the intervention group (Lowrance et al. 2007; Slope of SMDH = -0.61) | Increase from 10<sup>1.46</sup> resistant bacteria per g of faeces at 4 days since end of antibiotic administration to intervention group, to 10<sup>3.19</sup> resistant bacteria per g of faeces at 26 days since end of antibiotic administration to intervention group | Increase from 10<sup>2.98</sup> resistant bacteria per g of faeces at 4 days since end of antibiotic administration to intervention group, to 10<sup>3.74</sup> resistant bacteria per g of faeces at 26 days since end of antibiotic administration to intervention group | Slope of SMDH = -0.65 (95% CI -1.24 to -0.06) | 7 (4 completely randomised; 3 randomised complete block) | -----<br>**low** |

For outcomes measured during antibiotic administration, this judgement was arrived at by first downgrading for risk of bias (due to all 7 studies contributing to the overall effect having a high risk of bias). Subsequently, we downgraded for inconsistency since there was evidence of heterogeneity in the effect in I^2^ tests. We did not downgrade for indirectness since effects of the three major groups of antibiotics fed to cattle (MLSBs, tetracyclines, and sulfonamides) contributed to the pooled estimate of the effect during antibiotic administration.

For outcomes measured after antibiotic administration, this judgement was arrived at by first downgrading for risk of bias (due to all 7 studies contributing to the overall effect having a high risk of bias). We did not downgrade for heterogeneity since we did not detect heterogeneity either before or after accounting for the moderating effect of time. However, we did downgrade for indirectness since all estimates came from extended-spectrum cephalosporin interventions.

We did not upgrade any of our certainty judgements because none of the point estimates/slopes in our models were large according to interpretive guidelines (SMDH > 0.8), there was no clear evidence of straightforward dose-response effects in our tests for it, and there was no clear reason to believe all confounding would have overestimated the effect (plausible confounding).

## 4 Discussion

### 4.1 General interpretation of results

This systematic review and meta-analysis represents the most comprehensive synthesis to date assessing evidence for a relationship between antibiotic use and resistance in beef production. Our quantitative analysis in particular demonstrates that, overall, administering antibiotics to beef cattle increases the abundance of resistant phenotypes and genotypes associated with that antibiotic in their faeces. Moreover, this effect appears time-dependent — increasing in strength with days of continuous administration, and decreasing in strength with days since cessation of antibiotic administration. We are less certain about the exact size of these effects due to limitations of the evidence base included in the review (see below). However, our review clearly demonstrates the presence of these effects which is important given that the lack of a comprehensive synthesis has previously caused an overreliance on theory and even a questioning of the degree to which the use of antibiotics in animal food production results in the emergence of resistant bacteria^83^.

Demonstrating the presence of a positive overall effect of antibiotics in general (which we estimated to be small-to-medium in size; see below) represents a clear improvement over previous reviews. A previous WHO-commissioned systematic review indirectly implied such an effect existing by finding that broad interventions to restrict antibiotic use are associated with decreased relative abundance of resistant bacteria and genes^6,7^. A previous sys systematic review identified a positive effect of tylosin based on a semi-quantitative meta-analysis of four studies^8^. A more recent systematic review found non-significant associations in Salmonella but highlighted that the uncertainty of the evidence made their pooled estimates untrustworthy^9^. Overcoming some of the difficulties these syntheses encountered through a comprehensive synthesis and in-depth meta-analysis of individual participant data, we show clearly that a positive effect does exist in faeces. This is important because faeces are a primary vector through which antibiotic resistance emerging in beef production is likely to be transferred to humans and other animals through contamination of beef and the environment (though our synthesis was less able to make direct conclusions about effects in the environment).

Directly demonstrating the moderating effect of time also represents an improvement over previous reviews, which did not attempt such a meta-analysis. During antibiotic administration, we estimated that the effect size linearly increased from no effect at 10 days since antibiotic administration to a medium-to-large effect at 300 days of continuous antibiotic administration (Table 2). This is suggestive of a gradual increase in the abundance of resistance phenotypes and genotypes under long-term administration of subtherapeutic doses of antibiotics in feed for the purposes of prophylaxis and/or growth promotion (e.g. tylosin which constituted the majority of the interventions included in this part of the meta-analysis). After antibiotic administration, we estimated that the effect size linearly decayed from a medium-to-large effect at 10 days since antibiotic administration to no effect at 100 days since the cessation of antibiotic administration (Table 2). Such ‘resistance decay’ over similar timescales was previously observed in a systematic review and meta-analysis of human participants^84^, but ours is the first to observe it in livestock.

### 4.2 Limitations of the evidence included in the review

Whilst our review presents sufficient evidence to demonstrate the existence of time-dependent effects of antibiotic administration on antibiotic resistance, the exact size of these effects is less certain given issues in the underlying evidence base. Most obviously — despite being inclusive of different antibiotics in our review criteria — a limited number of studies of studies were eligible for inclusion in the review. A total of 33 studies were included in our systematic review, but we were not able to derive effects from either publications or upon request for most of these.

This meant that only 11 studies were able to be included in our meta-analysis of effects in faeces (only 2 of which had estimates of effects in faeces-impacted environments, precluding a meta-analysis of this outcome type). Aside from being related to potentially inappropriate methods of reporting the outcome (e.g. antimicrobial susceptibility testing; see below) and selective reporting, the limited number of studies available for meta-analysis brought its own issues. For example, it made our results more sensitive to leaving out certain studies and assuming a higher degree of correlation between multiple effect sizes from the same study.

There were also issues associated with the quality of included studies — with risk of bias, inconsistency, and indirectness being highlighted in the certainty assessment. On risk of bias, the almost universal (reported) blinding of investigators and the lack of pre-registration of studies introduced some risk that randomisation was undermined and effects were selectively reported, respectively. On inconsistency, significant heterogeneity was detected in effects measured during antibiotic administration, which may have reflected the diversity of interventions. On indirectness, effects measured after antibiotic administration were focused on ceftiofur, meaning the results were less generalisable to other antibiotics even as they were less heterogenous. There were also other quality issues such as the common use of antimicrobial susceptibility testing based on a very small number of isolates (which led to us excluding these outcomes from the meta-analysis), and the use of randomised complete block designs without recording blocking information (which meant we were not able to account for blocking when including some studies in the meta-analysis).

These limitations contributed to the conclusion of our certainty assessment that there was ‘low certainty’ in the exact size of the effects, which is somewhat consistent with a recent review^9^. However, we conclude that there is still sufficient evidence in the presence, if not the exact size and shape, of time-dependent effects. This is in view of the consistency with which narrative and quantitative synthesis point to positive effects on average, with biologically-plausible evidence that these effects may be larger in certain contexts (e.g. after many days of continuous antibiotic administration, immediately after cessation of antibiotic administration, or the administration of extended-spectrum cephalosporins). Moreover, issues such as the low quantity of studies, their small sample size, and not accounting for blocking would all be expected to widen confidence intervals and result in an *underestimated* effect^85^. We therefore conclude that despite limitations of the evidence included, there is sufficient evidence in at least the presence of an effect to warrant action — especially when considering the various practical, funding, institutional, and ethical reasons that limit the quantity and quality of studies available in this field.

### 4.3 Limitations of the review processes used

Our review represents the most comprehensive synthesis of its kind, and we addressed many of methodological difficulties either at the protocol design stage or during the process of conducting the review as we encountered them (with deviations fully reported and justified in Supplementary Information 2). Nonetheless, there are limitations in the final review processes used. For example, related to limitations of the evidence base, we included studies with different designs and quality issues (e.g. a high risk of bias). This likely contributed to increased heterogeneity, and lower certainty in the meta-analytical effects. Conversely, limiting our review to effects of direct selection (i.e. selection for resistance phenotypes and genotypes related only to the antibiotic class administered in the intervention) meant that we were not able to directly assesses effects on resistance in non-intervention antibiotic classes — even though they are likely correlated with effects on resistance to intervention antibiotics^86,87^. One notable manifestation of this is that we did not include monensin interventions in our synthesis^88^, since these studies typically focus on the effects of monensin on non-ionophore antibiotics which have clinical uses and breakpoints unlike ionophores^89^.

These points of potential under and overinclusiveness should be born in mind when interpreting our results, but should be regarded as limitations only. The heterogeneity introduced by including different study designs was ameliorated by the use of a standardised effect size (SMDH), and sensitivity and subgroup analyses that contextualise our results and alternative explanation for them. The lower certainty associated with including studies with a high risk of bias in our meta-analysis was less able to be ameliorated, but given all studies were judged to have a high risk of bias this approach was the only option and is at least consistent with that of previous reviews^6,8,9^. Aspects of potential underinclusiveness such as the focus on direct selection limit the generality of our conclusions, but were important to put some limits on this diverse evidence base to ensure the most causative evidence was synthesised (which was also why our eligibility criteria focused on controlled studies with baseline data). Therefore, our review methodology represents a balance between comprehensiveness with stringency; being inclusive in the face of a limited and varied evidence base, whilst prioritising the best available data for estimating the effect.

### 4.4 Implications of the results for practice, policy, and future research

A recent review^9^ argued that this evidence base was too uncertain to be able to inform policy and practice. However, our clear demonstration of the existence of time-dependent effects imply further policy action on the use of antibiotics in beef production is needed — especially as the evidence base we synthesised is most relevant to macrolides and extended-spectrum cephalosporins, which are classified as Highest Priority Critically Important Antimicrobials (HPCIA) and Critically Important Antimicrobials (CIA) by the World Health Organization^90^. For example, in the USA and Canada, the use of antibiotics such as tylosin for growth promotion is now ostensibly banned, but they can still be administered for extended periods for the purposes of preventing liver abscesses common in feedlot cattle^91^. Future North American legislators may therefore consider whether such legislation can be tightened, supported by antibiotic alternatives to treat and even prevent liver abscesses. In South America, the five largest meat producing countries (Argentina, Brazil, Chile, Colombia, and Uruguay) have tightened legislation in recent years, but this can still be improved by designating third- and fourth-generation cephalosporins as last resort antibiotics and preventing their use for prophylaxis, for example^92^. In contrast to the New World, the European Union has perhaps the most comprehensive restrictions in the world, banning the use of antibiotics for growth promotion, the routine use of antibiotics (e.g. to compensate for poor hygiene), and ostensibly the use of antibiotics for pro/metaphylaxis^93,94^. However, antibiotics can still be used pro/metaphylactically where the risk of infection or its spread is considered high^94^ — another potential point for legislative tightening. Short of banning antibiotics, policymakers might consider how our results about the moderating effect of time can be used to reduce the permitted duration of antibiotic feeding^69,95^, or antibiotic withdrawal periods currently based on the predicted decay of antibiotic concentration alone^96^.

At the same time, the low certainty in the exact size of the effect warrants attention from research teams, funders, and publishers to reduce research waste. Adopting and/or encouraging the more widespread use of blinding and pre-registration in these studies are obvious points for improvement that emerge from our risk of bias assessments. Similarly, our review highlights the need to reduce reliance on antimicrobial susceptibility testing and record/account for blocking factors when assessing effects in these studies. Finally, wider adoption of open research practices would improve transparency around unreported effects and facilitate the reuse of studies results in evidence synthesis and policy contexts. These include standardised reporting guidelines which are common in human trials and have previously been called for in this field too^8,97–100^, and openly publishing research data (and code) alongside their scientific papers at the point of publication.

### 4.5 Conclusion

Several controlled studies have been conducted over the past two decades to understand whether there is a cause-effect relationship between antibiotic use in the beef industry and antibiotic resistance. Our pre-registered systematic review and meta-analysis overcomes several of the limitations of previous syntheses to show that a causal and general relationship between antibiotic use in the beef industry and antibiotic resistance is likely. Moreover, this biologically plausible effect is likely to be moderated by time — increasing with the number of days of continuous antibiotic administration during antibiotic administration, and decreasing with days since the cessation of antibiotics after antibiotic administration. As in previous reviews, there remains some uncertainty in the exact size of these effects due to the risk of bias associated with these veterinary studies (at least when measured against the yardstick of medical studies). However, the evidence is sufficient to conclude that antibiotic use in the beef industry selects for antibiotic resistance and any future improvements to research practices in this field might therefore be better spent on studies that address solutions to this problem (e.g. antibiotic alternatives), rather than characterising it further.

## 5 CRediT authorship contribution statement

**Matt Lloyd Jones (guarantor)**: Conceptualization, Methodology, Software, Validation, Formal analysis, Investigation, Resources, Data Curation, Writing – Original Draft, Writing – Review & Editing, Visualization, Supervision, Project administration; **Alfredo Sánchez-Tójar:** Conceptualization, Methodology, Investigation, Software, Formal analysis, Resources, Writing - Review & Editing, Supervision, Validation, Project administration; **Alison Bethel:** Methodology, Investigation, Data Curation, Writing – Review & Editing; **Anne Frances Clare Leonard:** Conceptualization, Methodology, Investigation, Writing – Review & Editing, Supervision, Funding acquisition; **Emma Lamb:** Methodology, Investigation, Writing – Review & Editing; **Natalia Casanova:** Conceptualization, Methodology, Investigation, Writing – Review & Editing; **Johana Dominguez:** Conceptualization, Methodology, Investigation, Writing – Review & Editing; **María Paula Quiroga:** Conceptualization, Methodology, Investigation, Writing – Review & Editing; **Daniela Centron:** Conceptualization, Methodology, Investigation, Writing – Review & Editing, Funding acquisition; **Adriana Peralata Alonso:** Methodology, Investigation, Writing – Review & Editing; **Mariano Fernández-Miyakawa**: Conceptualization, Funding acquisition; **William H. Gaze:** Conceptualization, Methodology, Writing – Review & Editing, Supervision, Funding acquisition; **Alejandro Petroni:** Conceptualization, Methodology, Validation, Investigation, Writing – Review & Editing, Supervision, Funding acquisition; **Ruth Garside:** Conceptualization, Methodology, Validation, Investigation, Supervision, Writing – Review & Editing, Project administration

## Acknowledgements

We thank all the authors who upon request provided research data and guidance on its interpretation, enabling us to include their studies in our meta-analysis (or at least attempt to do so). These were Getahun Agga, John W. Schmidt, Tim McAllister, Harvey Morgan Scott, Kristin Hales, and Coltan Dornbach. We also thank all the authors who made to effort to find and provide research data that we did not eventually include in the meta-analysis. These were Lauren Wottlin, Raghavendra Amachawadi, Orhan Sahin, David Renter, Davin Holman, G Douglas Inglis, François Malouin, and Bill Epperson.

## 7 Transparency declarations

Mariano Fernández-Miyakawa started working as Head of R&D (Feed Additives) at VETANCO South America in September 2024 during the course of conducting this review.

## 8 Availability of data and materials

The input data for the meta-analysis is deposited at Open Science Framework (https://doi.org/10.17605/OSF.IO/2FYC8). This data can be used alongside the R code deposited at the lead author’s GitHub repository for this study (https://github.com/befriendabacterium/beefantibiotics-SRMA) to reproduce the meta-analysis. The input data, code, and its output data are permanently archived at Zenodo (https://doi.org/10.5281/zenodo.20466930).

