## Supplementary Data for "Effects of antibiotics on the abundance of antibiotic resistance determinants during and after antibiotic administration to beef cattle: A systematic review and meta-analysis of longitudinal studies"

Matt Lloyd Jonesᵃ, Alfredo Sánchez-Tójarᵇᶜᵈ, Alison Bethelᵇᵉ, Anne Leonardᵃ, Emma Lambᵃ, Natalia Casanovaᶠ, Johana Dominguezᶠ, María Paula Quirogaᵍʰᶦ, Daniela Centronᵍʰ, Adriana Peralta Alonsoʲ, Mariano Fernández-Miyakawaᶠ, Professor William Gazeᵃ, Professor Alejandro Petroniᵏ, Professor Ruth Garsideᵃ

*Running title: Antibiotic resistance during and after antibiotic use in beef cattle*

ᵃEuropean Centre for Environment and Human Health, Peter Lanyon Building University of Exeter Penryn TR10 8RD, United Kingdom

ᵇDepartment of Evolutionary Biology, Bielefeld University, Bielefeld, Germany

ᶜCNC-UC, Centre for Neuroscience and Cell Biology, University of Coimbra, Coimbra, Portugal

ᵈCIBB, Centre for Innovative Biomedicine and Biotechnology, University of Coimbra, Coimbra, Portugal

ᵉNIHR CLAHRC South West Peninsula, University of Exeter Medical School, South Cloisters, St Luke’s Campus, Exeter, Devon, United Kingdom

ᶠInstituto de Patobiología Veterinaria, Instituto Nacional de Tecnología Agropecuaria / Consejo Nacional de Investigaciones Científicas y Tecnológicas, INTA-CONICET

ᵍDepartamento de Microbiología, Parasitología e Inmunología, Facultad de Medicina, Universidad de Buenos Aires

ʰLaboratorio de Investigaciones en Mecanismos de Resistencia a Antibióticos (LIMRA), Instituto de Investigaciones en Microbiología y Parasitología Médica (IMPaM), CONICET - Universidad de Buenos Aires

ᶦClúster de Bioinformática, Instituto de Investigaciones en Microbiología y Parasitología Médica (IMPaM), CONICET - Universidad de Buenos Aires. Address (same for three affiliations): Paraguay 2155, piso 12º (zip code C1121ABG), Ciudad Autónoma de Buenos Aires, Argentina

ʲInstitution: Instituto de Ingeniería Rural (IIR), Instituto Nacional de Tecnología Agropecuaria (INTA)

ᵏServicio Antimicrobianos, Departamento de Bacteriología, Instituto Nacional de Enfermedades Infecciosas (INEI) - ANLIS “Dr. Carlos G. Malbrán”

**Table of contents**

***List of Supplementary Tables***

Supplementary Table 1: PRISMA 2020 Checklist. From: Page MJ, McKenzie JE, Bossuyt PM, Boutron I, Hoffmann TC, Mulrow CD, et al. The PRISMA 2020 statement: an updated guideline for reporting systematic reviews. BMJ 2021;372:n71. doi: 10.1136/bmj.n71. This work is licensed under CC BY 4.0. To view a copy of this license, visit https://creativecommons.org/licenses/by/4.0/	26

Supplementary Table 2: Protocol deviations and unregistered steps table, adapted from Willroth and Atherton (2024)	29

Supplementary Table 3: Reasons for excluding studies at full text stage	142

Supplementary Table 4: Certainty assessment table	226

***List of Supplementary Information***

Supplementary Information 1: Rationale for focusing on absolute, rather than relative abundance	25

Supplementary Information 2: Full eligibility criteria	42

Supplementary Information 3: Search strategy	58

Supplementary Information 4: Extracted data	72

Supplementary Information 5: Methods used to prepare the data for quantitative synthesis	79

Supplementary Information 6: Extracted data for meta-analysis	88

Supplementary Information 7: Aggregating estimates of the effect at the study level	125

Supplementary Information 8: Sensitivity testing methods	128

Supplementary Information 9: Reporting bias methods	131

Supplementary Information 10: Certainty assessment methods	133

Supplementary Information 11: Reasons for excluding studies from the meta-analysis	155

Supplementary Information 12: Reasons for excluding effects from the meta-analysis	156

Supplementary Information 13: Full RoB2 assessment	159

Supplementary Information 14: Full RoB2 assessment	160

Supplementary Information 15: Full ROBINSI assessment	161

Supplementary Information 16: Description of random effect estimates in the intercept-only models of the overall effect (hypothesis 1)	177

Supplementary Information 17: Description of random effect estimates in the meta-regression models of moderating effect of time (hypothesis 2)	192

Supplementary Information 18: Full results of subgroup analyses	206

***List of Supplementary Figures***

Supplementary Figure 1: PRISMA2020 flow diagram of the systematic review process for the results of our main, English-language searches of key bibliographic databases. *Specific databases searched via the Web of Science platform were SCI-EXPANDED, SSCI, A&HCI, CPCI-S, CPCI-SSH, ESCI. †Searched in English.	137

Supplementary Figure 2: PRISMA2020 flow diagram of the systematic review process for the results of our 2022 update of the English-language searches in the three bibliographic databases that produced the most results in our initial searches. *Specific databases searched via the Web of Science platform were SCI-EXPANDED, SSCI, A&HCI, CPCI-S, CPCI-SSH, ESCI. †Searched in English. Records removed for other reasons are records published before 2020, which should have been captured by the previous searches but escaped deduplication because of re-indexing by the bibliographic database providers.	138

Supplementary Figure 3: PRISMA2020 flow diagram of the systematic review process for the results of our 2024 update of the English-language searches in the three bibliographic databases that produced the most results in our initial searches. *Specific databases searched via the Web of Science platform were SCI-EXPANDED, SSCI, A&HCI, CPCI-S, CPCI-SSH, ESCI. †Searched in English. Records removed for other reasons are records published before 2022, which should have been captured by the previous searches but escaped deduplication because of re-indexing by the bibliographic database providers	139

Supplementary Figure 4: PRISMA2020 flow diagram of the systematic review process for the results of our 2025 update of the English-language searches in the three bibliographic databases that produced the most results in our initial searches. *Specific databases searched via the Web of Science platform were SCI-EXPANDED, SSCI, A&HCI, CPCI-S, CPCI-SSH, ESCI. †Searched in English. Records removed for other reasons are records published before 2024, which should have been captured by the previous searches but escaped deduplication because of re-indexing by the bibliographic database providers	140

Supplementary Figure 5: PRISMA2020 flow diagram of the systematic review process for the results of our additional English- and Spanish-language searches of Latin American databases. †Searched in English. ‡Searched in Spanish.	141

Supplementary Figure 6: Per-domain and overall risk of bias assessments for the 38 publications (reporting 33 studies) included in the review. Randomised trials are indicated by the † symbol and were assessed using the RoB 2 IPRG tool; cluster-randomised trials are indicated by the ‡ symbol and were assessed using the RoB 2 Cluster tool; non-randomised studies are indicated by * symbol. In order to present all study designs in one figure, domain were consolodated between the three risk tools according to their approximate equivalence where applicable. Hence D1 = Bias due to confounding (RoB 2 IPRG/Cluster D1; ROBINS-I D1); D2 = Bias due to selection of participants (RoB 2 Cluster D1b 'Timing of recruitment'; ROBINS-I D2); D3 = Bias due to classification of interventions (ROBINS-I only); D4 = Bias due to deviations from intended interventions (RoB 2 IPRG/Cluster D2; ROBINS-I D4); D5 = Bias due to missing data (RoB 2 IPRG/Cluster D3; ROBINS-I D5); D6 = Bias due to measurement of the outcome (RoB 2 IPRG/Cluster D4; ROBINS-I D6); D7 = Bias due to selection of the reported result (RoB 2 IPRG/Cluster D5; ROBINS-I D7). Similarly, judgements are consolodated into the following unified framework: Low (Yellow X; all tools); Some concerns/Moderate (Orange -; all tools); High/Serious (Dark orange; all tools); Critical (Dark;= red; ROBINS-I/non-randomised studies only).	158

Supplementary Figure 7: Forest plot of effects on absolute abundance (density) of antibiotic resistant bacteria in faeces pre-antibiotic administration, aggregated at the study level. Effects were aggregated using an intercept-only random effects model accounting for non-independence between effects from the same study, between temporally-autocorrelated effects from the same study arm, and residual variance as random effects.	163

Supplementary Figure 8: Forest plot of effects on logit-transformed relative abundance (logit-transformed proportion) of antibiotic resistant bacteria in faeces post-antibiotic administration, aggregated at the study level. Effects were aggregated using an intercept-only random effects model accounting for non-independence between effects from the same study, between temporally-autocorrelated effects from the same study arm, and residual variance as random effects.	164

Supplementary Figure 9: Forest plot of effects on logit-transformed relative abundance (logit-transformed proportion) of antibiotic resistant bacteria in faeces pre-antibiotic administration, aggregated at the study level. Effects were aggregated using an intercept-only random effects model accounting for non-independence between effects from the same study, between temporally-autocorrelated effects from the same study arm, and residual variance as random effects.	165

Supplementary Figure 10: Forest plot of effects on arcsine-transformed relative abundance (arcsine-transformed proportion) of bacteria in faeces post-antibiotic administration, aggregated at the study level. Effects were aggregated using an intercept-only random effects model accounting for non-independence between effects from the same study, between temporally-autocorrelated effects from the same study arm, and residual variance as random effects.	166

Supplementary Figure 11: Forest plot of effects on arcsine-transformed relative abundance (arcsine-transformed proportion) of antibiotic resistant bacteria in faeces pre-antibiotic administration, aggregated at the study level. Effects were aggregated using an intercept-only random effects model accounting for non-independence between effects from the same study, between temporally-autocorrelated effects from the same study arm, and residual variance as random effects.	167

Supplementary Figure 12: Forest plot of effects on absolute abundance of antibiotic resistant bacteria in pen surface soil post-antibiotic administration. Effects were aggregated at the study level using an intercept-only random effects model accounting for non-independence between effects from the same study, between temporally-autocorrelated effects from the same study arm, and residual variance as random effects. These were then pooled into an overall effect using an equal-effects model sensu Viechtbauer (2024b).	168

Supplementary Figure 13: Forest plot of effects on absolute abundance of antibiotic resistant bacteria in pen surface soil pre-antibiotic administration, aggregated at the study level. Effects were aggregated using an intercept-only random effects model accounting for meta-analytically non-independence between effects from the same study, between temporally-autocorrelated effects from the same study arm, and residual variance as random effects.)	169

Supplementary Figure 14: Forest plot of effects on relative abundance (logit-transformed proportion) of antibiotic resistant bacteria in pen surface soil post-antibiotic administration, aggregated at the study level. Effects were aggregated using an intercept-only random effects model accounting for non-independence between effects from the same study, between temporally-autocorrelated effects from the same study arm, and residual variance as random effects.	170

Supplementary Figure 15: Forest plot of effects on relative abundance (logit-transformed proportion) of antibiotic resistant bacteria in pen surface soil pre-antibiotic administration, aggregated at the study level. Effects were aggregated using an intercept-only random effects model accounting for non-independence between effects from the same study, between temporally-autocorrelated effects from the same study arm, and residual variance as random effects.	171

Supplementary Figure 16: Forest plot of effects on relative abundance (arcine-transformed proportion) of antibiotic resistant bacteria in pen surface soil post-antibiotic administration, aggregated at the study level. Effects were aggregated using an intercept-only random effects model accounting for non-independence between effects from the same study, between temporally-autocorrelated effects from the same study arm, and residual variance as random effects.	172

Supplementary Figure 17: Forest plot of effects on relative abundance (arcsine-transformed proportion) of antibiotic resistant bacteria in pen surface soil pre-antibiotic administration, aggregated at the study level. Effects were aggregated using an intercept-only random effects model accounting for non-independence between effects from the same study, between temporally-autocorrelated effects from the same study arm, and residual variance as random effects.	173

Supplementary Figure 18: Profile likelihood plots for each of the random effect estimates from the intercept-only model of the overall effect during the administration antibiotics. Where the random effect is accurately estimated, the random effect estimate should occur at the peak log-likelihood value (intersection of the two dotted lines). Profile likelihood plots shown are for the A) σ²₁ parameter, representing the random effect of study; B) σ²₂ parameter, representing the random effect of residual variance due to multiple effect sizes estimates per sampling event; C) τ² parameter, representing the random effect of study arm (outer term); D) ρ parameter, representing the random effect temporal autocorrelation within study arms (inner term)	178

Supplementary Figure 19: Profile likelihood plots for each of the random effect estimates from the intercept-only model of the overall effect during the administration antibiotics. Where the random effect is accurately estimated, the random effect estimate should occur at the peak log-likelihood value (intersection of the two dotted lines). Profile likelihood plots shown are for the A) σ²₁ parameter, representing the random effect of study; B) σ²₂ parameter, representing the random effect of residual variance due to multiple effect sizes estimates per sampling event; C) τ² parameter, representing the random effect of study arm (outer term); D) ρ parameter, representing the random effect temporal autocorrelation within study arms (inner term)	179

Supplementary Figure 20: Plots representing results of meta-analytical models testing our two hypotheses for effects measured during the administration of antibiotics. In these plots, colours of the points represent the studies from which they are derived, and their size is proportional to their inverse standard error (1/SE; small points are less precise estimates, larger points are more precise estimates). A) Orchard plot representing the modelled overall effect of antibiotic administration on the absolute abundance of resistant bacteria/resistance genes relating to the intervention antibiotic class in the targeted bacterial population. This overall effect alongside its heterogeneity estimate, are derived from the intercept-only model. In this plot, the overall effect (model intercept after accounting for random effects) is represented by the 'trunk' of the tree (black circle), with its confidence interval represented its 'branch' (shorter, thicker line through it), and its prediction interval represented by its 'twig' (longer, thinner line through it).B) Bubble plot representing the modelled relationship between linear timing of sampling (days of continuous antibiotic administration) and effect size (SMDH) from the meta-regression model. In this plot, the modelled effect of time is represented by the bold line, with the dashed line and dotted lines representing the 95% confidence and prediction intervals, respectively.	180

Supplementary Figure 21: Plots representing results of meta-analytical models testing our two hypotheses for effects on the absolute abundance of all determinants measured during the administration of antibiotics. In these plots, colours of the points represent the studies from which they are derived, and their size is proportional to their inverse standard error (1/SE; small points are less precise estimates, larger points are more precise estimates). A) Orchard plot representing the modelled overall effect of antibiotic administration on the absolute abundance of resistant bacteria/resistance genes relating to the intervention antibiotic class in the targeted bacterial population. This overall effect alongside its heterogeneity estimate, are derived from the intercept-only model. In this plot, the overall effect (model intercept after accounting for random effects) is represented by the 'trunk' of the tree (black circle), with its confidence interval represented its 'branch' (shorter, thicker line through it), and its prediction interval represented by its 'twig' (longer, thinner line through it). B) Bubble plot representing the modelled moderating effect of linear timing of sampling (days of continuous antibiotic administration) and residual heterogeneity from the meta-regression model. In this plot, the modelled effect of time is represented by the bold line, with the dashed line and dotted lines representing the 95% confidence and prediction intervals, respectively.	181

Supplementary Figure 22: Plots representing results of meta-analytical models testing our two hypotheses for effects on the absolute abundance of all determinants measured during the administration of antibiotics. In these plots, colours of the points represent the studies from which they are derived, and their size is proportional to their inverse standard error (1/SE; small points are less precise estimates, larger points are more precise estimates). A) Orchard plot representing the modelled overall effect of antibiotic administration on the absolute abundance of resistant bacteria/resistance genes relating to the intervention antibiotic class in the targeted bacterial population. This overall effect alongside its heterogeneity estimate, are derived from the intercept-only model. In this plot, the overall effect (model intercept after accounting for random effects) is represented by the 'trunk' of the tree (black circle), with its confidence interval represented its 'branch' (shorter, thicker line through it), and its prediction interval represented by its 'twig' (longer, thinner line through it). B) Bubble plot representing the modelled moderating effect of non-linear timing of sampling (days of continuous antibiotic administration) and residual heterogeneity from the meta-regression model. In this plot, the modelled effect of time is represented by the bold line, with the dashed line and dotted lines representing the 95% confidence and prediction intervals, respectively.	182

Supplementary Figure 23: Orchard plots representing the sensitivity of the modelled overall effect during the administration of antibiotics to leaving one study out. In these plots, the mean overall effect is represented by the 'trunk' of the tree (black circle), with its confidence interval represented its 'branch' (shorter, thicker line through it), and its prediction interval represented by its 'twig' (longer, thinner line through it). The mean and precision (1/standard error) of the individual effect sizes from studies included in the model is represented by the position and size of the coloured points, respectively.	184

Supplementary Figure 24: Orchard plots representing the sensitivity of the modelled overall effect during the administration of antibiotics to correcting for shared control non-independence and the removal of potentially problematic studies. In these plots, the mean overall effect is represented by the 'trunk' of the tree (black circle), with its confidence interval represented its 'branch' (shorter, thicker line through it), and its prediction interval represented by its 'twig' (longer, thinner line through it). The mean and precision (1/standard error) of the individual effect sizes from studies included in the model is represented by the position and size of the coloured points, respectively.	185

Supplementary Figure 25: Bubble plots representing the sensitivity of the modelled moderating effect of timing of sampling during the administration of antibiotics to leaving one study out. In these plots, the modelled effect of time is represented by the bold line, with the dashed line and dotted lines representing the 95% confidence and prediction intervals, respectively. The mean and precision (1/standard error) of the individual effect sizes from studies included in the model is represented by the position and size of the coloured points, respectively.	186

Supplementary Figure 26: Bubble plots representing the sensitivity of the modelled moderating effect of timing of sampling during the administration of antibiotics to correcting for shared control non-independence and the removal of potentially problematic studies. In these plots, the modelled effect of time is represented by the bold line, with the dashed line and dotted lines representing the 95% confidence and prediction intervals, respectively. The mean and precision (1/standard error) of the individual effect sizes from studies included in the model is represented by the position and size of the coloured points, respectively.	187

Supplementary Figure 27: Orchard plots representing the sensitivity of the modelled overall effect during the administration of antibiotics to assumptions about the correlation between effect sizes from the same study. Orchard plots of the original model and assuming correlations of 0, 0.2, 0.4, 0.6 and 0.8 are shown. In these plots, the mean overall effect is represented by the 'trunk' of the tree (black circle), with its confidence interval represented its 'branch' (shorter, thicker line through it), and its prediction interval represented by its 'twig' (longer, thinner line through it).	188

Supplementary Figure 28: Bubble plots representing the sensitivity of the moderating effect of timing of sampling on the effect during the administration of antibiotics to assumptions about the correlation between effect sizes from the same study. Bubble plots of the original model and assuming correlations of 0, 0.2, 0.4, 0.6 and 0.8 are shown. In these plots, the modelled effect of time is represented by the bold line, with the dashed line and dotted lines representing the 95% confidence and prediction intervals, respectively. The mean and precision (1/standard error) of the individual effect sizes from studies included in the model is represented by the position and size of the coloured points, respectively.	189

Supplementary Figure 29: Profile likelihood plots for each of the random effect estimates from the intercept-only model of the overall effect after the administration antibiotics. Where the random effect is accurately estimated, the random effect estimate should occur at the peak log-likelihood value (intersection of the two dotted lines). Profile likelihood plots shown are for the A) σ²₁ parameter, representing the random effect of study; B) σ²₂ parameter, representing the random effect of residual variance due to multiple effect sizes estimates per sampling event; C) τ² parameter, representing the random effect of study arm (outer term); D) ρ parameter, representing the random effect temporal autocorrelation within study arms (inner term)	193

Supplementary Figure 30: Profile likelihood plots for each of the random effect estimates from the intercept-only model of the overall effect after the administration antibiotics. Where the random effect is accurately estimated, the random effect estimate should occur at the peak log-likelihood value (intersection of the two dotted lines). Profile likelihood plots shown are for the A) σ²₁ parameter, representing the random effect of study; B) σ²₂ parameter, representing the random effect of residual variance due to multiple effect sizes estimates per sampling event; C) τ² parameter, representing the random effect of study arm (outer term); D) ρ parameter, representing the random effect temporal autocorrelation within study arms (inner term)	194

Supplementary Figure 31: Plots representing results of meta-analytical models testing our two hypotheses for effects measured during the administration of antibiotics. In these plots, colours of the points represent the studies from which they are derived, and their size is proportional to their inverse standard error (1/SE; small points are less precise estimates, larger points are more precise estimates). A) Orchard plot representing the modelled overall effect of antibiotic administration on the absolute abundance of resistant bacteria/resistance genes relating to the intervention antibiotic class in the targeted bacterial population. This overall effect alongside its heterogeneity estimate, are derived from the intercept-only model. In this plot, the overall effect (model intercept after accounting for random effects) is represented by the 'trunk' of the tree (black circle), with its confidence interval represented its 'branch' (shorter, thicker line through it), and its prediction interval represented by its 'twig' (longer, thinner line through it).B) Bubble plot representing the modelled relationship between non-linear timing of sampling (days of continuous antibiotic administration) and effect size (SMDH) from the meta-regression model. In this plot, the modelled effect of time is represented by the bold line, with the dashed line and dotted lines representing the 95% confidence and prediction intervals, respectively.	195

Supplementary Figure 32: Plots representing results of meta-analytical models testing our two hypotheses for effects on the absolute abundance of all determinants measured after the administration of antibiotics. In these plots, colours of the points represent the studies from which they are derived, and their size is proportional to their inverse standard error (1/SE; small points are less precise estimates, larger points are more precise estimates). A) Orchard plot representing the modelled overall effect of antibiotic administration on the absolute abundance of resistant bacteria/resistance genes relating to the intervention antibiotic class in the targeted bacterial population. This overall effect alongside its heterogeneity estimate, are derived from the intercept-only model. In this plot, the overall effect (model intercept after accounting for random effects) is represented by the 'trunk' of the tree (black circle), with its confidence interval represented its 'branch' (shorter, thicker line through it), and its prediction interval represented by its 'twig' (longer, thinner line through it). B) Bubble plot representing the modelled moderating effect of linear timing of sampling (days of continuous antibiotic administration) and residual heterogeneity from the meta-regression model. In this plot, the modelled effect of time is represented by the bold line, with the dashed line and dotted lines representing the 95% confidence and prediction intervals, respectively.	196

Supplementary Figure 33: Plots representing results of meta-analytical models testing our two hypotheses for effects on the absolute abundance of all determinants measured after the administration of antibiotics. In these plots, colours of the points represent the studies from which they are derived, and their size is proportional to their inverse standard error (1/SE; small points are less precise estimates, larger points are more precise estimates). A) Orchard plot representing the modelled overall effect of antibiotic administration on the absolute abundance of resistant bacteria/resistance genes relating to the intervention antibiotic class in the targeted bacterial population. This overall effect alongside its heterogeneity estimate, are derived from the intercept-only model. In this plot, the overall effect (model intercept after accounting for random effects) is represented by the 'trunk' of the tree (black circle), with its confidence interval represented its 'branch' (shorter, thicker line through it), and its prediction interval represented by its 'twig' (longer, thinner line through it). B) Bubble plot representing the modelled moderating effect of non-linear timing of sampling (days of continuous antibiotic administration) and residual heterogeneity from the meta-regression model. In this plot, the modelled effect of time is represented by the bold line, with the dashed line and dotted lines representing the 95% confidence and prediction intervals, respectively.	197

Supplementary Figure 34: Orchard plots representing the sensitivity of the modelled overall effect after the cessation of antibiotic administration to leaving one study out. In these plots, the mean overall effect is represented by the 'trunk' of the tree (black circle), with its confidence interval represented its 'branch' (shorter, thicker line through it), and its prediction interval represented by its 'twig' (longer, thinner line through it). The mean and precision (1/standard error) of the individual effect sizes from studies included in the model is represented by the position and size of the coloured points, respectively.	199

Supplementary Figure 35: Orchard plots representing the sensitivity of the modelled overall effect during the administration of antibiotics to correcting for shared control non-independence and the removal of potentially problematic studies. In these plots, the mean overall effect is represented by the 'trunk' of the tree (black circle), with its confidence interval represented its 'branch' (shorter, thicker line through it), and its prediction interval represented by its 'twig' (longer, thinner line through it). The mean and precision (1/standard error) of the individual effect sizes from studies included in the model is represented by the position and size of the coloured points, respectively.	200

Supplementary Figure 36: Bubble plots representing the sensitivity of the modelled moderating effect of timing of sampling after the cessation of antibiotic administration to leaving one study out. In these plots, the modelled effect of time is represented by the bold line, with the dashed line and dotted lines representing the 95% confidence and prediction intervals, respectively. The mean and precision (1/standard error) of the individual effect sizes from studies included in the model is represented by the position and size of the coloured points, respectively.	201

Supplementary Figure 37: Bubble plots representing the sensitivity of the modelled overall effect after the administration of antibiotics to correcting for shared control non-independence and the removal of potentially problematic studies. In these plots, the modelled effect of time is represented by the bold line, with the dashed line and dotted lines representing the 95% confidence and prediction intervals, respectively. The mean and precision (1/standard error) of the individual effect sizes from studies included in the model is represented by the position and size of the coloured points, respectively.	202

Supplementary Figure 38: Orchard plots representing the sensitivity of the modelled overall effect after the administration of antibiotics to assumptions about the correlation between effect sizes from the same study. Orchard plots of the original model and assuming correlations of 0, 0.2, 0.4, 0.6 and 0.8 are shown. In these plots, the mean overall effect is represented by the 'trunk' of the tree (black circle), with its confidence interval represented its 'branch' (shorter, thicker line through it), and its prediction interval represented by its 'twig' (longer, thinner line through it).	203

Supplementary Figure 39: Bubble plots representing the sensitivity of the moderating effect of timing of sampling on the effect after the administration of antibiotics to assumptions about the correlation between effect sizes from the same study. Bubble plots of the original model and assuming correlations of 0, 0.2, 0.4, 0.6 and 0.8 are shown. In these plots, the modelled effect of time is represented by the bold line, with the dashed line and dotted lines representing the 95% confidence and prediction intervals, respectively. The mean and precision (1/standard error) of the individual effect sizes from studies included in the model is represented by the position and size of the coloured points, respectively.	204

Supplementary Figure 40: Orchard plot representing results of the meta-analytical model testing for a subgroup effect of mode of administering the intervention groups (via feed or via injection). Points are effect sizes, with their colours representing the studies from which they are derived and their size being proportional to their inverse standard error (1/SE; small points are less precise estimates, larger points are more precise estimates). The pooled effect for each subgroup (coefficient after accounting for random effects) is represented by the 'trunk' of the tree (black circle), with its confidence interval represented its 'branch' (shorter, thicker line through it), and its prediction interval represented by its 'twig' (longer, thinner line through it). For the numbers above each subgroup, 'k' represents the number of effect sizes, whilst the number in brackets represents the number of studies from which they were derived.	209

Supplementary Figure 41: Orchard plot representing results of the meta-analytical model testing for a subgroup effect of whether or not intervention cattle were housed alongside control cattle. Points are effect sizes, with their colours representing the studies from which they are derived and their size being proportional to their inverse standard error (1/SE; small points are less precise estimates, larger points are more precise estimates). The pooled effect for each subgroup (coefficient after accounting for random effects) is represented by the 'trunk' of the tree (black circle), with its confidence interval represented its 'branch' (shorter, thicker line through it), and its prediction interval represented by its 'twig' (longer, thinner line through it). For the numbers above each subgroup, 'k' represents the number of effect sizes, whilst the number in brackets represents the number of studies from which they were derived.	210

Supplementary Figure 42: Orchard plot representing results of the meta-analytical model testing for a subgroup effect of whether or not another antibiotic was administered to intervention (and control) groups alongside the intervention antibiotic (i.e. presence of a cointervention). Points are effect sizes, with their colours representing the studies from which they are derived and their size being proportional to their inverse standard error (1/SE; small points are less precise estimates, larger points are more precise estimates). The pooled effect for each subgroup (coefficient after accounting for random effects) is represented by the 'trunk' of the tree (black circle), with its confidence interval represented its 'branch' (shorter, thicker line through it), and its prediction interval represented by its 'twig' (longer, thinner line through it). For the numbers above each subgroup, 'k' represents the number of effect sizes, whilst the number in brackets represents the number of studies from which they were derived.	211

Supplementary Figure 43: Bubble plot representing the results of meta-analytical models testing for an effect of dose. Points are effect sizes, with their colours representing the studies from which they are derived and their size being proportional to their inverse standard error (1/SE; small points are less precise estimates, larger points are more precise estimates). Plots the meta-regression fits of tests of dose (mg/kg) in the following formats: A) Daily dose of antibiotic administered B) Log10-transformed Daily dose of antibiotic administered C) Cumulative daily dose of antibiotic administered before each effect estimate D) Log10-transformed cumulative daily dose of antibiotic administered before each effect estimate.	212

Supplementary Figure 44: Orchard plot representing results of the meta-analytical model testing for a subgroup effect of the microorganism(s) targetted in the outcome assessment. Points are effect sizes, with their colours representing the studies from which they are derived and their size being proportional to their inverse standard error (1/SE; small points are less precise estimates, larger points are more precise estimates). The pooled effect for each subgroup (coefficient after accounting for random effects) is represented by the 'trunk' of the tree (black circle), with its confidence interval represented its 'branch' (shorter, thicker line through it), and its prediction interval represented by its 'twig' (longer, thinner line through it). For the numbers above each subgroup, 'k' represents the number of effect sizes, whilst the number in brackets represents the number of studies from which they were derived.	213

Supplementary Figure 45: Orchard plot representing results of the meta-analytical model testing for a subgroup effect of the method used to estimate the proportion of the targeted bacterial population that was antibiotic resistant in the outcome assessment. Points are effect sizes, with their colours representing the studies from which they are derived and their size being proportional to their inverse standard error (1/SE; small points are less precise estimates, larger points are more precise estimates). The pooled effect for each subgroup (coefficient after accounting for random effects) is represented by the 'trunk' of the tree (black circle), with its confidence interval represented its 'branch' (shorter, thicker line through it), and its prediction interval represented by its 'twig' (longer, thinner line through it). For the numbers above each subgroup, 'k' represents the number of effect sizes, whilst the number in brackets represents the number of studies from which they were derived.	214

Supplementary Figure 46: Orchard plot representing results of the meta-analytical model testing for a sub-subgroup effect of the antibiotic onto which samples were plated in CFU measures of the proportion of the targeted bacterial population that was antibiotic resistant in the outcome assessment. Points are effect sizes, with their colours representing the studies from which they are derived and their size being proportional to their inverse standard error (1/SE; small points are less precise estimates, larger points are more precise estimates). The pooled effect for each subgroup (coefficient after accounting for random effects) is represented by the 'trunk' of the tree (black circle), with its confidence interval represented its 'branch' (shorter, thicker line through it), and its prediction interval represented by its 'twig' (longer, thinner line through it). For the numbers above each subgroup, 'k' represents the number of effect sizes, whilst the number in brackets represents the number of studies from which they were derived.	215

Supplementary Figure 47: Orchard plot representing results of the meta-analytical model testing for a sub-subgroup effect of the antibiotic resistance genes targeted in qPCR measures of the proportion of the targeted bacterial population that was antibiotic resistant in the outcome assessment. Points are effect sizes, with their colours representing the studies from which they are derived and their size being proportional to their inverse standard error (1/SE; small points are less precise estimates, larger points are more precise estimates). The pooled effect for each subgroup (coefficient after accounting for random effects) is represented by the 'trunk' of the tree (black circle), with its confidence interval represented its 'branch' (shorter, thicker line through it), and its prediction interval represented by its 'twig' (longer, thinner line through it). For the numbers above each subgroup, 'k' represents the number of effect sizes, whilst the number in brackets represents the number of studies from which they were derived.	216

Supplementary Figure 48: Orchard representing results of the second meta-analytical model testing for a sub-subgroup effect of the relative (to the CLSI breakpoint) antibiotic concentration onto which samples were plated in CFU measures of the proportion of the targeted bacterial population that was antibiotic resistant in the outcome assessment(i.e. study breakpoint relative to CLSI breakpoint). Given the uneven spread of concentrations in the previous analysis, for this analysis we categorised antibiotuc concentrations/study breakpoints into four categories based on whether they were below, at, or above the CLSI breakpoint (or there was no CLSI breakpoint for the antibiotic-microorganism combination tested). Points are effect sizes, with their colours representing the studies from which they are derived and their size being proportional to their inverse standard error (1/SE; small points are less precise estimates, larger points are more precise estimates). The pooled effect for each subgroup (coefficient after accounting for random effects) is represented by the 'trunk' of the tree (black circle), with its confidence interval represented its 'branch' (shorter, thicker line through it), and its prediction interval represented by its 'twig' (longer, thinner line through it). For the numbers above each subgroup, 'k' represents the number of effect sizes, whilst the number in brackets represents the number of studies from which they were derived.	217

Supplementary Figure 49: Bubble plot representing results of the meta-analytical model testing for a sub-subgroup effect of the relative (to the CLSI breakpoint) antibiotic concentration onto which samples were plated in CFU measures of the proportion of the targeted bacterial population that was antibiotic resistant in the outcome assessment(i.e. study breakpoint relative to CLSI breakpoint). Points are effect sizes, with their colours representing the studies from which they are derived and their size being proportional to their inverse standard error (1/SE; small points are less precise estimates, larger points are more precise estimates).	218

Supplementary Figure 50: Bubble plot representing results of the meta-analytical model testing for a sub-subgroup effect of the absolute log10-transformed antibiotic concentration onto which samples were plated in CFU measures of the proportion of the targeted bacterial population that was antibiotic resistant in the outcome assessment (i.e. study breakpoint). Points are effect sizes, with their colours representing the studies from which they are derived and their size being proportional to their inverse standard error (1/SE; small points are less precise estimates, larger points are more precise estimates).	219

Supplementary Figure 51: Orchard plot representing results of the meta-analytical model testing for a subgroup effect of whether or not the outcomes underpinning effect sizes accounted for potential pen-level effects. Outcomes were classified according to the study design and analysis underpinning any effect sizes used in the meta-analysis. Briefly, 'Individual (pen effect not accounted for)' refers to effect sizes from studies in which there was a single pen per study arm/one pen for all study arms; 'Individual (pen effect not account for)' refers to extracted effect sizes from analyses at the individual level but which did not account for the fact that study arms are spread across different pens; 'Individual (pen as random effect)' refers to extracted effect sizes from analyses at the individual level but which did account for the fact that study arms are spread across different pens by using pen as a random effect in the model; 'Pen (individuals as pseudoreplicates)' refers to effect sizes from analyses at the pen level i.e. which average out individual-level variation by treating individuals as pseudoreplicates (this was the method we used to process raw data where available, hence it is the largest category). Points are effect sizes, with their colours representing the studies from which they are derived and their size being proportional to their inverse standard error (1/SE; small points are less precise estimates, larger points are more precise estimates). The pooled effect for each subgroup (coefficient after accounting for random effects) is represented by the 'trunk' of the tree (black circle), with its confidence interval represented its 'branch' (shorter, thicker line through it), and its prediction interval represented by its 'twig' (longer, thinner line through it). For the numbers above each subgroup, 'k' represents the number of effect sizes, whilst the number in brackets represents the number of studies from which they were derived.	220

Supplementary Figure 52: Orchard plot representing results of the meta-analytical model testing for a subgroup effect of study design. Points are effect sizes, with their colours representing the studies from which they are derived and their size being proportional to their inverse standard error (1/SE; small points are less precise estimates, larger points are more precise estimates). The pooled effect for each subgroup (coefficient after accounting for random effects) is represented by the 'trunk' of the tree (black circle), with its confidence interval represented its 'branch' (shorter, thicker line through it), and its prediction interval represented by its 'twig' (longer, thinner line through it). For the numbers above each subgroup, 'k' represents the number of effect sizes, whilst the number in brackets represents the number of studies from which they were derived.	221

Supplementary Figure 53: Publication bias plots	223

Supplementary Figure 54: Publication bias plots	224

Figure 1: Plots representing results of meta-analytical models testing our two hypotheses for effects measured after the administration of antibiotics. In these plots, points are effect sizes, with their colours representing the studies from which they are derived and their size being proportional to their inverse standard error (1/SE; small points are less precise estimates, larger points are more precise estimates). A) Orchard plot representing the modelled overall effect of antibiotic administration on the absolute abundance of resistant bacteria/resistance genes relating to the intervention antibiotic class in the targeted bacterial population that was resistant to the antibiotic class administered. This overall effect alongside its heterogeneity estimate, are derived from the intercept-only model. In this plot, the overall effect (model intercept after accounting for random effects) is represented by the 'trunk' of the tree (black circle), with its confidence interval represented its 'branch' (shorter, thicker line through it), and its prediction interval represented by its 'twig' (longer, thinner line through it). The prediction interval is however illegible as it is identical to the confidence interval when τ2=0, which is the case here (see Viechtbauer, 2025) B) Bubble plot representing the modelled relationship between timing of sampling (days of continuous antibiotic administration) and effect size (SMDH) from the meta-regression model. In this plot, the modelled effect of time is represented by the bold line, with the dashed line and dotted lines representing the 95% confidence and prediction intervals, respectively. The prediction interval is however illegible as it is identical to the confidence interval when τ2=0, which is the case here (see Viechtbauer, 2025).	242

### 1 Supplementary Methods

#### 1.1 Registration and reporting

Supplementary Information 1: Rationale for focusing on absolute, rather than relative abundance

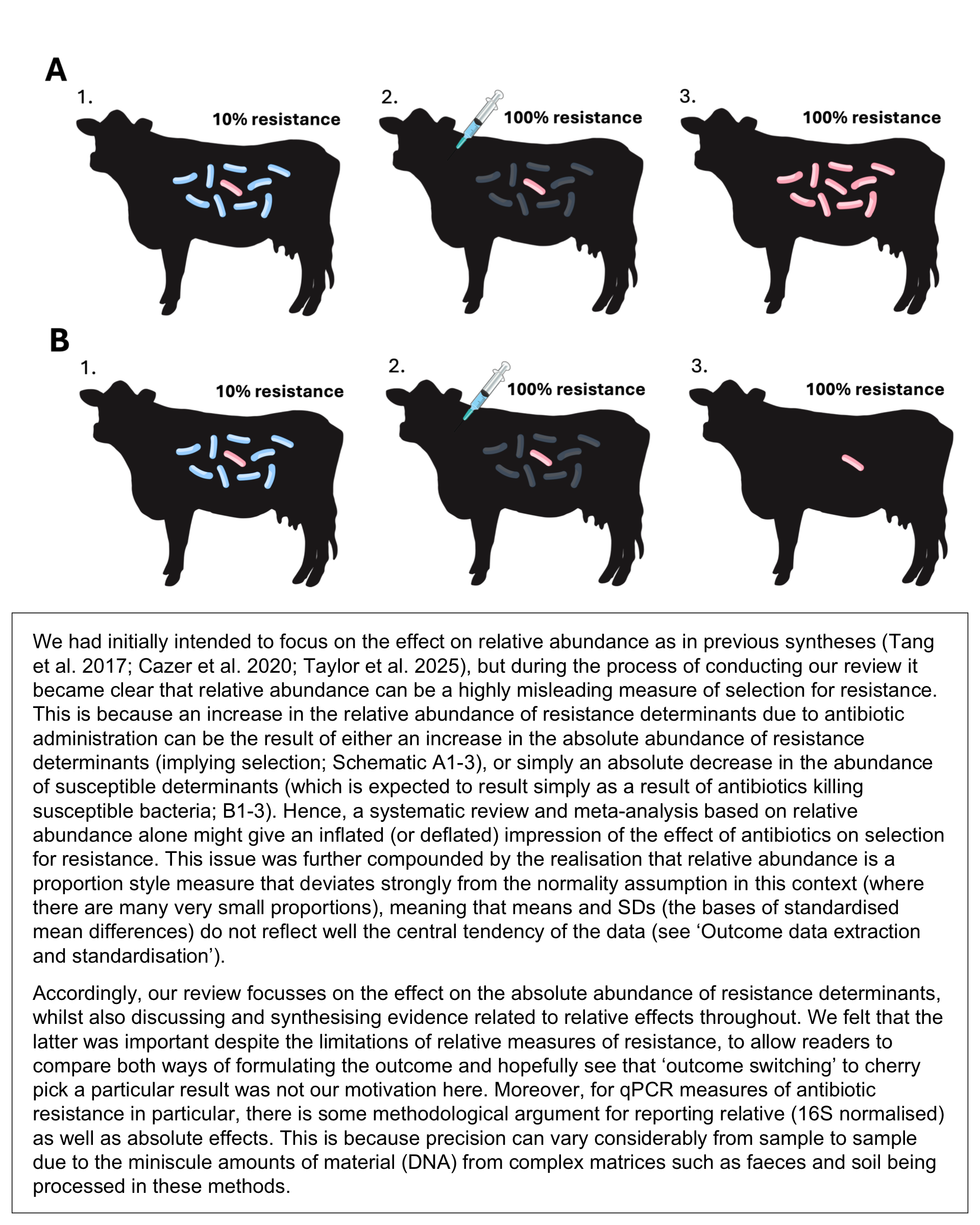

Illustrations used in schematic are from the NIH BioArt Source^1–4^.

Supplementary Table 1: PRISMA 2020 Checklist. From: Page MJ, McKenzie JE, Bossuyt PM, Boutron I, Hoffmann TC, Mulrow CD, et al. The PRISMA 2020 statement: an updated guideline for reporting systematic reviews. BMJ 2021;372:n71. doi: 10.1136/bmj.n71. This work is licensed under CC BY 4.0. To view a copy of this license, visit https://creativecommons.org/licenses/by/4.0/

| **Section and topic** | **Item #** | **Checklist item** | **Location where item is reported** |
| --- | --- | --- | --- |
| **TITLE** | | | |
| Title | 1 | Identify the report as a systematic review. | Title |
| **ABSTRACT** | | | |
| Abstract | 2 | See the PRISMA 2020 for Abstracts checklist (table 2). | Abstract |
| **INTRODUCTION** | | | |
| Rationale | 3 | Describe the rationale for the review in the context of existing knowledge. | Rationale |
| Objectives | 4 | Provide an explicit statement of the objective(s) or question(s) the review addresses. | Objectives |
| **METHODS** | | | |
| Eligibility criteria | 5 | Specify the inclusion and exclusion criteria for the review and how studies were grouped for the syntheses. | Methods: Eligibility criteria |
| Information sources | 6 | Specify all databases, registers, websites, organisations, reference lists and other sources searched or consulted to identify studies. Specify the date when each source was last searched or consulted. | Methods: Information sources & search strategy |
| Search strategy | 7 | Present the full search strategies for all databases, registers and websites, including any filters and limits used. | Methods: Information sources & search strategy |
| Selection process | 8 | Specify the methods used to decide whether a study met the inclusion criteria of the review, including how many reviewers screened each record and each report retrieved, whether they worked independently, and if applicable, details of automation tools used in the process. | Methods: Selection process   Methods: Synthesis Methods: Selection process |
| Data collection process | 9 | Specify the methods used to collect data from reports, including how many reviewers collected data from each report, whether they worked independently, any processes for obtaining or confirming data from study investigators, and if applicable, details of automation tools used in the process. | Methods: Data collection process and data items |
| Data items | 10a | List and define all outcomes for which data were sought. Specify whether all results that were compatible with each outcome domain in each study were sought (e.g. for all measures, time points, analyses), and if not, the methods used to decide which results to collect. | Methods: Data collection process and data items   Supporting Information 3 |
|  | 10b | List and define all other variables for which data were sought (e.g. participant and intervention characteristics, funding sources). Describe any assumptions made about any missing or unclear information. | Methods: Data collection process and data items   Supporting Information 3 |
| Study risk of bias assessment | 11 | Specify the methods used to assess risk of bias in the included studies, including details of the tool(s) used, how many reviewers assessed each study and whether they worked independently, and if applicable, details of automation tools used in the process. | Methods: Study risk of bias assessment |
| Effect measures | 12 | Specify for each outcome the effect measure(s) (e.g. risk ratio, mean difference) used in the synthesis or presentation of results. | Methods: Effect measures |
| Synthesis methods | 13a | Describe the processes used to decide which studies were eligible for each synthesis (e.g. tabulating the study intervention characteristics and comparing against the planned groups for each synthesis (item #5)). | Methods: Synthesis methods: Selection process |
|  | 13b | Describe any methods required to prepare the data for presentation or synthesis, such as handling of missing summary statistics, or data conversions. | Methods: Synthesis methods: Preparation of data |
|  | 13c | Describe any methods used to tabulate or visually display results of individual studies and syntheses. | Methods: Synthesis methods: Results of individual studies |
|  | 13d | Describe any methods used to synthesise results and provide a rationale for the choice(s). If meta-analysis was performed, describe the model(s), method(s) to identify the presence and extent of statistical heterogeneity, and software package(s) used. | Methods: Synthesis methods: Meta-analysis of the overall effect (hypothesis 1) |
|  | 13e | Describe any methods used to explore possible causes of heterogeneity among study results (e.g. subgroup analysis, meta-regression). | Methods: Synthesis methods: Meta-analysis of the moderation of the effect by time of sampling (hypothesis 2)   Methods: Synthesis methods: Subgroup and other alternative analyses |
|  | 13f | Describe any sensitivity analyses conducted to assess robustness of the synthesised results. | Methods: Synthesis methods: Sensitivity analyses |
| Reporting bias assessment | 14 | Describe any methods used to assess risk of bias due to missing results in a synthesis (arising from reporting biases). | Methods: Reporting bias assessment |
| Certainty assessment | 15 | Describe any methods used to assess certainty (or confidence) in the body of evidence for an outcome. | Methods: Certainty assessment |
| **RESULTS** | | | |
| Study selection | 16a | Describe the results of the search and selection process, from the number of records identified in the search to the number of studies included in the review, ideally using a flow diagram (see fig 1). | Results: Study selection |
|  | 16b | Cite studies that might appear to meet the inclusion criteria, but which were excluded, and explain why they were excluded. | Results: Study selection   Supplementary Table 3 |
| Study characteristics | 17 | Cite each included study and present its characteristics. | Results: Study characteristics |
| Risk of bias in studies | 18 | Present assessments of risk of bias for each included study. | Results: Risk of bias in studies |
| Results of individual studies | 19 | For all outcomes, present, for each study: (a) summary statistics for each group (where appropriate) and (b) an effect estimate and its precision (e.g. confidence/credible interval), ideally using structured tables or plots. | Results: Results of individual studies |
| Results of syntheses | 20a | For each synthesis, briefly summarise the characteristics and risk of bias among contributing studies. | Results: Results of syntheses: Effects measured during/after antibiotic administration: Primary analysis |
|  | 20b | Present results of all statistical syntheses conducted. If meta-analysis was done, present for each the summary estimate and its precision (e.g. confidence/credible interval) and measures of statistical heterogeneity. If comparing groups, describe the direction of the effect. | Results: Results of syntheses: Effects measured during/after antibiotic administration: Primary analysis |
|  | 20c | Present results of all investigations of possible causes of heterogeneity among study results. | Results: Results of syntheses: Effects measured during/after antibiotic administration: Primary analysis |
|  | 20d | Present results of all sensitivity analyses conducted to assess the robustness of the synthesised results. | Results: Results of syntheses: Effects measured during/after antibiotic administration: Sensitivity analysis |
| Reporting biases | 21 | Present assessments of risk of bias due to missing results (arising from reporting biases) for each synthesis assessed. | Results: Reporting bias |
| Certainty of evidence | 22 | Present assessments of certainty (or confidence) in the body of evidence for each outcome assessed. | Results: Certainty of evidence |
| **DISCUSSION** | | | |
| Discussion | 23a | Provide a general interpretation of the results in the context of other evidence. | Discussion: General interpretation of results |
| Discussion | 23b | Discuss any limitations of the evidence included in the review. | Discussion: Limitations of the evidence included in the review |
|  | 23c | Discuss any limitations of the review processes used. | Discussion: Limitations of the review processes used |
|  | 23d | Discuss implications of the results for practice, policy, and future research. | Discussion: Implications for practice, policy, and future research |
| **OTHER INFORMATION** | | | |
| Registration and protocol | 24a | Provide registration information for the review, including register name and registration number, or state that the review was not registered. | Methods: Registration and reporting |
|  | 24b | Indicate where the review protocol can be accessed, or state that a protocol was not prepared. | Methods: Registration and reporting |
|  | 24c | Describe and explain any amendments to information provided at registration or in the protocol. | Supplementary Box 1   Methods   Supporting Information 2 |
| Support | 25 | Describe sources of financial or non-financial support for the review, and the role of the funders or sponsors in the review. | Funding |
| Competing interests | 26 | Declare any competing interests of review authors. | Transparency declaration |
| Availability of data, code, and other materials | 27 | Report which of the following are publicly available and where they can be found: template data collection forms   data extracted from included studies   data used for all analyses   analytic code   any other materials used in the review. | Availability of data, code, and other materials |

Supplementary Table 2: Protocol deviations and unregistered steps table, adapted from Willroth and Atherton (2024)

| **Deviations** | | | | | | |
| --- | --- | --- | --- | --- | --- | --- |
| *No.* | *Details* | | *Original wording* | *New wording* | *Deviation Description* | *Reader Impact* |
| 1 | Type | Objectives | "Is there an increased prevalence of antibiotic resistance during and after antibiotic administration in beef cattle production?" | "In beef cattle (P), what is the effect of administering an antibiotic (I) versus not administering that antibiotic (C) on the abundance of resistance determinants (e.g. resistant bacteria, genes) for that antibiotic’s class in their faeces and environments impacted by it (O)?" | Adopted a PICO-structured research question (Morgan et al. 2018). We use the broad term 'abundance' to cover both absolute and relative abundance, given our review now focusses on the former but also includes the latter as a secondary outcome at the summary stage (New knowledge; see Deviation 3) | Improves clarity as research question is more specific and relatable to PICO criteria. |
|  | Reason | Multiple (Please Explain) |  |  |  |  |
|  | Timing | After data access |  |  |  |  |
| 2 | Type | Objectives | "H1. When an antibiotic is administered to beef cattle, there will be a higher prevalence of resistance to that antibiotic in the faeces and environments of the cattle to which it is administered, compared to that of non-administered cattle.    H2. The strength of this effect (H1) will increase over time during the period of antibiotic administration, and decrease over time after the cessation of antibiotic administration." | "H1. When an antibiotic is administered to beef cattle, there will be a higher abundance of resistance to that antibiotic in the faeces and environments of the cattle to which it is administered, compared to that of non-administered cattle.  H2. The strength of this effect (H1) will increase over time during the period of antibiotic administration, and decrease over time after the cessation of antibiotic administration." | Replaced 'prevalence' with 'abundance', given the former was a misnomer originally intended to refer to 'relative abundance' (Typo/Error). We use the broad term 'abundance' to cover both absolute and relative abundance, given our review now focusses on the former but also includes the latter as a secondary outcome at the summary stage (New knowledge; Deviation 3) | Should avoid confusion by correcting a misnomer ("prevalence") and encompassing both absolute and relative abundance. |
|  | Reason | Multiple (Please Explain) |  |  |  |  |
|  | Timing | After data access |  |  |  |  |
| 3 | Type | Eligibility Criteria | Too detailed to paste here - see PICOS_criteria.pdf at pre-registration. | Too detailed to paste here - see Methods: Eligibility Critieria (Main text) & Supporting Information 1: Full Eligibility Criteria (Supporting Data) | Minor changes to improve clarity in Eligibility Criteria:    - Simplified and corrected descriptions eligible study designs to randomised, cluster-randomised and non-randomised designs. These are not different to the original criteria and how they were applied but we felt it important to correct these to clarify what study designs were included (e.g. avoiding confusion in incorrect description of a cluster-randomised controlled design which was previously equated with a blocked design, and clarifying the distinction between before-and-after and prospective cohort studies.  - Population: No changes  - Intervention/exposure(s): Used term 'study arms' instead of 'groups' to avoid confusion with sub-study arm groups of cattle (e.g. pens across which the study arm is housed, replicates in a cluster-randomised trial). Clarified that indirect administration by distiller's grains is potentially eligible (see Unregistered Steps 1)  - Comparators/controls: Used term 'study arms' instead of 'groups' to avoid confusion with sub-study arm groups of cattle (e.g. pens across which the study arm is housed, replicates in a cluster-randomised trial)  - Outcomes: Used term 'relative abundance' instead of 'prevalence' to avoid confusion. Clarified how this is typically reported and how estimates of absolute abundance can be included as part of this (provided both numerator and denominator available). Described metagenomic and environmental outcomes as outcomes rather than 'secondary outcomes' to avoid confusion with relative abundance, which is considered a 'secondary outcome' in the final review. Clarified contextual wording regarding how metagenomic and environmental outcomes may be included in the review. | Should avoid confusion by correcting a misnomer ("prevalence") and encompassing both absolute and relative abundance. |
|  | Reason | Multiple (Please Explain) |  |  |  |  |
|  | Timing | After data access |  |  |  |  |
| 4 | Type | Eligibility Criteria | "Outcome measure(s)...The antibiotic resistance tested can be different between studies. Antibiotic resistance can be measured in one of two ways:    1. The prevalence (%) of phenotypic antibiotic resistance in tested isolates of an identified group of bacteria (at least to genus level; e.g. E. coli, Enterococcus, Salmonella, Campylobacter) isolated from a fresh faecal sample. Phenotypic antibiotic resistance can be established using broth dilution, agar dilution, disk diffusion, E-tests, selective plating or automated methods.    2. The prevalence (%) of key AMR genes (AMR genes per 16S genes) in DNA extracted from a fresh faecal sample and tested through real-time PCR/qPCR." | "Outcome: An estimate of either: 1) the absolute abundance of both resistant and all bacteria in the targeted population; or 2) the relative abundance of resistance in the targeted population. Estimates of absolute abundance refer to either culture-dependent methods to quantify the density of resistant and all bacteria in the target population (e.g. resistant cells per gram of faeces, all cells per gram of faeces measured via direct plating), or culture-independent methods to quantify the density of resistance and bacterial marker genes (e.g. genes per gram of faeces, 16S genes per gram of faeces measured via qPCR or metagenomics). Estimates of relative abundance can include direct plating and qPCR methods which may be presented or reformulated as relative abundance (e.g. resistance genes per 16S copies), as well as antimicrobial susceptibility testing methods which can only estimate relative abundance (e.g. proportion of 10 tested isolates that were resistant). The targeted resistance phenotype or genotype should relate specifically to the class of the antibiotic administered in the intervention (i.e. direct selection). The targeted bacterial populations should be from samples from fresh beef cattle faeces (direct from the rectum or deposited <24 hours before) or beef cattle associated environments (i.e. pen soil, water or feed). The bacterial populations in which resistance is assessed should be established bacteria rather than bacteria with which the animal has been experimentally colonised or infected (though it is fine for studies to use experimental colonisation/infection as a cointervention alongside an antibiotic intervention)." | The Outcome eligibility criterion was reworded for clarification purposes, but not fundamentally changed. We did not feel it necessary to fundamentally change the eligibility criteria by including studies that only measured the absolute abundance of resistance, given relative abundance information was sometimes all that was available and was useful for including the wider evidence base in the narrative synthesis. We also clarified that we were referring to effects on established bacteria rather than experimentally-infected bacteria (after encountering some studies like this during screening). | Minimal impact of this specific decision. Impact is mainly the loss of 1 study from the meta-analysis (Sharma et al. 2008) from not choosing to broaden our criteria to include studies solely measuring or reporting effects on the absolute abundance resistant determinants. However, this is only one study and there is a scientific argument for measuring absolute abundance of all determinants also, in order to contextualise effects (hence when most studies do this; see Supplementary Box 1). |
|  | Reason | New knowledge |  |  |  |  |
|  | Timing | After results known |  |  |  |  |
| 5 | Type | Information sources and search strategy | We will also search the following databases in Spanish and Portuguese:  • Lilacs  • SciELO  • MedicLATINA  • Latindex  • e-Revistas  • Sistema Nacional de Repositorios Digitales (Government of Argentina)" | "AB and APA also conducted English and/or Spanish-language searches of databases and grey literature sources with more focus on Latin American (among other) regions. This was in order to find relevant evidence from Latin America, which contains several of the world's largest beef producers and is experiencing a shift toward intensive production systems. These sources were LILACS (last searched in Spanish 12 February 2021) Google Académico (last searched in Spanish 6 April 2021), LA Referencia (last searched in English & Spanish 19 February 2021), SIDALC (last searched in English & Spanish 24 February 2021), REDIB (last searched in English & Spanish 18 February 2021), SciELO (last searched in Spanish 7 April 2021), SNRD (last searched in Spanish 17 February 2021), and Redalyc (last searched in English & Spanish 25 March 2021)." | MedicLATINA, LATINDEX, e-Revistas were removed from the Latin American search strategy and LA Referencia, REDIB and SciELO in consultation with and Information Specialist in Argentina based on access restrictions and suitability as alternatives/overlap. | No impact. |
|  | Reason | New knowledge |  |  |  |  |
|  | Timing | After results known |  |  |  |  |
| 6 | Type | Information sources and search strategy | "To ensure comprehensiveness we will use supplementary search methods. We will search websites and online catalogues as well as undertake a Google Scholar/Google Academico search in English, Spanish and Portuguese, we will hand search specific journals and undertake forwards and backwards citation searching of all of the final included papers.    Catalogues/Websites    • WorldCat – English, Spanish & Portuguese  • DEFRA – English only  • GOV.UK Antimicrobial Resistance research (https://www.gov.uk/health-and-social-care/antimicrobial-resistance) – English only  • Núcleo Básico de Revistas Científicas (CONICET database) – English, Spanish & Portuguese REDIB – English, Spanish & Portuguese  • SIDALC – English, Spanish & Portuguese  • SEDICI UNLP – English, Spanish & Portuguese    Hand-searching  • InfoVet (magazine) – English, Spanish & Portuguese  • Revista Veterinaria Argentina (magazine) – English, Spanish & Portuguese" | "AB and APA also conducted English and/or Spanish-language searches of databases and grey literature sources with more focus on Latin American (among other) regions. This was in order to find relevant evidence from Latin America, which contains several of the world's largest beef producers and is experiencing a shift toward intensive production systems. These sources were LILACS (last searched in Spanish 12 February 2021) Google Académico (last searched in Spanish 6 April 2021), LA Referencia (last searched in English & Spanish 19 February 2021), SIDALC (last searched in English & Spanish 24 February 2021), REDIB (last searched in English & Spanish 18 February 2021), SciELO (last searched in Spanish 7 April 2021), SNRD (last searched in Spanish 17 February 2021), and Redalyc (last searched in English & Spanish 25 March 2021)." | Only Google Scholar/Academico and REDIB were searched to keep the workload manageable and since it was considered these databases included at least some of the other sources. | No impact. |
|  | Reason | New knowledge |  |  |  |  |
|  | Timing | After results known |  |  |  |  |
| 7 | Type | Data collection process and data items | "The included outcomes measured in studies are the prevalence of phenotypic or genotypic bacterial antibiotic resistance (antibiotic resistance determinants per tested isolates or 16S genes) in cattle faeces and cattle-associated environments." | "For outcome(s) we sought information on the method and matrix (e.g. faeces) used to measure the abundance of resistance/all bacteria (e.g. direct enumeration), and the reported effects on absolute and/or relative abundance in the publication." | Only Google Scholar/Academico and REDIB were searched to keep the workload manageable and since it was considered these databases included at least some of the other sources. | We only fully realised the interpretability and statistical issues with relative abundance at the analysis stage and so decided to focus the synthesis on absolute abundance whilst drawing comparisons with effects on relative abundance throughout. This enables readers to first see that both ways of formulating the outcome produce qualitatively and quantitatively similar results, whilst also allowing us to use the most interpretively and statistically valid way of formulating the outcome for the main synthesis. |
|  | Reason | New knowledge |  |  |  |  |
|  | Timing | After results known |  |  |  |  |
| 8 | Type | Data collection process and data items | Too detailed to paste here - see data_extraction.pdf at pre-registration. | See Supporting Information 3. | Most of the originally intended data items were extracted or attempted to have been extracted; interested readers can compare our pre-registered and final lists. Main change is addition of items that we realised were important to also extract during the review and the omission/NAing of items that were unfeasible to obtain. | No impact - all of the core data items are extracted where possible, as well as additional items which enrich the analysis. |
|  | Reason | New knowledge |  |  |  |  |
|  | Timing | During data collection |  |  |  |  |
| 9 | Type | Study risk of bias assessment | "Risk of bias will be assessed on the study design for the outcome of a change in the prevalence of antibiotic resistance due to antibiotic administration (regardless of whether this is the focus of the original study design). Risk of bias will be assessed by 2 reviewers using the Cochrane's RoB 2 (experimental studies) and ROBINS-I (observational studies) tools (see https://www.riskofbias.info/ for details on these tools), with the alterations/additions/clarifications listed in the following sections.    Before deploying the tools fully, each reviewer will test the tools on each of 3 study types; 1 randomised controlled trial (RoB 2), 1 non-randomised controlled trial (ROBINS-I) and 1 cohort study (ROBINS-I). Reviewers will then discuss, adjust and clarify any areas of confusion in the use of the tools before their full deployment.    Following testing, each reviewer will assess the risk of bias for their full collection of assigned manuscripts (randomly allocated from each of the 3 types of study so that each reviewer has approximately the same number of each study type to assess). Studies will be designated as having a 'Low', 'Some concerns/Moderate' or 'High/Serious' risk of bias according the Cochrane tools. Disagreements will be resolved by consensus or through a third reviewer." | "Risk of bias was assessed for all included using Cochrane’s Risk of Bias 2 (RoB 2) tool for randomised controlled trials, and Cochrane’s ROBINS-I tool for non-randomised/observational studies. Risk of bias was assessed by a single reviewer (MLJ) but checked by a second using these tools at the publication, rather than individual outcome level. This made assessing these studies which frequently had multiple study arms, outcome measures and sampling points more manageable - taking advantage of the fact that there was typically some methodological unity between effect sizes reported in the same publication (e.g. one publication relating to a study might report culture-based outcomes whilst another might report culture-independent outcomes from the same samples). Hence, groups of outcomes reported in the same publication - for example, different resistance genes or relative and absolute measures of resistance (see next section) - were therefore considered simultaneously when assessing risk of bias (e.g. baseline differences in the outcome, measurement of the outcome)." | RoB 2 and ROBINS-I were used to assess risk of bias for randomised and non-randomised studies, respectively, as intended. However, we only had capacity for one reviewer to do this independently and another reviewer to check this. The guidance we originally drew up for completing these assessments was broadly followed, except that for found it more practical to negatively answer questions relating to cattle awareness of interventions rather than remove them, and tried to rely on more established Cochrane guidelines where it sufficed. | Readers may have slightly less confidence in our risk of bias assessments given they were conducted by one reviewer and checked by a second. However, even risk of bias assessments conducted independently by two reviewers have been shown to be inconsistent (Bertizzolo et al. 2019; Coon et al. 2024) |
|  | Reason | Plan not possible |  |  |  |  |
|  | Timing | During data collection |  |  |  |  |
| 10 | Type | Effect measures | Too detailed to paste here - see effectsize_calculation.pdf at pre-registration. | "To compare effects of antibiotic administration on antibiotic resistance across studies in a standardised way, we aimed to estimate the standardised mean difference accounting for heteroscedasticity (SMDH). Specifically, we sought to use the SMDH to estimate the difference between the mean absolute abundance of resistant bacteria/resistance genes between bacterial populations associated with beef cattle that had been administered antibiotics (the intervention group), and the mean absolute abundance of resistant bacteria/resistance genes in bacterial populations associated with beef cattle that had not been administered antibiotics, in terms of standard deviations (SDs). The SMDH (Bonett, 2008, 2009) was specifically chosen as the effect measure because - given the comprehensive scope of our synthesis - our eligibility criteria resulted in a set of included studies that used different methods with very different scales of measurement and errors (e.g. culture-based versus qPCR based measures, different target bacterial species, different target resistance genes). In contrast, SMD(H)s are designed to cope with such differences in method and scale of measurement by using the distance in terms of SD as the measure of the effect. SMDHs were never reported in publications themselves, so we sought to calculate them ourselves. The minimal level of data required for this was the means, standard deviations (SDs), and sample sizes of the absolute and relative abundance of resistant bacteria/resistance genes in the bacterial populations sampled, for both the intervention and control/comparator groups in each study for every time point measured in the study." | We had originally intended to use the log response ratio (or 'ratio of means') as the effect measure but this was ill suited to dealing with different methods of measuring the outcome with very different scales of measurements and errors.  In addition, there were sometimes zeros in the measures of resistance which are difficult to deal with using the LRR. Though these were unlikely to be true zeros and more likely samples in which the numbers fell below the detection limit of the resistance assay in question, we did not want to remove these since they likely still reflected true low proportions and their removal would therefore bias the distribution of effect sizes. The other option of adding an arbitrary pseudocount to deal with these cases would also bias our effect size distribution (Koricheva et al. 2013) | Readers may have the impression that we did this to achieve a particular result but we think it is clear that it is not appropriate to combine such different outcomes using the LRR, ignoring differences in the scale of measurement and standard deviations of each measure. Both are in the same category fo effect measure (Differences in Central Tendency). |
|  | Reason | New knowledge |  |  |  |  |
|  | Timing | After results known |  |  |  |  |
| 11 | Type | Synthesis methods | "Outcome measure(s)...The antibiotic resistance tested can be different between studies. Antibiotic resistance can be measured in one of two ways:    1. The prevalence (%) of phenotypic antibiotic resistance in tested isolates of an identified group of bacteria (at least to genus level; e.g. E. coli, Enterococcus, Salmonella, Campylobacter) isolated from a fresh faecal sample. Phenotypic antibiotic resistance can be established using broth dilution, agar dilution, disk diffusion, E-tests, selective plating or automated methods.    2. The prevalence (%) of key AMR genes (AMR genes per 16S genes) in DNA extracted from a fresh faecal sample and tested through real-time PCR/qPCR." | "The minimal level of data required for this [inclusion in the quantitative synthesis] was the means, standard deviations (SDs), and sample sizes of the absolute and relative abundance of resistant bacteria/resistance genes in the bacterial populations sampled, for both the intervention and control/comparator groups in each study for every time point measured in the study." | In line with Deviations 1-4, we specified that must provide a both measure of the effect on absolute and relative abundance of resistance to be included in the quantitative summary and synthesis (meta-analysis). | Minimal impact of this specific decision. Impact is mainly the loss of 3 studies (Zaheer et al., 2013; Muller et al. 2018; Holman et al., 2019) which were excluded because they only reported effects on relative abundance outcomes (see Supporting Information 4). However, we would argue that this minor loss of information is offset by the statistical and interpretability gains from also requiring information on absolute effects (we did contact the authors of these studies to try to obtain absolute data but were unsuccessful in these attempts). |
|  | Reason | New knowledge |  |  |  |  |
|  | Timing | After results known |  |  |  |  |
| 12 | Type | Synthesis methods | "Random Effects  • Within-study time since the start of antibiotic administration  • Unit-level ID (i.e. residual variance)" | "To test our first hypothesis, we fit intercept-only random effects models fit to ‘during’ and ‘after’ data subsets, implemented as multilevel meta-analytical models using the ‘rma.mv’ function in metafor. These models included the following three random effects:  1. Study ID to account for non-independence among effect sizes from the same study. We used the actual study (e.g. trial) rather than publication (i.e. scientific paper) as the study ID, since a few publications reported alternative outcomes from the same study.  2. Time of sampling nested within study arm to account for temporal autocorrelation between repeated measures taken within the same study arm (i.e. an intervention or control/comparator group). This was modelled with a continuous-time autoregressive structure (“struct = ‘CAR’” option in rma.mv). More information about this approach can be found in Viechtbauer (2024), with Stasielowicz (2022) offering an example of its implementation.  3. Unit-level observation ID to account for residual within-study variance. This is to account for any residual variance between multiple effect estimates from the same study, after accounting for the effects of study and repeated sampling within the same study arm (including alternative assessments of the outcome at the same time point within the same study arm). Examples of this approach and its implementation can be found in Konstantopoulos (2011), Credé, Roch and Kieszczynka (2010), Viechtbauer (2022a), Viechtbauer (2007)" | Changed the temporal autocorrelation random effect from time-within-study to time-within study arm, including study as a separate random effect (we chose study rather than publication as there were found to occasionally be multiple publications per study). We also modelled random effect of time within study arm with a continuous-autoregressive structure, which we had not initially been aware was appropriate way to model it (due to uneven spacing of timepoints). | Should improve confidence in results given that repeated measures within a study arm are almost certainly more temporally autocorrelated than across all a study's arms, and it was our original intention to model them as such (we just misspecified this) |
|  | Reason | Typo/Error |  |  |  |  |
|  | Timing | After results known |  |  |  |  |
| 13 | Type | Synthesis methods | "Unexplained variance in the models will be assessed by estimating and reporting total I² and I² for each random effect (Nakagawa & Santos, 2012) and total absolute heterogeneity (Q). If heterogeneity is identified, we will proceed to Stage 2, seeking to explain the heterogeneity using multilevel meta-regressions." | "The presence/absence of heterogeneity in each of the meta-analytical models was primarily assessed using both the Q test (Cochran, 1954). We also quantified heterogeneity using the I^2 statistic using the Jackson, White and Riley (2012) approach for multilevel meta-analytic models, which compares the variance-covariance matrix of the fixed effects of the model with and without random effects. I^2 estimates were interpreted sensu Higgins and Thompson (2002), using the same terminology to describe them when reporting results. Confidence and prediction intervals were also computed within metafor, meaning that prediction intervals were identical to the confidence interval when τ^2 = 0 (Viechtbauer, 2025)." | For simplicitly in an already complicated systematic review and meta-analysis, we only calculated the heterogeneity of the whole model (which was what we intended to base the estimate of heterogeneity on anyway) rather than for each random effect. We also did this using the Jackson et a. (2012)'s pseudo-I2 approach rather than the Nakagawa & Santos (2012) approach, as this was easier to implement for models with heterogeneous variances/inner-outer random effects terms like ours) We also clarified that how we interpreted I2 estimates. | There is some loss of detail on which random effects are driving heterogeneity but this also helps to avoid overwhelming the reader with less critical detail. |
|  | Reason | New knowledge |  |  |  |  |
|  | Timing | After results known |  |  |  |  |
| 14 | Type | Synthesis methods | "Fixed Effects  During:  • Time (days) since the start of antibiotic administration  • Used Daily Dose (UDD) of the antibiotic  • Class of antibiotic administered  After:   • Time (days) since the end of antibiotic administration  • Number of days with antibiotic administration  • Used Daily Dose (UDD) of the antibiotic  • Class of antibiotic administered" | "To test our second hypothesis, we tested the extent to which time of sampling could explain any heterogeneity identified in the intercept-only models. This was done by adding time of sampling as a fixed effect/moderator to the previously intercept-only (random effects) models, making them uni-moderator multilevel (mixed effects) meta-regressions. We tested both linear (‘H2 During Linear’ model & ‘H2 After Linear’ model) and non-linear relationships of time of sampling (‘H2 During Nonlinear’ model & ‘H2 After Nonlinear’ model), with the latter fit as a restricted cubic spline. Given the relatively small number of studies available for meta-analysis, the latter was fit with the minimum of 3 ‘knots’ (the points in the time series where different cubic polynomials are joined), resulting in 2 polynomials/coefficients fit to the effect of time (for implementation see Viechtbauer (2023); for a meta-analytical example see Shim and Lee (2019). " | The fixed effect of time remains as the main (sole) fixed effect we modelled in our meta-regressions (we just specified it with one variable (outcome_time_days) whose meaning becomes time since start/end after splitting the dataset into 'during' and 'after' subsets. We did not specify at protocol design stage how we would model the effect of time, but explored both linear and non-linear forms when conducting our meta-analysis as it was not clear which was more suited. | No impact, especially as we used a linear effect (the default) anyway. |
|  | Reason | Typo/Error |  |  |  |  |
|  | Timing | After data access |  |  |  |  |
| 15 | Type | Synthesis methods | "Fixed Effects  During:  • Time (days) since the start of antibiotic administration  • Used Daily Dose (UDD) of the antibiotic  • Class of antibiotic administered  After:   • Time (days) since the end of antibiotic administration  • Number of days with antibiotic administration  • Used Daily Dose (UDD) of the antibiotic  • Class of antibiotic administered" | "To test our second hypothesis, we tested the extent to which time of sampling could explain any heterogeneity identified in the intercept-only models. This was done by adding time of sampling as a fixed effect/moderator to the previously intercept-only (random effects) models, making them uni-moderator multilevel (mixed effects) meta-regressions. We tested both linear (‘H2 During Linear’ model & ‘H2 After Linear’ model) and non-linear relationships of time of sampling (‘H2 During Nonlinear’ model & ‘H2 After Nonlinear’ model), with the latter fit as a restricted cubic spline. Given the relatively small number of studies available for meta-analysis, the latter was fit with the minimum of 3 ‘knots’ (the points in the time series where different cubic polynomials are joined), resulting in 2 polynomials/coefficients fit to the effect of time (for implementation see Viechtbauer (2023); for a meta-analytical example see Shim and Lee (2019). " | We had originally intended to include used daily dose and antibiotic class as moderator but considering statistical advice to limit the number of moderators tested (Viechtbauer 2007) - especially relevant here where there are few studies relative to the number of potential moderators (overparameterisation) - we decided to avoid this in order to prioritise our primary hypothesis (H2) in the models (investigating the effects of dose and class in subgroup/alternative analyses instead). Morover, dose is highly correlated with days since start of feeding for the during subset, which is another reason not to include it alongside time of sampling. | Should improve confidence in results because eliminates multiple testing in primary model, reduces overparametrisaton, and autocorrelation between moderators. |
|  | Reason | Typo/Error |  |  |  |  |
|  | Timing | After data access |  |  |  |  |
| 16 | Type | Synthesis methods | "Planned Sensitivity analyses: Because phenotypic and genotypic approaches to measuring antibiotic resistance may produce fundamentally different results, we will also rerun the analyses with phenotypic and genotypic outcome measures in separate models." | "We conducted various subgroup analyses (i.e. of categorical moderators) and other alternative analyses (i.e. of non-categorical moderators) to explore alternative hypotheses for heterogeneity among effect sizes. These were conducted on the whole post-antibiotic administration dataset without splitting into ‘during’ and ‘after’ periods to maximise the power of subgroup analyses (representation in each subgroup) and because these analyses were considered as alternative moderators of the effect to the period and time of sampling. The alternative moderators considered fell into PICO(S) groupings and were:  • Population: Reproductive class; breed; weight; age; antibiotic treatment history of the cattle studied.  • Intervention(s) and control(s)/comparator(s): Antibiotic class administered in the intervention; antibiotic dose administered in the intervention (expressed as daily dose; log10-transformed daily dose, cumulative daily dose before each effect estimate, and log10-transformed cumulative daily dose before each effect estimate); whether or not the intervention and control/comparator groups were also administered other (cointervention) antibiotics; whether or not the intervention and control/comparator cattle were housed in the same enclosure.  • Outcome: Microorganisms targeted; method used to measure resistance.  • Study design: The overall design of the study (simplified to observational, completely randomised, or randomised complete block designs)" | As planned, we combined phenotypic (direct selection) and genotypic (qPCR) methods in the primary analyses, but tested for distinct effects in phenotypic and genotypic outcome assessments in a subgroup analysis of all data rather than a sensitivity analysis for each of the during and after subsets. This was because our sensitivity analyses were more focussed on things that could be seen as 'improvements' to our synthesis (and it was not clear that one of the genotypic or phenotypic data was 'better' than the other), and to maximise this power of this analysis by including all data (i.e. both during and after data, as we did for for all subgroup analyses). | No impact - essentially the same analysis is conducted and we considered breaking down the review outcomes at the level of subgroups in the certainty where a subgroup analysis provided strong evidence of subgroup effects. |
|  | Reason | Typo/Error |  |  |  |  |
|  | Timing | After data access |  |  |  |  |
| 17 | Type | Synthesis methods | "Given that some Intervention/exposure and Control groups may receive other antibiotics alongside the intervention antibiotic (e.g. Intervention/Exposure cattle receive tetracycline via injection over 3 consecutive days alongside in-feed monensin, whilst Control cattle only receive in-feed monensin), we will explore separating these ‘More controlled’ and ‘Less controlled’ studies into separate meta-analyses." | "We conducted various subgroup analyses (i.e. of categorical moderators) and other alternative analyses (i.e. of non-categorical moderators) to explore alternative hypotheses for heterogeneity among effect sizes. These were conducted on the whole post-antibiotic administration dataset without splitting into ‘during’ and ‘after’ periods to maximise the power of subgroup analyses (representation in each subgroup) and because these analyses were considered as alternative moderators of the effect to the period and time of sampling. The alternative moderators considered fell into PICO(S) groupings and were:  • Population: Reproductive class; breed; weight; age; antibiotic treatment history of the cattle studied.  • Intervention(s) and control(s)/comparator(s): Antibiotic class administered in the intervention; antibiotic dose administered in the intervention (expressed as daily dose; log10-transformed daily dose, cumulative daily dose before each effect estimate, and log10-transformed cumulative daily dose before each effect estimate); whether or not the intervention and control/comparator groups were also administered other (cointervention) antibiotics; whether or not the intervention and control/comparator cattle were housed in the same enclosure.  • Outcome: Microorganisms targeted; method used to measure resistance.  • Study design: The overall design of the study (simplified to observational, completely randomised, or randomised complete block designs)" | Following our logic of focussing sensitivity analyses on things that could be considered methodological 'improvements' to our synthesis, whilst reserving our subgroup analyses for things more inherent to the included studies with one category not necessarily 'better' than the other, we tested for the effect of cointerventions in a subgroup, rather than sensitivity analysis. | No impact - essentially the same analysis is conducted and we considered breaking down the review outcomes at the level of subgroups in the certainty where a subgroup analysis provided strong evidence of subgroup effects. |
|  | Reason | Typo/Error |  |  |  |  |
|  | Timing | After data access |  |  |  |  |
| 18 | Type | Synthesis methods | "Initially, we will only analyse data from interventions/exposures in which all of the cattle in each replicate are administered/not administered the intervention/exposure antibiotic. However, if there are a substantial number of studies in which only part of the group receive the intervention/exposure antibiotic, we will introduce these data into the models and add ‘proportion of animals treated’ as a moderator." | NA | When we did encounter this (Kanwar et al. 2014; Ohta et al. 2019; Levent et al. 2022) it only manifested as the treatment groups having 1 or 2'control' cattle in the pen alongside a separate control-only pen. Therefore we dealt with this simply excluding the 1 control animal from the pen mean calculations, and doing a subgroup analysis (rather than including proportion treated as a continuous moderator) to test whether the 'contamination' of treatment pens with control animals had any effect. | No impact - essentially the same analysis is conducted and we considered breaking down the review outcomes at the level of subgroups in the certainty where a subgroup analysis provided strong evidence of subgroup effects. |
|  | Reason | Typo/Error |  |  |  |  |
|  | Timing | After data access |  |  |  |  |
| 19 | Type | Synthesis methods | "Given that the number of 16S genes per bacterium can vary from 1-15 (see data_extraction.pdf for more details; Kembel et al., 2012; Sun et al., 2013; Větrovský & Baldrian, 2013), we will explore assuming different numbers of 16S genes in Control vs. Treatment groups when calculating qPCR/real-time PCR resistance prevalence/Log Response Ratio." | NA | We realised this analysis did not make sense because regardless of how many 16S genes you assume, if you assume the same number for each study then the genes/16S value will change uniformly across studies (and hence effect sizes and the meta-analytical results remain unchanged). There is no obvious logic by which number of 16S genes would systematically differ between studies (at least with the information we have) and so we abandoned this analysis. | No impact - this analysis was obviously misguided. |
|  | Reason | Typo/Error |  |  |  |  |
|  | Timing | After data access |  |  |  |  |
| 20 | Type | Synthesis methods | NA | "Given variation in effect sizes can be driven by (meta)analytical decisions (Gould et al., 2025), we subjected our candidate models for hypotheses 1 and 2 to several sensitivity analyses (Supporting Information 7). Briefly, these were: 1) a leave-one-out analysis; 2) removing/correcting potentially problematic studies; 3) assuming different degrees of correlation between effect sizes from the same study (Noble et al., 2017; O’Dea et al., 2019)." | These additional analyses were added either because they are fairly common (e.g. removing non-randomised studies) or in response to issues with the studies we encountered during the process of review. | Should increase confidence in our results as we subject them to additional sensitivity analyses. |
|  | Reason | Typo/Error |  |  |  |  |
|  | Timing | After data access |  |  |  |  |
| **Unregistered Steps** | | | | | | |
| *No.* | *Details* | | *Original wording* | *New wording* | *Deviation Description* | *Reader Impact* |
| 1 | Type | Eligibility Criteria | "Intervention(s)/exposure...Administration of Antibiotic X to an individual or a whole group. Administration of a named pharmaceutical antibiotic drug (“Antibiotic X” e.g. tetracycline) to every animal in an experimental or observational replicate (number of individuals in replicate ≥ 1). The antibiotic administered can be different between studies. The antibiotic may be administered in a single event or multiple events – provided injections occur on consecutive days. The antibiotic can be administered for any described purpose (e.g. experimental, growth promotion, disease prevention, disease treatment); delivered through any route (e.g. in-feed administration, injections); alongside administration of other antibiotics/non-antibiotic antimicrobials/drugs/substances (provided they are the same in intervention/exposure and control groups within the study)." | "Indirect administration of antibiotic contaminants of other media (e.g. the common practice of feeding cattle spent distiller’s grains from the fuel ethanol industry, which can contain virginiamycin as a contaminant) is eligible provided the dose of the contaminant antibiotic is quantified." | Specified how to deal with distillers' grains and similar studies that we had not previously anticipated. | Minimal impact - an extension of our existing eligibility criteria. Arguably the choice to include these studies could bias our synthesis towards no effect given that the doses of contaminating virginiamycin are probably relatively low. However, the one study of distiller's grains we included (Edrington et al. 2014), which included higher dose positive control arms also, was not included in the meta-analysis, and we would have explored dose-response effects in an alternative analyses anyway. |
|  | Timing | Select One |  |  |  |  |
| 2 | Type | Eligibility Criteria | "Outcome measure(s)...The antibiotic resistance tested can be different between studies. Antibiotic resistance can be measured in one of two ways:    1. The prevalence (%) of phenotypic antibiotic resistance in tested isolates of an identified group of bacteria (at least to genus level; e.g. E. coli, Enterococcus, Salmonella, Campylobacter) isolated from a fresh faecal sample. Phenotypic antibiotic resistance can be established using broth dilution, agar dilution, disk diffusion, E-tests, selective plating or automated methods.    2. The prevalence (%) of key AMR genes (AMR genes per 16S genes) in DNA extracted from a fresh faecal sample and tested through real-time PCR/qPCR." | "The bacterial populations in which resistance is assessed should be established bacteria rather than bacteria with which the animal has been experimentally colonised or infected (though it is fine for studies to use experimental colonisation/infection as a cointervention alongside an antibiotic intervention)." | Specified how to deal with experimental infection studies that we had not previously anticipated. | No impact - an extension of our existing eligibility criteria. Arguably measuring resistance in an experimentally infected strain might enable you to detect resistance better, but this relies on achieving high doses of colonisation/infection and is an assessment of resistance in an artificial population. |
|  | Timing | Select One |  |  |  |  |
| 3 | Type | Information sources & search strategy | "Outcome measure(s)...The antibiotic resistance tested can be different between studies. Antibiotic resistance can be measured in one of two ways:    1. The prevalence (%) of phenotypic antibiotic resistance in tested isolates of an identified group of bacteria (at least to genus level; e.g. E. coli, Enterococcus, Salmonella, Campylobacter) isolated from a fresh faecal sample. Phenotypic antibiotic resistance can be established using broth dilution, agar dilution, disk diffusion, E-tests, selective plating or automated methods.    2. The prevalence (%) of key AMR genes (AMR genes per 16S genes) in DNA extracted from a fresh faecal sample and tested through real-time PCR/qPCR." | "The bacterial populations in which resistance is assessed should be established bacteria rather than bacteria with which the animal has been experimentally colonised or infected (though it is fine for studies to use experimental colonisation/infection as a cointervention alongside an antibiotic intervention)." | Specified how to deal with experimental infection studies that we had not previously anticipated. | No impact - an extension of our existing eligibility criteria. Arguably measuring resistance in an experimentally infected strain might enable you to detect resistance better, but this relies on achieving high doses of colonisation/infection and is an assessment of resistance in an artificial population. |
|  | Timing | Select One |  |  |  |  |
| 4 | Type | Synthesis methods | NA | "Outcomes assessed by antimicrobial susceptibility testing were excluded due to statistical issues with using this data to quantitatively estimate the abundance of resistance that only became clear after obtaining study data (Supporting Information 2)." | We excluded outcomes assessed by antimicrobial susceptibility testing from the quantitative summary and synthesis, due to its potential to result in misleading effect sizes. Partly, this was because this method - consisting of isolating a sample of colonies and testing the susceptibility of each to an antibiotic or panel of antibiotics - only permits the estimation of the effect on the relative abundance of resistance, which is interpretatively problematic (see Supplementary Box 1). Moreover, we found that even estimates of relative abundance using this method were likely to be unreliable in these studies, given that the maximum number of isolates tested for each outcome was only 10. Aside from the accuracy issues associated with making any inference based on such an extremely small sample of a typical bacterial population of billions of cells per gram faeces, such sample sizes also result in very imprecise estimates of relative abundance. For example, assuming 95% confidence is needed and a conservative estimate of the true relative abundance of resistance as 0.50/50%, a sample size of 10 is only sufficient to estimate the proportion/percentage of resistant bacteria in the population within a margin of error of 0.32/32% (Manitz et al., 2021). Associated with this lack of precision, the resulting estimates of relative abundance will often be artificially similar or even identical across replicates (e.g. R1=0.1; R2=0.1; R3=0.1), resulting in zero SD in either the control or intervention group, and hence very inflated estimates of the standardised mean difference used to assess effect size. We therefore concluded that outcomes assessed by antimicrobial susceptibility testing did not allow us to answer our research question in the way we had hoped at the protocol design stage, and considered this a justifiable deviation from protocol (Supporting Information 2). | The main impact is the loss of 11 studies that only used antimicrobial suscpetibility testing, which is not an insignificant number. However we think it is clear that these studies do not even provide a reasonable estimate of relative abundance (which was not necessarily their primary intention, to be fair) that would be useful for meta-analysis. Hence we have kept them in the narrative summary only, favouring quality over quantity. |
|  | Timing | Select One |  |  |  |  |
| 5 | Type | Data collection process and data items | NA | Supporting Information 4: "For studies that housed cattle across several pens, we calculated the means and SDs of the log10-transformed data (and sample sizes) at the pen level. This was done by calculating the mean of animals within the same pen (considered pseudoreplicates), before calculating the mean and SD between these pen means (considered replicates and hence sample size considered the number of pens within a study arm). This was to account for dependence between samples/animals from the same pen - mirroring the way the original investigators typically did in their own analyses through averaging within pen or inclusion of pen as a random effect. For the “USMARC - Tylosin” study (Schmidt et al., 2020), using this method also enabled us to correct their reported estimates of the mean, SD, and sample size, which had not previously accounted for pen (for the remaining two studies with such unit-of-analysis errors, we could only correct the sample size as we could not obtain IPD; see below). For studies with the control/comparator and/or treatment/exposure groups housed together in a single pen, we were restricted to calculating means and SDs at the animal level. The potential sensitivity of our results to these less generalisable and conflated designs (Walker, 2018) was tested for implicitly through our Leave-One-Out sensitivity analysis (Quantitative summary and synthesis: Sensitivity analyses), as well as explicitly explored through a dedicated subgroup analysis (Quantitative summary and synthesis: Subgroup and other analyses)" | At protocol design stage, we did not fully anticipate the units of analysis differences between studies - with most studies housing animals across multiple pens and treating pens as the unit of analysis, and some studies treating animals as the unit of analysis (occasionally even where there were multiple pens). Therefore we dealt with this during the analysis by aggregating at the pen level where possible, and using sensitivity analyses to account for where we could not account for pen effects that were present/combining different units of analysis. | Minimal impact. We had to deal with this problem once we encountered it. It could be argued that we lose power by accounting for pen through aggregation if pen effects are not important, especially where the original authors did not do this. However, most studies account for pen so we are following thie consensus, and we could not do so using random effects to produce the means and standard deviations we needed for SMDH calculation. So this is a necessary and conservative addition to the protocol (especially as likely to bias towards the null, if anything). |
|  | Timing | During data collection |  |  |  |  |
| 6 | Type | Data collection process and data items | NA | Supporting Information 4: "Where relative abundance data was not reported in IPD or published figures, we derived it from the data on the absolute abundance of resistance determinants (numerator of relative abundance) and absolute abundance of all determinants (denominator of relative abundance). For IPD, this was done simply by dividing the absolute abundance of resistance determinants by the absolute abundance of all determinants (after unlogging the data). Where data had to be derived from publications and we only had the absolute abundance of resistance determinants (numerator of relative abundance) and total determinants (denominator of relative abundance), we calculated relative abundance from simulated data. For this we used the mean and SD of the log10-transformed data as terms in the ‘rnorm’ function (n=1000) from the base R package v4.5.2 (R Core Team, 2023) to simulate data for the numerator and denominator of relative abundance. The simulated numerator data was then divided by the simulated denominator data to calculate the relative abundance of resistance (after unlogging the data)." | We had not anticipated that we would not be able to directly calculate the mean and standard deviation of the relative abundance - now our secondary outcome - where studies had only reported the mean and standard deviation of the absolute abundance of resistance determinants (numerator) and all determinants (denominator). The only way to do this was to simulate the underlying data. | Readers may have less confidence in these results based on simulated data, but there was no other option and this is now a secondary outcome besides. |
|  | Timing | During data collection |  |  |  |  |
| 7 | Type | Data collection process and data items | NA | Supporting Information 4: "First, we extracted a numeric variable representing the time of sampling as reported in the study, which was simply the number of days since the start of the study that each measurement was taken. This variable (‘outcome_study_days’) was log10-transformed before being used to model temporal autocorrelation between longitudinal measurements within the same study arm via a random effect included in all models." | We did not previously specify that we would log10-transform the time of sampling data as we had not anticipated that it would be log-distributed. | Minimal impact. Readers may perceive that we did this to obtain a particular result for our meta-regressions but it is clear that the time of sampling data are not normally distributed so should be transformed. |
|  | Timing | After data access |  |  |  |  |
| 8 | Type | Data collection process and data items | Too detailed to paste here - see UDD_calculation.pdf at pre-registration. | "Daily antibiotic dose values extracted from publications that were not expressed in estimated milligrams per kilogram of the receiving animal’s body weight (mg/kg body weight) were standardised into this format (MLJ). Dosages expressed as milligram per pound of the receiving animal’s bodyweight were simply multiplied by 2.20462 to give mg/kg body weight. Dosages expressed as grams per tonne of feed/milligram per kilogram of feed, or parts per million were multiplied by 0.02, based on the estimate that cattle eat about 2% of their body weight per day (Rasby, 2013). Dosages expressed as milligram per head were divided by the estimated weight of cattle in each study. Where reported, we used the mean weight of cattle in the study. For studies which did not report the mean weight of cattle, we used the mean weight of cattle across studies that did report it. Once all dosages had been standardised into mg/kg body weight in this way, we also calculated the cumulative dosage administered before each measurement (by multiplying by the cumulative intervention length). We also created log10-transformed versions of the daily and cumulative dosage variables, as it was not entirely clear on which scale dosages were distributed. These four variables were used for a alternative meta-regressions of the effect of dose, rather than time." | We previously specified how we would calculate used daily dose (UDD) but not that we would explore different transformations of this data. | No impact. Appropriate to explore different transformations as distribution less clear cut than time of sampling for example. |
|  | Timing | After data access |  |  |  |  |
| 9 | Type | Synthesis methods | NA | "For studies that housed cattle across several pens, we calculated the means and SDs of the log10-transformed data (and sample sizes) at the pen level. This was done by calculating the mean of animals within the same pen (considered pseudoreplicates), before calculating the mean and SD between these pen means (considered replicates and hence sample size considered the number of pens within a study arm)...Some studies had randomised complete block designs, where cattle were blocked by potentially confounding characteristics such as age and weight before randomisation within each block, to reduce variability not due to the intervention. For these studies, where possible we calculated pen or animal means, SDs, and sample sizes within blocks (essentially treating each block as a sub-experiment)." | At protocol design stage, we did not fully anticipate the units of analysis differences between studies - with most studies housing animals across multiple pens and treating pens as the unit of analysis, and some studies treating animals as the unit of analysis (occasionally even where there were multiple pens). Therefore we dealt with this during the analysis by aggregating at the pen level where possible, and using sensitivity analyses to account for where we could not account for pen effects that were present/combining different units of analysis. | Minimal impact - differences between averaging within pen and using marginal effects as in original papers is typically minimal. Aggregating in this way is the only option to generate means and SDs, since you can only derive SEs for random effects and meta-analysis of marginal means is questionable. |
|  | Reason | New knowledge |  |  |  |  |
|  | Timing | After data access |  |  |  |  |
| 10 | Type | Synthesis methods | NA | "2.8.3 Results of individual studies  Before formally testing our hypotheses with the subset of included in the meta-analysis, the results of all studies included in the systematic review were first included in a narrative synthesis. This synthesis sought to understand the range of effects or associations reported in the relevant literature as a whole, regardless of whether these studies were suitable for inclusion in a meta-analysis. For this, we drew on the whole evidence base including studies using methods that can assess effects on the absolute and relative abundance of resistance (direct plating, qPCR and metagenomics), as well as studies using methods that can only assess effects on the relative abundance of resistance (antimicrobial susceptibility testing). This synthesis consisted of descriptive statistics of study characteristics underpinned by the data extracted into the study characteristics table, though we avoided explicit vote counting given its severe limitations as a synthesis method (Friedman, 2001; Grainger, Stewart and Haddaway, 2022).  To complement the narrative synthesis of the results of individual studies, we also quantitatively synthesised the results of individual studies included in the meta-analysis. For this, we meta-analytically aggregated the effect sizes (SMDHs) at the study level to produce a forest plot sensu Viechtbauer (2022b). We produced separate forest plots for effects measured in faeces and environmental samples, and within these two groupings outcomes measures pre-antibiotic administration (i.e. baseline measurements) or post-antibiotic administration (Supporting Information 6)." | Methods for summarising Results of individual studies via narrative synthesis and forest plots not specified at design stage. | Readers may have slightly less confidence in the objectivity of our summaries of results of individual studies, given it was not pre-specified. However, this initial synthesis is done in a standard way using a Study Characteristics table and discussing the PECO characteristics of the included studies, guided be pre-specified data extraction, as well as using forest plots. |
|  | Timing | During data collection |  |  |  |  |
| 11 | Type | Synthesis methods | NA | "We conducted various subgroup analyses (i.e. of categorical moderators) and other alternative analyses (i.e. of non-categorical moderators) to explore alternative hypotheses for the heterogeneity identified. These were conducted on the whole post-antibiotic administration dataset without splitting into ‘during’ and ‘after’ periods to maximise the power of subgroup analyses (representation in each subgroup) and because these analyses were considered as alternative moderators of the effect to the period and time of sampling." | We did not previously specify any subgroup analyses, conflating sensitivity and subgroup analyses somewhat in our protocol. Some of the sensitivity analyses (and antibiotic class/dose effects) were executed as subgroup/alternative analyses for the reasons described above, whilst others were added as exploratory analyses. We decided to do subgroup analyses on the whole dataset to preserve power, seeing these as alternative hypotheses to H1 and H2. Subgroup analyses fell into PICO categories though we were not able to execute some (e.g. Population ones) due to limited data/reporting. | No impact. Appropriate to explore different transformations as distribution less clear cut than time of sampling for example. |
|  | Timing | After data access |  |  |  |  |
| 12 | Type | Synthesis methods | NA | "Given our hypotheses centered around the effect of antibiotic administration on antibiotic resistance during and after antibiotic administration (see ‘Introduction: Objectives’), we first split post-antibiotic administration data for faecal outcomes into sub-datasets relating to 1) effects measured during antibiotic administration and 2) effects measured after antibiotic administration (defined as at least 1 day since the last antibiotic administration event). Pre-antibiotic administration data were excluded from the synthesis because including them would result in inaccurate estimates of the effect due to antibiotic administration (since antibiotics had not yet been administered for these samples). Data from environmental samples were excluded from the synthesis because their scarcity precluded meta-analysis (see Results)." | Although implied, it was not clear in our pre-registration that we would exclude baseline data from the main analysis, so we clarified this here. We already made clear that we anticipated having to exclude environmental samples from the main synthesis. | Minimal impact. For the 'after' antibiotic administration subset in particular, it could be argued that the loss of baseline data changes the modelled effect of time because by shaving off part of the distribution where the effect sizes lie around 0 at baseline (which might result in a more U-shape relationship with time similar to a the pharmacokinetic relationship of time and antibiotic concentration. However, including baseline data would heavily zero skew the data and contaminate the overall effect in particular with estimates before the intervention has even be administered. It seems more sensible therefore to exclude them and conclude that the relationship is linear within the time range measured (it might still have been U-shaped if there had been measurements available very soon after antibiotic administration, for example). |
|  | Reason | Typo/Error |  |  |  |  |
|  | Timing | After results known |  |  |  |  |
| 12 | Type | Certainty assessment | NA | "We assessed certainty in the size of the overall effect during (“During H1” model), the linear moderating effect of time during (“During H2” model), the overall effect after (“After H1” model), and the linear moderating effect of time after (“After H2” model). We used the standard GRADE approach (Schünemann et al., 2013; Schünemann et al., 2023) to assess certainty separately for each of these four review outcomes from the meta-analysis (Supporting Information 9)." | We did not previously include any methods for Certainty Assessment due to an oversight at protocol design stage. Especially given we had not realised the importance of certainty assessment at protocol design stage and hence not pre-registered methods for it, we tried to keep our implementation of the GRADE approach as simple and aligned with Cochrane guidance as possible (Schünemann et al. 2023a). For example, we used standard definitions for defining the size of the effect and its uncertainty and incorporated our pre-planned risk of bias and publication bias assessments into this framework straightforwardly. | Should improve interpretability especially with Summary of Findings table. As this was not pre-planned, readers may have slightly less confidence in its objectivity but given we assessed the four review outcomes (rather than an unplanned subgroup analysis) and followed guidance this should give readers more confidence. |
|  | Timing | After data access |  |  |  |  |

#### 1.2 Eligibility criteria

Supplementary Information 2: Full eligibility criteria

**Study designs to be included**

Study designs MUST include at least one measurement of antibiotic resistance to the administered antibiotic taken BEFORE the antibiotic course begins, and at least one from DURING or AFTER the antibiotic course. The included study designs are listed below. For the purposes of this systematic review and meta-analysis, all studies – whether experimental or observational - MUST contain a within-study, ‘No administration of Antibiotic X’ control or comparator group.

*Specific definitions of study designs used in the inclusion/exclusion criteria below are those of NICE and available at: https://www.nice.org.uk/process/pmg4/chapter/appendix-d-glossary-of-study-designs#focus-group-study*

**Inclusion criteria:**

- **Randomised controlled trials** – Experimental studies in which the intervention (antibiotic administration) is randomised on an individual animal basis. Individual animals are randomly assigned to the intervention (antibiotic administration) or control (no antibiotic administration) arms. Sometimes this may be done as ‘block randomisation’, where animals are first blocked into groups according potential nuisance variables (e.g. age, weight), and then randomisation to intervention/control is then performed within the group (blocks). The outcome of interest (antibiotic resistance) is then measured before the intervention (antibiotic administration). The intervention (antibiotic administration) then begins and the outcome of interest (antibiotic resistance) is measured at one or more points after it has begun.
- **Cluster randomised controlled trials.** Experimental studies in which randomisation occurs at the level of the group (e.g. a feedlot). Groups are randomly assigned to the replicate intervention (antibiotic administration) and control (no antibiotic administration) arms. Sometimes this may be done as ‘block randomisation’, where groups are first blocked into larger groups according potential nuisance variables (e.g. age, weight), and then randomisation to intervention/control is then performed within groups (blocks). The outcome of interest (antibiotic resistance) is then measured before the intervention (antibiotic administration). The intervention (antibiotic administration) then begins and the outcome of interest (antibiotic resistance) is measured at one or more points after it has begun.
- **Non-randomised controlled designs.** Experimental studies in which there is no random assignment of individuals or clusters to intervention/control arms of cattle before the intervention (antibiotic administration). This may include **non-randomised controlled trials** or **before-and-after** studies in which the investigators assign the experimental units to intervention/control arms non-randomly, or **prospective cohort (longitudinal) studies** in which someone other than the investigators assigns the study units to intervention/control arms non-randomly (e.g. farmers choosing to treat some animals with antibiotics due to the presentation of symptoms.)

**Exclusion criteria:**

- **Retrospective cohort (longitudinal) studies.** Cohort studies that the measure the outcome of interest (antibiotic resistance) in both exposure groups after the exposure (antibiotic administration) has taken place. These studies are excluded because they do not contain a measure before the intervention of interest (antibiotic administration) took place. It is therefore not possible to estimate the relationship between possible cause (antibiotic administration) and effect (antibiotic resistance).
- **Case-control studies.** Retrospective, comparative observational studies in which the investigator selects individuals/groups in which the outcome of interest (antibiotic resistance) has been observed and individuals/groups in which it has not, and then collects data to ascertain what level of the exposure of interest (antibiotic administration) occurred in those individuals/groups. These studies are excluded because they do not contain a measure before the intervention of interest (antibiotic administration) took place. It is therefore not possible to estimate the relationship between possible cause (antibiotic administration) and effect (antibiotic resistance).
- **Interrupted time series.** Studies with more than two observations made on the same individuals or groups, with at least one of the observations being before the intervention (antibiotic administration) event. These studies are excluded because they do not contain a control group.
- **Correlation studies.** Observational studies in which the outcome of interest (antibiotic resistance) is measured at a single point in time after the exposure (antibiotic administration) has taken place. These studies are excluded because they are purely descriptive and do not contain a measure before the intervention of interest (antibiotic administration) took place. It is therefore not possible to estimate the relationship between possible cause (antibiotic administration) and effect (antibiotic resistance).
- **Cross-sectional study.** Observational studies in which the outcome of interest (antibiotic resistance) and/or the exposure of interest (antibiotic administration) is measured at a single point in time after the exposure (antibiotic administration) has taken place. These studies are generally used to determined the prevalence of the outcomes or exposures across the population. These studies are excluded because they are purely descriptive and do not contain a measure before the intervention of interest (antibiotic administration) took place. It is therefore not possible to estimate the relationship between possible cause (antibiotic administration) and effect (antibiotic resistance).
- **Surveys.** Quantitative surveys are the same as cross-sectional studies. Qualitative surveys are excluded because they do not allow for the planned quantitative systematic review/meta-analysis of the outcome of interest (antibiotic resistance) or the relationship between possible cause (antibiotic administration) and effect (antibiotic resistance).
- **Systematic review.** – A study summarising the literature using explicit and systematic methods to identify, appraise and summarise the literature according to predetermined criteria. These are excluded, as they are not experimental or observational studies.
- **Meta-analyses.** A study using statistics to compare the findings of multiple studies. These are excluded, as they are not primary experimental or observational studies.

**Population**

**Inclusion criteria:**

- **Weaned beef cattle**. *Bos taurus* species being raised for meat production during the study (though importantly, this CAN include dairy cattle enrolled for beef production or cattle described as ‘veal’ (since there is no legal definition in some countries e.g. USA). *Bos taurus* also includes the sub-species *Bos taurus indicus* (e.g. Zebu, Brahman). Beef cattle must be weaned i.e. feeding only on solid food, whether grass, legumes, silage, hay, grain, soy, other ingredients, or a mixture. Beef cattle can be grazing cattle (e.g. raised on pasture) or manually fed cattle (e.g. raised in a feedlot).

**Exclusion criteria:**

- **Dairy cattle.** Cattle being raised only for dairy production during the study.
- **Non-cattle**. Any species of animal that is not *Bos taurus*.

**Intervention(s), exposure**

**Inclusion criteria:**

- **Administration of Antibiotic X to an individual or a whole group.** Administration of a named pharmaceutical antibiotic drug (“Antibiotic X” e.g. tetracycline) to every animal in an experimental or observational replicate (number of individuals in replicate ≥ 1). The antibiotic administered can be different between studies. The antibiotic may be administered in a single event or multiple events – provided injections occur on consecutive days. The antibiotic can be administered for any described purpose (e.g. experimental, growth promotion, disease prevention, disease treatment); delivered through any route (e.g. in-feed administration, injections); alongside administration of other antibiotics/non-antibiotic antimicrobials/drugs/substances (provided they are the same in intervention and control groups within the study). Indirect administration of antibiotic contaminants of other media (e.g. the common practice of feeding cattle spent distiller’s grains from the fuel ethanol industry, which can contain virginiamycin as a contaminant) is eligible provided the dose of the contaminant antibiotic is quantified.
- *More controlled example: A group of 10 cattle that all receive tetracycline via injection over 3 consecutive days, and no other antibiotics.*
- *Less controlled example: A group of 10 cattle that all receive tetracycline via injection over 3 consecutive days, alongside in-feed monensin.*
- **Administration of Antibiotic X to part of a group.** Administration of a named pharmaceutical antibiotic drug (“Antibiotic X” e.g. tetracycline) to a subset of the total animals in an experimental or observational replicate (number of individuals in replicate ≥ 1). The antibiotic administered can be different between studies. The antibiotic may be administered in a single event or multiple events – provided injections occur on consecutive days. The antibiotic can be administered for any described purpose (e.g. experimental, growth promotion, disease prevention, disease treatment); delivered through any route (e.g. in-feed administration, injections); alongside administration of other antibiotics/non-antibiotic antimicrobials/drugs/substances (provided they are the same in intervention and control groups within the study). Indirect administration of antibiotic contaminants of other media is again eligible provided dose is quantified.
- *More controlled example: An individual in a group of 10 cattle that receives tetracycline via injection over 3 consecutive days, and no other antibiotics.*
- *Less controlled example: An individual in a group of 10 cattle that receives tetracycline via injection over 3 consecutive days, alongside in-feed monensin.*

**Exclusion criteria:**

- **Unclear whether antibiotics have been administered during the study.** Study arms for which it is not clearly stated whether antibiotics were administered during the study.
- **Undefined course of antibiotics during the study.** Antibiotic administration that is not well described – for example, no information on the antibiotic administered, dose, number or timing of all administration events, number of individuals treated.

**Comparator(s)/control**

**Inclusion criteria:**

- **No administration of Antibiotic X to an individual or a whole group.** These are control individuals or groups in which every animal in an experimental or observational replicate (number of individuals in replicate ≥ 1) does NOT receive Antibiotic X. Administration of other antibiotics/non-antibiotic antimicrobials/drugs/substances is permitted, provided they are the same as those in the intervention group(s) in the study.
- *More controlled example: A group of 10 cattle that receives no antibiotic administration.*
- *Less controlled example: A group of 10 cattle that receives no tetracycline administration, but receives in-feed monensin.*

**Exclusion criteria:**

- **Unclear whether antibiotics have been administered during the study.** Study arms for which it is not clearly stated whether antibiotics were administered during the study.

**Outcome measure(s)**

**Inclusion criteria (faecal outcomes):**

- **Antibiotic resistance to Antibiotic X in fresh cattle faeces.** Antibiotic resistance must be measured for the class of the antibiotic that is administered in the intervention group (“Antibiotic X”) i.e. we are interested only in direct selection. The antibiotic resistance tested can be different between studies. Antibiotic resistance can be measured in one of two ways:
  1. Relative abundance (0-1 if expressed as a proportion or 1-100% if expressed as a percentage) of phenotypic antibiotic resistance in tested isolates of an identified group of bacteria (at least to genus level; e.g. E. coli, Enterococcus, Salmonella, Campylobacter) isolated from a fresh faecal sample. Phenotypic antibiotic resistance can be established using broth dilution, agar dilution, disk diffusion, E-tests, selective plating or automated methods.
  2. Relative abundance (0-1 if expressed as a proportion or 1-100% if expressed as a percentage) of key AMR genes (AMR genes per 16S genes) in DNA extracted from a fresh faecal sample and tested through real-time PCR/qPCR.

The above may be directly reported and the subject of the original analysis or, more commonly, the absolute abundance of resistant and all bacteria in the target population (i.e. the numerators and denominators used to calculate relative abundance) reported separately (e.g. in published figures). Estimates of absolute abundance are typically expressed as cells/genes per mass of sample.

‘Fresh’ faeces means direct samples from cattle rectums (e.g. faecal grab/swab) or faeces that has been deposited on the day of sampling (though these might be refrigerated/frozen/preserved prior to antibiotic resistance testing). Regarding the latter, preferably, defecation is observed directly on the day of sampling. Alternatively, faeces can be attributed to a particular date without observation, provided the enclosure has been cleared/cleaned of faeces within the last 24 hours – for studies that do not clear/clean cattle enclosures daily, this means only samples taken a day after a clearing event has taken place. Where the appropriate experimental replicate is a group (e.g. pen, feedlot), it is acceptable for faeces from different cattle in the group to have been mixed before antibiotic resistance is tested (because in this case, individual cattle are pseudoreplicates) - as an alternative to averaging over individual samples or including pen as a random effect, for example. However, faeces should not be mixed between replicates.

**Inclusion criteria (metagenomic outcomes):**

In addition to, or in absence of, measurement of the primary outcome (bacterial antimicrobial resistance in faeces), a study may be included if it contains metagenomic outcomes. Study design, Population, Intervention, Control, and Outcome criteria other than those pertaining to the method of assessing antimicrobial resistance remain the same. The antibiotic resistance method eligibility criteria to be used in place of those for phenotypic/qPCR methods are:

- **Antibiotic resistance measured via metagenomics.** Metagenomic and sequencing methods provide a powerful way to measure antibiotic resistance genes. These can be included provided 16S genes are quantified in order to estimate relative abundance in a similar way as for qPCR approaches. Metagenomic outcomes will be included in the narrative synthesis only, given that meta-analysing them is beyond the scope of this systematic review for the following reasons:
  1. Metagenomic and sequencing approaches are more suitable for measuring the richness and composition of resistance genes, rather than abundance/prevalence (to which more sensitive and targeted phenotypic and real-time PCR/qPCR approaches are more suited).
  2. Such a meta-analysis requires extracting raw sequence data and combined bioinformatic analysis. Although sequence data is usually available in sequence repositories, the bioinformatics methods used for combined analysis differ substantially from those used in meta-analysis. This data will therefore be collected where it is available and analysed in a separate study if feasible.

**Inclusion criteria (environmental outcomes):**

In addition to, or in absence of, measurement of the primary outcome (bacterial antimicrobial resistance in faeces), a study may be included if it contains environmental sample types. Study design, Population, Intervention, Control, and Outcome criteria other than those pertaining to sample type remain the same. The sample type eligibility criteria to be used in place of those for fresh faeces are:

- **Antibiotic resistance in deposited cattle faeces (environmental).**. Deposited faeces means faeces that left the rectum of cattle over 24 hours ago and can be traced back to one experimental or observational replicate (i.e. not mixed with other replicates). This could be faecal pats, manure or slurry from a replicate individual or group, for example. In order for a sample to be included, there must be detail given on the timing of cleaning and/or clearing of deposited faeces in the cattle enclosure (e.g. in a variable ‘time since cleaning’), as this will greatly affect the environmental prevalence of antibiotic resistant bacteria.
- **Antibiotic resistance in cattle-associated soil (environmental).** Cattle-associated soil means soil or mud that can be traced back to one experimental or observational replicate (i.e. not mixed with other replicates). In order for a sample to be included, there must be detail given on the timing of cleaning and/or clearing of deposited faeces in the cattle enclosure (e.g. in a variable ‘time since cleaning’), as this will greatly affect the environmental prevalence of antibiotic resistant bacteria.
- **Antibiotic resistance in cattle-associated water (environmental).** Cattle-associated water means either:
  1. A static water source (artificial or natural) from within cattle enclosures, that can be traced back to one experimental or observational replicate (i.e. not mixed with other replicates). For example, a cattle trough or pond used for keeping cattle hydrated.
  2. A moving watercourse (artificial or natural) that can be traced back to one experimental or observational replicate (i.e. not mixed with other replicates). For example, wastewater, irrigation canal or river water that is downstream of only one replicate. This is most likely to be the case where a whole beef production facility is the unit of replication and each facility has a unique downstream flow (because it is difficult to attribute downstream water to one replicate where there are multiple replicates on one site).
- **Antibiotic resistance in cattle-associated air (environmental).** Cattle-associated air can either be:
  1. Air (indoor or outdoor) from within cattle enclosures that can be traced back to one experimental or observational replicate (i.e. not mixed with other replicates). For example, air from inside a cattle shed or an outdoor cattle pen.
  2. Air from outside of cattle enclosures (artificial or natural) that can be traced back to one experimental or observational replicate (i.e. not mixed with other replicates). For example, air downwind of a field or feedyard. This is most likely to be the case where a whole beef production facility is the unit of replication and each facility has a unique downwind flow (because it is difficult to attribute downwind air to one replicate where there are multiple replicates on one site).
- In order for a sample to be included, there must be detail given on the timing of cleaning and/or clearing of deposited faeces in the cattle enclosure (e.g. in a variable ‘time since cleaning’), as this will greatly affect the environmental prevalence of antibiotic resistant bacteria.

**Exclusion criteria (all outcomes):**

- **Microbiological analyses that cannot detect antibiotic resistance.**
- **Number of experimental or observation replicates (individuals/groups) per intervention/control groups not indicated.** This is necessary to calculate the effect size. Note that this refers to the number of replicate individuals/groups of cattle, rather than pseudoreplicates such as the number of faecal samples tested per individual/group or the number of bacterial isolates tested per faecal sample (though these will be considered in Risk of Bias assessment)
- **Minimum inhibitory concentration(s) and/or resistance cut-off concentration used not indicated (phenotypic).** Phenotypic tests of antibiotic resistance for which the minimum inhibitory concentration of the antibiotic and/or the threshold used to classify something as ‘resistant’ are not reported (e.g. CLSI or EUCAST breakpoint, ECOFF cut-off value).
- **Mean and standard deviation of the relative abundance of resistance per intervention/control group not reported or able to be calculated (phenotypic).** This is necessary to combine with the number of replicates per intervention/control groups in order to calculate the effect size (see Analysis Plan). These would be phenotypic tests of the relative abundance of antibiotic resistance for which it is not possible to calculate the mean and standard deviation of the percentage of tested bacterial isolates that were found to be resistant in the sample. This must be either directly reported (e.g. in text/table/figure) or possible to calculate from data provided in the main/supplementary materials/upon request.
- **Mean and standard deviation of the prevalence (%) of resistance per intervention/control group not reported or able to be calculated (genotypic).** This is combined with the number of replicates per intervention/control groups in order to calculate the effect size (see Analysis Plan). These would be genotypic tests of the relative abundance of antibiotic resistance for which it is not possible to calculate the mean and standard deviation of the number of antibiotic resistance genes per 16S genes in the sample. This must be either directly reported (e.g. in text/table/figure) or possible to calculate from data provided in the main/supplementary materials/upon request.
- **Resistance in non-bacterial microorganisms** (e.g. archaea, fungi, protists, viruses)
- **Resistance in non-microorganisms** (e.g. helminths, ticks)
- **Resistance that is only specific to non-antibiotics** (e.g. resistance genes that are specific to antifungals or antivirals, detergents, metals)

#### 1.3 Information sources & search strategy

Supplementary Information 3: Search strategy

**Main searches conducted via University of Exeter**

| **Database** | **July.Sept.2020** | **Nov.2022** | **Mar.2024** | **Apr.2025** |
| --- | --- | --- | --- | --- |
| Agricola | x |  |  |  |
| CAB Abstracts | x | x | x | x |
| Environment Complete | x |  |  |  |
| Explore | x |  |  |  |
| Global Health | x |  |  |  |
| Google Scholar | x |  |  |  |
| Lilacs | x |  |  |  |
| Medline | x | x |  | x |
| PQDT | x |  |  |  |
| Scopus | x |  |  |  |
| VetMed | x |  |  |  |
| Web of Science | x | x | x | x |

*Agricola*

Keyword Anywhere(cattle or cow or cows or bovine or steer or steers or calf or calves) AND Keyword Anywhere(resistance or multiresistance) AND Keyword Anywhere(feed or infeed or diet or faeces or feces or feedlot)(DATE=2010-2020)

*CAB Abstracts via Ovid*

```
1     (cattle* or cow or cows or bovine* or steer or steers or calf or calves).tw. (625127)
2     (cattle* or cow or cows or bovine* or steer or steers or calf or calves).ti. (313168)
3     exp cattle/ (513764)
4     (feed* or infeed* or diet* or faec* or fec*).tw. (1175841)
5     exp feedlots/ (3989)
6     ((bact* or microb* or antibiotic* or antimicrob* or drug* or multidrug*) adj3 (resist* or multiresist*)).tw. (93095)
7     1 or 3 (626000)
8     4 or 5 (1175841)
9     6 and 7 and 8 (1846)
10     (resist* or multiresist*).ti. (189918)
11     2 and 10 (3261)
12     (bact* or microb* or antibiotic* or antimicrob* or drug* or multidrug*).tw. (1646685)
13     11 and 12 (2103)
14     9 or 13 (3455)
```

*Environment Complete via EBSCO*

Search 1

TI(cattle* OR cow or cows OR bovine* OR steer or steers or calf or calves)
OR
AB(cattle* OR cow or cows OR bovine* OR steer or steers or calf or calves)
OR
DE "CATTLE" OR DE "COWS"

AND
TI(feed* or infeed* or diet* or faec* or fec*)
OR
AB(feed* or infeed* or diet* or faec* or fec*)

AND
TI(bact* or microb* or antibiotic* or antimicrob* or drug*) N3 resist*
OR
AB(bact* or microb* or antibiotic* or antimicrob* or drug*) N3 resist*
OR
TI(bact* or microb* or antibiotic* or antimicrob* or drug*) N3 multiresist*
OR
AB(bact* or microb* or antibiotic* or antimicrob* or drug*) N3 multiresist*

Search 2

TI(cattle* OR cow or cows OR bovine* OR steer or steers or calf or calves)
AND
TI(resist* or multiresist*)

*Explore via the British Library (restricted to reports or PhD theses)*

resist* OR multiresist
AND
cattle* OR cow OR cows OR bovine* OR steer OR steers OR calf OR calves
AND
feed* OR infeed* OR diet* OR faec* OR fec*

*Global Health (via Ovid)*

```
1     (cattle* or cow or cows or bovine* or steer or steers or calf or calves).tw. (91583)
2     cattle/ (57869)
3     1 or 2 (91583)
4     (feed* or infeed* or diet* or faec* or fec*).tw. (523367)
5     feedlots/ (646)
6     4 or 5 (523367)
7     (bact* or microb* or antibiotic* or antimicrob*).tw. (702643)
8     (resist* or multiresist*).tw. (258326)
9     3 and 6 and 7 and 8 (1051)
10     (cattle* or cow or cows or bovine* or steer or steers or calf or calves).ti. (30612)
11     (resist* or multiresist*).ti. (88938)
12     10 and 11 (670)
13     9 or 12 (1538)
```

*Google Scholar via Publish or Perish (restricted to 1000 references)*

Cattle and feedlots in title = 238 hits
Cattle and feedlots in keywords = 940 hits

Both results were downloaded and duplicates removed leaving 884 which were imported into main EndNote library

*LILACS*

cattle or cow or cows or bovine or steer or steers or calf or calves
AND
resistance or multiresistance

*MEDLINE (via Ovid)*

```
1     (cattle* or cow or cows or bovine* or steer or steers or calf or calves).tw. (348341)
2     (cattle* or cow or cows or bovine* or steer or steers or calf or calves).ti. (169018)
3     exp cattle/ (348018)
4     (feed* or infeed* or diet* or faec* or fec*).tw. (1059436)
5     ((bact* or microb* or antibiotic* or antimicrob* or drug* or multidrug*) adj3 (resist* or multiresist*)).tw. (218870)
6     1 or 3 (470194)
7     (resist* or multiresist*).ti. (313099)
8     2 and 7 (2026)
9     exp Drug Resistance, Bacterial/ (89779)
10     5 or 9 (265474)
11     4 and 6 and 10 (1103)
12     8 or 11 (2814)
```

*ProQuest Dissertations and Theses Global*

(cattle* or cow or cows or bovine* or steer or steers or calf or calves).ti.
OR
(cattle* or cow or cows or bovine* or steer or steers or calf or calves).ab
AND
(feed* or infeed* or diet* or faec* or fec*).ti .
OR
(feed* or infeed* or diet* or faec* or fec*).ab
AND
(resist* OR multiresist*).tw

*SCOPUS*

Search 1

(cattle* or cow or cows or bovine* or steer or steers or calf or calves).ti.
OR
(cattle* or cow or cows or bovine* or steer or steers or calf or calves).ab
AND
(feed* or infeed* or diet* or faec* or fec*).ti .
OR
(feed* or infeed* or diet* or faec* or fec*).ab
AND
((bact* or microb* or antibiotic* or antimicrob* or drug* or multidrug*) w/3 (resist*).ti
OR
((bact* or microb* or antibiotic* or antimicrob* or drug* or multidrug*) w/3 (multiresist*).ti
OR
((bact* or microb* or antibiotic* or antimicrob* or drug* or multidrug*) w/3 (resist*).ab
OR
((bact* or microb* or antibiotic* or antimicrob* or drug* or multidrug*) w/3 (multiresist*).ab

Search 2

(cattle* or cow or cows or bovine* or steer or steers or calf or calves).ti.
AND
(resist* OR multiresist*).ti
AND
(bact* or microb* or antibiotic* or antimicrob* or drug* or multidrug*).ti
OR
(bact* or microb* or antibiotic* or antimicrob* or drug* or multidrug*).ab

*VetMed via CABI (with thanks to external colleague at Cornell University Library)*

title:(cattle* or cow or cows or bovine* or steer or steers or calf or calves)
AND
title:(resist* or multiresist*)
AND
title:(bact* or microb* or antibiotic* or antimicrob* or drug* or multidrug*)

*Web of Science*

Search 1

```
TI=(cattle* OR cow or cows OR bovine* OR steer or steers or calf or calves)
AB=(cattle* OR cow or cows OR bovine* OR steer or steers or calf or calves)
KP=(cattle* OR cow or cows OR bovine* OR steer or steers or calf or calves)
AND
TI=(feed* or infeed* or diet* or faec* or fec*)
AB=(feed* or infeed* or diet* or faec* or fec*)
KP=(feed* or infeed* or diet* or faec* or fec*)
AND
TI=(bact* or  microb*  or  antibiotic*  or  antimicrob* or drug*) NEAR/3 resist*
AB=(bact* or  microb*  or  antibiotic*  or  antimicrob* or drug*) NEAR/3 resist*
KP=(bact* or  microb*  or  antibiotic*  or  antimicrob* or drug*) NEAR/3 resist*
OR
TI=(bact* or  microb*  or  antibiotic*  or  antimicrob* or drug*) NEAR/3 multiresist*
AB=(bact* or  microb*  or  antibiotic*  or  antimicrob* or drug*) NEAR/3 multiresist*
KP=(bact* or  microb*  or  antibiotic*  or  antimicrob* or drug*) NEAR/3 multiresist*
```

Search 2

TI=(cattle* OR cow or cows OR bovine* OR steer or steers or calf or calves)
AND
TI=resist* or multiresist*

**Latin American searches conducted by INTA**

| **Database** | **Date** |
| --- | --- |
| LILACS | 22 Feb 2021 |
| Google Académico | Mar 2021 |
| La Referencia | 22 Feb 2021 |
| SIDALC | 22 Feb 2021 |
| REDIB | 22 Feb 2021 |
| SciELO | Apr 2021 |
| Redalyc | Mar 2021 |
| SNRD (INTA institutional repository) | 22 Feb 2021 |

*LILACS*

Subject: Health (: Medicine, Public Health, Dentistry, Nursing, Veterinary Medicine, Sanitary Engineering, Pharmacy and Chemistry, Biology, Nutrition, Psychology, Ecology and Environment, etc.)

Geographic coverage: Latin America and the Caribbean

Chronological coverage: since 1982

Language coverage: Spanish, Portuguese, English (and French): the documents are indexed and retrieved by any of the three languages. The documents that are available in several languages will be indexed individually in each of them (duplication of the same document).

Language used: controlled: DeCS thesaurus - Descriptors in Health Sciences (MeSH translation):

Content coverage: general, includes meeting papers, theses, technical reports and health legislation

Search 1 (461 hits)

resistencia
AND
ganado vacuno

Search 2 (62 hits)

resistencia
AND
ganado vacuno
AND
antibioticos

Search 3 (55 hits)

resistencia
AND
ganado vacuno
AND
antimicrobianos

All 578 records were downloaded, resulting in **578 records** for double-screening.

*Google Académico (via Publish or Perish)*

Search 1 - Title (22 hits)

Bovino OR vacuno AND feedlot

resistencia AND feedlot AND antibiotic = > 1000 hits (first 1000 downloaded)
resistencia AND feedlot AND antimicrobiano = 289 hits (first 1000 downloaded)

Search 2 - Keywords (>1000 hits - 1000 is the retrieval limit of Publish or Perish)

Bovino OR vacuno AND feedlot

Search 3 - Keywords (>1000 hits - 1000 is the retrieval limit of Publish or Perish)

resistencia AND feedlot AND antibiotic

Search 4 - Keywords (289 hits)

resistencia AND feedlot AND antimicrobiano

All 2311 results were downloaded, books and and duplicates removed. AP then individually screened out articles clearly not related to the topic, resulting in **231 records** for double-screening.

*La Referencia*

Geographical scope: Latin America and Spain Language: Spanish, English and Portuguese Language used: natural and controlled language Note: The database extends the search to similar words or in another language.

Search (168 hits):

All fields (subject, title and table of contents ) : antimicrobiano OR antibiotico AND
All fields (subject, title and table of contents ): bovino OR vacuno AND
All fields (subject, title and table of contents ): resistencia

All 168 records were downloaded, resulting in **168 records** for double-screening.

*SIDALC*

Limited to databases accessible within Argentina (but not filtered to Argentinian publications). Thesaurus used: AGROVOC

Terms used:

- GANADO
- BOVINO
- BOVINOS
- SUSCEPTIBILIDAD
- ANTIMICROBIANA
- ANTIBIOTICOS
- ANTIMICROBIANOS
- RESISTENCIA A LA ENFERMEDAD
- RESISTENCIA A MEDICAMENTOS
- RESISTENCIA A LOS ANTIBIOTICOS
- RESISTENCIA A MEDICAMENTOS

Search 1 (47 hits):

(All Fields:BOVINOS AND All Fields:ANTIMICROBIANOS AND All Fields:RESISTENCIA)"

Search 2 (42 hits):

(All Fields:BOVINOS AND All Fields:ANTIBIOTICOS AND All Fields:RESISTENCIA)"

All results were downloaded and duplicates removed, resulting in **87 records** for screening.

*REDIB*

Geographic scope: Ibero-American Search language used: natural language, keywords (not thesaurus)

All fields: resist* AND
All fields: bovin* OR vacun* AND
All fields: antimicrob* OR antibiot*)

All 85 records were downloaded, resulting in **85 records** for double-screening.

*SCIELO*

Subject: general science Chronological coverage: since 1995 to present Geographic coverage: Latin America and the Caribbean, Spain and Africa Language coverage: Spanish, Portuguese, English

Search (43 hits):

(resisten*) AND (antimicrobian*) OR (antibiotic*) AND (bovino) OR (vacuno)

All 43 records were downloaded, resulting in **43 records** for double-screening.

*Sistema Nacional de Repositorios Digitales (SNRD)*

https://repositoriosdigitales.mincyt.gob.ar/

Affiliated repositories at time of search: 44 (instituciones) Temática: General sciences Search language used: natural language and controlled language (thesaurus used by different institutions) Language used: The documents are indexed and retrieved in both Spanish and English, interchangeably.

Search (29 hits)

Todos los Campos: antimicrobiano OR antibiotico AND
Todos los Campos: vacuno OR bovino AND
Todos los Campos: resistencia

All 29 records were downloaded, resulting in **29 records** for double-screening.

*Redalyc*

Publications that include: articles with reference Geographical scope: Latin America and the Caribbean, Spain and Portugal Languages included in the search: Spanish, English and Portuguese

Search 1 (134 hits)

resistencia AND antimicrobianos

Search 1 (4 hits)

resistencia AND antibacterianos

All results were downloaded, books and and duplicates removed. AP then individually screened out articles that do not refer to cattle, resulting in **6 records** for double-screening.

**Search updates**

A SST was created in December 2021 with the 34 initially included papers from the searches carried out in July 2020, and the forward and backward citation searching initially carried out in December 2021. 33/34 articles were retrieved from the database searching with one additional article included from forward citation searching. No database retrieved unique articles but Medline retrieved 32, Web of Science 30 and CAB Abstracts 29 so it was decided to search only these databases in any update searches. The searches carried out via INTA retrieved no further included articles so it was decided not to search these sources for any update searches.

**Citation searches**

We initially performed citation searches using Web of Science, SCOPUS, Google Scholar, and Citationchaser^5^ in December 2021. However, when we compared the new included studies emerging from these citation searches to those emerging from the first search updates subsequently performed in November 2022, we found that citation searching did not find articles that could not have otherwise been found by a search update. This was likely because our search strategy was broad and the included studies were all peer-reviewed articles published in well-indexed scientific journals. We therefore did not conduct citation searches on the final set of included studies, since they appeared to perform the same function as search updates, which were simpler to run and more efficient (i.e. we were able to capture the same included studies whilst running fewer searches and screening fewer records).

##Data collection process and data items

Supplementary Information 4: Extracted data

**Study characteristics table (simplified)**

- **Study no:** Arbitrary number assigned to each study (e.g. randomised controlled trial)
- **Study (code)**: Shorthand/computer-readable name assigned to each study in the format ‘Institution_Interventions’. Different antibiotic interventions in studies with multiple are separated by camel case.
- **Study (Research institute):** Longhand/human readable name assigned to each study in the format ‘Institution - Interventions’. Different antibiotic interventions in studies with multiple are separated by semicolons.
- **Study design:** Design name (randomised block, completely randomised design, or non-randomised design); number of animals in intervention/control group(s) and number of pens in which they were housed (pens are typically considered the unit of analysis and are for the meta-analysis where possible, though this is not true for every study’s analysis or extracted means).
- **Population:** Type of production system and reproductive class (e.g. feedlot steers, unspecified calves/cows/heifers/bulls); approximate age at enrollment (in months)
- **Eligible intervention(s):** Antibiotic interventionss from the study that are eligible for inclusion in the systematic review, in the form ‘Antibiotic (dose)’ with each antibiotic intervention separated with a semicolon. Dose is reported in mg/kg, converted from other formats or estimated (“~”) based on approximate consumption for certain in-feed antibiotics.
- **Eligible control(s)**: Antibiotics administered in the control group(s) that are eligible for comparison with the intervention group(s), if any. Typically, but not always, these are in-feed antibiotics such as tylosin and monensin, where given.
- **Publication no.:** Arbitrary number assigned to each publication (e.g. randomised controlled trial). Among the publications included in this review, there are sometimes multiple publications reporting different outcomes from the same study (e.g. trial).
- **Publication(s):** Publication in which the outcome matrix/eligible outcomes were reported. Separate publications related to the same study are reported in sub-rows of the study row.
- **Outcome matrix:** The matrix from which the target bacteria or their DNA were isolated (e.g. fresh faeces, pen surface soil, feed, water).
- **Eligible outcome(s):** Outcome methods and target organisms that are eligible for inclusion in the systematic review. For direct enumeration, the
- **Reported effect(s) - Enrichment in absolute abundance (per mass of faecal matrix) of faecal resistance determinants in intervention relative to control:** Narrative descriptive of the effect of the intervention(s) on the absolute abundance of resistant bacteria/resistance genes related to the antibiotic used in that intervention(s), relative to the control(s). Temporal trends are reported approximately.
- **Reported effect(s) - Enrichment in relative abundance (per population of bacteria) of resistance determinants in intervention relative to control:** Narrative descriptive of the effect of the intervention(s) on the relative of resistant bacteria/resistance genes related to the antibiotic used in that intervention(s), relative to the control(s). Temporal trends are reported approximately.

**Study characteristics table (full)**

- **Originalorder:** Original order in which entered
- **Second_reviewer:** Name of the second reviewer who checked the data extraction
- **Search_source:** Search from which this publication/study is derived.
- **Study_no:** Arbitrary number assigned to each study (e.g. randomised controlled trial)
- **Study**: Shorthand/computer-readable name assigned to each study in the format ‘Institution_Interventions’. Different antibiotic interventions in studies with multiple are separated by camel case.
- **Study_site:** Site/station and institution where the study was conducted.
- **Study_year:** Actual or approximate year in which study was conducted.
- **Study_year_actualorfirstpublication:** Whether ‘Study_year’ is the actual year (reported in a publication) or the year of the first publication (where actual year is not reported).
- **Location:** Location of the study in form ‘City, State/Municipality, Country’
- **Study design:** Design name (randomised block, completely randomised design, or non-randomised design); number of animals in intervention/control group(s) and number of pens in which they were housed (pens are typically considered the unit of analysis and are for the meta-analysis where possible, though this is not true for every study’s analysis or extracted means).
- **Blocking:** Factors by which the animals were blocked before randomisation, if any.
- **Accounted for block in analysis?:** Text denoting whether or not block was accounted for in the original analysis reported in the publication(s).
- **Correct unit of analysis (housing):** What the correct unit of analysis in terms of housing is in the study. Where intervention animals are all housed in a single pen (with or without the control animals), this is ‘animal’ since there is no replication. Where intervention animals are housed across multiple pens, this is ‘Enclosure (pen)’ since pen-level dependencies between animals should be accounted for. Where there is one animal per pen, this is ‘Animal/Enclosure (pen)’ since the choice is equivalent.
- **Accounted for housing in analysis?:** Text denoting whether or not housing was accounted for in the original analysis reported in the publication(s).
- **Production system:** Production system in which the study takes place/that the research station is simulating.
- **Population:** Type of production system and reproductive class (e.g. feedlot steers, unspecified calves/cows/heifers/bulls); approximate age at enrollment (in months)
- **Intervention:** Antibiotic interventions from the study, including both those that are eligible for inclusion in the systematic review and those that are not (see ‘Reason for excluding from meta-analysis (study arm)’ column). Interventions are described in the form: Antibiotic (brand); labelled dose (approximate dose in mg/kg body weight); number of continuous daily doses administered over course of the intervention; mode of administration
- **Cointerventions/Control:** Antibiotics administered in the control group(s) (if any) that are also given to the intervention/exposure group(s) in addition to the intervention antibiotic (cointerventions and control groups are therefore equivalent as per our eligibility criteria). Typically, but not always, these are in-feed antibiotics such as tylosin and monensin, where given.. Cointerventions are described in the form: Antibiotic (brand); labelled dose (approximate dose in mg/kg body weight); number of continuous daily doses administered over course of the intervention; mode of administration.
- **Other cointerventions:** Other, non-antibiotic cointerventions administered to both the cointervention/control group(s) and the intervention/exposure group(s). For example, an additional probiotic or mineral supplement that distinguishes a study arm.
- **Intervention replicates:** Number of replicates in the intervention arm, with the unit of replication (animal or enclosure) defined.
- **Control replicates:** Number of replicates in the control arm, with the unit of replication (animal or enclosure) defined.
- **Publication no.:** Arbitrary number assigned to each publication (e.g. randomised controlled trial). Among the publications included in this review, there are sometimes multiple publications reporting different outcomes from the same study (e.g. trial).
- **Publication(s):** Publication in which the outcome matrix/eligible outcomes were reported. Separate publications related to the same study are reported in sub-rows of the study row.
- **Outcome (as reported in publication):** Antibiotic resistance determinants (target bacteria for culture-based methods and target resistance genes for culture-independent methods) defining the intervention, the form in which they were reported (relative/absolute abundance), the matrix from which they were isolated (e.g. fresh faeces, pen surface soil, feed, water), and the specific method used to assess resistance.
- **Sampling regimen:** Number of days over which longitudinal sampling took place, samples taken over this period with respect to the antibiotic administration regime.
- **Intervention trend (as reported in publication)**: Narrative descriptive of trend in the reported outcome in the intervention(s) group represented by this entry.
- **Control trend (as reported in publication)**: Narrative descriptive of trend in the reported outcome in the control group represented by this entry.
- **Reported effect(s) - Enrichment in absolute abundance (per mass of faecal matrix) of faecal resistance determinants in intervention relative to control:**
- **Reported effect(s) - Enrichment in relative abundance (per population of bacteria) of resistance determinants in intervention relative to control:** Narrative descriptive of the effect of the intervention(s) on the relative of resistant bacteria/resistance genes related to the antibiotic used in that intervention(s), relative to the control(s). Temporal trends are reported approximately.
- **Reason for excluding from meta-analysis (study):** Reason for excluding this study from the meta-analysis, if applicable. There may be multiple reasons, with the primary one listed first.
- **Reason for excluding from meta-analysis (study arm):** Reason for excluding this study arm from the meta-analysis, if applicable. There may be multiple reasons, with the primary one listed first.

#### 1.4 Synthesis methods

##### 1.4.1 Preparation of data

Supplementary Information 5: Methods used to prepare the data for quantitative synthesis

**Outcome data extraction and standardisation**

*Individual participant data*

To calculate the effect size for each effect combination in each study, the minimal level of data required was the mean, SD, and sample size of the absolute abundance of resistance determinants (i.e. density of resistant bacteria belonging to the taxonomic group targeted, density of resistance genes; the numerator for calculating relative abundance) and absolute abundance of all determinants (i.e. total density of bacteria belonging to the taxonomic group targeted, density of 16S genes; denominator for calculating relative abundance). The latter was required alongside the former to conduct complementary analyses related to relative effects (Supplementary Box 1). The means and SDs were required on a log scale since microbiological count data tends to be log10-normally distributed.

For studies that housed cattle across several pens, we calculated the means and SDs of the log10-transformed data (and sample sizes) at the pen level. This was done by calculating the mean absolute abundance of resistance determinants across animals within the same pen (considered pseudoreplicates), before calculating the mean and SD across these pen means (considered replicates and hence sample size considered the number of pens within a study arm). This was to account for dependence between samples/animals from the same pen — mirroring the way the original analyses had accounted for psuedoreplication (through averaging within pen or including pen as a random effect), or accounting for it where it had not. We were able to achieve the latter for the “USMARC - Tylosin” study^6^, for which we were able to use IPD to calculate estimates of the mean, SD and sample size accounting for psuedoreplication, even though the originally-reported estimates had not. For the remaining two studies with this pseudoreplication issue, we could only correct the sample size as we could not obtain their respective IPD (see below). For studies with the control and/or treatment groups housed together in a single pen, we were restricted to calculating means and SDs at the animal level. The potential sensitivity of our results to these less generalisable and conflated designs^7^ was tested for implicitly through our Leave-One-Out sensitivity analysis (see below), as well as explicitly explored through a dedicated subgroup analysis (Methods: Synthesis Methods: Subgroup and other alternative analyses)

Some studies had randomised complete block designs, where cattle were blocked by potentially confounding characteristics such as age and weight before randomisation within each block, to reduce variability not due to the intervention. For these studies, where possible we calculated pen or animal means, SDs, and sample sizes within blocks (essentially treating each block as a sub-experiment). However, only two of these six studies - Texas A&M_TylTylprobiotic^8^ and Texas A&M_probiotic^9^ - provided enough for this approach. Furthermore, for “Texas A&M - Tylosin with probiotic”^9^, we could not use this information to calculate pen SDs within block, since there was no replication of study arms within blocks (i.e. only a single pen per study arm per block). Hence, we could only account for blocking for one of the six randomised complete block studies, and thus tested the sensitivity of our results to this (see below).

*Data derived from published figures*

For the two studies for which study data were only available in the form of published figures - “Texas A&M - Ceftiofur”^10^ and “Lethbridge - Tylosin”^11^ - MLJ manually extracted log10-transformed means and error bar limits from these figures using WebPlotDigitizer version 5.2^12^. Extracted means required no conversion. Extracted error bar limits were used to calculate SD in R by calculating the distance between the mean and the error bar limit where the error bar already represented SD, and converting this to SD by multiplying by the square root of the sample size (see below) where it represented standard error (SE). In one (1/97) case, due to the small values and the slight imprecision inherent in extracting data from a figure, the extracted values fell slightly below zero. These were converted to zero, since negative values are not possible for this type of count data.

The number of pens, rather than number of animals in each study arm, was treated as the sample size for these two studies. This decision was made because in one study the sample size was unclear^10^, whilst in the other the number of animals appeared to have been incorrectly reported as the sample size^11^. Using the number of pens therefore avoided overweighting these two studies in the meta-analysis, partially correcting potential unit-of-analysis errors (i.e. pseudoreplication). The potentially underestimated SDs associated with unit-of-analysis errors^7^ could not be corrected, but the sensitivity of our results to these was tested in a sensitivity analysis (Quantitative summary and synthesis: Subgroup and other analyses).

*Relative abundance (secondary outcome) data*

Means, SDs, and sample sizes for relative abundance were calculated or extracted from published figures in the same way as for absolute abundance described above, with additional pre-processing.

Where relative abundance data was not reported in IPD or published figures, we derived it from the data on the absolute abundance of resistance determinants (numerator of relative abundance) and absolute abundance of all determinants (denominator of relative abundance). For IPD, this was done simply by dividing the absolute abundance of resistance determinants by the absolute abundance of all determinants (after back-transforming the log-transformed data). Where data had to be derived from publications and we only had the absolute abundance of resistance determinants (numerator of relative abundance) and total determinants (denominator of relative abundance), we calculated relative abundance from simulated data. For this we used the mean and SD of the log10-transformed data as terms in the ‘rnorm’ function (n=1000) from the base R package v4.5.2^13^ to simulate data for the numerator and denominator of relative abundance. The simulated numerator data was then divided by the simulated denominator data to calculate the relative abundance of resistance (after back-transforming the log-transformed data).

Finally, we attempted to normalise the relative abundance (proportion) data before the calculation of means and SDs through logit and arcsine-square root transformation. This was because, as is common with proportion data^14^, the relative abundance data extracted deviated strongly from normality due to very low relative abundances of some determinants (positive skew), leading to means and SDs that do not reflect well the central tendency of the data well and hence effect sizes derived from them may be less informative. Logit transformation was performed using the ‘logit’ function from the car R package v3.1.5^15^ using 0.025 as the truncation factor. Arcsine-square root transformation was performed using the ‘transf.asin’ function of the metafor R package v4.8.0^16^. In practice these transformations had suboptimal success in normalising the data, and because of this and other reported limitations of these transformations^17,18^, we performed sensitivity analyses by testing effect sizes derived from logit-transformed and arcsine-square root transformed data.

Alongside the issues of interpretability associated with relative effects, these various statistical issues associated with the relative abundance data also contributed to our decision to refocus our synthesis on absolute abundance. Nonetheless, for reasons of transparency, relative abundance was kept as a secondary outcome in our initial narrative and quantitative summaries of effects despite these issues (Supplementary Box 1).

*Calculation of the SMDH*

Finally, the above-derived means, SDs and sample sizes of the intervention and control arms across each study’s longitudinal sampling regimen were translated into SMDHs using the ‘escalc’ function of the metafor R package v4.8.0^16^. This resulted in four versions of the SMDH used acros

- **SMDH of absolute abundance of resistance all determinants:** The primary outcome measure used across all analyses in our quantitative synthesis.
- **SMDH of absolute abundance of all determinants:** A secondary outcome measure used across all analyses in our quantitative synthesis to contextualise the primary outcome results.
- **SMDH of relative abundance of resistance determinants (logit-transformed):** A secondary outcome measure used in our analysis of the Results of individual studies for reasons of transparency, despite statistical issues (see above).
- **SMDH of relative abundance of resistance determinants (arcsine-transformed):** A secondary outcome measure used in our analysis of the Results of individual studies for reasons of transparency, despite statistical issues (see above).

*Validation*

For each effect, data preparation was validated by comparing the most relevant plot from the original publication to longitudinal plots of the processed abundance data for the control and intervention groups. In almost all cases, the plots of the processed abundance closely approximated the original plots (Figure only available on request due to copyright associated with the original plots). Slight differences can be accounted for by the fact that the processing steps used in the original publication were not always clear (none of the included studies shared reproducible data and code as part of the original publication), and/or that the nearest matching plot in the original publication reported marginal means from mixed models.

Plots of the “USMARC - Tylosin with monensin”^19^ study data were quite different from the original plots - likely because the nearest matching plot was a plot of marginal means and their SEs across two trials (i.e. represented aggregate measures across two trials), whereas we had to separately calculate pen means and SDs for each block for our meta-analysis (see above). Combined with the fact that not all of the processing steps used in the original publication were clear, this probably explains this difference. To a much lesser extent, plots of the “Texas A&M - Tylosin”^9^ study data also differed from the original plot, which was again probably due to our processing necessarily differing from theirs. Specifically, upon receiving the data, it appeared that there was no replication within weight blocks (a point of ambiguity, but not necessarily invalid in a randomised complete block design) and that earlier and later timepoints appeared to have measured in different pens (i.e. again, not necessarily invalid, but meant that more dependence than existed would be assumed when assuming temporal autocorrelation within study arms in an analysis). This meant that we had to average across weight blocks when processing the data, rather than calculating means within each block. We also treated the final two timepoints in the longitudinal series as separate, continuous timepoints - rather than combining them into one categorical ‘End’ timepoint as they had in the original study, explaining differences in the last timepoint(s). Nonetheless, we consider these choices conservative since they assumed less power than the original analyses, and the general patterns between figures in the publication^9^ and plotted data were very similar, which should have resulted in similar effect estimates. Moreover, the sensitivity of our results to particular studies such as these was tested in our Leave-One-Out Sensitivity Analysis (Methods: Synthesis Methods: Subgroup and other alternative analyses).

We also produced longitudinal plots of the SMDHs derived from the processed data, to validate that they aligned with the expected patterns in the control and intervention data.

**Metadata extraction and standardisation**

Other variables related to timing of sampling, antibiotic dose and publication bias were also extracted and/or (re)calculated and used as moderators in meta-regressions for our primary, alternative, and publication bias analyses. For the IPD, these were extracted as part of processing the outcome data in custom R scripts, filling in missing metadata manually using information from the publications and/or discussion with the authors, before writing the final CSV (see below). For the data derived from published figures, these metadata were extracted directly into an Excel spreadsheet. These were then combined into one dataframe, before standardising and/or deriving new estimates from several variables through a R script, to permit their use in meta-analysis.

*Timing of sampling*

Central to our meta-analytical approach was engaging with the dependence of effects on the timing of their assessment in these longitudinal studies, which required extracting data related to the timing of sampling in several forms.

First, we extracted a numeric variable representing the timing of sampling as reported in the study i.e. the number of days since the start of the study that each measurement was taken. This variable (‘outcome_study_days’) was log10-transformed before being used to model temporal autocorrelation between longitudinal measurements within the same study arm via a random effect included in all models.

Second, we created a numeric variable reflecting either the number of days since the start or the end of antibiotic administration, depending on whether the measurement was taken during antibiotic administration (<1 day since the last administration event) or after antibiotic administration (>1 day since the last administration event), respectively (with measures taken pre-antibiotic administration coded as NA). This variable (‘outcome_time_days’) was log10-transformed before being used as the fixed effect in our hypothesis 2 models (meta-regressions).

Finally, we created two categorical variables reflecting whether the measurement was taken pre- or post- antibiotic administration, or during or after antibiotic administration. This was used to split the dataset four ways for each sample type (e.g. faecal, pen soil), since our approach to quantitative summary and synthesis involved modelling each of these subsets separately (see ‘Quantitative summary and synthesis’).

*Antibiotic dose variable*

Daily antibiotic dose values extracted from publications that were not expressed in estimated milligrams per kilogram of the receiving animal’s body weight (mg/kg body weight) were standardised into this format. Dosages expressed as milligram per pound of the receiving animal’s bodyweight were multiplied by 2.20462 to give mg/kg body weight. Dosages expressed as grams per tonne of feed/milligram per kilogram of feed, or parts per million were multiplied by 0.02, based on the estimate that cattle eat about 2% of their body weight per day^20^. Dosages expressed as milligram per head were divided by the estimated weight of cattle in each study. Where reported, we used the mean weight of cattle in the study. For studies which did not report the mean weight of cattle, we used the mean weight of cattle across studies that did report it. Once all dosages had been standardised into mg/kg body weight in this way, we also calculated the cumulative dosage administered before each measurement (by multiplying by the cumulative intervention length). We also created log10-transformed versions of the daily and cumulative dosage variables, as it was not entirely clear on which scale dosages were distributed. These four variables were used for alternative meta-regressions of the effect of dose, rather than time. The results of these analyses informed our decision of whether to upgrade the outcomes based on a dose-response effect in the GRADE Certainty assessment.

*Publication (reporting) bias variables*

The approach of Nakagawa^21^, was followed to calculate the variables needed for meta-regression tests of small-study effects (i.e. via adjusted Egger’s regressions) and time-lag (i.e. decline effects) publication bias.

Effective sample size using previously published methods^22,23^ using the following formula:

$$4\tilde{n}_{i}=\frac{4n_{1i}n_{2i}}{n_{1i}+n_{2i}},$$

where $n_{1i}$ is the sample size of the control group and $n_{2i}$ is the sample size of the treatment group. This calculation of sample size accounts for unbalanced sampling in control and treatment groups, avoiding artefactual patterns in funnel plots/Egger’s regressions^21^.

The year of publication of each paper from which the effect sizes were derived was mean-centered using the base R function ‘scale’ to facilitate estimate interpretation following Nakagawa^21^.

Supplementary Information 6: Extracted data for meta-analysis

**Meta-analytical dataset**

1. **id:** Arbitrary ID assigned to each observation. Factor variable with as many levels as rows.
2. **study_publicationDOI:** DOI of the publication from which the observation is derived. Factor variable whose number is levels is equal to the number of included publications.
3. **study_publicationID:** Shorthand/computer-readable name assigned to each study in the format ‘Institution_Interventions’. Different antibiotic interventions in studies with multiple are separated by camel case. Factor variable whose number is levels is equal to the number of included publications.
4. **study_studyID:** Shorthand/computer-readable name assigned to each study in the format ‘Institution_Interventions’. Different antibiotic interventions in studies with multiple are separated by camel case. Factor variable whose number is levels is equal to the number of included studies.
5. **study_studydesign:** Design name (randomised blockd design, completely randomised design, or non-randomised design); number of animals in intervention/control group(s) and number of pens in which they were housed (pens are typically considered the unit of analysis and are for the meta-analysis where possible, though this is not true for every study’s analysis or extracted means). Factor variable whose number is levels is equal to the number of included studies.
6. **population_reproductiveclass: R**eproductive class of the animals in the study (e.g. steer, heifer). Character variable with multiple reproductive classes separated by semicolons where applicable.
7. **population_breed:** Breed of the animals in the study. Where multiple classes, they are separated by semicolons. Character variable with multiple breeds separated by semicolons where applicable.
8. **population_weight_kg:** Mean weight of the animals in the study if reported in a publication (standard deviation not recorded though frequently reported). Numeric variable expressed in kilograms (converted from other units where necessary).
9. **population_minage_days:** Minimum age of the animals in the study if reported in a publication. Numeric variable expressed in days (converted from other units where necessary).
10. **population_maxage_days:** Maximum age of the animals in the study if reported in a publication, converted to days. Numeric variable expressed in days (converted from other units where necessary).
11. **population_antibioticfreebefore:** Whether or not the animals in study were free from antibiotic administration prior to the study. Factor variable with levels ‘Unknown’, ‘True’ (antibiotic-free), ‘False’ (antibiotic-exposed).
12. **intervention_name:** Name of the intervention, as described in the publication and/or the raw data obtained. Factor variable whose number is levels is equal to the number of included study arms.
13. **intervention_pens_number:** Number of pens across which the intervention arm is housed (also considered the appropriate number of replicates/unit of replication). Numeric variable with a value equal to the total number of pens divided by the number of study arms.
14. **intervention_animals_perpen:** Number of animals in each of the intervention pens. Where this varies, the mean value is used. Numeric variable with a value equal to the total number of animals divided by the number of pens per study arm.
15. **intervention_total_animals:** Total number of animals in the intervention arm. Numeric variable equal to the total number of animals in the study divided by the total number of study arms.
16. **intervention_proportion_perpen:** Proportion of animals receiving the intervention, per pen. Numeric value between 0 and 1.
17. **intervention_purpose:** Typical purpose of the antibiotic intervention outside of the study context. Feeding of antibiotics such as tylosin for growth promotion purposes is distinguished from prophylaxis for liver abscesses, though the distinction is sometimes blurry. Factor variable with levels ‘growthpromotion’, ‘prophylaxis’, and ‘metaphylaxis’.
18. **intervention_antibiotic_name:** Name of the antibiotic used in the intervention. Factor variable equal to the number of different antibiotics used across all included study arms.
19. **intervention_antibiotic_class:** Class of the antibiotic used in the intervention. Factor variable equal to the number of different antibiotic classes used across all included study arms. *Used as a moderator in a subgroup analysis.*
20. **intervention_antibiotic_brand:** Name of the antibiotic brand used in the intervention. Factor variable equal to the number of different antibiotic brands used across all included study arms.
21. **intervention_mode:** Mode of administering the antibiotic used in the intervention. Factor variable with levels ‘feed’ and ‘injection’.
22. **intervention_length_days:** Length of the antibiotic intervention (continuous administration period of the first administration event - where there are multiple, intermittent administration periods only observations after the first and before subsequent administration periods are included). Numeric variable expressed in days.
23. **intervention_dosage_value:** Dose of the antibiotic intervention as reported in publication(s). Numeric value expressed in the original reported units (see next column). This column is used when converting all dosages ‘mg_kg’ in the four forms expressed in ‘intervention_perdayUDD_mgkg’, ‘intervention_perdayUDDlog10_mgkg’, ‘intervention_cumulativeUDD_mgkg’, and ‘intervention_cumulativeUDDlog10_mgkg’ columns. *Used to calculate a moderator used in a subgroup analysis.*
24. **intervention_dosage_unit:** Units of the dose of the antibiotic intervention recorded in the ‘intervention_dosage_unit’ column. Factor variable with levels ‘g_tonne_feed’, ‘ppm_feed’, ‘mg_kg_bodyweight’, ‘mg_lb_bodyweight’, and ‘mg_head’. This column is used when converting all dosages ‘mg_kg’ in the four forms expressed in ‘intervention_perdayUDD_mgkg’, ‘intervention_perdayUDDlog10_mgkg’, ‘intervention_cumulativeUDD_mgkg’, and ‘intervention_cumulativeUDDlog10_mgkg’ columns. *Used to calculate a moderator used in a subgroup analysis.*
25. **intervention_cointervention_TF:** Whether or not a cointervention antibiotic was given alongside the intervention antibiotic. Given our eligibility criteria, this should be equal to the value of ‘control_cointervention_TF**’.** Logical variable with values ’TRUE’ (cointervention administered) and ’FALSE (cointervention not administered). *Used as a moderator in a subgroup analysis.*
26. **cointervention1_purpose:** Typical purpose of the first antibiotic cointervention outside of the study context. Feeding of antibiotics for growth promotion purposes is distinguished from prophylaxis for liver abscesses, though the distinction is sometimes blurry. Factor variable with levels ‘growthpromotion’, ‘prophylaxis_LA’ (prophylaxis for liver abscesses), ‘prophylaxis_BRD’ (prophylaxis for bovine respiratory disease), and ‘metaphylaxis’.
27. **cointervention1_antibiotic_name:** Name of the antibiotic used in the first cointervention. Factor variable equal to the number of different antibiotics used across all included study arms.
28. **cointervention1_antibiotic_class:** Name of the antibiotic used in the first cointervention. Factor variable equal to the number of different antibiotic classes used across all included study arms.
29. **cointervention1_antibiotic_brand:** Name of the antibiotic brand used in the first intervention. Factor variable equal to the number of different antibiotic brands used across all included study arms.
30. **cointervention1_mode:** Mode of administering the antibiotic used in the first intervention. Factor variable with levels ‘feed’ and ‘injection’.
31. **cointervention1_length_days:** Length of the first antibiotic cointervention (continuous administration period of the first administration event - where there are multiple, intermittent administration periods only observations after the first and before subsequent administration periods are included). Numeric variable expressed in days.
32. **cointervention1_dosage_value:** Dose of the first antibiotic intervention as reported in publication(s). Numeric value expressed in the original reported units (see next column). This column is used when converting all dosages ‘mg_kg’ in the four forms expressed in ‘intervention_perdayUDD_mgkg’, ‘intervention_perdayUDDlog10_mgkg’, ‘intervention_cumulativeUDD_mgkg’, and ‘intervention_cumulativeUDDlog10_mgkg’ columns.
33. **cointervention1_dosage_unit:** Units of the dose of the first antibiotic cointervention recorded in the ‘intervention_dosage_unit’ column. Factor variable with levels ‘g_tonne_feed’, ‘ppm_feed’, ‘mg_kg_bodyweight’, ‘mg_lb_bodyweight’, and ‘mg_head’. This column is used when converting all dosages ‘mg_kg’ in the four forms expressed in ‘intervention_perdayUDD_mgkg’, ‘intervention_perdayUDDlog10_mgkg’, ‘intervention_cumulativeUDD_mgkg’, and ‘intervention_cumulativeUDDlog10_mgkg’ columns.
34. **cointervention2_purpose:** Typical purpose of the second antibiotic cointervention outside of the study context. Feeding of antibiotics for growth promotion purposes is distinguished from prophylaxis for liver abscesses, though the distinction is sometimes blurry. Factor variable with levels ‘growthpromotion’, ‘prophylaxis_LA’ (prophylaxis for liver abscesses), ‘prophylaxis_BRD’ (prophylaxis for bovine respiratory disease), and ‘metaphylaxis’.
35. **cointervention2_antibiotic_name:** Name of the antibiotic used in the second cointervention. Factor variable equal to the number of different antibiotics used across all included study arms.
36. **cointervention2_antibiotic_class:** Name of the antibiotic used in the second cointervention. Factor variable equal to the number of different antibiotic classes used across all included study arms.
37. **cointervention2_antibiotic_brand:** Name of the antibiotic brand used in the second intervention. Factor variable equal to the number of different antibiotic brands used across all included study arms.
38. **cointervention2_mode:** Mode of administering the antibiotic used in the second intervention. Factor variable with levels ‘feed’ and ‘injection’.
39. **cointervention2_length_days:** Length of the second antibiotic cointervention (continuous administration period of the first administration event - where there are multiple, intermittent administration periods only observations after the first and before subsequent administration periods are included). Numeric variable expressed in days.
40. **cointervention2_dosage_value:** Dose of the second antibiotic intervention as reported in publication(s). Numeric value expressed in the original reported units (see next column). This column is used when converting all dosages ‘mg_kg’ in the four forms expressed in ‘intervention_perdayUDD_mgkg’, ‘intervention_perdayUDDlog10_mgkg’, ‘intervention_cumulativeUDD_mgkg’, and ‘intervention_cumulativeUDDlog10_mgkg’ columns.
41. **cointervention2_dosage_unit:** Units of the dose of the second antibiotic cointervention recorded in the ‘intervention_dosage_unit’ column. Factor variable with levels ‘g_tonne_feed’, ‘ppm_feed’, ‘mg_kg_bodyweight’, ‘mg_lb_bodyweight’, and ‘mg_head’. This column is used when converting all dosages ‘mg_kg’ in the four forms expressed in ‘intervention_perdayUDD_mgkg’, ‘intervention_perdayUDDlog10_mgkg’, ‘intervention_cumulativeUDD_mgkg’, and ‘intervention_cumulativeUDDlog10_mgkg’ columns.
42. **control_name:** Name of the control, as described in the publication and/or the raw data obtained. Factor variable whose number is levels is equal to the number of included study arms.
43. **control_pens_number:** Number of pens across which the control arm is housed (also considered the appropriate number of replicates/unit of replication). Numeric variable with a value equal to the total number of pens divided by the number of study arms.
44. **control_animals_perpen:** Number of animals in each of the control pens. Where this varies, the mean value is used. Numeric variable with a value equal to the total number of animals divided by the number of pens per study arm.
45. **control_total_animals:** Total number of animals in the control arm. Numeric variable equal to the total number of animals in the study divided by the total number of study arms.
46. **control_proportion_perpen:** Proportion of animals receiving the control treatment, per pen. Numeric value between 0 and 1.
47. **control_cointervention_TF:** Whether or not a cointervention antibiotic was given alongside the control antibiotic. Given our eligibility criteria, this should be equal to the value of ‘intervention_cointervention_TF**’.** Logical variable with values ’TRUE’ (cointervention administered) and ’FALSE (cointervention not administered).
48. **outcome_date_yyyymmdd:** Date on which this observation of the outcome was made, if reported. Date variable in YYYYMMDD format.
49. **outcome_study_days:** Number of days into the study that this observation of the outcome was made. Numeric variable expressed as log10-transformed days+1 (pseudocount to avoid infinity values for zero).
50. **outcome_timesinceintervention_start_days:** Number of days since the start of antibiotic administration that this observation of the outcome was made (for observations taken during ongoing antibiotic administration). *Used as a moderator in the primary meta-analysis (H2).* Numeric variable expressed as days.
51. **outcome_timesinceintervention_end_days:** Number of days since the end of antibiotic administration that this observation of the outcome was made (for observations taken after the cessation of antibiotic administration). *Used as a moderator in the primary meta-analysis (H2).* Numeric variable expressed as days.
52. **outcome_organism:** Shorthand denotation of the target microorganisms for this observation. Factor variable with classes ‘ecoli’ (*E. coli*), ‘ent/entero’ (*Enterococcus spp.*), ‘salm’ (*Salmonella spp.) and ’*bacteria’ (bacteria generally i.e. qPCR).
53. **outcome_units:** Units in which the outcome is reported. Factor variable with levels ‘cfuml’ (CFU/ml), ‘cfug’ (CFU/gram), ‘cfuswab’ (CFU/swab), ‘genespergram’ (genes per gram of faeces for qPCR measures).
54. **outcome_resistance_target:** Shorthand denotation of the target antibiotic (direct enumeration) or gene (qPCR) used to assess resistance for this observation. Factor variable whose number of levels is equal to the number of different resistance targets used across all included study arms.
55. **outcome_resistance_antibiotic:** Target antibiotic used to assess resistance for direct enumeration assessments. Factor variable whose number of levels is equal to the number of different resistance targets used across all included study arms.
56. **outcome_resistance_studybreakpointmgL:** Breakpoint used to assess resistance for direct enumeration assessments. Factor variable whose number of levels is equal to the number of different resistance targets used across all included study arms.
57. **outcome_resistance_CLSIorNARMSbreakpointmgL:** Breakpoint type (of the value in column ‘outcome_resistance_studybreakpointmgL’) used to assess resistance for direct enumeration assessments. Factor variable whose number of levels is equal to the number of different resistance targets used across all included study arms.
58. **outcome_series_id:** Arbitrary ID denoting the longitudinal series to which the observation belongs within this study. A longitudinal series is considered a group of repeated measurements of the same outcome within the same study arm (i.e. there may multiple series per study arm if there are multiple, alternative measures of the resistance outcome). Numeric variable whose maximum value within each study represents the number of series meeting the eligibility criteria for the meta-analysis.
59. **outcome_sample_type:** The matrix from which the target bacteria or their DNA were isolated (e.g. fresh faeces, pen surface soil, feed, water). *Used to split the dataset by sample matrix before meta-analysis.*
60. **outcome_sample_instrument:** Instrument used to obtain the sample in which the outcome was assessed. Logical variable with levels ‘grab/gloves’ (e.g. faecal or soil grab sample), ‘bag’ (bag sample), ’’swab” (swab sample).
61. **outcome_sample_subtype:** Additional detail on how the sampling instrument was applied/how sampling was performed.. Factor variable with levels ‘per_rectum’ (i.e. one sample per rectum of each sampled animal) and ‘per_quadrant’ (i.e. one sample per quadrant of each sampled enclosure).
62. **outcome_measurement_type:** Type of outcome measurement/method of assessing resistance. Factor variable with levels ‘Direct plating’ and ‘Quantitative PCR’. *Used as the moderator in a subgroup analysis.*
63. **outcome_unitofanalysis:** Unit of analysis at which the means and standard deviations were (re)calculated, either by the original investigators (extracted) or by the evidence synthesis team (from raw). Factor variable with levels ‘animal’ (where all animals in same study arm are in the same pen or all animals in the study are in the same pen) and ‘pen’ (where study animals are housed across multiple pens). *Used as the moderator in a subgroup analysis.*
64. **outcome_origanalysisaccountedforpen:** Whether or not the original analysis accounted for pen, if applicable (i.e. if animals were housed across multiple pens). Logical variable with values ‘TRUE’ (pen was accounted for via random effects and/or aggregating at the pen level) and ‘FALSE’ (pen was not accounted for/animals were treated as independent despite co-housing).
65. **outcome_metaanalysisaccountedforpen:** Whether or not the meta-analysis analysis accounted for pen, if possible (i.e. if a pen variable was included in the datasets provided upon request). Logical variable with values ‘TRUE’ (pen was accounted for via random effects and/or aggregating at the pen level) and ‘FALSE’ (pen was not accounted for/animals were treated as independent despite co-housing).
66. **outcome_origanalysisaccountedforblocking:** Whether or not the original analysis accounted for blocking, if applicable (i.e. if animals were blocked before randomisation/a random block design was used). Logical variable with values ‘TRUE’ (block was accounted for via random effects and/or aggregating at the pen level) and ‘FALSE’ (block was not accounted for/animals were treated as independent despite blocking).
67. **outcome_metaanalysisaccountedforblocking:** Whether or not the meta-analysis analysis accounted for block, if possible (i.e. if a block variable was included in the datasets provided upon request). Logical variable with values ‘TRUE’ (block was accounted for via random effects and/or aggregating at the pen level) and ‘FALSE’ (block was not accounted for/animals were treated as independent despite co-housing).
68. **outcome_from_raworextracted:** Whether the means, standard deviations and replicates in the following columns were derived from an IPD-style processing of raw data, or extracted from figures in the publications. Factor variable with levels ‘raw’ and ‘extracted’.
69. **outcome_actual_totalresistancedeterminantslog10_intervention_mean:** IPD-derived mean of the log10-transformed absolute abundance of resistance determinants (i.e. density of resistant bacteria or resistance genes) in the intervention/exposure group. Numeric variable with NAs for data that was extracted from publications (i.e. not derived from IPD).
70. **outcome_actual_totalresistancedeterminantslog10_intervention_sd:** IPD-derived standard deviation of the log10-transformed absolute abundance of resistance determinants (i.e. density of resistant bacteria or resistance genes) in the intervention/exposure group. Numeric variable with NAs for data that was extracted from publications (i.e. not derived from IPD).
71. **outcome_actual_totalresistancedeterminantslog10_intervention_reps:** IPD-derived number of replicates (sample size) of the log10-transformed absolute abundance of resistance determinants (i.e. density of resistant bacteria or resistance genes) in the intervention/exposure group. Numeric variable with NAs for data that was extracted from publications (i.e. not derived from IPD).
72. **outcome_actual_totaldeterminantslog10_intervention_mean:** IPD-derived mean of the log10-transformed absolute abundance of total determinants (i.e. density of all target bacteria or target genes) in the intervention/exposure group. Numeric variable with NAs for data that was extracted from publications (i.e. not derived from IPD).
73. **outcome_actual_totaldeterminantslog10_intervention_sd:** IPD-derived standard deviation of the log10-transformed absolute abundance of total determinants (i.e. density of all target bacteria or target genes) in the intervention/exposure group. Numeric variable with NAs for data that was extracted from publications (i.e. not derived from IPD).
74. **outcome_actual_totaldeterminantslog10_intervention_reps:** IPD-derived number of replicates (sample size) of the log10-transformed absolute abundance of total determinants (i.e. density of all target bacteria or target genes) in the intervention/exposure group. Numeric variable with NAs for data that was extracted from publications (i.e. not derived from IPD).
75. **outcome_actual_logitproportionresistancedeterminants_intervention_mean:** IPD-derived mean of the logit-transformed proportion of resistance determinants (i.e. absolute abundance of resistance determinants divided by the absolute abundance of all determinants) in the intervention/exposure group. Numeric variable with NAs for data that was extracted from publications (i.e. not derived from IPD).
76. **outcome_actual_logitproportionresistancedeterminants_intervention_sd:** IPD-derived standard deviation of the logit-transformed proportion of resistance determinants (i.e. absolute abundance of resistance determinants divided by the absolute abundance of all determinants) in the intervention/exposure group. Numeric variable with NAs for data that was extracted from publications (i.e. not derived from IPD).
77. **outcome_actual_logitproportionresistancedeterminants_intervention_reps:** IPD-derived number of replicates (sample size) of the arcsine-transformed proportion of resistance determinants (i.e. absolute abundance of resistance determinants divided by the absolute abundance of all determinants) in the intervention/exposure group. Numeric variable with NAs for data that was extracted from publications (i.e. not derived from IPD).
78. **outcome_actual_arcsinproportionresistancedeterminants_intervention_mean:** IPD-derived mean of the arcsin-transformed proportion of resistance determinants (i.e. absolute abundance of resistance determinants divided by the absolute abundance of all determinants) in the intervention/exposure group. Numeric variable with NAs for data that was extracted from publications (i.e. not derived from IPD).
79. **outcome_actual_arcsinproportionresistancedeterminants_intervention_sd:** IPD-derived standard deviation of the arcsin-transformed proportion of resistance determinants (i.e. absolute abundance of resistance determinants divided by the absolute abundance of all determinants) in the intervention/exposure group. Numeric variable with NAs for data that was extracted from publications (i.e. not derived from IPD).
80. **outcome_actual_arcsinproportionresistancedeterminants_intervention_reps:** IPD-derived number of replicates (sample size) of the arcsin-transformed proportion of resistance determinants (i.e. absolute abundance of resistance determinants divided by the absolute abundance of all determinants) in the intervention/exposure group. Numeric variable with NAs for data that was extracted from publications (i.e. not derived from IPD).
81. **outcome_actual_totalresistancedeterminantslog10_control_mean:** IPD-derived mean of the log10-transformed absolute abundance of resistance determinants (i.e. density of resistant bacteria or resistance genes) in the control group. Numeric variable with NAs for data that was extracted from publications (i.e. not derived from IPD).
82. **outcome_actual_totaldeterminantslog10_control_mean:** IPD-derived mean of the log10-transformed absolute abundance of total determinants (i.e. density of all target bacteria or target genes) in the control group. Numeric variable with NAs for data that was extracted from publications (i.e. not derived from IPD).
83. **outcome_actual_logitproportionresistancedeterminants_control_mean:** IPD-derived mean of the logit-transformed proportion of resistance determinants (i.e. absolute abundance of resistance determinants divided by the absolute abundance of all determinants) in the control group. Numeric variable with NAs for data that was extracted from publications (i.e. not derived from IPD).
84. **outcome_actual_arcsinproportionresistancedeterminants_control_mean:** IPD-derived mean of the arcsin-transformed proportion of resistance determinants (i.e. absolute abundance of resistance determinants divided by the absolute abundance of all determinants) in the control group. Numeric variable with NAs for data that was extracted from publications (i.e. not derived from IPD).
85. **outcome_actual_totalresistancedeterminantslog10_control_sd:** IPD-derived standard deviation of the log10-transformed absolute abundance of resistance determinants (i.e. density of resistant bacteria or resistance genes) in the control group. Numeric variable with NAs for data that was extracted from publications (i.e. not derived from IPD).
86. **outcome_actual_totaldeterminantslog10_control_sd:** IPD-derived standard deviation of the log10-transformed absolute abundance of total determinants (i.e. density of all target bacteria or target genes) in the control group. Numeric variable with NAs for data that was extracted from publications (i.e. not derived from IPD).
87. **outcome_actual_logitproportionresistancedeterminants_control_sd:** IPD-derived standard deviation of the logit-transformed proportion of resistance determinants (i.e. absolute abundance of resistance determinants divided by the absolute abundance of all determinants) in the control group. Numeric variable with NAs for data that was extracted from publications (i.e. not derived from IPD).
88. **outcome_actual_arcsinproportionresistancedeterminants_control_sd:** IPD-derived standard deviation of the arcsin-transformed proportion of resistance determinants (i.e. absolute abundance of resistance determinants divided by the absolute abundance of all determinants) in the control group. Numeric variable with NAs for data that was extracted from publications (i.e. not derived from IPD).
89. **outcome_actual_totalresistancedeterminantslog10_control_reps:** IPD-derived number of replicates (sample size) of the log10-transformed absolute abundance of resistance determinants (i.e. density of resistant bacteria or resistance genes) in the control group. Numeric variable with NAs for data that was extracted from publications (i.e. not derived from IPD).
90. **outcome_actual_totaldeterminantslog10_control_reps:** IPD-derived number of replicates (sample size) of the log10-transformed absolute abundance of total determinants (i.e. density of all target bacteria or target genes) in the control group. Numeric variable with NAs for data that was extracted from publications (i.e. not derived from IPD).
91. **outcome_actual_logitproportionresistancedeterminants_control_reps:** IPD-derived number of replicates (sample size) of the logit-transformed proportion of resistance determinants (i.e. absolute abundance of resistance determinants divided by the absolute abundance of all determinants) in the control group. Numeric variable with NAs for data that was extracted from publications (i.e. not derived from IPD).
92. **outcome_actual_arcsinproportionresistancedeterminants_control_reps:** IPD-derived number of replicates (sample size) of the arcsin-transformed proportion of resistance determinants (i.e. absolute abundance of resistance determinants divided by the absolute abundance of all determinants) in the control group. Numeric variable with NAs for data that was extracted from publications (i.e. not derived from IPD).
93. **outcome_interventionssharingcontrol:** Number of interventions that share the control group. Integer variable equal to the number of intervention arms divided by the number of control arms. *Used in a sensitivity analysis.*
94. **outcome_plot_actualormarginal:** Whether the means and standard deviations extracted from published plots/figures are actual or marginal (i.e. adjusted for other factors in the model). They should all be marginal as we excluded marginal means and standard deviations from the meta-analysis because there was no way to derive standard deviations for them. Factor variable with classes ‘actual’ and ’marginal.
95. **outcome_plot_varbartype:** Type of variability/error bar extracted from the published plots/figures. Factor variable with classes ‘std_error’ (standard error).
96. **outcome_plot_totaldeterminantslog10_intervention_mean:** Plot extraction-derived mean of the log10-transformed absolute abundance of total determinants (i.e. density of all target bacteria or target genes) in the intervention/exposure group. Numeric variable with NAs for data that was extracted from publications (i.e. not derived from IPD).
97. **outcome_plot_totaldeterminantslog10_intervention_errorbarlimit:** Plot extraction-derived limit of the error bar the log10-transformed absolute abundance of total determinants (i.e. density of all target bacteria or target genes) in the intervention/exposure group. Can be the upper or lower limit, whatever is most convenient to extracted, as this is converted to an absolute distance from the mean before using it to calculate the standard deviation, taking into account ’**outcome_plot_varbartype’.** Numeric variable with NAs for data that was extracted from publications (i.e. not derived from IPD).
98. **outcome_plot_totaldeterminantslog10_intervention_reps:** Plot extraction-derived number of replicates (sample size) of the log10-transformed absolute abundance of total determinants (i.e. density of all target bacteria or target genes) in the intervention/exposure group. Numeric variable with NAs for data that was extracted from publications (i.e. not derived from IPD).
99. **outcome_plot_totaldeterminantslog10_control_mean:** Plot extraction-derived mean of the log10-transformed absolute abundance of total determinants (i.e. density of all target bacteria or target genes) in the control group. Numeric variable with NAs for data that was extracted from publications (i.e. not derived from IPD).
100. **outcome_plot_totaldeterminantslog10_control_errorbarlimit:** Plot extraction-derived limit of the error bar the log10-transformed absolute abundance of total determinants (i.e. density of all target bacteria or target genes) in the control group. Can be the upper or lower limit, whatever is most convenient to extracted, as this is converted to an absolute distance from the mean before using it to calculate the standard deviation, taking into account **outcome_plot_varbartype** Numeric variable with NAs for data that was extracted from publications (i.e. not derived from IPD).
101. **outcome_plot_totaldeterminantslog10_control_reps:** Plot extraction-derived number of replicates (sample size) of the log10-transformed absolute abundance of total determinants (i.e. density of all target bacteria or target genes) in the control group. Numeric variable with NAs for data that was extracted from publications (i.e. not derived from IPD).
102. **outcome_plot_totalresistancedeterminantslog10_intervention_mean:** Plot extraction-derived mean of the log10-transformed absolute abundance of resistance determinants (i.e. density of resistant bacteria or resistance genes) in the intervention/exposure group. Numeric variable with NAs for data that was extracted from publications (i.e. not derived from IPD).
103. **outcome_plot_totalresistancedeterminantslog10_intervention_errorbarlimit:** Plot extraction-derived limit of the error bar the log10-transformed absolute abundance of resistance determinants (i.e. density of resistant bacteria or resistance genes) in the intervention/exposure group. Can be the upper or lower limit, whatever is most convenient to extracted, as this is converted to an absolute distance from the mean before using it to calculate the standard deviation, taking into account **outcome_plot_varbartype** Numeric variable with NAs for data that was extracted from publications (i.e. not derived from IPD).
104. **outcome_plot_totalresistancedeterminantslog10_intervention_reps:** Plot extraction-derived number of replicates (sample size) of the log10-transformed absolute abundance of resistance determinants (i.e. density of resistant bacteria or resistance genes) in the intervention/exposure group. Numeric variable with NAs for data that was extracted from publications (i.e. not derived from IPD).
105. **outcome_plot_totalresistancedeterminantslog10_control_mean:** Plot extraction-derived mean of the log10-transformed absolute abundance of resistance determinants (i.e. density of resistant bacteria or resistance genes) in the control group. Numeric variable with NAs for data that was extracted from publications (i.e. not derived from IPD).
106. **outcome_plot_totalresistancedeterminantslog10_control_errorbarlimit:** Plot extraction-derived limit of the error bar the log10-transformed absolute abundance of resistance determinants (i.e. density of resistant bacteria or resistance genes) in the control group. Can be the upper or lower limit, whatever is most convenient to extracted, as this is converted to an absolute distance from the mean before using it to calculate the standard deviation, taking into account **outcome_plot_varbartype** Numeric variable with NAs for data that was extracted from publications (i.e. not derived from IPD).
107. **outcome_plot_totalresistancedeterminantslog10_control_reps:** Plot extraction-derived number of replicates (sample size) of the log10-transformed absolute abundance of resistance determinants (i.e. density of resistant bacteria or resistance genes) in the control group. Numeric variable with NAs for data that was extracted from publications (i.e. not derived from IPD).
108. **outcome_plot_proportionorpercentage:** Type of proportion data extracted from the published plots/figures. Factor variable with classes ‘proportion’ (proportion) or ‘percentage’ (percentage; must be converted to proportion by dividing by 100 before further analysis).
109. **outcome_plot_proportionresistancedeterminants_intervention_mean:** Plot extraction-derived mean of the proportion of resistance determinants (i.e. absolute abundance of resistance determinants divided by the absolute abundance of all determinants) in the intervention/exposure group. Numeric variable with NAs for data that was extracted from publications (i.e. not derived from IPD).
110. **outcome_plot_proportionresistancedeterminants_intervention_errorbarlimit:** Plot extraction-derived error bar limit of the proportion of resistance determinants (i.e. absolute abundance of resistance determinants divided by the absolute abundance of all determinants) in the intervention/exposure group. Can be the upper or lower limit, whatever is most convenient to extracted, as this is converted to an absolute distance from the mean before using it to calculate the standard deviation, taking into account **outcome_plot_varbartype** Numeric variable with NAs for data that was extracted from publications (i.e. not derived from IPD).
111. **outcome_plot_proportionresistancedeterminants_intervention_reps**: Plot extraction-derived number of replicates (sample size) of the proportion of resistance determinants (i.e. absolute abundance of resistance determinants divided by the absolute abundance of all determinants) in the intervention/exposure group. Numeric variable with NAs for data that was extracted from publications (i.e. not derived from IPD).
112. **outcome_plot_proportionresistancedeterminants_control_mean:** Plot extraction-derived mean of the proportion of resistance determinants (i.e. absolute abundance of resistance determinants divided by the absolute abundance of all determinants) in the control group. Numeric variable with NAs for data that was extracted from publications (i.e. not derived from IPD).
113. **outcome_plot_proportionresistancedeterminants_control_errorbarlimit:** Plot extraction-derived error bar limit of the proportion of resistance determinants (i.e. absolute abundance of resistance determinants divided by the absolute abundance of all determinants) in the control group. Can be the upper or lower limit, whatever is most convenient to extracted, as this is converted to an absolute distance from the mean before using it to calculate the standard deviation, taking into account **outcome_plot_varbartype** Numeric variable with NAs for data that was extracted from publications (i.e. not derived from IPD).
114. **outcome_plot_proportionresistancedeterminants_control_reps:** Plot extraction-derived number of replicates (sample size) of the proportion of resistance determinants (i.e. absolute abundance of resistance determinants divided by the absolute abundance of all determinants) in the control group. Numeric variable with NAs for data that was extracted from publications (i.e. not derived from IPD).
115. **notes_1:** First notes column containing notes after data processing via IPD or plot extraction. Character variable with free-text entry.

- *—*
- *Variables after this point are derived from programmatic processing.*

116. **outcome_plot_totaldeterminantslog10_intervention_var:** The ‘variability’ of the log10-transformed absolute abundance of total determinants (i.e. density of all target bacteria or target genes) in the intervention/exposure group, derived by extracting the absolute distance between **outcome_plot_totaldeterminantslog10_intervention_mean’** and **outcome_plot_totaldeterminantslog10_intervention_errorbarlimit’** (‘tidy_outcomedata.R’). Numeric variable expressed on a log10-scale.
117. **outcome_plot_totaldeterminantslog10_control_var:** The ‘variability’ of the log10-transformed absolute abundance of total determinants (i.e. density of all target bacteria or target genes) in the control group, derived by extracting the absolute distance between **outcome_plot_totaldeterminantslog10_control_mean** and**outcome_plot_totaldeterminantslog10_control_errorbarlimit** (‘tidy_outcomedata.R’).Numeric variable expressed on a log10-scale.
118. **outcome_plot_totalresistancedeterminantslog10_intervention_var:** The ‘variability’ of the log10-transformed absolute abundance of resistance determinants (i.e. density of resistant bacteria or resistance genes) in the intervention/exposure group, derived by extracting the absolute distance between **outcome_plot_totaldeterminantslog10_intervention_mean** and **outcome_plot_totaldeterminantslog10_intervention_errorbarlimit** (‘tidy_outcomedata.R’).Numeric variable expressed on a log10-scale.
119. **outcome_plot_totalresistancedeterminantslog10_control_var:** The ‘variability’ of the log10-transformed absolute abundance of resistance determinants (i.e. density of resistant bacteria or resistance genes) in the control group, derived by extracting the absolute distance between **outcome_plot_totaldeterminantslog10_control_mean** and **outcome_plot_totaldeterminantslog10_control_errorbarlimit** (‘tidy_outcomedata.R’).Numeric variable expressed on a log10-scale.
120. **outcome_plot_proportionresistancedeterminants_intervention_var:** The ‘variability’ of the proportion of resistance determinants (i.e. absolute abundance of resistance determinants divided by the absolute abundance of all determinants) in the intervention/exposure group, derived by extracting the absolute distance between **outcome_plot_totaldeterminantslog10_intervention_mean** and **outcome_plot_totaldeterminantslog10_intervention_errorbarlimit** (‘tidy_outcomedata.R’).Numeric variable between 0 and 1.
121. **outcome_plot_proportionresistancedeterminants_control_var:** The ‘variability’ of the proportion of resistance determinants (i.e. absolute abundance of resistance determinants divided by the absolute abundance of all determinants) in the control group, derived by extracting the absolute distance between **outcome_plot_totaldeterminantslog10_control_mean** and **outcome_plot_totaldeterminantslog10_control_errorbarlimit** (‘tidy_outcomedata.R’). Numeric variable between 0 and 1.
122. **intervention_mean_totalresistancedeterminants:** The mean of the log10-transformed absolute abundance of resistance determinants (i.e. density of resistant bacteria or resistance genes) in the intervention/exposure group. Derived by simply moving the means derived from IPD and from plot extraction into one column (‘tidy_outcomedata.R’). Numeric variable expressed on a log10-scale.
123. **intervention_sd_totalresistancedeterminants:** The standard deviation of the log10-transformed absolute abundance of resistance determinants (i.e. density of resistant bacteria or resistance genes) in the intervention/exposure group. Derived by simply moving the standard deviations derived from IPD to this column, and converting variability (**outcome_plot_totalresistancedeterminantslog10_intervention_var**)to standard deviation in the appropriate way depending on the error bar type in **outcome_plot_varbartype**, moving into the same column (‘tidy_outcomedata.R’). Numeric variable expressed on a log10-scale.
124. **intervention_reps_totalresistancedeterminants:** The number of replicates (sample size) of the log10-transformed absolute abundance of resistance determinants (i.e. density of resistant bacteria or resistance genes) in the intervention/exposure group. Derived by simply moving the means derived from IPD and from plot extraction into one column (‘tidy_outcomedata.R’). Numeric variable expressed on a log10-scale.
125. **control_mean_totalresistancedeterminants:** The mean of the log10-transformed absolute abundance of resistance determinants (i.e. density of resistant bacteria or resistance genes) in the control group. Derived by simply moving the means derived from IPD and from plot extraction into one column (‘tidy_outcomedata.R’). Numeric variable expressed on a log10-scale.
126. **control_sd_totalresistancedeterminants:** The standard deviation of the log10-transformed absolute abundance of resistance determinants (i.e. density of resistant bacteria or resistance genes) in the control group. Derived by simply moving the standard deviations derived from IPD to this column, and converting variability (**outcome_plot_totalresistancedeterminantslog10_intervention_var**)to standard deviation in the appropriate way depending on the error bar type in **outcome_plot_varbartype**, moving into the same column (‘tidy_outcomedata.R’). Numeric variable expressed on a log10-scale.
127. **control_reps_totalresistancedeterminants:** The number of replicates (sample size) of the log10-transformed absolute abundance of resistance determinants (i.e. density of resistant bacteria or resistance genes) in the control group. Derived by simply moving the means derived from IPD and from plot extraction into one column (‘tidy_outcomedata.R’). Numeric variable expressed on a log10-scale.
128. **intervention_mean_totaldeterminants:** The mean of the log10-transformed absolute abundance of total determinants (i.e. density of all target bacteria or target genes) in the intervention/exposure group. Derived by simply moving the means derived from IPD and from plot extraction into one column (‘tidy_outcomedata.R’). Numeric variable expressed on a log10-scale.
129. **intervention_sd_totaldeterminants:** The standard deviation of the log10-transformed absolute abundance of total determinants (i.e. density of all target bacteria or target genes) in the intervention/exposure group. Derived by simply moving the standard deviations derived from IPD to this column, and converting variability (**outcome_plot_totalresistancedeterminantslog10_intervention_var**)to standard deviation in the appropriate way depending on the error bar type in **outcome_plot_varbartype**, moving into the same column (‘tidy_outcomedata.R’). Numeric variable expressed on a log10-scale.
130. **intervention_reps_totaldeterminants:** The number of replicates (sample size) of the log10-transformed absolute abundance of total determinants (i.e. density of all target bacteria or target genes) in the intervention/exposure group. Derived by simply moving the means derived from IPD and from plot extraction into one column (‘tidy_outcomedata.R’). Numeric variable expressed on a log10-scale.
131. **control_mean_totaldeterminants:** The mean of the log10-transformed absolute abundance of total determinants (i.e. density of all target bacteria or target genes) in the control group. Derived by simply moving the means derived from IPD and from plot extraction into one column (‘tidy_outcomedata.R’). Numeric variable expressed on a log10-scale.
132. **control_sd_totaldeterminants:** The standard deviation of the log10-transformed absolute abundance of total determinants (i.e. density of all target bacteria or target genes) in the control group. Derived by simply moving the standard deviations derived from IPD to this column, and converting variability (**outcome_plot_totalresistancedeterminantslog10_intervention_var**)to standard deviation in the appropriate way depending on the error bar type in **outcome_plot_varbartype**, moving into the same column (‘tidy_outcomedata.R’). Numeric variable expressed on a log10-scale.
133. **control_reps_totaldeterminants:** The number of replicates (sample size) of the log10-transformed absolute abundance of total determinants (i.e. density of all target bacteria or target genes) in the control group. Derived by simply moving the means derived from IPD and from plot extraction into one column (‘tidy_outcomedata.R’). Numeric variable expressed on a log10-scale.
134. **proportiondata_origin:** Whether the proportion data was derived from raw data or from simulating the underlying absolute abundance of resistance and total determinants (using means and standard deviations extracted from plots) and then dividing one by the other. This is distinct from whether or not the outcome data in general were from raw or extracted (‘outcome_from_raworextracted’), since only plot extracted outcome data that are not already in the proportion format needed to be derived from simulation in this way. Factor variable with levels ‘raw’ and ‘simulated’ (NAs for data derived from IPD).
135. **intervention_mean_logitpropres:** The mean of the logit-transformed proportion of resistance determinants (i.e. absolute abundance of resistance determinants divided by the absolute abundance of all determinants) in the intervention/exposure group. Derived by simply moving the means derived from IPD and calculated from simulated data based on extracted resistance/total determinants into one column (‘tidy_outcomedata.R’). Numeric variable expressed on a log10-scale.
136. **intervention_sd_logitpropres:** The standard deviation of the logit-transformed proportion of resistance determinants (i.e. absolute abundance of resistance determinants divided by the absolute abundance of all determinants) in the intervention/exposure group. Derived by simply moving the means derived from IPD and calculated from simulated data based on extracted resistance/total determinants into one column (‘tidy_outcomedata.R’). Numeric variable expressed on a log10-scale.
137. **intervention_reps_logitpropres:** The number of replicates (sample size) of the logit-transformed proportion of resistance determinants (i.e. absolute abundance of resistance determinants divided by the absolute abundance of all determinants) in the intervention/exposure group. Derived by simply moving the means derived from IPD and calculated from simulated data based on extracted resistance/total determinants into one column (‘tidy_outcomedata.R’). Numeric variable expressed on a log10-scale.
138. **intervention_mean_arcsinpropres:** The mean of the arcsin-transformed proportion of resistance determinants (i.e. absolute abundance of resistance determinants divided by the absolute abundance of all determinants) in the intervention/exposure group. Derived by simply moving the means derived from IPD and calculated from simulated data based on extracted resistance/total determinants into one column (‘tidy_outcomedata.R’). Numeric variable expressed on a log10-scale.
139. **intervention_sd_arcsinpropres:** The standard deviation of the arcsin-transformed proportion of resistance determinants (i.e. absolute abundance of resistance determinants divided by the absolute abundance of all determinants) in the intervention/exposure group. Derived by simply moving the means derived from IPD and calculated from simulated data based on extracted resistance/total determinants into one column (‘tidy_outcomedata.R’). Numeric variable expressed on a log10-scale.
140. **intervention_reps_arcsinpropres:** The number of replicates (sample size) of the arcsin-transformed proportion of resistance determinants (i.e. absolute abundance of resistance determinants divided by the absolute abundance of all determinants) in the intervention/exposure group. Derived by simply moving the means derived from IPD and calculated from simulated data based on extracted resistance/total determinants into one column (‘tidy_outcomedata.R’). Numeric variable expressed on a log10-scale.
141. **control_mean_logitpropres:** The mean of the logit-transformed proportion of resistance determinants (i.e. absolute abundance of resistance determinants divided by the absolute abundance of all determinants) in the control group. Derived by simply moving the means derived from IPD and calculated from simulated data based on extracted resistance/total determinants into one column (‘tidy_outcomedata.R’). Numeric variable expressed on a log10-scale.
142. **control_sd_logitpropres:** The standard deviation of the logit-transformed proportion of resistance determinants (i.e. absolute abundance of resistance determinants divided by the absolute abundance of all determinants) in the control group. Derived by simply moving the means derived from IPD and calculated from simulated data based on extracted resistance/total determinants into one column (‘tidy_outcomedata.R’). Numeric variable expressed on a log10-scale.
143. **control_reps_logitpropres:** The number of replicates (sample size) of the logit-transformed proportion of resistance determinants (i.e. absolute abundance of resistance determinants divided by the absolute abundance of all determinants) in the control group. Derived by simply moving the means derived from IPD and calculated from simulated data based on extracted resistance/total determinants into one column (‘tidy_outcomedata.R’). Numeric variable expressed on a log10-scale.
144. **control_mean_arcsinpropres:** The mean of the arcsin-transformed proportion of resistance determinants (i.e. absolute abundance of resistance determinants divided by the absolute abundance of all determinants) in the control group. Derived by simply moving the means derived from IPD and calculated from simulated data based on extracted resistance/total determinants into one column (‘tidy_outcomedata.R’). Numeric variable expressed on a log10-scale.
145. **control_sd_arcsinpropres:** The standard deviation of the arcsin-transformed proportion of resistance determinants (i.e. absolute abundance of resistance determinants divided by the absolute abundance of all determinants) in the control group. Derived by simply moving the means derived from IPD and calculated from simulated data based on extracted resistance/total determinants into one column (‘tidy_outcomedata.R’). Numeric variable expressed on a log10-scale.
146. **control_reps_arcsinpropres:** The number of replicates (sample size) of the arcsin-transformed proportion of resistance determinants (i.e. absolute abundance of resistance determinants divided by the absolute abundance of all determinants) in the control group. Derived by simply moving the means derived from IPD and calculated from simulated data based on extracted resistance/total determinants into one column (‘tidy_outcomedata.R’). Numeric variable expressed on a log10-scale.
147. **outcome_timeofbaseline_days:** The number of days into the study (‘outcome_study_days’) - usually after arrival to the feedlot/facility - when the baseline measure was taken. Numeric variable expressed in days.
148. **pre_or_post_intervention period:** Whether this observation was taken pre or post intervention/exposure (includes both ‘during’ and after’). Factor variable with levels ‘pre’ and ‘post’. *Used to split the data before quantitatively summarising ‘Results of individual studies’.*
149. **period:** Whether this observation was taken before, during, or after intervention/exposure. Factor variable with levels ‘before’, ‘during’, and ‘after’. *Used to split the data before meta-analysis.*
150. **intervention_perdayUDD_mgkg:** Per-day used daily doses of antibiotic administered in the intervention/exposure, calculated by standardising ‘intervention_dosage_value’ to ‘mg_kg’ (where necessary based on ‘intervention_dosage_unit’). Numeric variable expressed on a raw scale.
151. **intervention_perdayUDDlog10_mgkg:** Log10-transformed per-day used daily doses of antibiotic administered in the intervention/exposure, calculated by log10-transforming ’**intervention_perdayUDD_mgkg’** Numeric variable expressed on a log10 scale.
152. **intervention_cumulativeUDD_mgkg:** Cumulative used daily doses of antibiotic administered in the intervention/exposure up until the point of observation, calculated by multiplying **intervention_perdayUDD_mgkg** by the **outcome_timesinceintervention_start_days** Numeric variable expressed on a raw scale.
153. **intervention_cumulativeUDDlog10_mgkg:** Cumulative used daily doses of antibiotic administered in the intervention/exposure up until the point of observation, calculated by log10-transforming ’**intervention_cumulativeUDD_mgkg’.** Numeric variable expressed on a log10 scale.
154. **outcome_frommixedpen_TF:** Whether or not the intervention/exposure arm was housed in the same pen as the control arm. Dichotomised version of ‘intervention_proportion_perpen’. Logical variable with levels TRUE (housed with control arm) and FALSE (housed separately from control arm). *Used as the moderator in a subgroup analysis.*
155. **study_colour:** Unique colour type for each study’s effect sizes in figures, derived by randomly generating and assigning colours from a palette (‘tidy_metadata.R’). Factor variable with each level representing a unique hex code for each study.
156. **publication_pch:** Unique point type for each study’s effect sizes in figures, derived by randomly generating and assigning colours from a palette (‘tidy_metadata.R’). Factor variable with each level representing a unique hex code for each study.
157. **n_tilda:**
158. **inv_n_tilda:**
159. **sqrt_inv_n_tilda:**
160. **publication_year:** Year of publication, derived by parsing the number from ‘study_publicationID’ (‘tidy_metadata.R’). Numeric variable.
161. **publication_year_mc:** Mean-centered year of publication, derived by scaling ‘publication_year’ (‘tidy_metadata.R’). Numeric variable.
162. **study_studyID_pubready:** Publication-ready studyID, derived by formatting ‘study_studyID’ in a human-readable way (‘tidy_metadata.R’). Factor variable whose number is levels is equal to the number of included studies.
163. **study_publicationID_pubready:** Publication-ready publicationID, derived by formatting ‘study_publicationID’ in a human-readable way (‘tidy_metadata.R’). Factor variable whose number is levels is equal to the number of included publications.
164. **study_and_pubs_plotready:** Publication-ready study and publicationID, derived by combining **study_studyID_pubready** and **study_publicationID_pubready**, separated by a newline ‘(’tidy_metadata.R’). Multiple publications from the same study are grouped in this way. Factor variable whose number is levels is equal to the number of included studies.
165. **cointerventions_other_name:** Other, non-antibiotic cointerventions. This variable was created by grepping ‘intervention_name’ to distinguish study arms that are identical other than having an additional cointervention (e.g. probiotic alongside the antibiotic) because they have not been distinguished elsewhere. Character variable.
166. **study_block:** Block to which the intervention arm belongs. This variable was created by grepping ‘’intervention_name’ to distinguish study arms that are identical other than belonging to a different block (e.g. weight block, sub-trial) because they have not been distinguished elsewhere. Character variable.
167. **cluster_none:** Cluster to which the observation belongs, down to the level of study days, used for aggregating within-study observations. This ‘control cluster’ should equal the number of observations (rows) in the dataframe. Derived by concatenating strings from other columns that define the observation. Character variable.
168. **cluster_intervoutcome:** Cluster to which the observation belongs, down to the level of resistance target (i.e. intervention-outcome combination or effect), used for aggregating within-study observations. Derived by concatenating strings from other columns that define the observation. Character variable.
169. **cluster_studyarm:** Cluster to which the observation belongs, down to the level of study arm, used for aggregating within-study observations. Derived by concatenating strings from other columns that define the observation. Character variable.
170. **cluster_organism:** Cluster to which the observation belongs, down to the level of microorganism, used for aggregating within-study observations. Derived by concatenating strings from other columns that define the observation. Character variable.
171. **cluster_resistancetarget:** Cluster to which the observation belongs, down to the level of resistance target, used for aggregating within-study observations. Derived by concatenating strings from other columns that define the observation. Character variable.
172. **cluster_sampletype:** Cluster to which the observation belongs, down to the level of sample type, used for aggregating within-study observations. Derived by concatenating strings from other columns that define the observation. Character variable.
173. **yi_totalresistancedeterminants:** Standardised mean difference (accounting for heterogeneity) in the absolute abundance of resistance between the intervention/exposure and control group. Calculated by applying metafor’s ‘escalc’ function to the ‘intervention_mean_totalresistancedeterminants’, ‘control_mean_totalresistancedeterminants’, ‘intervention_sd_totalresistancedeterminants, ’control_sd_totalresistancedeterminants’, ‘intervention_reps_totalresistancedeterminants’ , and ‘control_reps_totalresistancedeterminants’ columns. Numeric variable.
174. **vi_totalresistancedeterminants:** Sampling variance of the standardised mean difference (accounting for heterogeneity) in the absolute abundance of resistance between the intervention/exposure and control group. Calculated by applying metafor’s ‘escalc’ function to the ‘intervention_mean_totalresistancedeterminants’, ‘control_mean_totalresistancedeterminants’, ‘intervention_sd_totalresistancedeterminants, ’control_sd_totalresistancedeterminants’, ‘intervention_reps_totalresistancedeterminants’ , and ‘control_reps_totalresistancedeterminants’ columns. Numeric variable.
175. **ni_totalresistancedeterminants:** Sample size of the standardised mean difference (accounting for heterogeneity) in the absolute abundance of resistance between the intervention/exposure and control group. Calculated by applying metafor’s ‘escalc’ function to the ‘intervention_mean_totalresistancedeterminants’, ‘control_mean_totalresistancedeterminants’, ‘intervention_sd_totalresistancedeterminants, ’control_sd_totalresistancedeterminants’, ‘intervention_reps_totalresistancedeterminants’ , and ‘control_reps_totalresistancedeterminants’ columns. Numeric variable.
176. **yi_totaldeterminants:** Standardised mean difference (accounting for heterogeneity) in the absolute abundance of total determinants between the intervention/exposure and control group. Calculated by applying metafor’s ‘escalc’ function to the ‘intervention_mean_totaldeterminants’, ‘control_mean_totaldeterminants’, ‘intervention_sd_totaldeterminants, ’control_sd_totalrdeterminants’, ‘intervention_reps_totaldeterminants’ , and ‘control_reps_totaldeterminants’ columns. Numeric variable.
177. **vi_totaldeterminants:** Sampling variance of the standardised mean difference (accounting for heterogeneity) in the absolute abundance of total determinantsv between the intervention/exposure and control group. Calculated by applying metafor’s ‘escalc’ function to the ‘intervention_mean_totaldeterminants’, ‘control_mean_totaldeterminants’, ‘intervention_sd_totaldeterminants, ’control_sd_totalrdeterminants’, ‘intervention_reps_totaldeterminants’ , and ‘control_reps_totaldeterminants’ columns. Numeric variable.
178. **ni_totaldeterminants:** Sample size of the standardised mean difference (accounting for heterogeneity) in the absolute abundance of total determinants between the intervention/exposure and control group. Calculated by applying metafor’s ‘escalc’ function to the ‘intervention_mean_totaldeterminants’, ‘control_mean_totaldeterminants’, ‘intervention_sd_totaldeterminants, ’control_sd_totalrdeterminants’, ‘intervention_reps_totaldeterminants’ , and ‘control_reps_totaldeterminants’ columns. Numeric variable.
179. **yi_logitpropres:** Standardised mean difference (accounting for heterogeneity) in the logit-transformed relative abundance of resistance determinants between the intervention/exposure and control group. Calculated by applying metafor’s ‘escalc’ function to the ‘intervention_mean_logitpropres’, ‘control_mean_logitpropres’, ‘intervention_sd_logitpropres, ’control_sd_logitpropres’, ‘intervention_reps_logitpropres’ , and ‘control_reps_logitpropres’ columns. Numeric variable.
180. **vi_logitpropres:** Sampling variance of the standardised mean difference (accounting for heterogeneity)in the logit-transformed relative abundance of resistance determinants between the intervention/exposure and control group. Calculated by applying metafor’s ‘escalc’ function to the ‘intervention_mean_logitpropres’, ‘control_mean_logitpropres’, ‘intervention_sd_logitpropres, ’control_sd_logitpropres’, ‘intervention_reps_logitpropres’ , and ‘control_reps_logitpropres’ columns. Numeric variable.
181. **ni_logitpropres:** Sample size of the standardised mean difference (accounting for heterogeneity) in the logit-transformed relative abundance of resistance determinants between the intervention/exposure and control group. Calculated by applying metafor’s ‘escalc’ function to the ‘intervention_mean_logitpropres’, ‘control_mean_logitpropres’, ‘intervention_sd_logitpropres, ’control_sd_logitpropres’, ‘intervention_reps_logitpropres’ , and ‘control_reps_logitpropres’ columns. Numeric variable.
182. **yi_arcsinpropres:** Standardised mean difference (accounting for heterogeneity) in the arcsin-transformed relative abundance of resistance determinants between the intervention/exposure and control group. Calculated by applying metafor’s ‘escalc’ function to the ‘intervention_mean_arcsinpropres’, ‘control_mean_arcsinpropres’, ‘intervention_sd_arcsinpropres, ’control_sd_arcsinpropres’, ‘intervention_reps_arcsinpropres’ , and ‘control_reps_arcsinpropres’ columns. Numeric variable.
183. **vi_arcsinpropres:** Sampling variance of the standardised mean difference (accounting for heterogeneity) in the arcsin-transformed relative abundance of resistance determinants between the intervention/exposure and control group. Calculated by applying metafor’s ‘escalc’ function to the ‘intervention_mean_arcsinpropres’, ‘control_mean_arcsinpropres’, ‘intervention_sd_arcsinpropres, ’control_sd_arcsinpropres’, ‘intervention_reps_arcsinpropres’ , and ‘control_reps_arcsinpropres’ columns. Numeric variable.
184. **ni_arcsinpropres:** Sample size of the standardised mean difference (accounting for heterogeneity) in the arcsin-transformed relative abundance of resistance determinants between the intervention/exposure and control group. Calculated by applying metafor’s ‘escalc’ function to the ‘intervention_mean_arcsinpropres’, ‘control_mean_arcsinpropres’, ‘intervention_sd_arcsinpropres, ’control_sd_arcsinpropres’, ‘intervention_reps_arcsinpropres’ , and ‘control_reps_arcsinpropres’ columns. Numeric variable.
185. **unitlevel_id:** Arbitrary ID (row number) used to account for residual variance as a random effect.
186. **outcome_time_days:** Time column produced by log10-transforming and combining the ‘outcome_timesinceintervention_start_days’ (during) and ‘’outcome_timesinceintervention_end_days’ (after) columns into one, so that they may be specified as a moderator using the same model term in the meta-regressions. Combining them in this way despite them representing different things is acceptable because ‘during’ and ‘after’ data are modelled in separate meta-regressions. Numeric variable expressed on a log10 scale.

##### 1.4.2 Results of individual studies

Supplementary Information 7: Aggregating estimates of the effect at the study level

To produce the four types of forest plots, we first split the meta-analytical dataset into measures of antibiotic resistance in faeces and measures of antibiotic resistance in environmental samples, and within these two groupings outcomes measures pre-antibiotic administration (i.e. baseline measurements) or post-antibiotic administration.

We then fit intercept-only random effects models using each of the four datasets as a basis for aggregating effects at the study level and summarising the overall effect across in these four distinct datasets. The random effects used in these models were the same as those in the meta-analysis (see ‘Methods: Synthesis methods: Meta-analysis’ in the main text). The response variables in these models were the SMDHs representing the effect on the absolute abundance of resistance determinants, weighted by their associated sampling variances. We also fit equivalent models using SMDHs representing the effect on relative abundance of resistance determinants (logit-transformed and arcsine square root-transformed), and their associated sampling variances, as this was the form in which we had initially intended to analyse the review outcomes (Supplementary Box 1).

Finally, the variance-covariance matrix of these models were used to aggregate effects at the study level in a second, equal-effects model^24^ for each subset (‘H1 Aggregated Faeces Pre’ model, ‘H1 Aggregated Faeces Post’ model, ‘H1 Aggregated Enviro Pre’ model & ‘H1 Aggregated Enviro Post’ model). The results were then presented in forest plots built using the ‘forest’ function of the metafor R package, to facilitate summarising the results of individual studies. As well as allowing us to gain an initial quantitative summary of study-level effects pre- and post-antibiotic administration (and create an associated forest plot), these intercept-only models allowed us to estimate overall effects, after accounting for various forms of non-independence between effect sizes from the same study/study arm. This allowed us straightforwardly to validate that any effects detected post-administration did not exist pre-administration, as well as to compare the effects in faeces with that in environmental samples. It also allowed us to explicitly compare the effects on absolute abundance with those on relative abundance, allowing readers to make their own judgement about this protocol deviation (Supplementary Box 1).

##### 1.4.3 Sensitivity analyses

Supplementary Information 8: Sensitivity testing methods

**Leave-one-out-analysis**

In our first set of sensitivity analyses, we reran our meta-analytical models leaving one of the previously-included studies out at a time (i.e. a leave-one-out meta-analysis). This allowed us to get a sense of whether individual studies were disproportionately affecting our overall estimates (hypothesis 1) and the modelled moderating effect of time (hypothesis 2).

**Removal and correction of potentially problematic studies and effect sizes**

In our second set of sensitivity analyses, we removed or corrected studies/study arms whose effect sizes we thought might be problematic before rerunning our models.

We removed the following groups of potentially problematic studies/study arms:

1. **Studies that were not randomised or not correctly randomised** as non-randomised studies are very likely to be confounded and therefore generally offer lower quality evidence for a causative effect^25^.
2. **Studies for which we had to extract data.** As already mentioned, extracting means and SDs is less precise than calculating them from IPD. The 3 studies we excluded here were the Lethbridge_Chlortetsul study^26^, the Lethbridge_Tyl study^11^ and the ’TexasA&M - Ceftiofur”^10^ study..
3. **Studies for which we were not able to fully account for pen** as SMDHs were potentially overestimated for these studies^7^. The 2 studies we removed were the “Texas A&M - Ceftiofur”^10^ study and the “Lethbridge - Tylosin”^11^ study.
4. **Studies for which we were not able to account for blocking** as SMDHs were potentially underestimated for these studies^7^. The 5 studies we removed were the “USMARC - Chlortetracycline” study^27^, “TexasA&M - Ceftiofur; Tulathromycin” study^28^, “USMARC - Tylosin” study^6^, ” WKU - Tylosin” study^29^, and the “TexasA&M - Tylosin with probiotic”^9^ study.

In another analysis, we corrected for shared-control non-independence where necessary (i.e. for studies in which several intervention groups shared one control group) by penalising the sample sizes of intervention groups (dividing by the number of interventions with which it shared a control), before re-calculating SMDHs and rerunning our analyses.

**Assumed correlation between effect sizes from the same study**

Since all studies included contributed multiple effect sizes, our third set of sensitivity analyses tested the effect of varying the assumed correlation between effect sizes within a study. This was done by refitting our candidate models using variance-covariance matrices that assumed differing degrees of correlation between effect sizes from the same study^30,31^.

#### 1.5 Reporting bias assessment

Supplementary Information 9: Reporting bias methods

To test for small-study effects for each of the four review outcomes, we first tested for sample size publication bias by testing for a relationship between the effective sample size and the residuals of the four models used in our primary analysis. Effective sample size was chosen over variance to account for unbalanced sampling in the control and treatment groups underpinning the standardised mean differences. This was done by creating a funnel plot of model residuals against effective sample size and visually inspecting it. However, given the unreliability of visual methods for detecting sample size publication bias, especially where there is significant heterogeneity^32–34^, we also conducted formal tests by fitting adjusted Egger’s regressions of the relationship between effect size and the square root of the inverse of effective sample size. Its inverse was taken such that larger values represent lower accuracy (as for variance), and the square root transformation was applied to normalise the distribution of sample sizes^21^.

We also investigated potential time-lag publication bias (i.e. decline effects) by plotting and regressing of the mean-centered year of publication against effect size. Where we found evidence of these forms of publication bias in regressions, we subsequently intended to perform a additional sensitivity analyses consisting of adding these moderators to our candidate models and observing how results changed.

#### 1.6 Certainty assessment

Supplementary Information 10: Certainty assessment methods

We assessed certainty in the size of the four pooled effects estimated during meta-analysis. To begin our implementation of the GRADE approach^35^, each of the four bodies of evidence underpinning the four models was assigned an initial “high” level of certainty, given that all but one study included in the quantitative synthesis (meta-analysis) was (correctly) randomised^36^. We then considered whether to downgrade certainty for each body of evidence according to the five standard downgrading criteria, and subsequently whether to upgrade certainty based on three standard upgrading criteria.

The five standard downgrading criteria considered were risk of bias, inconsistency, indirectness, imprecision, and publication bias. For risk of bias, we considered there to be ‘Serious limitations’ and downgraded certainty by one level if most studies contributing to the estimate had ‘Some concerns’ or a ‘High risk’ of bias overall. For inconsistency, we considered there to be ‘Serious limitations’ and downgraded by one level if there was quantitative evidence of (residual) between-study heterogeneity in the model based on the I${}^{2}$ test for heterogeneity. For indirectness, we considered there to be ‘Serious limitations’ and downgraded certainty by one level if the body of evidence was biased towards a restricted version of our eligibility criteria. For imprecision, we considered there to be ‘Serious limitations’ and downgraded by one level if the confidence interval of the estimate included an SMDH of 0 (no effect). For publication bias, we considered there to be ‘Serious limitations’ and downgraded by one level if we had found quantitative evidence of publication bias in the body of evidence (see Reporting biases).

The three standard upgrading domains considered were large effects, dose–response, plausible confounding. For large effects, we upgraded certainty by one level if the SMDH was 0.8 or above, which is considered ‘large’ effect according to the rules of thumb established by this effect size’s originator^37^. For dose-response, we upgraded certainty by one level if we had found evidence of a dose-response relationship in the alternative analysis using dose as a moderator (note we did this as a single analysis across all post-antibiotic administration, rather than separately for ‘during’ and ‘after’ subsets, to maximise power; see ‘Subgroup and other analysis’). For plausible confounding, we upgraded by one level in the exceptional case that all confounding would be expected to make the observed relationships stronger (e.g. positive overall effects more positive, direction of the relationship with timing of sampling more pronounced).

This process resulted in a final certainty of evidence score following the GRADE schema - high (starting score), moderate, low, or very low - which was included in a standard Summary of Findings table. This table also included illustrative comparative differences; examples of the mean absolute abundance of antibiotic resistance in pairs of control (without antibiotic) and intervention (with antibiotic) samples that had a similar SMDH to the point estimates from our models. For the overall effects (Outcomes 1 and 3), we illustrated the comparative difference with an individual pair of measurements from control and intervention arms in one of the included studies, which had a similar SMDH to the pooled SMDH from the meta-analysis. For the moderating effect of time (Outcomes 2 and 4), we illustrated the comparative difference with a pair of all longitudinal measurements from control and intervention arms in one of the included studies, which had a similar slope of SMDH to the pooled slope of SMDH from the meta-regression.

### 2 Supplementary Results

#### 2.1 Study selection

Supplementary Figure 1: PRISMA2020 flow diagram of the systematic review process for the results of our main, English-language searches of key bibliographic databases. *Specific databases searched via the Web of Science platform were SCI-EXPANDED, SSCI, A&HCI, CPCI-S, CPCI-SSH, ESCI. †Searched in English.

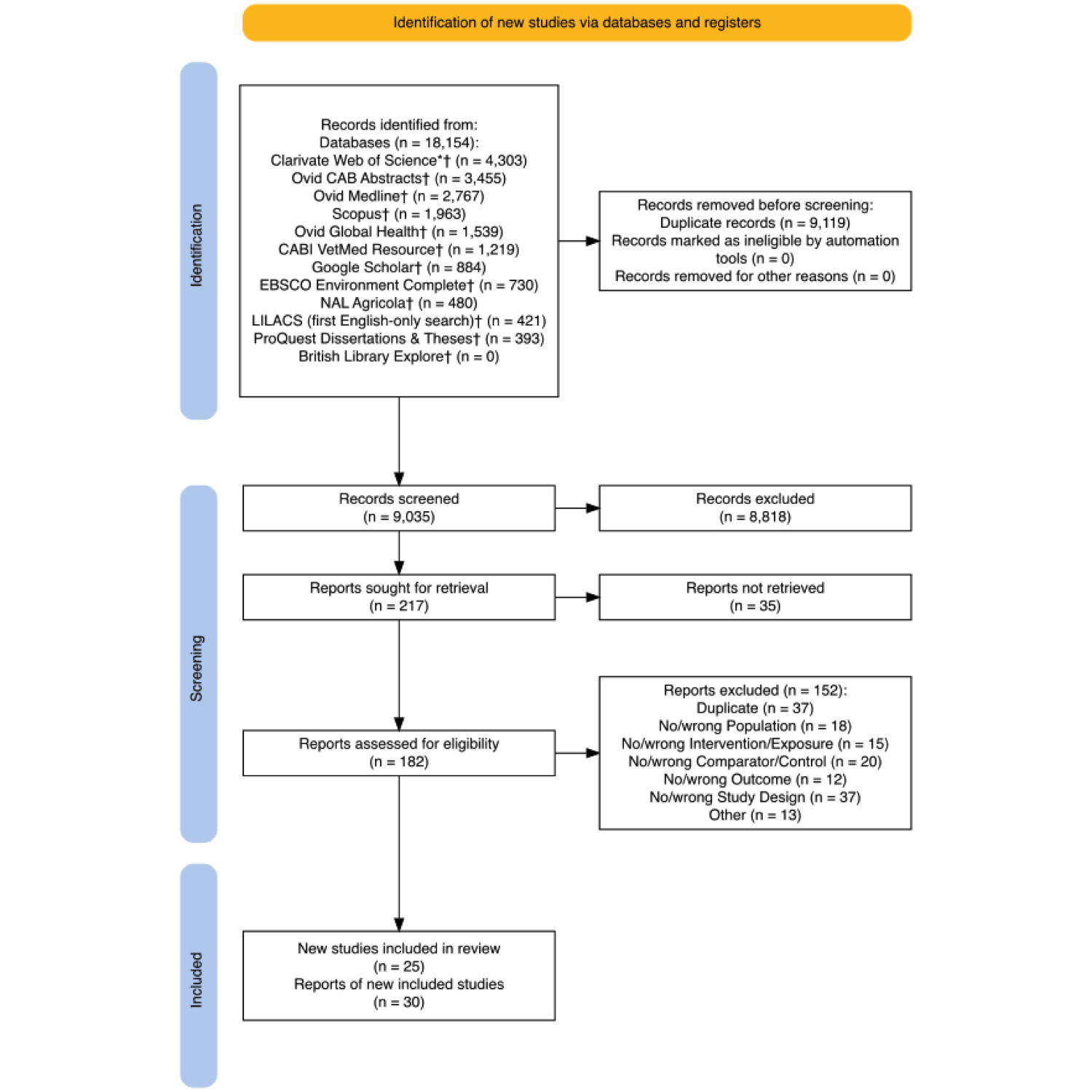

Supplementary Figure 2: PRISMA2020 flow diagram of the systematic review process for the results of our 2022 update of the English-language searches in the three bibliographic databases that produced the most results in our initial searches. *Specific databases searched via the Web of Science platform were SCI-EXPANDED, SSCI, A&HCI, CPCI-S, CPCI-SSH, ESCI. †Searched in English. Records removed for other reasons are records published before 2020, which should have been captured by the previous searches but escaped deduplication because of re-indexing by the bibliographic database providers.

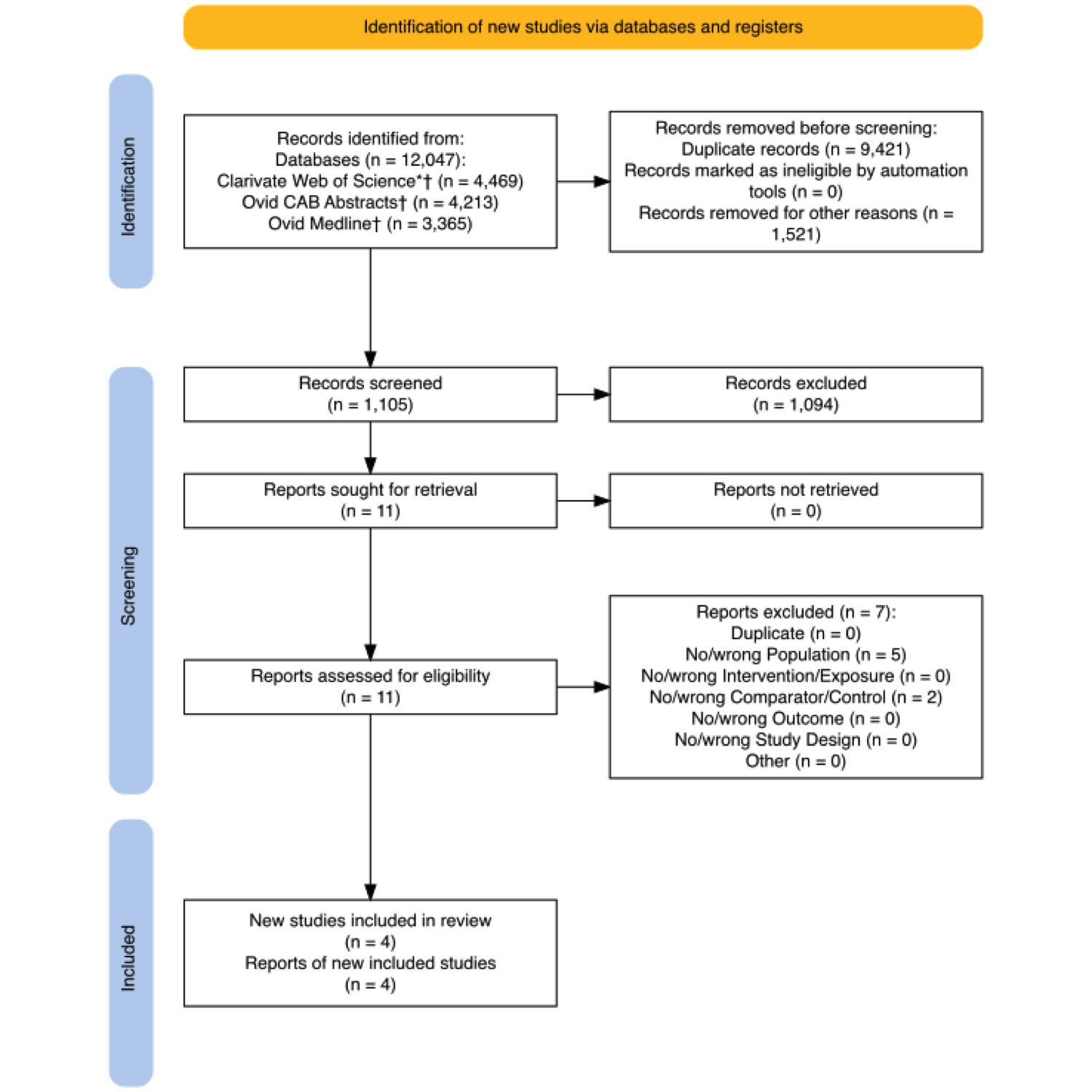

Supplementary Figure 3: PRISMA2020 flow diagram of the systematic review process for the results of our 2024 update of the English-language searches in the three bibliographic databases that produced the most results in our initial searches. *Specific databases searched via the Web of Science platform were SCI-EXPANDED, SSCI, A&HCI, CPCI-S, CPCI-SSH, ESCI. †Searched in English. Records removed for other reasons are records published before 2022, which should have been captured by the previous searches but escaped deduplication because of re-indexing by the bibliographic database providers

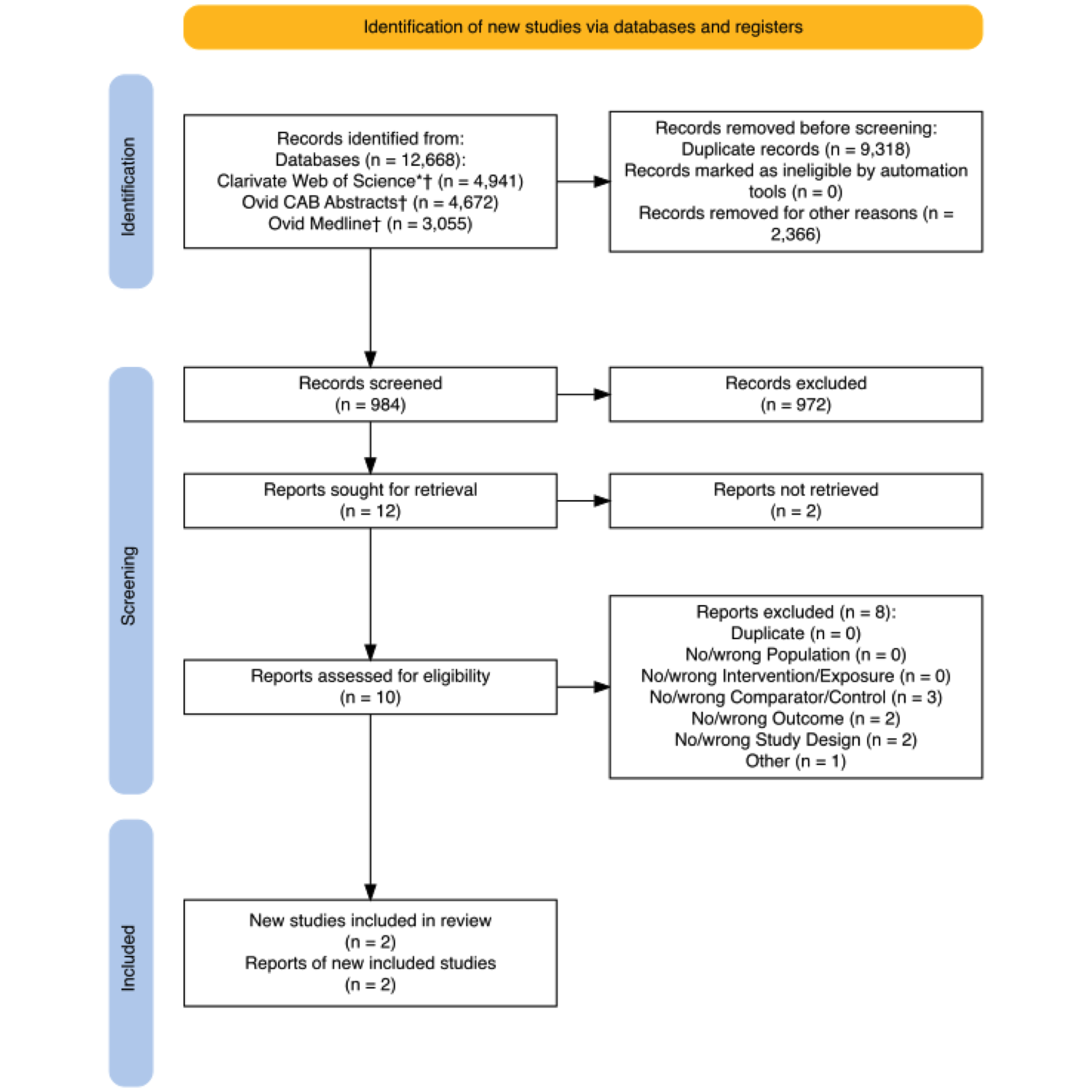

Supplementary Figure 4: PRISMA2020 flow diagram of the systematic review process for the results of our 2025 update of the English-language searches in the three bibliographic databases that produced the most results in our initial searches. *Specific databases searched via the Web of Science platform were SCI-EXPANDED, SSCI, A&HCI, CPCI-S, CPCI-SSH, ESCI. †Searched in English. Records removed for other reasons are records published before 2024, which should have been captured by the previous searches but escaped deduplication because of re-indexing by the bibliographic database providers

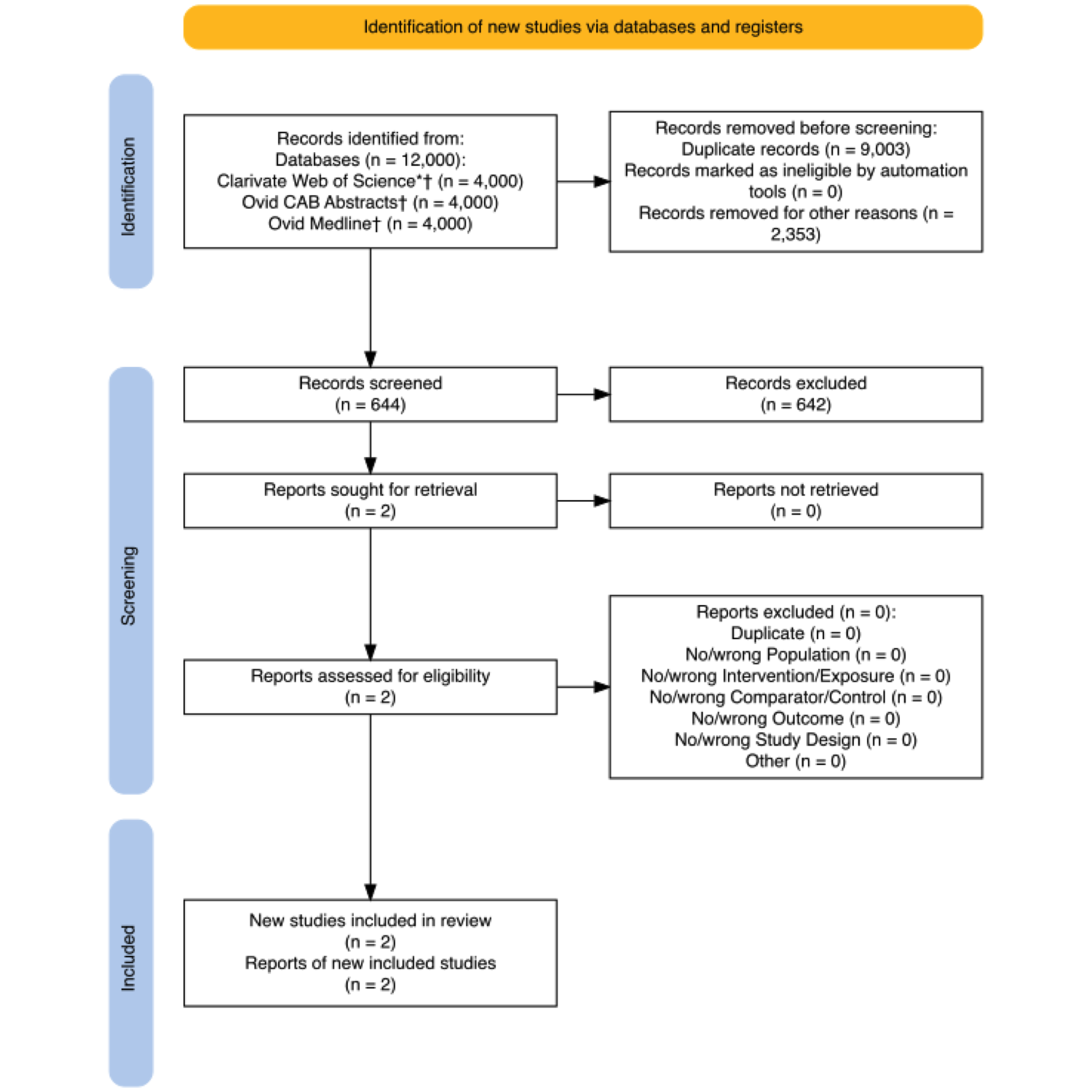

Supplementary Figure 5: PRISMA2020 flow diagram of the systematic review process for the results of our additional English- and Spanish-language searches of Latin American databases. †Searched in English. ‡Searched in Spanish.

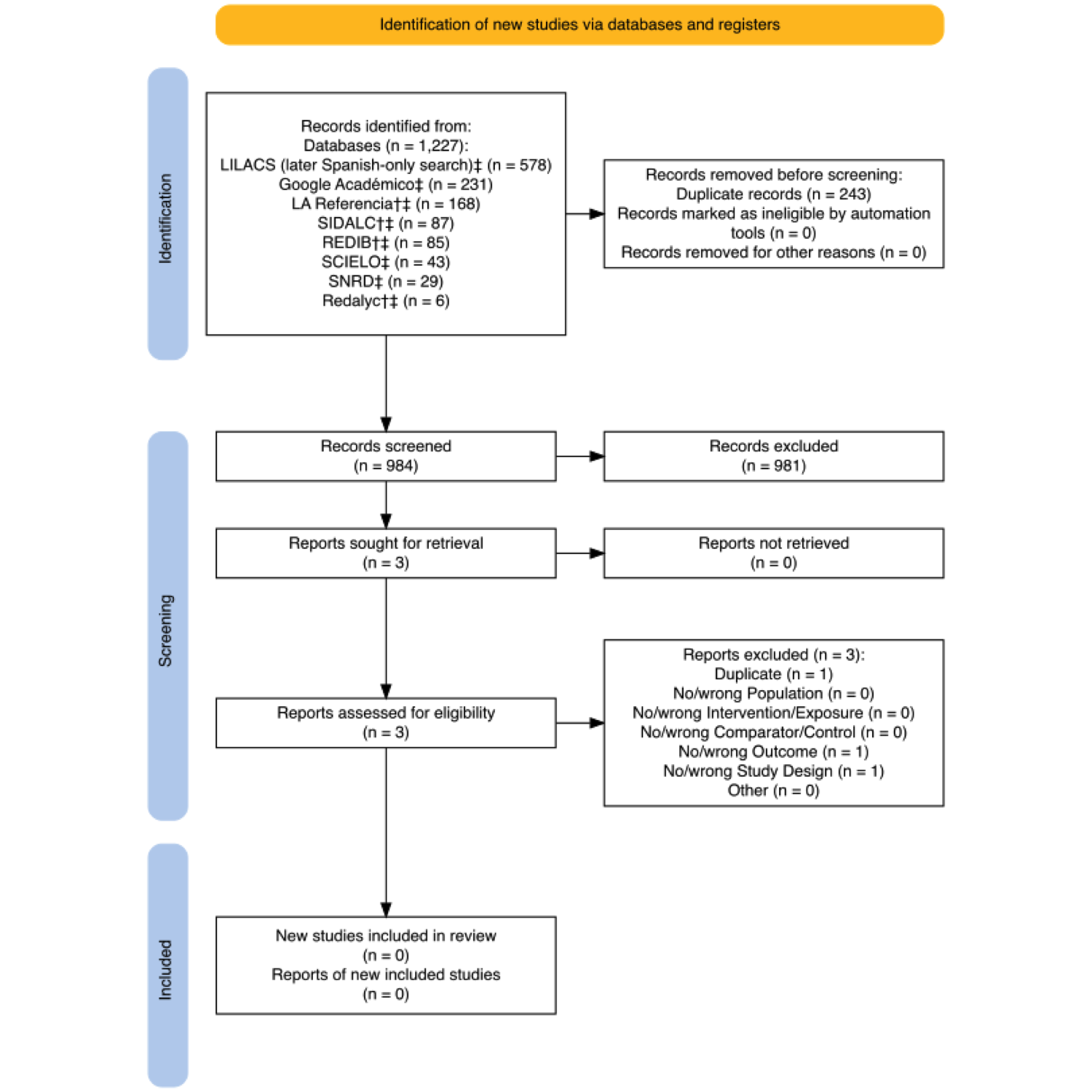

Supplementary Table 3: Reasons for excluding studies at full text stage

| **no.** | **authors** | **year** | **title** | **doi** | **PMID** | **final_exclusion_reason** | **final_exclusion_reason_elaboration** | **further_elaboration** |
| --- | --- | --- | --- | --- | --- | --- | --- | --- |
| 1 | Agga, G. E.; Schmidt, J. W.; Arthur, T. M. | 2016 | Effects of in-feed chlortetracycline prophylaxis in beef cattle on animal health and antimicrobialresistant Escherichia coli | 10.1128/aem.01928-16 | 27736789 | Duplicate | Duplicate record. |  |
| 2 | Agga, Getahun E.; Schmidt, John W.; Arthur, Terrance M. | 2016 | Effects of in-feed chlortetracycline prophylaxis of beef cattle on animal health and antimicrobial-resistant Escherichia coli | 10.1128/aem.01928-16 | 27736789 | Duplicate | Duplicate record. |  |
| 3 | Alali, W. Q.; Scott, H. M.; Norby, B.; Gebreyes, W.; Loneragan, G. H. | NA | Quantification of the Bla(CMY-2) in Feces from Beef Feedlot Cattle Administered Three Different Doses of Ceftiofur in a Longitudinal Controlled Field Trial | 10.1089/fpd.2009.0271 | 19622032 | Duplicate | Duplicate record. |  |
| 4 | Alexander, T. W.; Yanke, L. J.; Topp, E.; Olson, M. E.; Read, R. R.; Morck, D. W.; McAllister, T. A. | NA | Effect of subtherapeutic administration of antibiotics on the prevalence of antibiotic-resistant Escherichia coli bacteria in feedlot cattle | 10.1128/aem.00489-08 | 18502931 | Duplicate | Duplicate record. |  |
| 5 | Benedict, K. M.; Gow, S. P.; Booker, C. W.; McAllister, T. A.; Morley, P. S. | NA | ASSOCIATIONS BETWEEN ANTIMICROBIAL USE AND ANTIMICROBIAL RESISTANCE IN ESCHERICHIA COLI SAMPLED FROM INDIVIDUAL FEEDLOT CATTLE | 10.1371/journal.pone.0143995 | 26633649 | Duplicate | Duplicate record. | Published in PLOS One. |
| 6 | Catry, B.; Dewulf, J.; Maes, D.; Pardon, B.; Callens, B.; Vanrobaeys, M.; Opsomer, G.; de Kruif, A.; Haesebrouck, F. | NA | Effect of Antimicrobial Consumption and Production Type on Antibacterial Resistance in the Bovine Respiratory and Digestive Tract | 10.1371/journal.pone.0146488 | 26820134 | Duplicate | Duplicate record. |  |
| 7 | Checkley, Sylvia Lee | 2008 | The epidemiology of antimicrobial resistance in fecal <i>Escherichia coli</i> isolates of feedlot cattle in western Canada | NA | NA | Duplicate | Duplicate record. |  |
| 8 | Derycke, J.; Boivin, R. | 1982 | SELECTION AND PERSISTENCE OF CALF FECAL ESCHERICHIA-COLI SIMULTANEOUSLY RESISTANT TO TETRACYCLINE, STREPTOMYCIN AND SULFONAMIDES AFTER LONG-TERM ADMINISTRATION OF OXYTETRACYCLINE | NA | NA | Duplicate | Duplicate record. |  |
| 9 | Dodd, C. C.; Renter, D. G.; Shi, X. R.; Alam, M. J.; Nagaraja, T. G.; S; erson, M. W. | NA | Prevalence and Persistence of Salmonella in Cohorts of Feedlot Cattle | 10.1089/fpd.2010.0777 | 21381942 | Duplicate | Duplicate record. |  |
| 10 | Gronvold, A. M. R.; Mao, Y. J.; L'Abee-Lund, T. M.; Sorum, H.; Sivertsen, T.; Yannarell, A. C.; Mackie, R. I. | NA | Fecal microbiota of calves in the clinical setting: Effect of penicillin treatment | 10.1016/j.vetmic.2011.05.040 | 21705160 | Duplicate | Duplicate record. |  |
| 11 | Hocher, A. | 1983 | Prevalence of salmonellae and enteropathogenic E. coli and antibiotic-resistant E. coli in the intestinal flora of weaned calves of Dissertation [German] | NA | NA | Duplicate | Duplicate record. |  |
| 12 | Hocher, A. | 1981 | Occurrence of salmonellae and enteropathogenic E. coli and of antibiotic-resistant E. coli in the intestinal flora of weaned calves. of thesis [German] | NA | NA | Duplicate | Duplicate record. |  |
| 13 | Inglis, G. D.; Morck, D. W.; McAllister, T. A.; Entz, T.; Olson, M. E.; Yanke, L. J.; Read, R. R. | NA | Temporal prevalence of antimicrobial resistance in Campylobacter spp. from beef cattle in Alberta feedlots | 10.1128/AEM.02830-05 | 16751519 | Duplicate | Duplicate record. |  |
| 14 | Kanwar, N.; Scott, H. M.; Norby, B.; Loneragan, G. H.; Vinasco, J.; McGowan, M.; Cottell, J. L.; Chengappa, M. M.; Bai, J. F.; Boerlin, P. | 2013 | Effects of ceftiofur and chlortetracycline treatment strategies on antimicrobial susceptibility and on tet(A), tet(B), and blaCMY-2 resistance genes among E. coli isolated from the feces of feedlot cattle | 10.1371/journal.pone.0080575 | 24260423 | Duplicate | Duplicate record. |  |
| 15 | Kanwar, Neena; Scott, Harvey M. | 2013 | Treatment strategies impacting ceftiofur resistance among enteric bacteria in cattle | NA | NA | Duplicate | Duplicate record. |  |
| 16 | Keijser, B. J. F.; Agamennone, V.; van den Broek, T. J.; Caspers, M.; van de Braak, A.; Bomers, R.; Havekes, M.; Schoen, E.; van Baak, M.; Mioch, D.; Bomers, L.; Montijn, R. C. | NA | Dose-dependent impact of oxytetracycline on the veal calf microbiome and resistome | 10.1186/s12864-018-5419-x | 30660184 | Duplicate | Duplicate record. |  |
| 17 | Kyselkova, M.; Jirout, J.; Chronakova, A.; Vrchotova, N.; Bradley, R.; Schmitt, H.; Elhottova, D. | NA | Cow excrements enhance the occurrence of tetracycline resistance genes in soil regardless of their oxytetracycline content | 10.1016/j.chemosphere.2013.08.058 | 24053942 | Duplicate | Duplicate record. |  |
| 18 | Lefebvre, B.; Diarra, M. S.; Gigure, K.; Roy, G.; Michaud, S.; Malouin, F. | 2005 | Antibiotic resistance and hypermutability of Escherichia coli O157 from feedlot cattle treated with growth-promoting agents | 10.4315/0362-028x-68.11.2411 | 16300081 | Duplicate | Duplicate record. |  |
| 19 | Lefebvre, Brigitte; Diarra, Moussa S.; Gigure, Karine; Roy, Gabriel; Michaud, Sophie; Malouini, Franis | 2005 | Antibiotic Resistance and Hypermutability of Escherichia coil O157 from Feed lot Cattle Treated with Growth-Promoting Agents | 10.4315/0362-028x-68.11.2411 | 16300081 | Duplicate | Duplicate record. |  |
| 20 | Levent, G.; Schlochtermeier, A.; Ives, S. E.; Norman, K. N.; Lawhon, S. D.; Loneragan, G. H.; Anderson, R. C.; Vinasco, J.; Scott, H. M. | NA | Population Dynamics of Salmonella enterica within Beef Cattle Cohorts Followed from Single-Dose Metaphylactic Antibiotic Treatment until Slaughter | 10.1128/AEM.01386-19 | 31519659 | Duplicate | Duplicate record. |  |
| 21 | McEwen, S. A.; Poppe, C.; Popa, M.; Werber, D.; O'Connor, A.; Dunlop, H. | 2000 | Therapeutic and in-feed antimicrobial use and resistance in fecal commensals of swine and cattle | NA | NA | Duplicate | Duplicate record. |  |
| 22 | Mirzaagha, P.; Louie, M.; Sharma, R.; Yanke, L. J.; Topp, E.; McAllister, T. A. | NA | Distribution and characterization of ampicillin-and tetracycline-resistant Escherichia coli from feedlot cattle fed subtherapeutic antimicrobials | 10.1186/1471-2180-11-78 | 21504594 | Duplicate | Duplicate record. |  |
| 23 | Mller, Hans Christian; Bibber-Krueger, Cadra L. Van; Ogunrinu, Olanrewaju J.; Amachawadi, Raghavendra G.; Scott, H. Morgan; Drouillard, James S. | 2018 | Effects of intermittent feeding of tylosin phosphate during the finishing period on feedlot performance, carcass characteristics, antimicrobial resistance, and incidence and severity of liver abscesses in steers | 10.1093/jas/sky166 | 29718254 | Duplicate | Duplicate record. |  |
| 24 | Morley, P. S.; Dargatz, D. A.; Hyatt, D. R.; Dewell, G. A.; Patterson, J. G.; Burgess, B. A.; Wittum, T. E. | NA | Effects of Restricted Antimicrobial Exposure on Antimicrobial Resistance in Fecal Escherichia coli from Feedlot Cattle | 10.1089/fpd.2010.0632 | 2103427 | Duplicate | Duplicate record. |  |
| 25 | Salaheen, S.; Kim, S. W.; Cao, H. L.; Wolfgang, D. R.; Hovingh, E.; Karns, J. S.; Haley, B. J.; Van Kessel, J. A. S. | NA | Antimicrobial Resistance Among Escherichia coli Isolated from Veal Calf Operations in Pennsylvania | 10.1089/fpd.2018.2530 | 30673353 | Duplicate | Duplicate record. |  |
| 26 | Schmidt, J. W.; Vikram, A.; Miller, E.; Jones, S. A.; Arthur, T. M. | NA | In-Feed Tylosin Phosphate Administration to Feedlot Cattle Minimally Affects Antimicrobial Resistance | 10.4315/0362-028X.JFP-19-342 | 31971461 | Duplicate | Duplicate record. |  |
| 27 | Schmidt, JW; Griffin, D; Kuehn, LA; ... | 2013 | Influence of therapeutic ceftiofur treatments of feedlot cattle on fecal and hide prevalences of commensal Escherichia coli resistant to expanded-spectrum | 10.1128/AEM.03592-12 | 23354706 | Duplicate | Duplicate record. |  |
| 28 | Sharma, R.; Larney, F. J.; Chen, J.; Yanke, L. J.; Morrison, M.; Topp, E.; McAllister, T. A.; Yu, Z. T. | NA | Selected Antimicrobial Resistance during Composting of Manure from Cattle Administered Sub-Therapeutic Antimicrobials | 10.2134/jeq2007.0638 | 19202027 | Duplicate | Duplicate record. |  |
| 29 | Sharma, R.; Munns, K.; Alexander, T.; Entz, T.; Mirzaagha, P.; Yanke, L. J.; Mulvey, M.; Topp, E.; McAllister, T. | NA | Diversity and distribution of commensal fecal Escherichia coli bacteria in beef cattle administered selected subtherapeutic antimicrobials in a feedlot setting | 10.1128/AEM.00704-08 | 18723654 | Duplicate | Duplicate record. |  |
| 30 | Thornton Smith, Ashley B.; Renter, David G. | 2017 | Epidemiology and Quinolone-Susceptibilities of <i>Salmonella </i> and <i>Campylobacter</i> in Feedlot Cattle | 10.1089/fpd.2017.2282 | 28768141 | Duplicate | Duplicate record. |  |
| 31 | Weinroth, M. D.; Martin, J. N.; Doster, E.; Geornaras, I.; Parker, J. K.; Carlson, C. R.; Metcalf, J. L.; Morley, P. S.; Belk, K. E. | NA | Investigation of tylosin in feed of feedlot cattle and effects on liver abscess prevalence, and fecal and soil microbiomes and resistomes | 10.1093/jas/skz306 | 31563955 | Duplicate | Duplicate record. |  |
| 32 | Weinroth, M. D.; Scott, H. M.; Norby, B.; Loneragan, G. H.; Noyes, N. R.; Rovira, P.; Doster, E.; Yang, X.; Woerner, D. R.; Morley, P. S.; Belk, K. E. | NA | Effects of Ceftiofur and Chlortetracycline on the Resistomes of Feedlot Cattle | 10.1128/AEM.00610-18 | 29728379 | Duplicate | Duplicate record. |  |
| 33 | Weinroth, Margaret; Belk, Keith E. | 2016 | Effects of antibiotic treatment strategies on feedlot cattle resistome and microbiome | NA | NA | Duplicate | Duplicate record. |  |
| 34 | Xu, S. W.; Sura, S.; Zaheer, R.; Wang, G.; Smith, A.; Cook, S.; Olson, A. F.; Cessna, A. J.; Larney, F. J.; McAllister, T. A. | NA | Dissipation of Antimicrobial Resistance Determinants in Composted and Stockpiled Beef Cattle Manure | 10.2134/jeq2015.03.0146 | 27065400 | Duplicate | Duplicate record. |  |
| 35 | Xu, ShanWei; Amarakoon, I. D.; Zaheer, R.; Smith, A.; Srinivas, Sura; Wang, G.; Reuter, T.; Zvomuya, F.; Cessna, A. J.; Larney, F. J.; McAllister, T. A. | 2018 | Dissipation of antimicrobial resistance genes in compost originating from cattle manure after direct oral administration or post-excretion fortification of antimicrobials | 10.1080/10934529.2017.1404337 | 29215973 | Duplicate | Duplicate record. |  |
| 36 | Xu, Shanwei; Amarakoon, Inoka D.; Zaheer, Rahat; Smith, Alanna; Sura, Srinivas; Wang, George; Reuter, Tim; Zvomuya, Francis; Cessna, Allan J.; Larney, Francis J.; McAllister, Tim A. | 2018 | Dissipation of antimicrobial resistance genes in compost originating from cattle manure after direct oral administration or post-excretion fortification of antimicrobials | 10.1080/10934529.2017.1404337 | 29215973 | Duplicate | Duplicate record. |  |
| 37 | Minihan, D.; O'Mahony, M.; Whyte, P. | 2006 | The antimicrobial resistance profiles of Campylobacter and E. coli O157 isolates from a cohort of clinically normal feedlot cattle | NA | NA | Duplicate | Duplicate record. |  |
| 38 | Morley, P. S.; Dargatz, D. A.; Hyatt, D. R.; Dewell, G. A.; Patterson, G.; Wittum, T. E.; Smith, R. A. | 2007 | Effects of rearing practices that restrict antimicrobial exposure in feedlot cattle on antimicrobial resistance in enteric bacteria | 10.1089/fpd.2010.0632 | 21034271 | Duplicate | Duplicate record. |  |
| 39 | Alexander, T. W.; Yanke, J. L.; Reuter, T.; Topp, E.; Read, R. R.; Selinger, B. L.; McAllister, T. A. | NA | Longitudinal characterization of antimicrobial resistance genes in feces shed from cattle fed different subtherapeutic antibiotics | 10.1186/1471-2180-11-19 | 21261985 | No/wrong Study Design | No baseline measure before administration of intervention antibiotic. | There are no faecal deposits from the intervention cattle before they have been administered antibiotics (i.e. no baseline measure). |
| 40 | Benedict, K. M.; Gow, S. P.; McAllister, T. A.; Booker, C. W.; Hannon, S. J.; Checkley, S. L.; Noyes, N. R.; Morley, P. S. | NA | Antimicrobial Resistance in Escherichia coli Recovered from Feedlot Cattle and Associations with Antimicrobial Use | 10.1371/journal.pone.0143995 | 26633649 | No/wrong Study Design | Cross-sectional study of different farms. |  |
| 41 | Berge, AC; Hancock, DD; Sischo, WM; ... | 2010 | Geographic, farm, and animal factors associated with multiple antimicrobial resistance in fecal Escherichia coli isolates from cattle in the western United States | 10.2460/javma.236.12.1338 | 20550450 | No/wrong Study Design | Cross-sectional study of different farms. |  |
| 42 | Carlson, C.; Martin, J.; Woerner, D.; Morley, P.; Geornaras, G.; Belk, K. | 2017 | Characterization of Enterococci, Salmonella Spp., and Generic Escherichia Coli Isolated From the Feces of Cattle Fed Rations with and without Tylosin Phosphate | 10.221751/rmc2016.014 | NA | No/wrong Study Design | No baseline measure before administration of intervention antibiotic. |  |
| 43 | Catry, B.; Dewulf, J.; Maes, D.; Pardon, B.; Callens, B.; Vanrobaeys, M.; Opsomer, G.; Kruif, A. de; Haesebrouck, F. | 2016 | Effect of antimicrobial consumption and production type on antibacterial resistance in the bovine respiratory and digestive tract | 10.1371/journal.pone.0146488 | 26820134 | No/wrong Study Design | Cross-sectional study of different farms. |  |
| 44 | Checkley, S. L.; Campbell, J. R.; Chirino-Trejo, M.; Janzen, E. D.; McKinnon, J. J. | 2008 | Antimicrobial resistance in generic fecal Escherichia coli obtained from beef cattle on arrival at the feedlot and prior to slaughter, and associations with volume of total individual cattle antimicrobial treatments in one western Canadian feedlot | NA | 18505198 | No/wrong Study Design | Although there are two interventions arms and a control arm, the individual antibiotic treatments (cointerventions) are uncontrolled across the experiment making deducing the effect of the interventions difficulty (especially given individual treatments likely to covary with study arm) |  |
| 45 | Chen, J.; Fluharty, F. L.; St-Pierre, N.; Morrison, M.; Yu, Z. | NA | Technical note: Occurrence in fecal microbiota of genes conferring resistance to both macrolide-lincosamide-streptogramin B and tetracyclines concomitant with feeding of beef cattle with tylosin | 10.2527/jas.2007-0705 | 18469042 | No/wrong Study Design | No baseline measure before administration of intervention antibiotic. |  |
| 46 | English, A.; Echeverry, A.; Sarturi, J.; Opheim, T.; Nightingale, K.; Miller, M.; Brashears, M. | 2017 | Antimicrobial Resistance Patterns of Enterococcus Isolated from Feedlot Cattle after Feeding Direct-Fed Microbials in Diets with and Without Tylosin | 10.22175/rmc2017.129 | NA | No/wrong Study Design | No baseline measure before administration of intervention antibiotic. |  |
| 47 | Erwerth, W.; Mehlhorn, G.; Methling, W. | NA | Dynamics of drug resistance of the bacterial flora of the air of calf houses [particularly micrococci and Escherichia coli] | NA | 3303655 | No/wrong Study Design | No baseline measure before administration of intervention antibiotic. |  |
| 48 | Gedek, B.; Schal, E. | 1976 | The occurrence of drug-resistant colicinogenic strains of E. coli in the faecal flora of calves. 2. Colicin types formed with simultaneous resistance to antibiotics and sulphonamides | NA | 773050 | No/wrong Study Design | Unsystematic sampling from a trial for descriptive study. |  |
| 49 | Gedek, B.; Schl, E. | 1976 | Zum Auftreten von arzneimittelresistenten, colicinogenen E. coliStmmen in der Faecalflora von Klbern: I. Mitteilung: Hufigkeit unter Antibiotikaverabreichung in therapeutischer und nutritiver Dosierung | NA | NA | No/wrong Study Design | Unsystematic sampling from a trial for descriptive study. |  |
| 50 | Gedek, B.; Schl, E. | 1976 | Zum Auftreten von arzneimittelresistenten, colicinogenen E. coliStmmen in der Faecalflora von Klbern: II. Mitteilung: Art der gebildeten Colicintypen bei gleichzeitigem Vorliegen von Resistenz gegenber Antibiotika und Sulfonamiden | NA | NA | No/wrong Study Design | Unsystematic sampling from a trial for descriptive study. |  |
| 51 | Gosling, R. J.; Mueller-Doblies, D.; Martelli, F.; Nunez-Garcia, J.; Kell, N.; Rabie, A.; Wales, A. D.; Davies, R. H. | NA | Observations on the distribution and persistence of monophasic Salmonella Typhimurium on infected pig and cattle farms | 10.1016/j.vetmic.2018.10.032 | 30473358 | No/wrong Study Design | Cross-sectional study of different farms. |  |
| 52 | Gunn, G. J.; Hall, M.; Low, J. C. | NA | Comparison of antibiotic resistance for Escherichia coli populations isolated from groups of diarrhoeic and control calves | 10.1016/s1090-0233(02)00244-7 | NA | No/wrong Study Design | Cross-sectional study of different farms. |  |
| 53 | Inglis, GD; Morck, DW; McAllister, TA; Entz, T; ... | 2006 | Temporal prevalence of antimicrobial resistance in Campylobacter spp. from beef cattle in Alberta feedlots | 10.1128/AEM.02830-05 | 16751519 | No/wrong Study Design | No baseline measure before administration of intervention antibiotic. | First sample taken 'within 1 day after arrival at the feedlot' which is likely after feeding (chlorteteracycline and oxytetracycline) begins and the injection of oxytetracycline upon arrival |
| 54 | Jacob, M. E.; Fox, J. T.; Narayanan, S. K.; Drouillard, J. S.; Renter, D. G.; Nagaraja, T. G. | NA | Effects of feeding wet corn distillers grains with solubles with or without monensin and tylosin on the prevalence and antimicrobial susceptibilities of fecal foodborne pathogenic and commensal bacteria in feedlot cattle | NA 10.2527/jas.2007-0091 | 18192558 | No/wrong Study Design | No baseline measure before administration of intervention antibiotic. | "Several limitations to our study include the small number of sample collection days, low statistical power to detect differences, and lack of baseline prevalence and antimicrobial susceptibility estimates" |
| 55 | Larsen, H. E.; Larsen, J. L. | NA | [Occurrence and distribution of drug resistant Escherichia coli in feces from cattle and swine] | NA | 4580598 | No/wrong Study Design | Cross-sectional study of different farms. |  |
| 56 | Layton, H. W.; Langworth, B. F.; Jarolmen, H.; Simkins, K. L. | 1975 | Influence of chlortetracycline and chlortetracycline + sulfamethazine supplemented feed on the incidence, persistence and antibacterial susceptibility of Salmonella typhimurium in experimentally inoculated calves | 10.1111/j.1439-0450.1975.tb00611.x | NA | No/wrong Study Design | No baseline measure before administration of intervention antibiotic. | First sample taken at 1 day after administration to intervention group. |
| 57 | Lee, C.; Zaheer, R.; Munns, K.; Holman, D. B.; Domselaar, G. van; Zovoilis, A.; McAllister, T. A. | 2023 | Effect of antimicrobial use in conventional versus natural cattle feedlots on the microbiome and resistome | 10.3390/microorganisms11122982 | 38138126 | No/wrong Study Design | Cross-sectional study of different farms. |  |
| 58 | Massot, M.; Couffignal, C.; Clermont, O.; D'Humieres, C.; Chatel, J.; Plault, N.; Andremont, A.; Caron, A.; Mentre, F.; Denamur, E. | NA | Day-to-Day Dynamics of Commensal Escherichia coli in Zimbabwean Cows Evidence Temporal Fluctuations within a Host-Specific Population Structure | 10.1128/aem.00659-17 | 28411228 | No/wrong Study Design | No baseline measure before administration of intervention antibiotic. | First sample taken at 2 days after administration to intervention group. |
| 59 | Merle, R.; Weise, S.; Gorisek, L.; Baer, J.; Robe, C.; Friese, A.; Jensen, K. C. | 2023 | The therapy frequency of antibiotics and phenotypical resistance of Escherichia coli in calf rearing sites in Germany | 10.3389/fvets.2023.1152246 | 37275609 | No/wrong Study Design | Cross-sectional study of different farms. |  |
| 60 | Mirzaagha, P.; Louie, M.; Ranjana, Sharma; Yanke, L. J.; Topp, E.; McAllister, T. A. | 2011 | Distribution and characterization of ampicillin- and tetracycline-resistant Escherichia coli from feedlot cattle fed subtherapeutic antimicrobials | 10.1186/1471-2180-11-78 | 21504594 | No/wrong Study Design | No baseline measure before administration of intervention antibiotic. |  |
| 61 | Mollenkopf, Dixie Francis; Wittum, Thomas | 2017 | Epidemiology of -lactamase-producing <i>Enterobacteriaceae </i> in Humans and Livestock | NA | NA | No/wrong Study Design | Cross-sectional study of different farms. |  |
| 62 | Moncada M, Paola M. | 2017 | Efecto del uso de Lactobacillus salivarius como suplemento en la dieta final de ganado de carne en patrones de prevalencia y resistencia antimicrobiana de Campylobacter spp | NA | NA | No/wrong Study Design | No baseline measure before administration of intervention antibiotic. |  |
| 63 | Morley, PS; Dargatz, DA; Hyatt, DR; ... | 2011 | Effects of Restricted Antimicrobial Exposure on Antimicrobial Resistance in Fecal Escherichia coli from Feedlot Cattle | 10.1089/fpd.2010.0632 | 21034271 | No/wrong Study Design | Cross-sectional study of different farms. | We explored whether there were any pens within the feedlots studied that could be used in an eligible comparison but it was not possible. |
| 64 | Noyes, N. R.; Benedict, K. M.; Gow, S. P.; Waldner, C. L.; Reid-Smith, R. J.; Booker, C. W.; McAllister, T. A.; Morley, P. S. | NA | Modelling considerations in the analysis of associations between antimicrobial use and resistance in beef feedlot cattle | 10.1017/S0950268815002423 | 26541938 | No/wrong Study Design | Modelling study |  |
| 65 | Peak, N.; Knapp, C. W.; Yang, R. K.; Hanfelt, M. M.; Smith, M. S.; Aga, D. S.; Graham, D. W. | NA | Abundance of six tetracycline resistance genes in wastewater lagoons at cattle feedlots with different antibiotic use strategies | 10.1111/j.1462-2920.2006.01123.x | 17227419 | No/wrong Study Design | Cross-sectional study of different farms. |  |
| 66 | Rovira, P.; McAllister, T.; Lakin, S. M.; Cook, S. R.; Doster, E.; Noyes, N. R.; Weinroth, M. D.; Yang, X.; Parker, J. K.; Boucher, C.; Booker, C. W.; Woerner, D. R.; Belk, K. E.; Morley, P. S. | NA | Characterization of the Microbial Resistome in Conventional and "Raised Without Antibiotics" Beef and Dairy Production Systems | 10.3389/fmicb.2019.01980 | 31555225 | No/wrong Study Design | No baseline measure before administration of intervention antibiotic. |  |
| 67 | Sharma, Ranjana; Larney, Francis J.; Jing, Chen; Yanke, L. Jay; Morrison, Mark; Topp, Edward; McAllister, Tim A.; Zhongtang, Yu | 2009 | Selected Antimicrobial Resistance during Composting of Manure from Cattle Administered Sub-Therapeutic Antimicrobials | 10.2134/jeq2007.0638 | 19202027 | No/wrong Study Design | No baseline measure before administration of intervention antibiotic. | This study is in manure so would need a sample of the manure before antibiotic administration. |
| 68 | Vidovic, S.; Korber, D. R. | NA | Prevalence of Escherichia coli O157 in Saskatchewan cattle: Characterization of isolates by using random amplified polymorphic DNA PCR, antibiotic resistance profiles, and pathogenicity determinants | 10.1128/AEM.02791-05 | 16751550 | No/wrong Study Design | Cross-sectional study of different farms. | Point prevalence study. |
| 69 | Weinroth, M.; Rovira, P.; Yang, X.; Parker, J.; Morley, P. S.; Belk, K. E. | 2018 | Characterization of Antibiotic Resistance Transmission in the Beef Production System | 10.221751/rmc2018.132 | NA | No/wrong Study Design | No baseline measure before administration of intervention antibiotic. |  |
| 70 | Weinroth, Margaret D.; Scott, H. Morgan; Norby, Bo; Loneragan, Guy H.; Noyes, Noelle R.; Rovira, Pablo; Doster, Enrique; Xiang, Yang; Woerner, Dale R.; Morley, Paul S.; Belka, Keith E. | 2018 | Effects of Ceftiofur and Chlortetracycline on the Resistomes of Feedlot Cattle | 10.1128/AEM.00610-18 | 29728379 | No/wrong Study Design | No baseline measure before administration of intervention antibiotic. | Despite being described as 'Day 0', day 0 is actually after antibiotic administration: "Samples evaluated in this study were collected on day 0 and day 26, which was only 6 days after the final in-feed administration of CTC" |
| 71 | Weinroth, Margaret; Belk, Keith E. | 2019 | Shotgun Metagenomic and 16S rRNA Gene Sequencing Investigation of Antimicrobial Resistance in the Beef Supply Chain | NA | NA | No/wrong Study Design | No baseline measure before administration of intervention antibiotic. | See associated publication(s). |
| 72 | Weinroth, MD; Martin, JN; Doster, E; ... | 2019 | Investigation of tylosin in feed of feedlot cattle and effects on liver abscess prevalence, and fecal and soil microbiomes and resistomes | 10.1093/jas/skz306 | 31563955 | No/wrong Study Design | Cross-sectional study of different farms. |  |
| 73 | Woeriner, D. R.; Ransom, J. R.; Sofos, J. N.; Scanga, J. A.; Smith, G. C.; Belk, K. E. | 2006 | Preharvest processes for microbial control in cattle | NA | NA | No/wrong Study Design | No baseline measure before administration of intervention antibiotic. | "Samples were collected, within a time span of 10 days for all treatments, randomly from 25 cattle per pen, 12 to 48h before transport to a commercial slaughter facility." |
| 74 | Wu, R. B.; Alexander, T. W.; Li, J. Q.; Munns, K.; Sharma, R.; McAllister, T. A. | NA | Prevalence and diversity of class 1 integrons and resistance genes in antimicrobial-resistant Escherichia coli originating from beef cattle administered subtherapeutic antimicrobials | 10.1111/j.1365-2672.2011.05066.x | 21645183 | No/wrong Study Design | No baseline measure before administration of intervention antibiotic. | Tetr E. coli isolated from rectal faecal samples that were collected on day 169 and 197 (steers have been administrated antimicrobials for 169 and 197 days) were chosen and includedin this study. |
| 75 | Xu, Shanwei; Sura, Srinivas; Zaheer, Rahat; George, Wang; Smith, Alanna; Cook, Shaun; Olson, Andrew F.; Cessna, Allan J.; Larney, Francis J.; McAllister, Tim A. | 2016 | Dissipation of Antimicrobial Resistance Determinants in Composted and Stockpiled Beef Cattle Manure | 10.2134/jeq2015.03.0146 | 27065400 | No/wrong Study Design | No baseline measure before administration of intervention antibiotic. | This study is in manure so would need a sample of the manure before antibiotic administration. |
| 76 | Gow, Sheryl Pamela | 2007 | Investigation of antimicrobial resistance and antimicrobial use in Western Canadian cow -calf herds | NA | NA | No/wrong Study Design | Cross-sectional study of different farms. |  |
| 77 | Vikram, A.; Rovira, P.; Agga, G. E.; Arthur, T. M.; Bosilevac, J. M.; Wheeler, T. L.; Morley, P. S.; Belk, K. E.; Schmidt, J. W. | NA | Impact of "Raised without Antibiotics" Beef Cattle Production Practices on Occurrences of Antimicrobial Resistance | 10.1128/AEM.01682-17 | 28887421 | No/wrong Study Design | Cross-sectional study of different farms. | We explored whether there were any pens within the feedlots studied that could be used in an eligible comparison but it was not possible. |
| 78 | Alexlander, T. W.; Reuter, T.; Sharma, R.; Yanke, L. J.; Topp, E.; McAllister, T. A. | NA | Longitudinal Characterization of Resistant Escherichia coli in Fecal Deposits from Cattle Fed Subtherapeutic Levels of Antimicrobials | 10.1128/AEM.00944-09 | 19801481 | No/wrong Study Design | No baseline measure before administration of intervention antibiotic. | There are no faecal deposits from the intervention cattle before they have been administered antibiotics (i.e. no baseline measure). |
| 79 | Ripamonti, B.; Tirloni, E.; Stella, S.; Bersani, C.; Agazzi, A.; Maroccolo, S.; Savoini, G. | 2013 | Effects of a species-specific probiotic formulation on multiresistant Escherichia coli isolates from the gut of veal calves | NA | NA | No/wrong Population | Unweaned cattle. |  |
| 80 | Aust, MO; Godlinski, F; Travis, GR; Hao, X; ... | 2008 | Distribution of sulfamethazine, chlortetracycline and tylosin in manure and soil of Canadian feedlots after subtherapeutic use in cattle | 10.1016/j.envpol.2008.03.011 | 18440678 | No/wrong Population | There is no distinct population because the faeces of all cattle are mixed into one massive deposit. |  |
| 81 | Berge, A. C. B.; Moore, D. A.; Sischo, W. M. | NA | Prevalence and antimicrobial resistance patterns of Salmonella enterica in preweaned calves from dairies and calf ranches | 10.2460/ajvr.67.9.1580 | 10.2460/ajvr.67.9.1580 | No/wrong Population | Unweaned cattle. |  |
| 82 | Berge, Anna Catharina Bjornsdotter; Sischo, William M. | 2004 | Spatial, temporal and management-specific factors influencing antibiotic resistance and carbohydrate fermentation patterns in bovine enteric <i>Escherichia coli</i> and the clinical consequences of limiting antibiotic use in pre-weaned calves | NA | NA | No/wrong Population | Unweaned cattle. |  |
| 83 | Beyi, A. F.; Brito-Goulart, D.; Hawbecker, T.; Ruddell, B.; Hassall, A.; Dewell, R.; Dewell, G.; Sahin, O.; Zhang, Q.; Plummer, P. J. | 2021 | Enrofloxacin Alters Fecal Microbiota and Resistome Irrespective of Its Dose in Calves | 10.3390/microorganisms9102162 | 34683483 | No/wrong Population | Dairy cattle. |  |
| 84 | Beyi, A. F.; Brito-Goulart, D.; Hawbecker, T.; Slagel, C.; Ruddell, B.; Hassall, A.; Dewell, R.; Dewell, G.; Sahin, O.; Zhang, Q.; Plummer, P. J. | 2021 | Danofloxacin Treatment Alters the Diversity and Resistome Profile of Gut Microbiota in Calves | 10.3390/microorganisms9102023 | 34683343 | No/wrong Population | Dairy cattle. |  |
| 85 | De Rycke, J.; Boivin, R. | 1982 | [Selection and persistence in the slaughter calf fecal flora of Escherichia coli simultaneously resistant to tetracycline, streptomycin and sulfonamides following long term treatment with oxytetracycline] | NA | NA | No/wrong Population | Unweaned cattle. |  |
| 86 | Embden, J. D. A. van; Groothuis, D. G.; Soedirman, N. | 1982 | The effect of virginiamycin in animal feed on drug resistance in the streptococcal flora of calves | NA | NA | No/wrong Population | Unweaned cattle. |  |
| 87 | Gay, E.; Bour, M.; Cazeau, G.; Jarrige, N.; Martineau, C.; Madec, J. Y.; Haenni, M. | NA | Antimicrobial Usages and Antimicrobial Resistance in Commensal Escherichia coli From Veal Calves in France: Evolution During the Fattening Process | 10.3389/fmicb.2019.00792 | 31031738 | No/wrong Population | Unweaned cattle. |  |
| 88 | Gedek, B.; Schal, E. | 1976 | The occurrence of drug-resistant colicinogenic strains of E. coli in the faecal flora of calves. 1. Frequency with antibiotics given in therapeutic and nutritive doses | NA | 773050 | No/wrong Population | Unweaned cattle. | Veal? |
| 89 | Gibbons, J. F. | 2011 | Antimicrobial resistance and antimicrobial use in farm-animal production in Ireland | NA | NA | No/wrong Population | Dairy cattle. |  |
| 90 | Goulart, D. B.; Beyi, A. F.; Wu, Z.; Adiguzel, M. C.; Schroeder, A.; Singh, K.; Xu, C.; Ocal, M. M.; Dewell, R.; Dewell, G. A.; Plummer, P. J.; Zhang, Q.; Sahin, O. | 2022 | Effect of Danofloxacin Treatment on the Development of Fluoroquinolone Resistance in Campylobacter jejuni in Calves | 10.3390/antibiotics11040531 | NA | No/wrong Population | Dairy cattle. |  |
| 91 | Goulart, D. B.; Beyi, A. F.; Wu, Z.; Adiguzel, M. C.; Wilson, S.; Xu, C.; Pang, J.; Dewell, R.; Dewell, G. A.; Plummer, P. J.; Zhang, Q.; Sahin, O. | 2022 | Influence of Single Dose Enrofloxacin Injection on Development of Fluoroquinolone Resistance in Campylobacter jejuni in Calves | 10.3390/antibiotics11101407 | NA | No/wrong Population | Dairy cattle. |  |
| 92 | Grnvold, Anne-Mette R.; Mao, Yuejian; LAbe-Lund, Trine M.; Srum, Henning; Sivertsen, Tore; Yannarell, Anthony C.; Mackie, Roderick I. | 2011 | Fecal microbiota of calves in the clinical setting: Effect of penicillin treatment | 10.1016/j.vetmic.2011.05.040 | 21705160 | No/wrong Population | Dairy cattle. |  |
| 93 | Hirsh, D. C.; Wiger, N. | 1978 | The effect of tetracycline upon the spread of bacterial resistance from calves to man | 10.2527/jas1978.4651437x | NA | No/wrong Population | Unweaned cattle. |  |
| 94 | Howe, K.; Linton, A. H.; Osborne, A. D. | 1976 | A longitudinal study of Escherichia coli in cows and calves with special reference to the distribution of O-antigen types and antibiotic resistance | 10.1111/j.1365-2672.1976.tb04181.x | 780335 | No/wrong Population | Dairy cattle. |  |
| 95 | Keijser, B. J. F.; Agamennone, V.; Broek, T. J. van den; Caspers, M.; Braak, A. van de; Bomers, R.; Havekes, M.; Schoen, E.; Baak, M. van; Mioch, D.; Bomers, L.; Montijn, R. C. | 2019 | Dose-dependent impact of oxytetracycline on the veal calf microbiome and resistome | 10.1186/s12864-018-5419-x | 30660184 | No/wrong Population | Unweaned cattle. |  |
| 96 | Kyselkov, Martina; Jirout, Ji; Chrokov, Alica; Vrchotov, Nadda; Bradley, Robert; Schmitt, Heike; Elhottov, Dana | 2013 | Cow excrements enhance the occurrence of tetracycline resistance genes in soil regardless of their oxytetracycline content | 10.1016/j.chemosphere.2013.08.058 | 24053942 | No/wrong Population | Dairy cattle. |  |
| 97 | Linton, A. H.; Howe, K.; Osborne, A. D. | 1975 | The effects of feeding tetracycline, nitrovin and quindoxin on the drug-resistance of coli-aerogenes bacteria from calves and pigs | 10.1111/j.1365-2672.1975.tb00530.x | 1095540 | No/wrong Population | Unweaned cattle. |  |
| 98 | Liu, Jinxin; Call, Douglas R. | 2016 | The role of excreted antibiotics in the establishment of persistent on-farm reservoirs of antibiotic-resistant bacteria | NA | NA | No/wrong Population | Dairy cattle. |  |
| 99 | Pereira, R. V.; Siler, J. D.; Cummings, K. J.; Davis, M. A.; Warnick, L. D. | NA | Effect of heifer-raising practices on E-coli antimicrobial resistance and Salmonella prevalence in heifer raisers | 10.1017/S0950268815000357 | 25904042 | No/wrong Population | Dairy cattle. |  |
| 100 | Schonecker, L.; Schnyder, P.; Overesch, G.; Schupbach-Regula, G.; Meylan, M. | 2019 | Associations between antimicrobial treatment modalities and antimicrobial susceptibility in Pasteurellaceae and E. coli isolated from veal calves under field conditions | 10.1016/j.vetmic.2019.07.015 | 31500731 | No/wrong Population | Unweaned cattle. | Veal? |
| 101 | Shawver, S.; Wepking, C.; Ishii, S.; Strickland, M. S.; Badgley, B. D. | 2021 | Application of manure from cattle administered antibiotics has sustained multi-year impacts on soil resistome and microbial community structure | 10.1016/j.soilbio.2021.108252 | NA | No/wrong Population | Dairy cattle. | See Wepking et al. (2019) 10.1111/ele.13390 |
| 102 | Agga, G. E.; Schmidt, J. W.; Arthur, T. M. | NA | Antimicrobial-Resistant Fecal Bacteria from Ceftiofur-Treated and Nonantimicrobial-Treated Comingled Beef Cows at a Cow-Calf Operation | 10.1089/mdr.2015.0259 | 36910206 | No/wrong Intervention/Exposure | No distinct exposure/intervention group. A subgroup of the 'ceftiofur-treated' group that received no other antimicrobials can theoretically be compared to the 'antimicrobial-free' control group, but it is lumped in to a larger 'ceftiofur-treated' group with ceftiofur-treated animals that also receive other antimicrobials, so it is not possible to distinguish the effect of ceftiofur. |  |
| 103 | Anderson, M. A.; Whitlock, J. E.; Harwood, V. J. | NA | Diversity and distribution of Escherichia coli genotypes and antibiotic resistance phenotypes in feces of humans, cattle, and horses | 10.1128/AEM.01029-06 | 16950903 | No/wrong Intervention/Exposure | No antimicrobial intervention is defined. |  |
| 104 | Aslam, M.; Stanford, K.; McAllister, T. A. | NA | Characterization of antimicrobial resistance and seasonal prevalence of Escherichia coli O157:H7 recovered from commercial feedlots in Alberta, Canada | 10.1111/j.1472-765X.2010.02798.x | 20102509 | No/wrong Intervention/Exposure | There is no antibiotic intervention - the study compares different feedlots. |  |
| 105 | Edrington, TS; MacDonald, JC; Farrow, RL; ... | 2010 | Influence of wet distiller's grains on prevalence of Escherichia coli O157: H7 and Salmonella in feedlot cattle and antimicrobial susceptibility of generic Escherichia coli | 10.1089/fpd.2009.0417 | 20132029 | No/wrong Intervention/Exposure | Distiller's grains exposure without quantification of virginiamycin content. | Distillers grains sometimes contain virginiamycin but this is not quantified and explicitly explored here as in later included publication by Edrington et al. (2014) |
| 106 | Grispoldi, L.; Bertero, F.; Franceschini, S.; Mastrosimone, F.; Sechi, P.; Iulietto, M. F.; Ceccarelli, M.; Cenci-Goga, B. T. | 2017 | Prevalence and characterisation of shigatoxigenic Escherichia coli isolated from beef cattle fed with prebiotics | 10.4081/ijfs.2017.6958 | 29564239 | No/wrong Intervention/Exposure | No antibiotic intervention/exposure. | Prebiotic is the intervention. |
| 107 | Humphry, R. W.; Gunn, G. J.; Fenlon, D.; Low, J. C. | 2005 | Variations in antimicrobial resistance: a longitudinal study of E. coli populations from cattle | NA | NA | No/wrong Intervention/Exposure | No antibiotic intervention/exposure. | Unclear whether no intervention/exposure or no comparator/control but given is on a outside herd in Scotland assume they aren't receiving antibiotics and no intervention/exposure most likely. There is only one group besides |
| 108 | Klotz, P.; Higgins, P. G.; Schaubmar, A. R.; Failing, K.; Leidner, U.; Seifert, H.; Scheufen, S.; Semmler, T.; Ewers, C. | NA | Seasonal Occurrence and Carbapenem Susceptibility of Bovine Acinetobacter baumannii in Germany | 10.3389/fmicb.2019.00272 | 30853949 | No/wrong Intervention/Exposure | No distinct exposure/intervention group. |  |
| 109 | Knox, Natalie Cecile; Krause, Denis | 2013 | An exploration of ecological concepts in the context of antimicrobial resistance and the use of phytochemical compounds within the ruminant gut microbiome | NA | NA | No/wrong Intervention/Exposure | No antibiotic intervention/exposure. | Phytochemical compounds. |
| 110 | Minihan, D.; O'Mahony, M.; Whyte, P.; Fanning, S.; Fitzsimons, T.; Collins, J. D. | 2006 | The antimicrobial resistance profiles of Campylobacter and E. coli O157 isolates from a cohort of clinically normal feedlot cattle | NA | NA | No/wrong Intervention/Exposure | No distinct exposure/intervention group. | There are antibiotics given but not in a controlled way; there is no defined intervention group. |
| 111 | Nesemeier, B.; Ekiri, A. B.; L; blom, D.; Doetkott, D.; Khaitsa, M. L. | 2015 | Prevalence and antimicrobial resistance of Salmonella enterica shed from range and feedlot cattle from post-weaning to slaughter | NA | NA | No/wrong Intervention/Exposure | No antibiotic intervention/exposure. | None mentioned at least. |
| 112 | Pilaszek, J.; Truszczynski, M. | 1980 | A POSSIBILITY OF ACQUIRING AND LOOSING RESISTANCE TO ANTIBIOTICS IN BOVINE MYCOPLASMA STRAINS .2. ACQUIRING AND LOOSING RESISTANCE TO CHLORAMPHENICOL, OXYTETRACYCLINE, NEOMYCIN AND STREPTOMYCIN | NA | NA | No/wrong Intervention/Exposure | No antibiotic intervention/exposure. | In-vitro study of Mycoplasma strains. |
| 113 | Salaheen, S.; Kim, SeonWoo; Cao, HuiLin; Wolfgang, D. R.; Hovingh, E.; Karns, J. S.; Haley, B. J.; Kessel, J. A. S. van | 2019 | Antimicrobial resistance among Escherichia coli isolated from veal calf operations in Pennsylvania | 10.1089/fpd.2018.2530 | 30673353 | No/wrong Intervention/Exposure | No antibiotic intervention/exposure. | None mentioned at least. |
| 114 | Wells, JE; Shackelford, SD; Berry, ED; ... | 2009 | Prevalence and Level of Escherichia coli O157:H7 in Feces and on Hides of Feedlot Steers Fed Diets with or without Wet Distillers Grains with Solubles | 10.4315/0362-028x-72.8.1624 | 19722393 | No/wrong Intervention/Exposure | Distiller's grains exposure without quantification of virginiamycin content. | Distillers grains |
| 115 | Werckenthin, C.; Seidl, S.; Riedl, J.; Kiossis, E.; Wolf, G.; Stolla, R.; Kaaden, O. R. | NA | Escherichia coli isolates from young calves in Bavaria: In vitro susceptibilities to 14 anti-microbial agents | 10.1046/j.1439-0450.2002.00538.x | NA | No/wrong Intervention/Exposure | No antibiotic intervention/exposure. | None mentioned at least. |
| 116 | Alexander, T. W.; Jin, X.; Li, Q.; Cook, S.; McAllister, T. A. | NA | Characterization of tetracycline resistance genes in Escherichia coli isolated from feedlot cattle administered therapeutic or subtherapeutic levels of tetracycline | 10.1139/cjm-2012-0660 | 23586754 | No/wrong Comparator/Control | No intervention antibiotic-free comparator group. | Chlortetracycline is compared with chlortetracycline with oxytetracycline and since both groups receive a tetracycline this is not an eligible control. |
| 117 | Berge, A. C. B.; Atwill, E. R.; Sischo, W. M.; Menzies, F. D.; Reid, S. W. J. | 2001 | Assessing dynamics of antibiotic resistance in faecal Escherichia coli and in young calves using cluster analysis techniques | 10.1016/S0167-5877(03)00191-0 | 14519339 | No/wrong Comparator/Control | No antibiotic-free comparator group: |  |
| 118 | Caproni Junior, L.; Umehara, O.; Costa, E. O.; Benites, N. R.; Melville, P. A.; Raemdonck, D. | 2002 | Clinical efficacy of danofloxacin 18% in the treatment of naturally occurring infectious diseases in cattle | 10.1590/1808-1657v69n4p0072002 | NA | No/wrong Comparator/Control | No antibiotic-free comparator group. |  |
| 119 | Da Costa, Marcio Carvalho | 2010 | Prevalence and molecular characterization of <i>Clostridium difficile</i> in veal calves and feedlot beef cattle | NA | NA | No/wrong Comparator/Control | No antibiotic-free comparator group. |  |
| 120 | Davedow, T.; Narvaez-Bravo, C.; Zaheer, R.; S; erson, H.; Rodas-Gonzalez, A.; Klima, C.; Booker, C. W.; Hannon, S. J.; Bras, A. L.; Gow, S.; McAllister, T. | NA | Investigation of a Reduction in Tylosin on the Prevalence of Liver Abscesses and Antimicrobial Resistance in Enterococci in Feedlot Cattle | 10.3389/fvets.2020.00090 | 32185186 | No/wrong Comparator/Control | No antibiotic-free comparator group. | 'CON' group is actually a positive control that receives tylosin throughout the whole period, whilst the FIRST-78% and LAST-75% groups overlap with no sampling in the first 25% period where they do not. |
| 121 | Doster, E.; Pinnell, L. J.; Noyes, N. R.; Parker, J. K.; Anderson, C. A.; Booker, C. W.; Hannon, S. J.; McAllister, T. A.; Gow, S. P.; Belk, K. E.; Morley, P. S. | 2022 | Evaluating the effects of antimicrobial drug use on the ecology of antimicrobial resistance and microbial community structure in beef feedlot cattle | 10.3389/fmicb.2022.970358 | 36583056 | No/wrong Comparator/Control | No antibiotic-free comparator group. | Cohort study of antimicrobial-exposed cattle. |
| 122 | Erwin, S.; Foster, D. M.; Jacob, M. E.; Papich, M. G.; Lanzas, C. | NA | The effect of enrofloxacin on enteric Escherichia coli: Fitting a mathematical model to in vivo data | 10.1371/journal.pone.0228138 | 32004337 | No/wrong Comparator/Control | No antibiotic-free comparator group. | There is some real data in this modelling study but no enrofloxacin-free comparator group. |
| 123 | Ferguson, K. M.; Jacob, M. E.; Theriot, C. M.; Callahan, B. J.; Prange, T.; Papich, M. G.; Foster, D. M. | NA | Dosing Regimen of Enrofloxacin Impacts Intestinal Pharmacokinetics and the Fecal Microbiota in Steers | 10.3389/fmicb.2018.02190 | 30283418 | No/wrong Comparator/Control | No antibiotic-free comparator group. | "steers received either a single subcutaneous injection of 12.5 mg/kg enrofloxacin...or three subcutaneous injections of 5.0 mg/kg enrofloxacin administered once each day on 3 sequential days" |
| 124 | Foster, D. M.; Jacob, M. E.; Farmer, K. A.; Callahan, B. J.; Theriot, C. M.; Kathariou, S.; Cernicchiaro, N.; Prange, T.; Papich, M. G. | NA | Ceftiofur formulation differentially affects the intestinal drug concentration, resistance of fecal Escherichia coli, and the microbiome of steers | 10.1371/journal.pone.0223378 | 31584976 | No/wrong Comparator/Control | No antibiotic-free comparator group. | "...animals received one of two treatments (n = 6 steers per treatment)—subcutaneous injection of ceftiofur crystalline-free acid (CCFA; Excede®; 6.6 mg/kg) as a single dose at the base of the ear, or ceftiofur hydrochloride (CHCL; Excenel®; 2.2 mg/kg) subcutaneously in the neck every 24 hours for 3 treatments. |
| 125 | Gall; , JC; Hyatt, DR; Crupper, SS; ... | 2001 | Prevalence, antibiotic susceptibility, and diversity ofEscherichia coli O157: H7 isolates from a longitudinal study of beef cattle feedlots | 10.1128/AEM.67.4.1619-1627.2001 | 11282614 | No/wrong Comparator/Control | No antibiotic-free comparator group. |  |
| 126 | Halleran, J. L.; Callahan, B. J.; Jacob, M. E.; Sylvester, H. J.; Prange, T.; Papich, M. G.; Foster, D. M. | 2021 | Effects of danofloxacin dosing regimen on gastrointestinal pharmacokinetics and fecal microbiome in steers | 10.1038/s41598-021-90647-z | 34045586 | No/wrong Comparator/Control | No antibiotic-free comparator group. | "First, steers administered the lower dose of danofloxacin twice will have lower gastrointestinal concentrations when compared to the steers whom received the single, high dose of danofloxacin." |
| 127 | Halleran, Jennifer; Sylvester, Hannah; Jacob, Megan; Callahan, Benjamin; Baynes, Ronald; Foster, Derek | 2024 | Impact of florfenicol dosing regimen on the phenotypic and genotypic resistance of enteric bacteria in steers | 10.1038/s41598-024-55591-8 | 38418677 | No/wrong Comparator/Control | No antibiotic-free comparator group. | "Twelve steers were administered either two intramuscular (20 mg/kg q 48 h; n = 6) or a single subcutaneous dose (40 mg/kg, n = 6)." |
| 128 | Hordijk, J.; Mevius, D. J.; Kant, A.; Bos, M. E. H.; Gravel; , H.; Bosman, A. B.; Hartskeerl, C. M.; Heederik, D. J. J.; Wagenaar, J. A. | 2013 | Within-farm dynamics of ESBL/AmpC-producing Escherichia coli in veal calves: a longitudinal approach | 10.1093/jac/dkt219 | 23759504 | No/wrong Comparator/Control | No antibiotic-free comparator group. | "Multiple antimicrobial group treatments (of all animals within the herd) were administered on all farms (Figure 1)." |
| 129 | Hoyle, D.; Davison, H. C.; Knight, H.; Yates, C. M.; Dobay, O.; Gunn, G. J.; Amyes, S. G. B.; Woolhouse, M. E. J. | NA | Molecular characterisation of bovine faecal Escherichia coli shows persistence of defined ampicillin resistant strains and the presence of class 1 integrons on an organic beef farm | 10.1016/j.vetmic.2006.01.006 | 16490325 | No/wrong Comparator/Control | No antibiotic-free comparator group. |  |
| 130 | Kleinhenz, K. E.; Mollenkopf, D. F.; Heider, L. C.; Funk, J. A.; Wittum, T. E.; Smith, R. A. | 2010 | Impact of Ceftiofur Use on the Recovery of Salmonella and Resistant E. coli in Dairy and Beef Cattle Populations | 10.21423/aabppro20104152 | NA | No/wrong Comparator/Control | No antibiotic-free comparator group. | Seems to be studying ceftiofur-treated cattle with no clear comparator. |
| 131 | Larsen, H. E.; Larsen, J. L. | 1975 | Persistance of resistant Escherichia coli in a calf herd (Danish) | NA | 1094405 | No/wrong Comparator/Control | No antibiotic-free comparator group. |  |
| 132 | Lee, M. K.; Billington, S. J.; Joens, L. A. | 2004 | Potential virulence and antimicrobial susceptibility of Campylobacter jejuni isolates from food and companion animals | 10.1089/fpd.2004.1.223 | 15992284 | No/wrong Comparator/Control | No antibiotic-free comparator group. | Includes beef cattle that are probably antimicrobial treated without a clear comparator group. |
| 133 | Long, N. S.; Wells, J. E.; Berry, E. D.; Legako, J. F.; Woerner, D. R.; Loneragan, G. H.; Broadway, P. R.; Carroll, J. A.; Sanchez, N. C. B.; Fernando, S. C.; Bacon, C. M.; Helmuth, C. L.; Smock, T. M.; Manahan, J. L.; Hoffman, A. A.; Hales, K. E. | 2022 | Metaphylactic antimicrobial effects on occurrences of antimicrobial resistance in Salmonella enterica, Escherichia coli and Enterococcus spp. measured longitudinally from feedlot arrival to harvest in high-risk beef cattle | 10.1111/jam.15691 | 35766106 | No/wrong Comparator/Control | No antibiotic-free comparator group. | Although there's a sterile saline negative control group (CON), it actually received therapeutic antimicrobial treatment ad-libitum in response to symptoms of BRD. Whilst ethically understandable the ad-lib nature makes it difficult to compare with intervention groups as these cointerventions are not standardised across study arms. "Cattle were monitored daily for visual signs of BRD and were eligible for up to three additional therapeutic antimicrobial treatments as needed. Therapeutic treatment antimicrobials consisted of enrofloxacin (Baytril 100; Bayer Animal Health), tildiprosin (Zuprevo; Merck Animal Health) and danofloxacin (Advocin; Zoetis) with a 3 day, 7 day and 0 day post-treatment interval (PTI) respectively. " |
| 134 | Minihan, D.; Whyte, P.; O'Mahony, M.; Cowley, D.; O'Halloran, F.; Corcoran, D.; Fanning, S.; Collins, J. D. | NA | Phenotypic and genotypic anti-microbial resistance profiles of Campflobacters from untreated feedlot cattle and their environment | 10.1111/j.1439-0450.2006.00933.x | 16629986 | No/wrong Comparator/Control | No distinct exposure/intervention group. | There are antibiotics given but not in a controlled way; there is no defined intervention group. |
| 135 | Morris, C.; Wickramasingha, D.; Abdelfattah, E. M.; Pereira, R. V.; Okello, E.; Maier, G. | 2023 | Prevalence of antimicrobial resistance in fecal Escherichia coli and Enterococcus spp. isolates from beef cow-calf operations in northern California and associations with farm practices | 10.3389/fmicb.2023.1086203 | 36910206 | No/wrong Comparator/Control | No antibiotic-free comparator group. | Convenience sample of cattle that all likely received antimicrobials. |
| 136 | Noyes, N. R.; Yang, X.; Linke, L. M.; Magnuson, R. J.; Dettenwanger, A.; Cook, S.; Geornaras, I.; Woerner, D. E.; Gow, S. P.; McAllister, T. A.; Yang, H.; Ruiz, J.; Jones, K. L.; Boucher, C. A.; Morley, P. S.; Belk, K. E. | NA | Resistome diversity in cattle and the environment decreases during beef production | 10.7554/eLife.13195 | 26952213 | No/wrong Comparator/Control | No distinct antibiotic-free comparator group. | Cohort study of antimicrobial-exposed cattle. |
| 137 | Noyes, Noelle Robertson; Morley, Paul S. | 2016 | The epidemiology and ecology of antimicrobial use and resistance in North American beef production systems | NA | NA | No/wrong Comparator/Control | No distinct antibiotic-free comparator group. | Cohort study of antimicrobial-exposed cattle. |
| 138 | Rao, S.; Van Donkersgoed, J.; Bohaychuk, V.; Besser, T.; Song, X. M.; Wagner, B.; Hancock, D.; Renter, D.; Dargatz, D.; Morley, P. S. | NA | Antimicrobial Drug Use and Antimicrobial Resistance in Enteric Bacteria Among Cattle from Alberta Feedlots | 10.1089/fpd.2009.0400 | 19958100 | No/wrong Comparator/Control | No antibiotic-free comparator group. |  |
| 139 | Siegel, D.; Huber, W. G.; Enloe, F. | 1974 | Continuous non-therapeutic use of antibacterial drugs in feed and drug resistance of the Gram-negative enteric florae of food-producing animals | 10.1128/AAC.6.6.697 | 4615630 | No/wrong Comparator/Control | No antibiotic-free comparator group. | This one is a bit confusing as the authors speak of comparing animals with and without exposure to antibacterial drugs but it actually becomes clear that the 'without' group (Montana cattle) have 'minimal' not 'no' exposure "Farm animals continuously fed antibacterial drugs are compared to animals having minimal or no exposure to antibacterial agents." |
| 140 | Venter, B. J. | 1984 | The presence and persistence of transmissible drug resistance factors in the coliform intestinal flora of calves reared in an intensive system | NA | NA | No/wrong Comparator/Control | No antibiotic-free comparator group. | The intervention is a proprietary cocktail of antibiotics given to 'most' of the calves but this isn't compared with those who didn't receive it. |
| 141 | Webb, A. L.; Selinger, L. B.; Taboada, E. N.; Inglis, G. D. | NA | Subtype-Specific Selection for Resistance to Fluoroquinolones but Not to Tetracyclines Is Evident in Campylobacter jejuni Isolates from Beef Cattle in Confined Feeding Operations in Southern Alberta, Canada | 10.1128/AEM.02713-17 | 29352087 | No/wrong Comparator/Control | No antibiotic-free comparator group. |  |
| 142 | Lefebvre, B.; Malouin, F.; Roy, G.; Giguere, K.; Diarra, M. S. | NA | Growth performance and shedding of some pathogenic bacteria in feedlot cattle treated with different growth-promoting agents | 10.4315/0362-028x-69.6.1256 | 16786843 | No/wrong Outcome | Relative or absolute resistance (numerator+denominator) not assessed. | Includes a measure of MICs of some isolates but it seems this is based on antimicrobial susceptibility testing data already published in Lefebvre et al. (2005) and this paper focusses on effects of antibiotics on growth performance. |
| 143 | Edrington, T. S.; Looper, M. L.; Duke, S. E.; Callaway, T. R.; Genovese, K. J.; Anderson, R. C.; Nisbet, D. J. | NA | Effect of Ionophore Supplementation on the Incidence of Escherichia coli O157:H7 and Salmonella and Antimicrobial Susceptibility of Fecal Coliforms in Stocker Cattle | 10.1089/fpd.2006.3.284 | 16972777 | No/wrong Outcome | No measure of direct selection. | No measure of ionophore resistance (likely because no breakpoint) |
| 144 | Agga, G. E.; Galloway, H. O. | 2023 | Dynamics of extended-spectrum beta-lactamase-producing, third-generation cephalosporin-resistant and tetracycline-resistant Escherichia coli in feedlot cattle with or without tylosin administration | 10.1016/j.jfp.2023.100144 | 37597606 | No/wrong Outcome | Asssed indirect selection only. | Only indirect selection to cephalosporin and tetracycline assessed (antimicrobial susceptibility testing) - would need a measure of resistance to MLSBs (direct selection) to be eligible. |
| 145 | Bessegatto, J. A.; Paulino, L. R.; Lisboa, J. A. N.; Alfieri, A. A.; Montemor, C. H.; Medeiros, L. P.; Kobayashi, R. K. T.; Weese, J. S.; Costa, M. C. | NA | Changes in the fecal microbiota of beef cattle caused by change in management and the use of virginiamycin as a growth promoter | 10.1016/j.rvsc.2017.06.011 | 28675873 | No/wrong Outcome | Assessed effect on composition. |  |
| 146 | Bessegatto, Jos Antonio | 2016 | Mudanas na microbiota fecal de bovinos de corte causadas pelo confinamento e pela ingesto de virginiamicina | NA | NA | No/wrong Outcome | Assessed effect on composition. |  |
| 147 | Costa, M. C.; Staempfli, H.; Arroyo, L. G.; Pearl, D. L.; Weese, S. J. | NA | PREVALENCE OF TETRACYCLINE RESISTANCE GENES IN CLOSTRIDIUM DIFFICILE ISOLATED FROM CALVES PRIOR AND POST TREATMENT WITH OXYTETRACYCLINE | NA | NA | No/wrong Outcome | Assessed effect on prevalence of target organism across cattle. | Prevalence of C. difficile |
| 148 | Credille, Brent; Berghaus, Roy D.; Miller, Ella Jane; Credille, Allison; Schrag, Nora F. D.; Naikare, Hemant | 2023 | Antimicrobial Metaphylaxis and its Impact on Health, Performance, Antimicrobial Resistance, and Contextual Antimicrobial Use in High-Risk Beef Stocker Calves | 10.1093/jas/skad417 | 38126883 | No/wrong Outcome | Assessed only antibiotic resistance profiles via antimicrobial susceptibility testing, not relative abundance of specific resistance phenotypes. | Resistance patterns' reported descriptively for each study arm (Table 5) but not relative abundance of resistance to individual antibiotics for each study arm. |
| 149 | Dodd, CC; Renter, DG; Shi, X; Alam, MJ; ... | 2011 | Prevalence and Persistence of Salmonella in Cohorts of Feedlot Cattle | 10.1089/fpd.2010.0777 | 21381942 | No/wrong Outcome | Assessed effect on prevalence of target organism across cattle. | Prevalence of Salmonella. |
| 150 | Gorham, PE; Carroll, LH; McAskill, JW; ... | 1990 | Tilmicosin as a single injection treatment for respiratory disease of feedlot cattle | NA | 17423706 | No/wrong Outcome | Assessed effect on health. |  |
| 151 | Hunter, J. E. B.; Shelley, J. C.; Walton, J. R.; Hart, C. A.; Bennett, M. | 1992 | Apramycin resistance in Escherichia coli: possible transfer to Salmonella typhimurium in calves | 10.1017/s0950268800049748 | 1582469 | No/wrong Outcome | Assessed only antibiotic resistance profiles via antimicrobial susceptibility testing, not relative abundance of specific resistance phenotypes. |  |
| 152 | Levent, Gizem; Schlochtermeier, Ashlynn; Ives, Samuel E.; Norman, Keri N.; Lawhon, Sara D.; Loneragan, Guy H.; Anderson, Robin C.; Vinasco, Javier; Morgan Scott, H. | 2019 | Population Dynamics of Salmonella enterica within Beef Cattle Cohorts Followed from Single-Dose Metaphylactic Antibiotic Treatment until Slaughter | 10.1128/AEM.01386-19 | 31519659 | No/wrong Outcome | Assessed effect on prevalence of target organism across cattle. | Prevalence of Salmonella with some MIC testing but not quantifying relative abundance of specific resistance phenotypes. Was done in included follow-up study Levent et al. 2022. |
| 153 | Lussier, Pamela | 2010 | Characterization of putative extended-spectrum -lactamases (ESBL) producing <i>Escherichia coli</i> isolated from feedlot cattle in southern Alberta | NA | NA | No/wrong Outcome | Descriptive/mechanistic assessment of antibiotic resistance. |  |
| 154 | Murray, S. A.; Holbert, A. C.; Norman, K. N.; Lawhon, S. D.; Sawyer, J. E.; Scott, H. M. | NA | Macrolide-susceptible probiotic Enterococcus faecium ST296 exhibits faecal-environmental-oral microbial community cycling among beef cattle in feedlots | 10.1111/lam.13269 | 31883125 | No/wrong Outcome | Descriptive/mechanistic assessment of antibiotic resistance. | Assessment of effect on absolute abundance is reported in later included publication Murray et al. (2022) |
| 155 | Rodriguez-Palacios, A.; Pickworth, C.; Loerch, S.; LeJeune, J. T. | NA | Transient Fecal Shedding and Limited Animal-to-Animal Transmission of Clostridium difficile by Naturally Infected Finishing Feedlot Cattle | 10.1128/AEM.02736-10 | 21441320 | No/wrong Outcome | Assessed effect on prevalence of target organism across cattle. | Prevalence of C. difficile plus some descriptive/mechanistic genetic work. |
| 156 | Thomas, M.; Webb, M.; Ghimire, S.; Blair, A.; Olson, K.; Fenske, G. J.; Fonder, A. T.; Christopher-Hennings, J.; Brake, D.; Scaria, J. | NA | Metagenomic characterization of the effect of feed additives on the gut microbiome and antibiotic resistome of feedlot cattle | 0.1038/s41598-017-12481-6 | 28947833 | No/wrong Outcome | Assessed effect on composition. | There is some resistome work too but it is presence/absence based not relative/absolute abundance based. |
| 157 | Swaffham Prior, Lord Soulsby of | 1999 | NA | NA | NA | Other | Opinion piece |  |
| 158 | Anonymous | NA | Feedyard dust can include drugs, bacteria | NA | NA | Other | This is a promo piece covering a scientific paper. |  |
| 159 | Berge, A. C. | 2017 | Antibiotic reduction in animal production | NA | NA | Other | Press piece. |  |
| 160 | Bernier Gosselin, Veronique; Fernandez, Javier E.; Ollagnier, Catherine; Morel, Isabelle; Siegenthaler, Raphael; Collaud, Alexandra; Meylan, Mireille; Perreten, Vincent | 2024 | Whole-genome sequencing-based antimicrobial resistance and shedding dynamics of Escherichia coli isolated from calves before and after antimicrobial group treatments | 10.1128/spectrum.03214-23 | 38441467 | Other | There is no replication for investigating the parenteral treatments because drug and timing combination is unique to each of the 8 individuals. |  |
| 161 | Dealy, J.; Moeller, M. W. | 1977 | Influence of bambermycins on Salmonella infection and antibiotic resistance in calves | 10.2527/jas1977.445734x | 324962 | Other | Resistance assessed in an experimentally-infected strain. | We clarified our eligibility criteria to specify that we are interested in effects on established microbiota. |
| 162 | Evangelisti, D. G.; English, A. R.; Girard, A. E.; Lynch, J. E.; Solomons, I. A. | 1975 | Influence of subtherapeutic levels of oxytetracycline on Salmonella typhimurium in swine, calves and chickens | 10.1128/AAC.8.6.664 | 1108781 | Other | Resistance assessed in an experimentally-infected strain. | We clarified our eligibility criteria to specify that we are interested in effects on established microbiota. |
| 163 | Girard, A. E.; English, A. R.; Evangelisti, D. G.; Lynch, J. E.; Solomons, I. A. | 1976 | Influence of subtherapeutic levels of a combination of neomycin and oxytetracycline on Salmonella typhimurium in swine, calves and chickens | 10.1128/AAC.10.1.89 | 791090 | Other | Resistance assessed in an experimentally-infected strain. |  |
| 164 | Hinton, M.; Rixson, P. D.; Allen, V.; Linton, A. H. | NA | The persistence of drug resistant Escherichia coli strains in the majority faecal flora of calves | 10.1017/s0022172400065128 | 6392420 | Other | No replication. | There are only 5 calves with different antimicrobial and other exposures so very descriptive/associative. |
| 165 | Morley, P. S.; Dargatz, D. A.; Hyatt, D. R.; Dewell, G. A.; Patterson, G.; Wittum, T. E. | 2007 | Effects of rearing practices that restrict antimicrobial exposure in feedlot cattle on antimicrobial resistance in enteric bacteria | 10.1089/fpd.2010.0632 | 21034271 | Other | Conference abstract of study published elsewhere. |  |
| 166 | NA | NA | Ceftriaxone-resistant Salmonella infection from antibiotic-treated cattle | 10.1056/NEJM200004273421703 | 10781620 | Other | Case report |  |
| 167 | Pohl, P. | 1983 | Effects and consequences of antibiotic supplementation on resistance of Enterobacteriaceae in animals | 10.5169/seals-590594 | NA | Other | Opinion piece |  |
| 168 | Slyter, L. L.; Kern, D. L.; Weaver, J. M.; Rumsey, T. S. | 1973 | Chlortetracycline and antibiotic resistance of fecal bacteria from heifers | NA |  | Other | Conference abstract of study published elsewhere. |  |
| 169 | Xu, S.; Amarakoon, I. D.; Zaheer, R.; Sura, S.; Reuter, T.; Zvomuya, F.; Cessna, A. J.; Larney, F. J.; McAllister, T. A. | NA | Dissipation of antimicrobials and resistance genes in compost originating from cattle manure after direct oral administration or post-excretion fortification of antimicrobials | NA |  | Other | Conference abstract of study published elsewhere. |  |
| 170 | McEwen, S. A.; Poppe, C.; Popa, M.; Werber, D.; O'Connor, A.; Dunlop, H. | 2000 | Therapeutic and in-feed antimicrobial use and resistance in fecal commensals of swine and cattle | NA | NA | Other | Conference abstract of study published elsewhere. | Published as O'Connor et al. (2002) |

Supplementary Information 11: Reasons for excluding studies from the meta-analysis

22 of the 33 studies included in the narrative synthesis were excluded from the meta-analysis.

7 studies were excluded because the data presented in published figures was insufficient to permit their inclusion in the meta-analysis despite apparently being otherwise eligible (and authors were not able to provide raw data). Of these studies, 3^38–40^ were excluded because they only reported effects on relative abundance outcomes (despite using absolute abundance data to calculate these measures) in the paper and could not provide data to permit the calculation of effects on absolute abundance upon request. Conversely, 1 of these studies was excluded because they only reported effects on absolute abundance and could not provide data to permit thec calculation of effects on relative abundance^41^. The remaining 3 either presented their results without estimates of sampling error^42,43^ or relative to baseline^44^.

Similarly, 11 studies reported across 13 publications^43,45–56^ were excluded because they only assessed the outcome via antimicrobial susceptibility testing which meant that they only assessed the effect on relative abundance (Box 1).

A further 3 studies were excluded because they only assessed the outcome via metagenomics and their raw data and metadata were not sufficiently shared besides^59^.

Finally, 1 study^60^ was excluded because it became clear when processing the data obtained from study authors that the study design contained insufficient repeated sampling to permit its inclusion in the meta-analysis.

Supplementary Information 12: Reasons for excluding effects from the meta-analysis

39 repeatedly-measured effects from 11 studies reported in 12 publications were included in the meta-analysis. The 12 publications are a subset of the 17 publications relating to studies included in the meta-analysis reported in the main study characteristics table (Table 1), because 4 publications were unsuitable for meta-analysis. One study^44^ was unsuitable because the qPCR resistance outcomes were reported relative to baseline and data was not available to recalculate it in terms of absolute and relative abundance. Another^61^ was unsuitable because only marginal means were reported in the publication and data was not available to calculate raw means and SDs. Finally, two studies^52,53^ were unsuitable because they used antimicrobial susceptibility testing to assess the resistance outcome in samples that were elsewhere assessed via direct enumeration^62^ and qPCR^63^ (and hence were included in the meta-analysis).

This left 34 effects for potential inclusion, 5 of which were split into 2 effects due to blocking, resulting in in 39 effects. Two of these were removed before meta-analysis because they were only represented by one study and hence could not be meta-analysed. These were the feed and water samples from two studies^6,27^. This resulted in 39 effects in the meta-analysis. All of the repeated measures within these longitudinally-sampled effects were included in the meta-analysis where possible (occasionally effect sizes were not estimatable), with the exception of 3 of the 257 measurements where the SMDH could not be calculated. These were measurements from 2 studies^19,63^ where the intervention/exposure or control SD was zero due to the identical abundance values across replicates due to imputing values below a detection limit^19^, or where the intervention/exposure or control SD was incalculable because there was only one replicate of data available after processing the raw data.

#### 2.2 Risk of bias in studies

Supplementary Figure 6: Per-domain and overall risk of bias assessments for the 38 publications (reporting 33 studies) included in the review. Randomised trials are indicated by the † symbol and were assessed using the RoB 2 IPRG tool; cluster-randomised trials are indicated by the ‡ symbol and were assessed using the RoB 2 Cluster tool; non-randomised studies are indicated by * symbol. In order to present all study designs in one figure, domain were consolodated between the three risk tools according to their approximate equivalence where applicable. Hence D1 = Bias due to confounding (RoB 2 IPRG/Cluster D1; ROBINS-I D1); D2 = Bias due to selection of participants (RoB 2 Cluster D1b 'Timing of recruitment'; ROBINS-I D2); D3 = Bias due to classification of interventions (ROBINS-I only); D4 = Bias due to deviations from intended interventions (RoB 2 IPRG/Cluster D2; ROBINS-I D4); D5 = Bias due to missing data (RoB 2 IPRG/Cluster D3; ROBINS-I D5); D6 = Bias due to measurement of the outcome (RoB 2 IPRG/Cluster D4; ROBINS-I D6); D7 = Bias due to selection of the reported result (RoB 2 IPRG/Cluster D5; ROBINS-I D7). Similarly, judgements are consolodated into the following unified framework: Low (Yellow X; all tools); Some concerns/Moderate (Orange -; all tools); High/Serious (Dark orange; all tools); Critical (Dark;= red; ROBINS-I/non-randomised studies only).

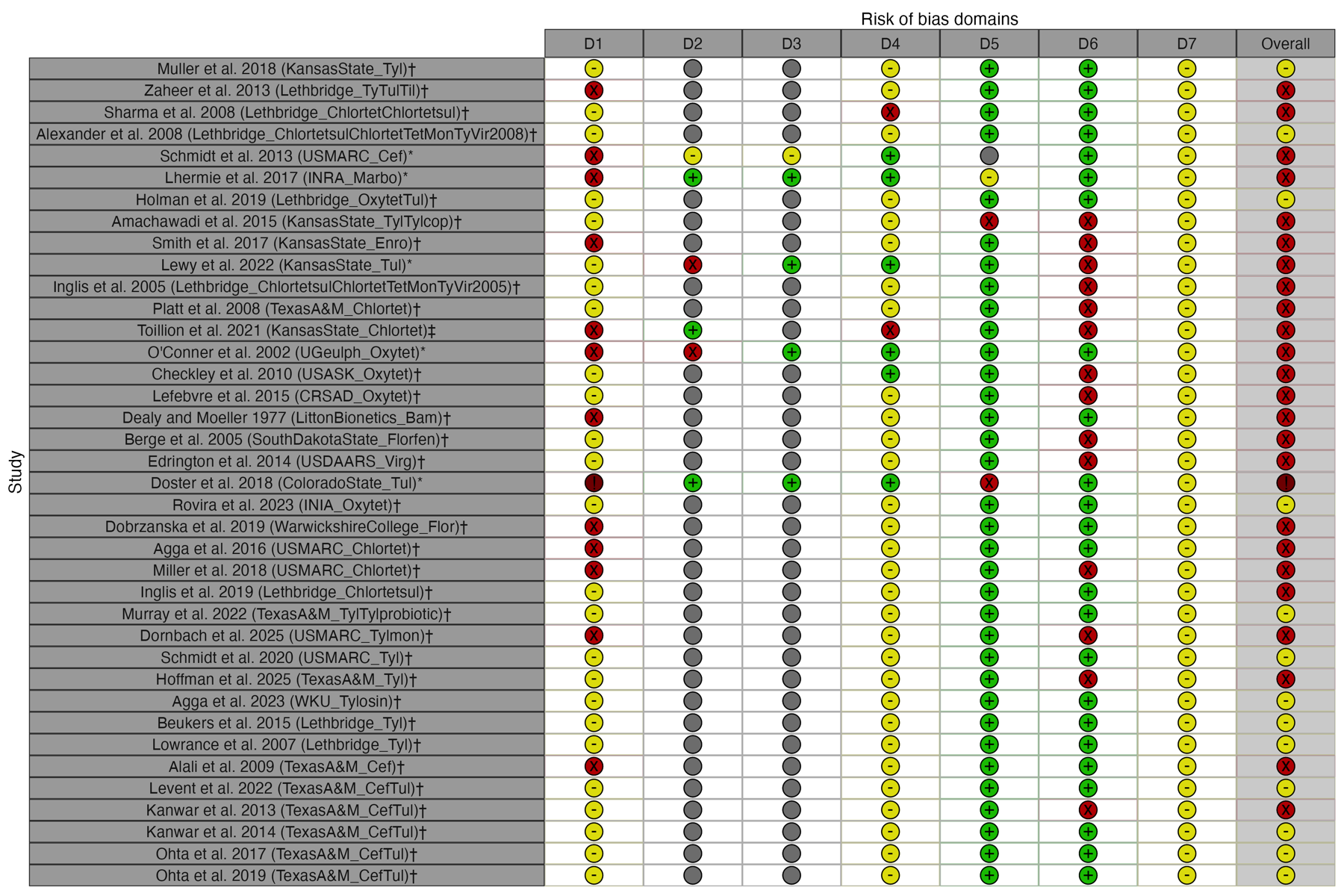

Supplementary Information 13: Full RoB2 assessment

See ‘ROB2_IRPG_beta_v9_GazePetroniWP1_final_120125_v2.xlsm’’.

Supplementary Information 14: Full RoB2 assessment

See ‘ROB2_cluster_beta_v3_GazePetroniWP1_final_120125_v2(Toillion2022only).xlsm’.

Supplementary Information 15: Full ROBINSI assessment

See ‘ROBINS-I_v1_19sep2016_allassessments.docx’.

#### 2.3 Results of individual studies

Supplementary Figure 7: Forest plot of effects on absolute abundance (density) of antibiotic resistant bacteria in faeces pre-antibiotic administration, aggregated at the study level. Effects were aggregated using an intercept-only random effects model accounting for non-independence between effects from the same study, between temporally-autocorrelated effects from the same study arm, and residual variance as random effects.

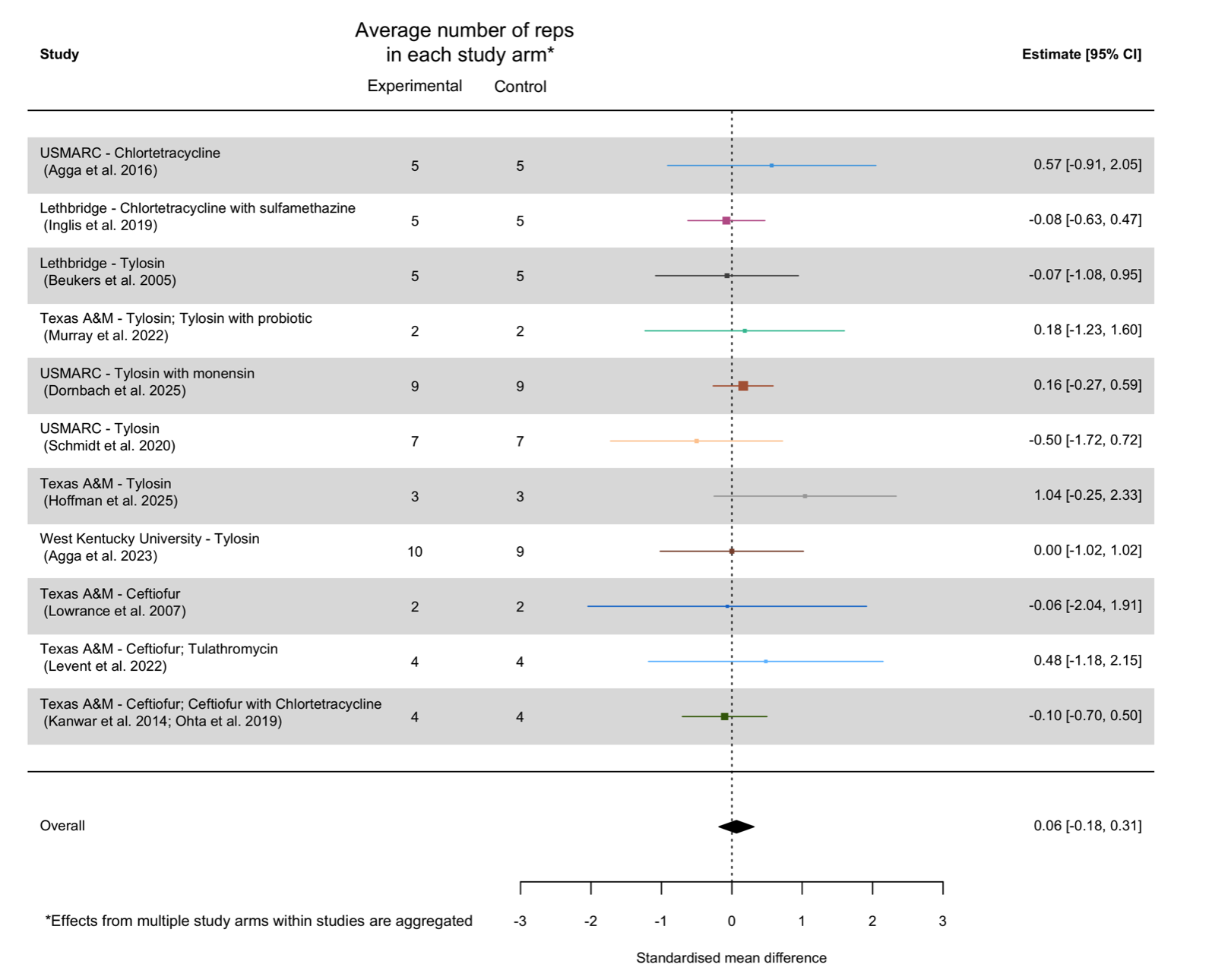

Supplementary Figure 8: Forest plot of effects on logit-transformed relative abundance (logit-transformed proportion) of antibiotic resistant bacteria in faeces post-antibiotic administration, aggregated at the study level. Effects were aggregated using an intercept-only random effects model accounting for non-independence between effects from the same study, between temporally-autocorrelated effects from the same study arm, and residual variance as random effects.

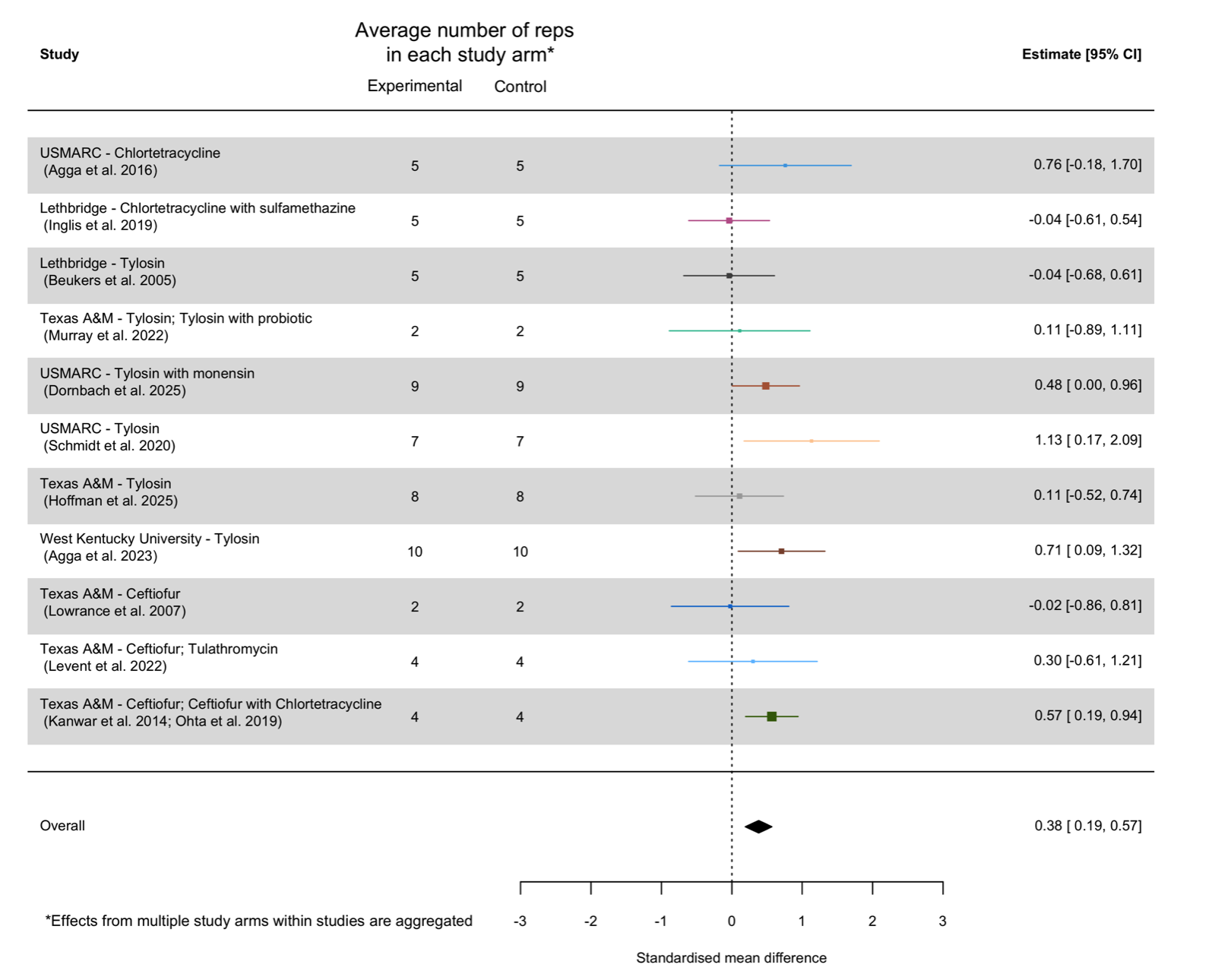

Supplementary Figure 9: Forest plot of effects on logit-transformed relative abundance (logit-transformed proportion) of antibiotic resistant bacteria in faeces pre-antibiotic administration, aggregated at the study level. Effects were aggregated using an intercept-only random effects model accounting for non-independence between effects from the same study, between temporally-autocorrelated effects from the same study arm, and residual variance as random effects.

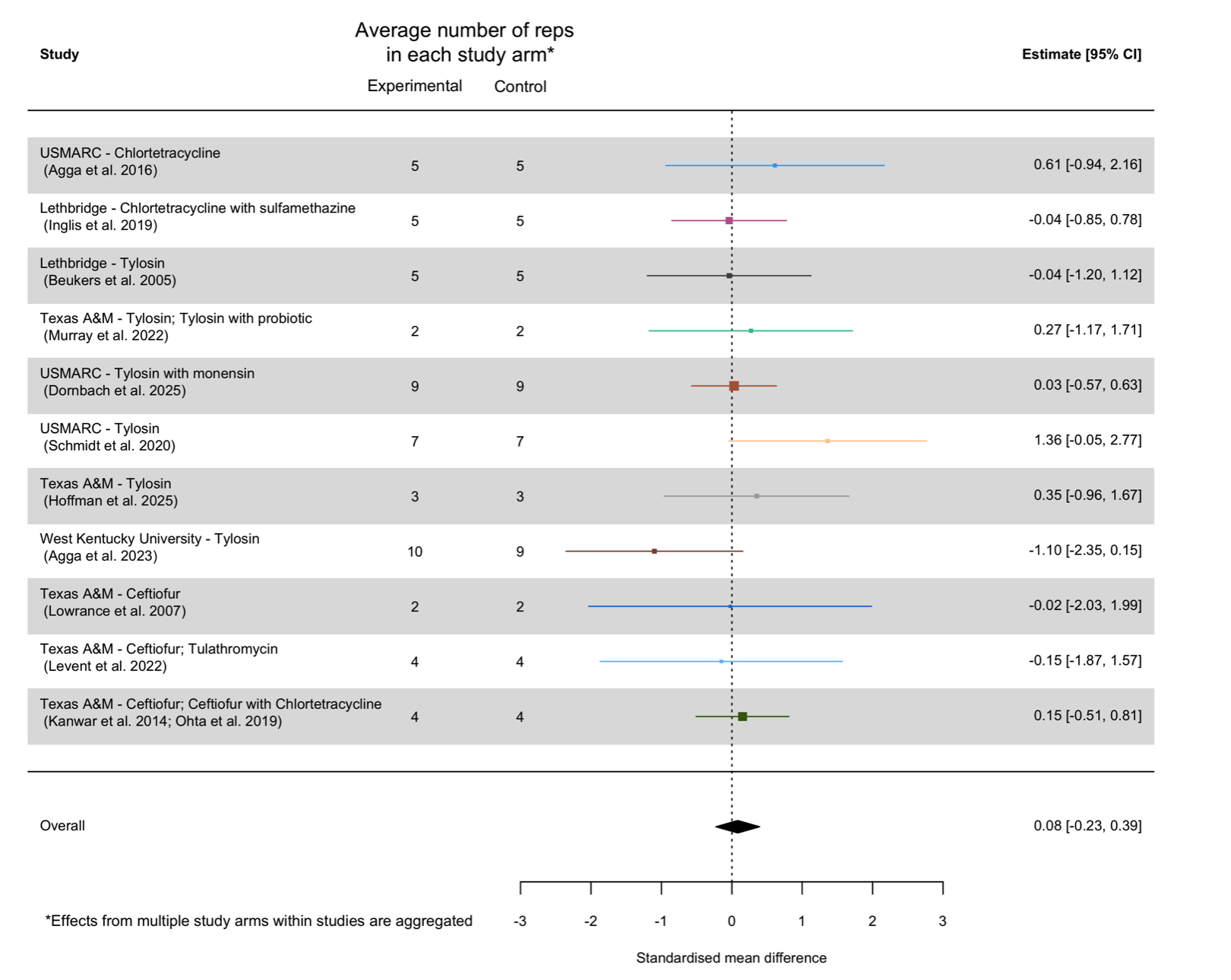

Supplementary Figure 10: Forest plot of effects on arcsine-transformed relative abundance (arcsine-transformed proportion) of bacteria in faeces post-antibiotic administration, aggregated at the study level. Effects were aggregated using an intercept-only random effects model accounting for non-independence between effects from the same study, between temporally-autocorrelated effects from the same study arm, and residual variance as random effects.

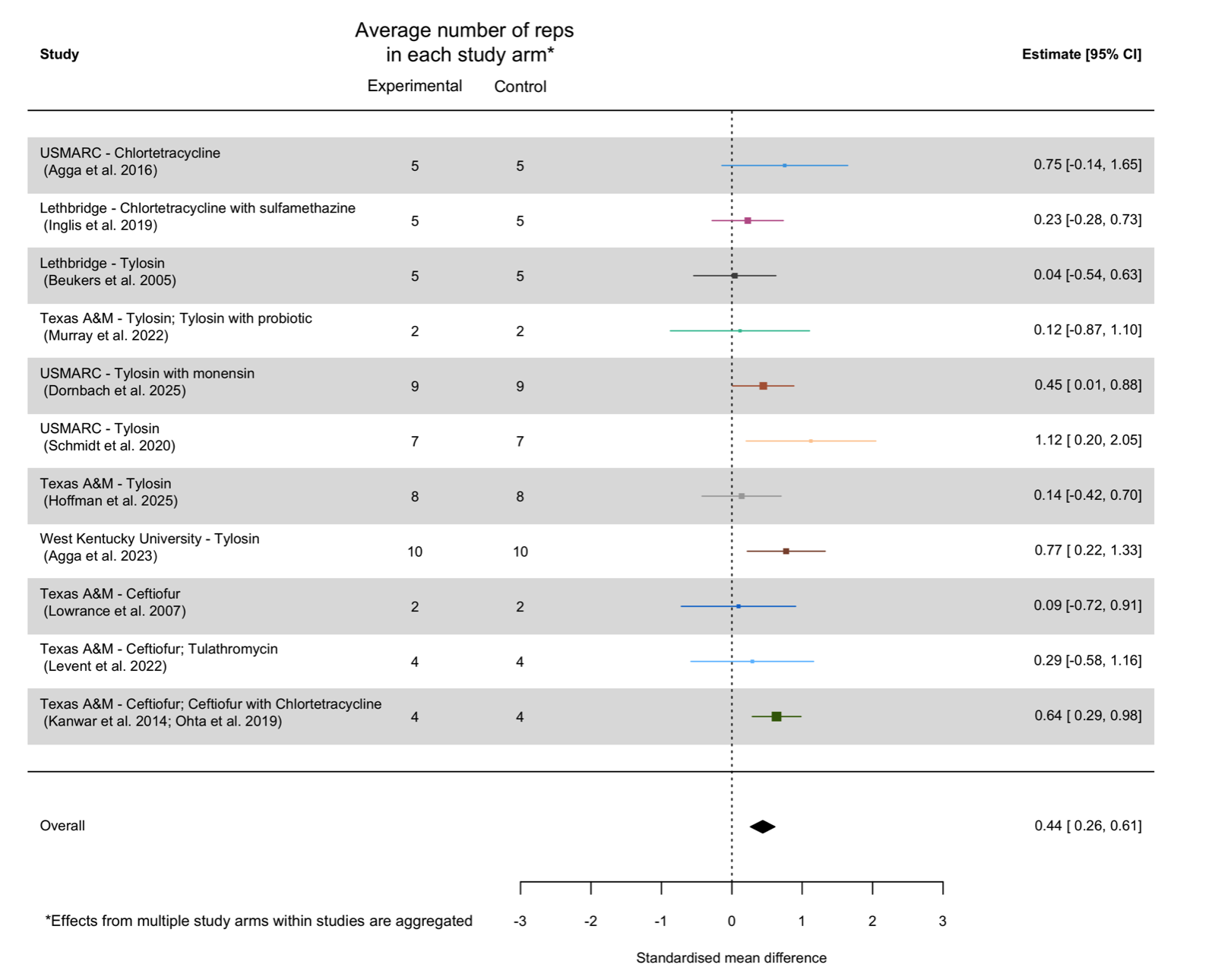

Supplementary Figure 11: Forest plot of effects on arcsine-transformed relative abundance (arcsine-transformed proportion) of antibiotic resistant bacteria in faeces pre-antibiotic administration, aggregated at the study level. Effects were aggregated using an intercept-only random effects model accounting for non-independence between effects from the same study, between temporally-autocorrelated effects from the same study arm, and residual variance as random effects.

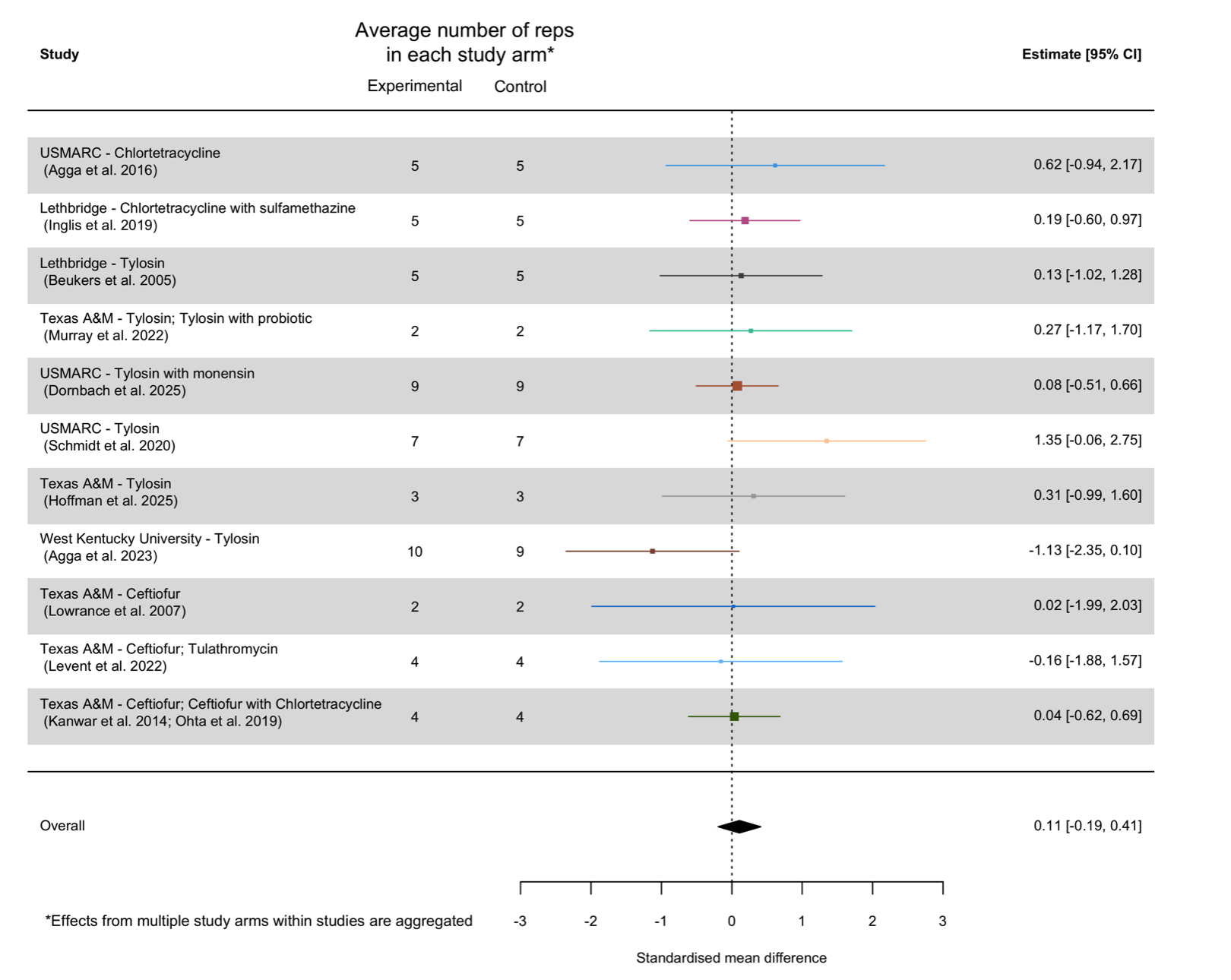

Supplementary Figure 12: Forest plot of effects on absolute abundance of antibiotic resistant bacteria in pen surface soil post-antibiotic administration. Effects were aggregated at the study level using an intercept-only random effects model accounting for non-independence between effects from the same study, between temporally-autocorrelated effects from the same study arm, and residual variance as random effects. These were then pooled into an overall effect using an equal-effects model sensu Viechtbauer (2024b).

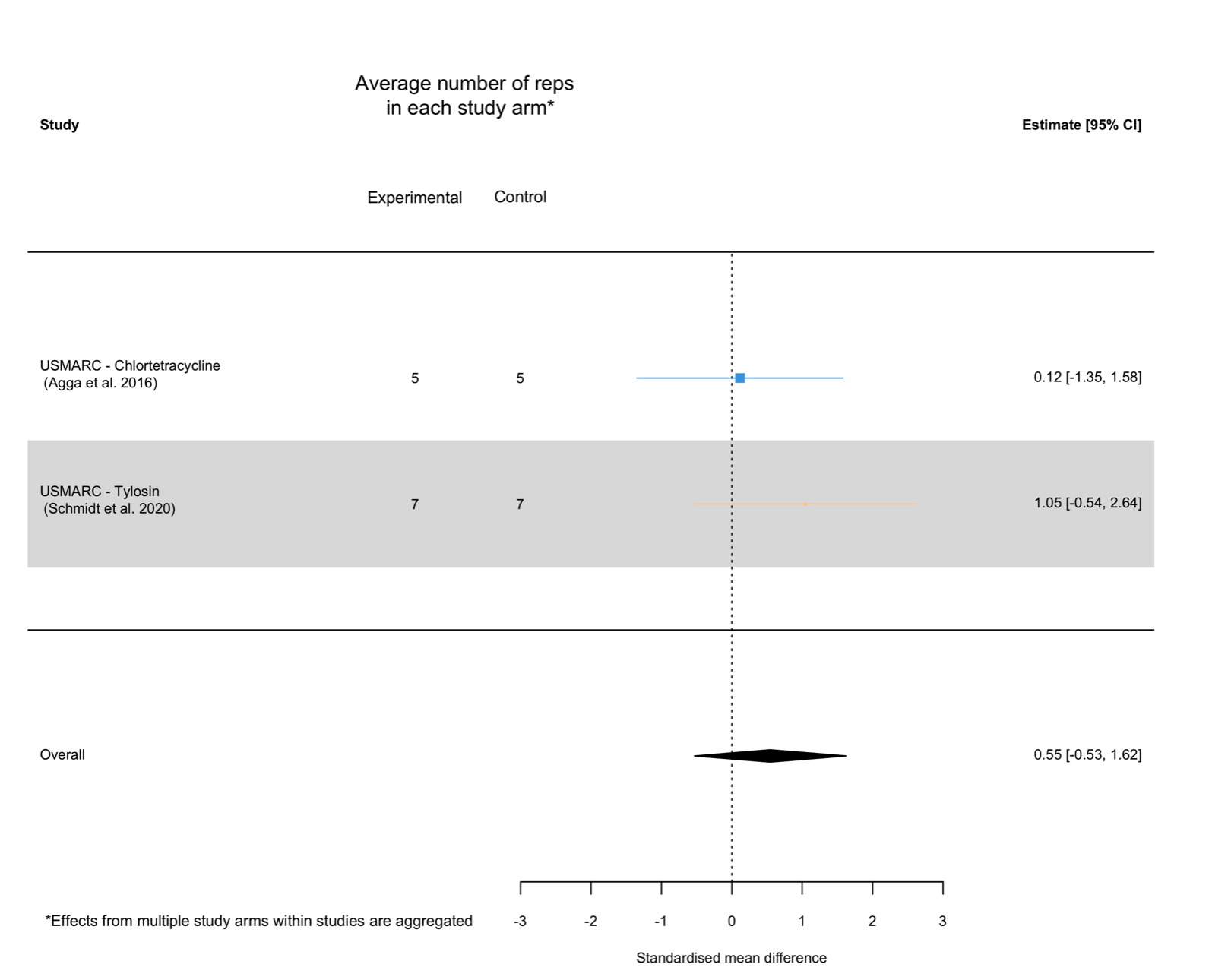

Supplementary Figure 13: Forest plot of effects on absolute abundance of antibiotic resistant bacteria in pen surface soil pre-antibiotic administration, aggregated at the study level. Effects were aggregated using an intercept-only random effects model accounting for meta-analytically non-independence between effects from the same study, between temporally-autocorrelated effects from the same study arm, and residual variance as random effects.)

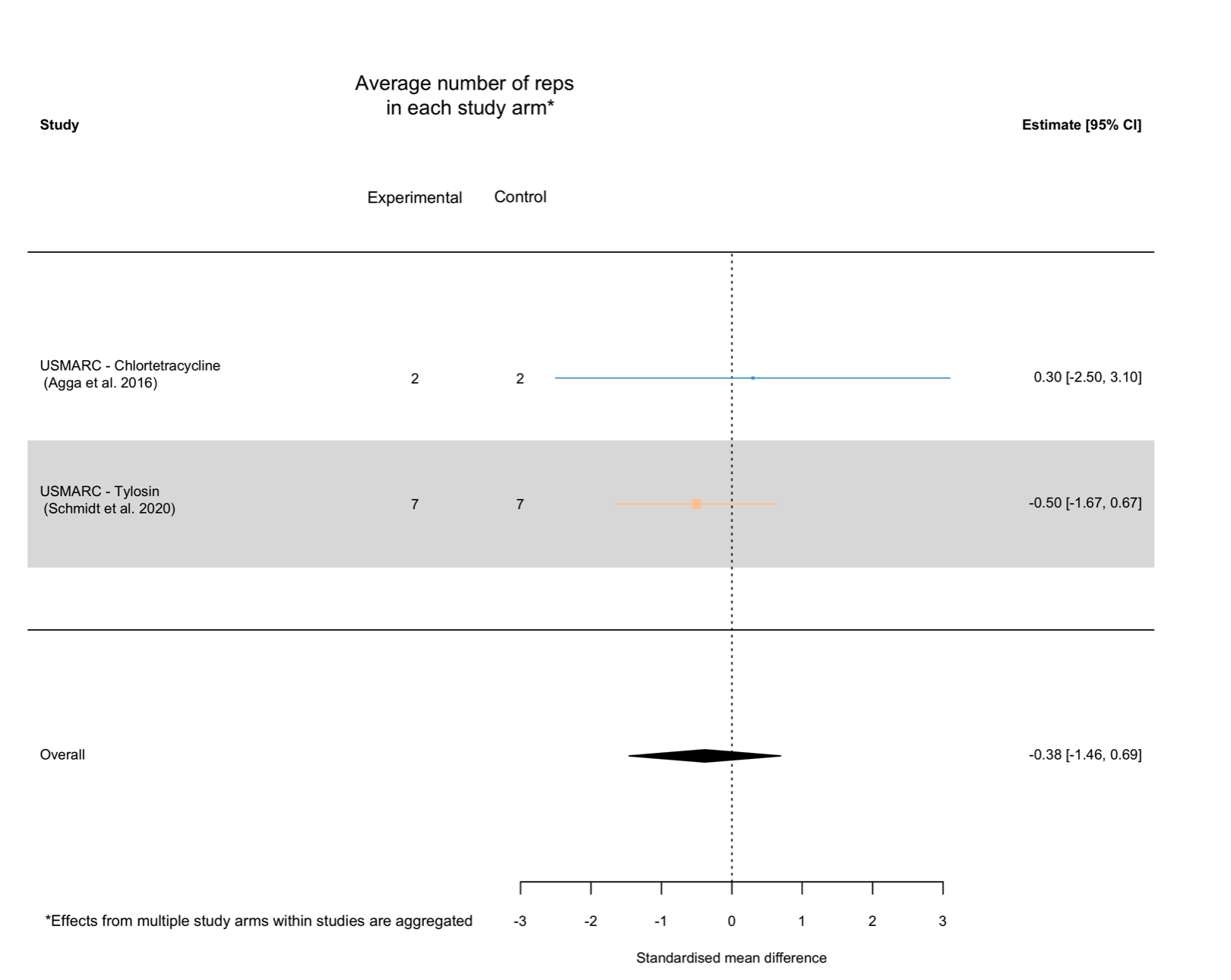

Supplementary Figure 14: Forest plot of effects on relative abundance (logit-transformed proportion) of antibiotic resistant bacteria in pen surface soil post-antibiotic administration, aggregated at the study level. Effects were aggregated using an intercept-only random effects model accounting for non-independence between effects from the same study, between temporally-autocorrelated effects from the same study arm, and residual variance as random effects.

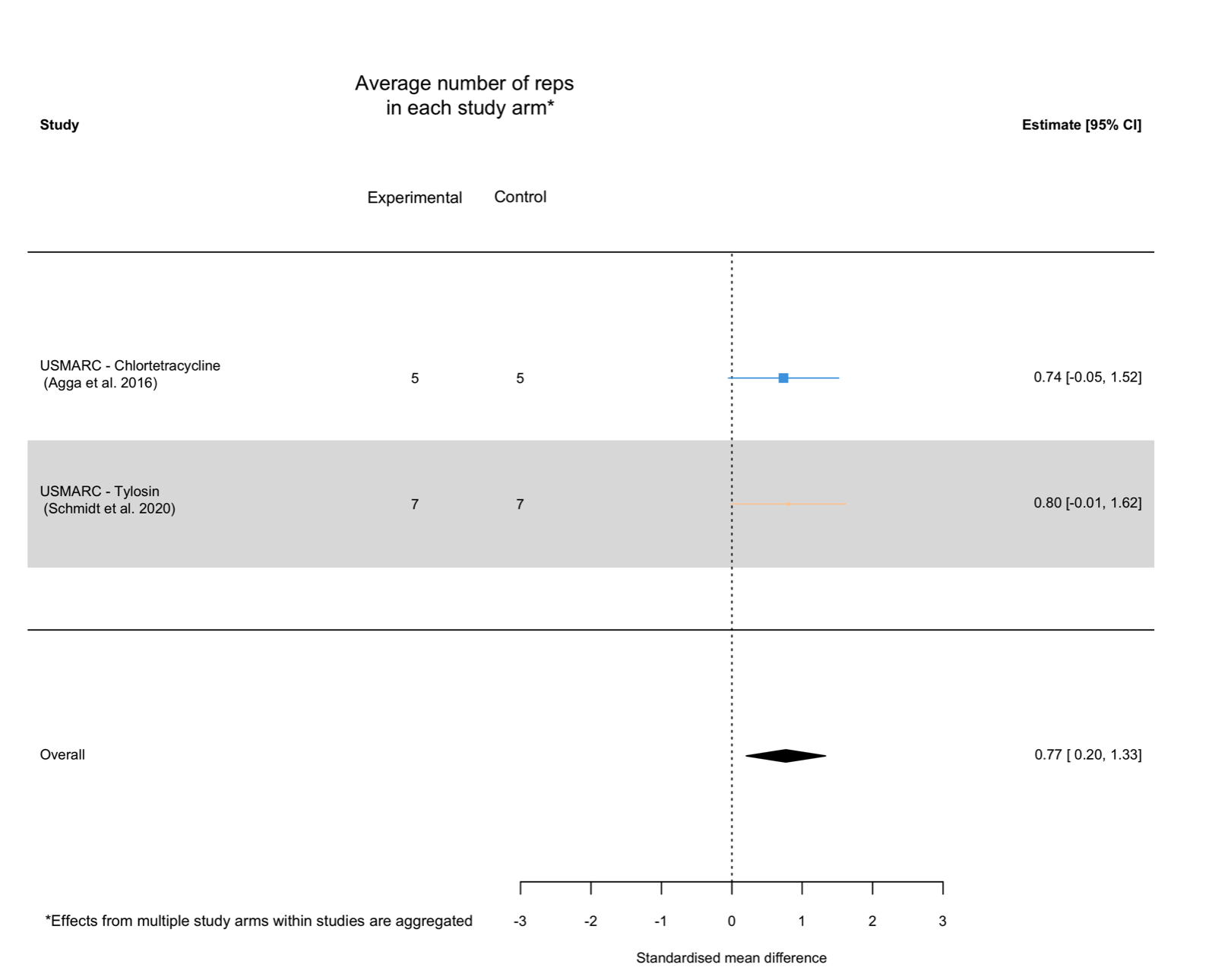

Supplementary Figure 15: Forest plot of effects on relative abundance (logit-transformed proportion) of antibiotic resistant bacteria in pen surface soil pre-antibiotic administration, aggregated at the study level. Effects were aggregated using an intercept-only random effects model accounting for non-independence between effects from the same study, between temporally-autocorrelated effects from the same study arm, and residual variance as random effects.

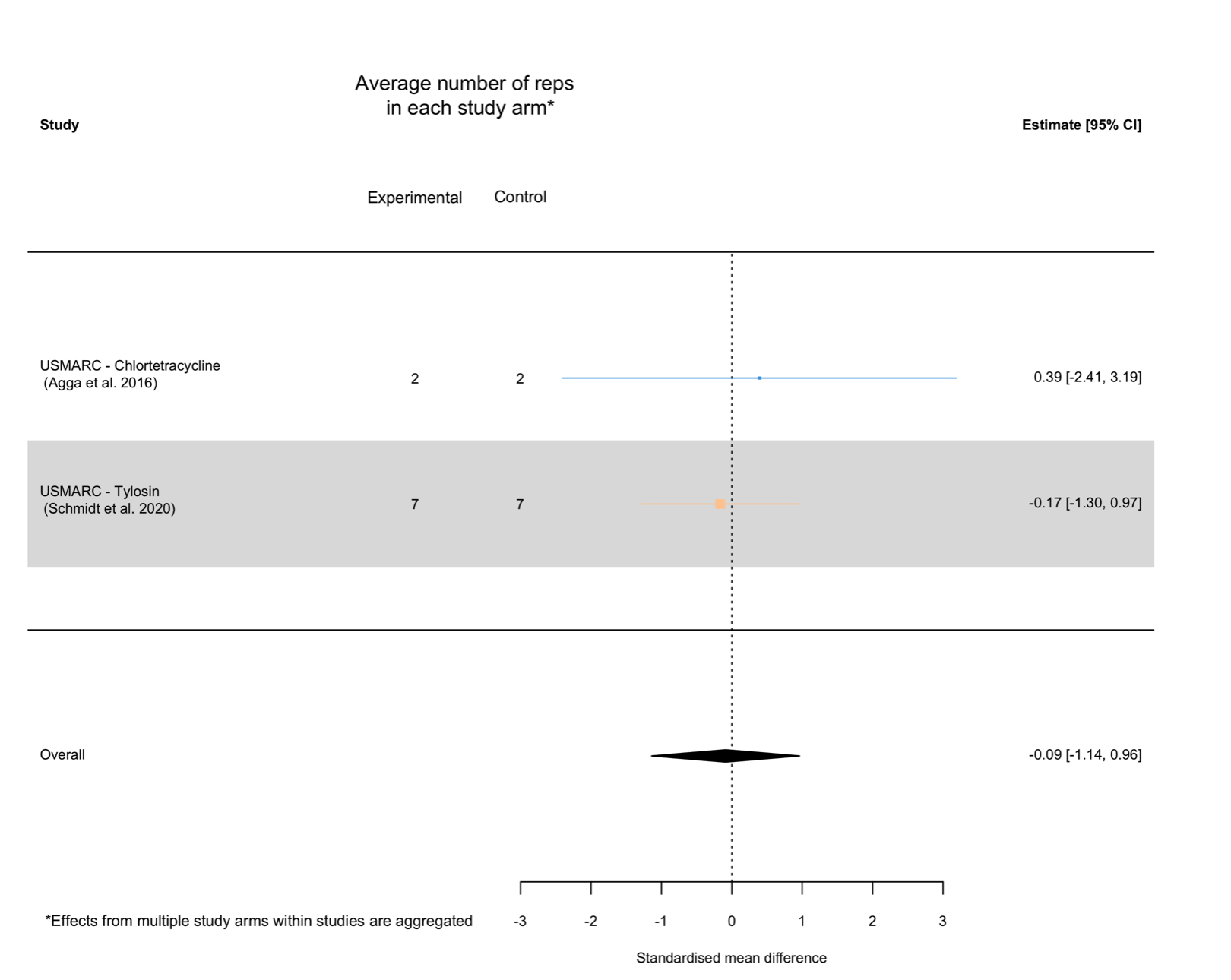

Supplementary Figure 16: Forest plot of effects on relative abundance (arcine-transformed proportion) of antibiotic resistant bacteria in pen surface soil post-antibiotic administration, aggregated at the study level. Effects were aggregated using an intercept-only random effects model accounting for non-independence between effects from the same study, between temporally-autocorrelated effects from the same study arm, and residual variance as random effects.

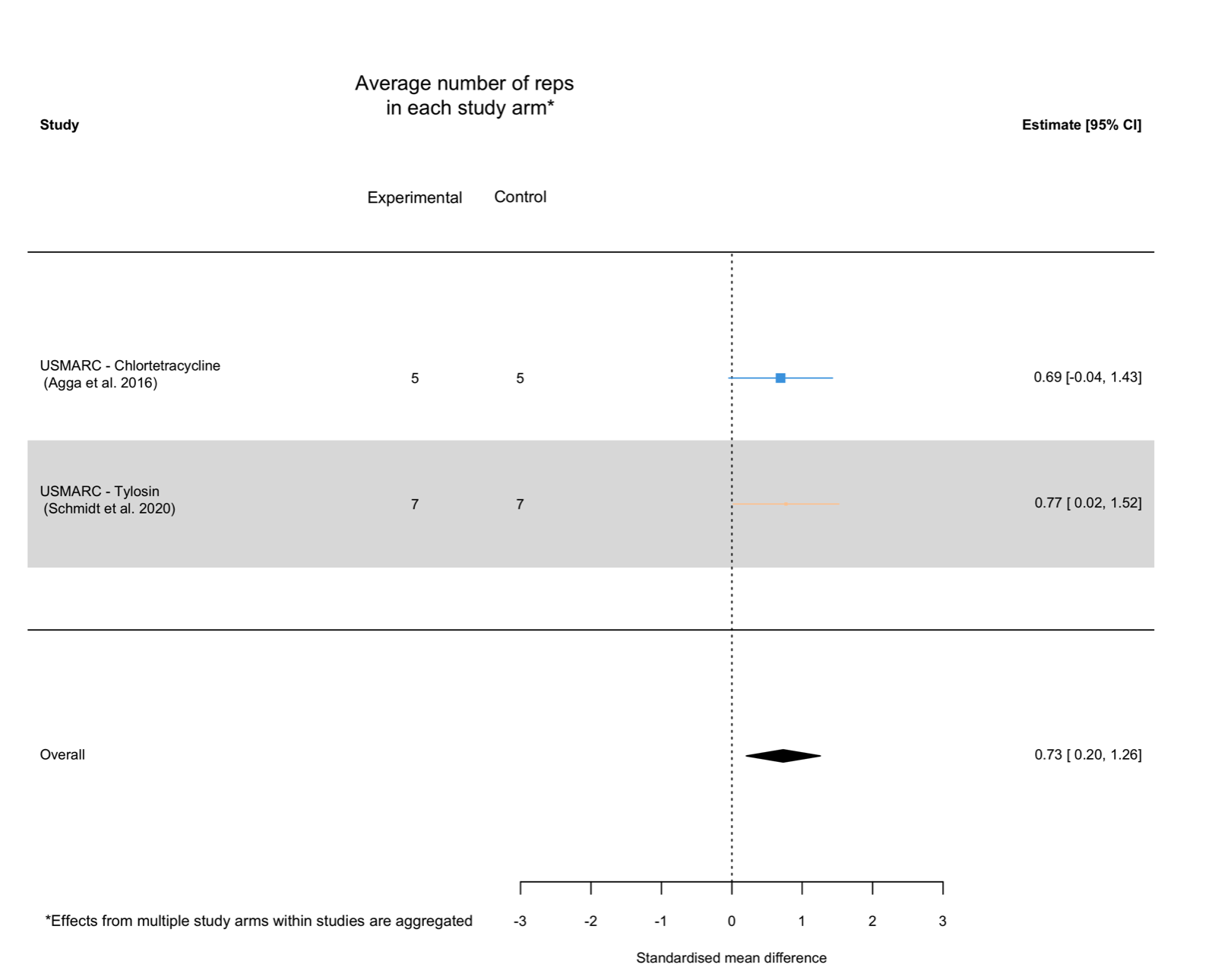

Supplementary Figure 17: Forest plot of effects on relative abundance (arcsine-transformed proportion) of antibiotic resistant bacteria in pen surface soil pre-antibiotic administration, aggregated at the study level. Effects were aggregated using an intercept-only random effects model accounting for non-independence between effects from the same study, between temporally-autocorrelated effects from the same study arm, and residual variance as random effects.

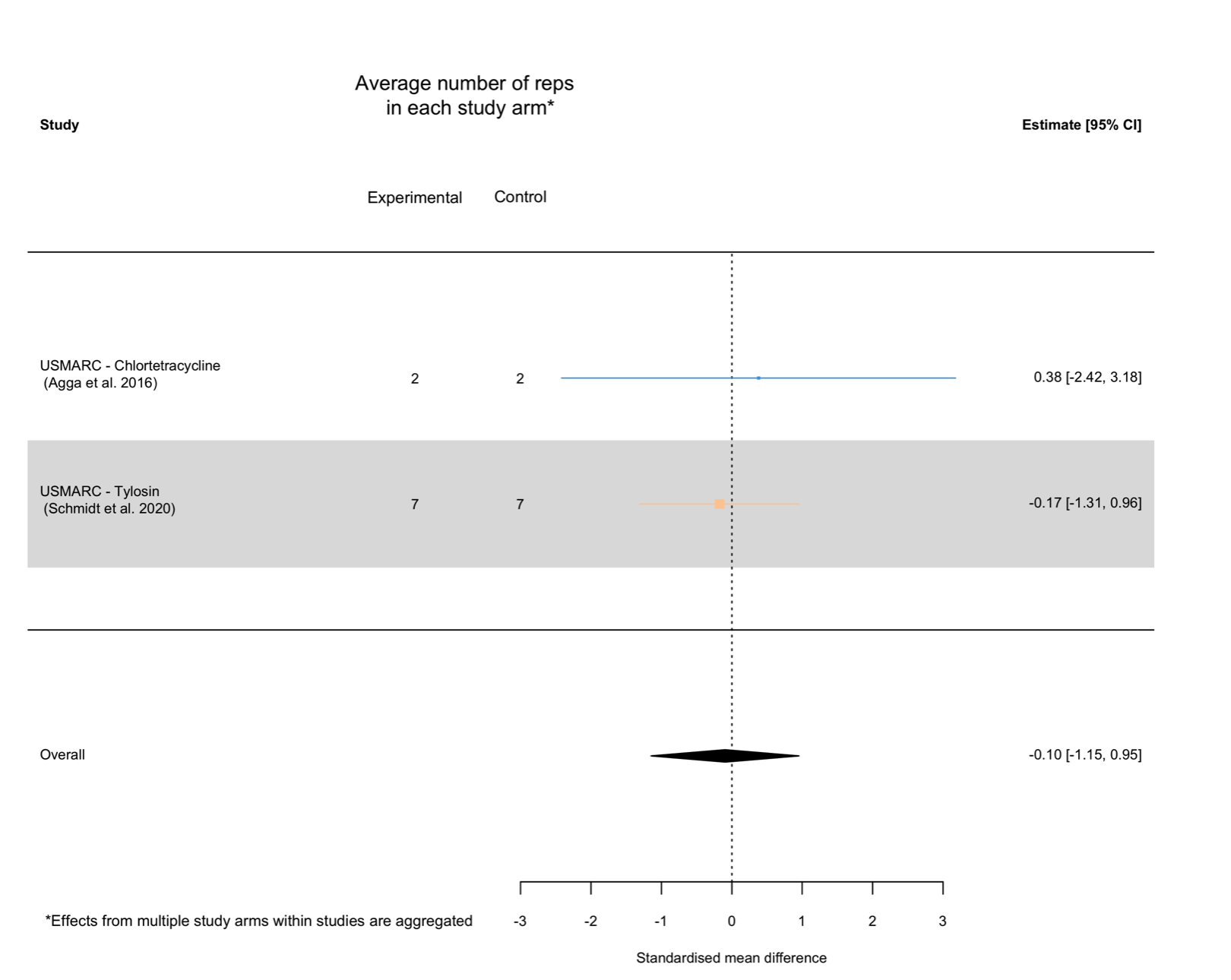

#### 2.4 Results of syntheses (hypotheses 1 & 2)

##### 2.4.1 Effects measured during antibiotic administration

###### 2.4.1.1 Primary analysis

Supplementary Information 16: Description of random effect estimates in the intercept-only models of the overall effect (hypothesis 1)

Random effect estimates of model of the overall effect during the administration of antibiotics indicated that the strongest random effect was of temporal autocorrelation between repeated measures/effect estimates from the same study arm ($\tau^{2}$ = 0.2; $\rho$ = 0.09). There was no detectable effect of study ($\sigma_{1}^{2}$ = 0), and minimal residual variance ($\sigma_{1}^{2}$ = 0). Profile likelihood plots indicated that these random effect estimates were accurate (Supplementary Figure 18).

Random effect estimates for the meta-regression model with a linear effect of timing of sampling were similar to those of intercept-only models. There was moderate temporal autocorrelation between repeated measures of antibiotic resistance taken within the same study arm ($\tau^{2}$ = 0.15; $\rho$ = 0.13), no detectable effect of study ($\sigma_{1}^{2}$ = 0), and minimal residual variance ($\sigma_{1}^{2}$ = 0). Profile likelihood plots indicated that these random effect estimates were accurate (Supplementary Figure 19).

Supplementary Figure 18: Profile likelihood plots for each of the random effect estimates from the intercept-only model of the overall effect during the administration antibiotics. Where the random effect is accurately estimated, the random effect estimate should occur at the peak log-likelihood value (intersection of the two dotted lines). Profile likelihood plots shown are for the A) σ²₁ parameter, representing the random effect of study; B) σ²₂ parameter, representing the random effect of residual variance due to multiple effect sizes estimates per sampling event; C) τ² parameter, representing the random effect of study arm (outer term); D) ρ parameter, representing the random effect temporal autocorrelation within study arms (inner term)

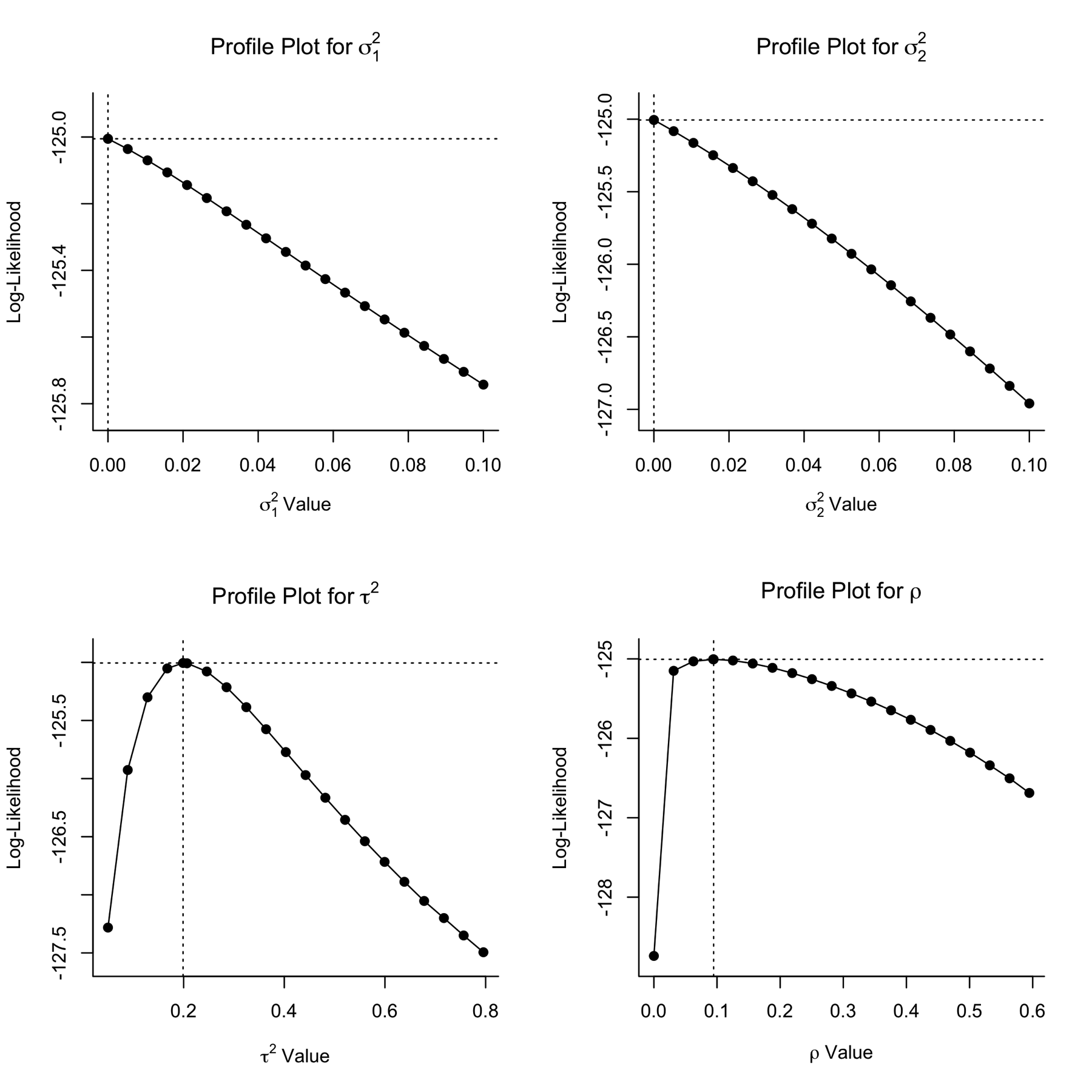

Supplementary Figure 19: Profile likelihood plots for each of the random effect estimates from the intercept-only model of the overall effect during the administration antibiotics. Where the random effect is accurately estimated, the random effect estimate should occur at the peak log-likelihood value (intersection of the two dotted lines). Profile likelihood plots shown are for the A) σ²₁ parameter, representing the random effect of study; B) σ²₂ parameter, representing the random effect of residual variance due to multiple effect sizes estimates per sampling event; C) τ² parameter, representing the random effect of study arm (outer term); D) ρ parameter, representing the random effect temporal autocorrelation within study arms (inner term)

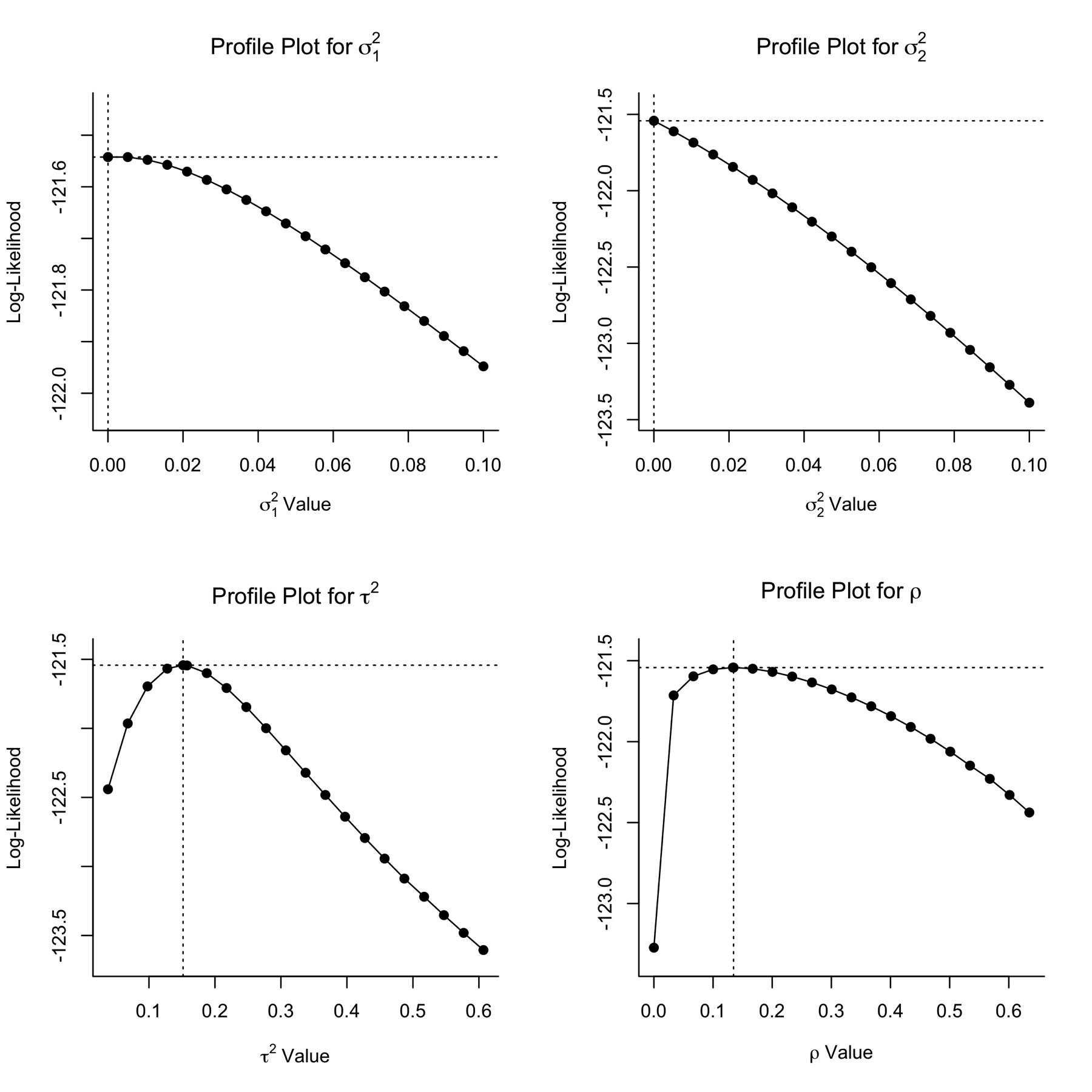

Supplementary Figure 20: Plots representing results of meta-analytical models testing our two hypotheses for effects measured during the administration of antibiotics. In these plots, colours of the points represent the studies from which they are derived, and their size is proportional to their inverse standard error (1/SE; small points are less precise estimates, larger points are more precise estimates). A) Orchard plot representing the modelled overall effect of antibiotic administration on the absolute abundance of resistant bacteria/resistance genes relating to the intervention antibiotic class in the targeted bacterial population. This overall effect alongside its heterogeneity estimate, are derived from the intercept-only model. In this plot, the overall effect (model intercept after accounting for random effects) is represented by the 'trunk' of the tree (black circle), with its confidence interval represented its 'branch' (shorter, thicker line through it), and its prediction interval represented by its 'twig' (longer, thinner line through it).B) Bubble plot representing the modelled relationship between linear timing of sampling (days of continuous antibiotic administration) and effect size (SMDH) from the meta-regression model. In this plot, the modelled effect of time is represented by the bold line, with the dashed line and dotted lines representing the 95% confidence and prediction intervals, respectively.

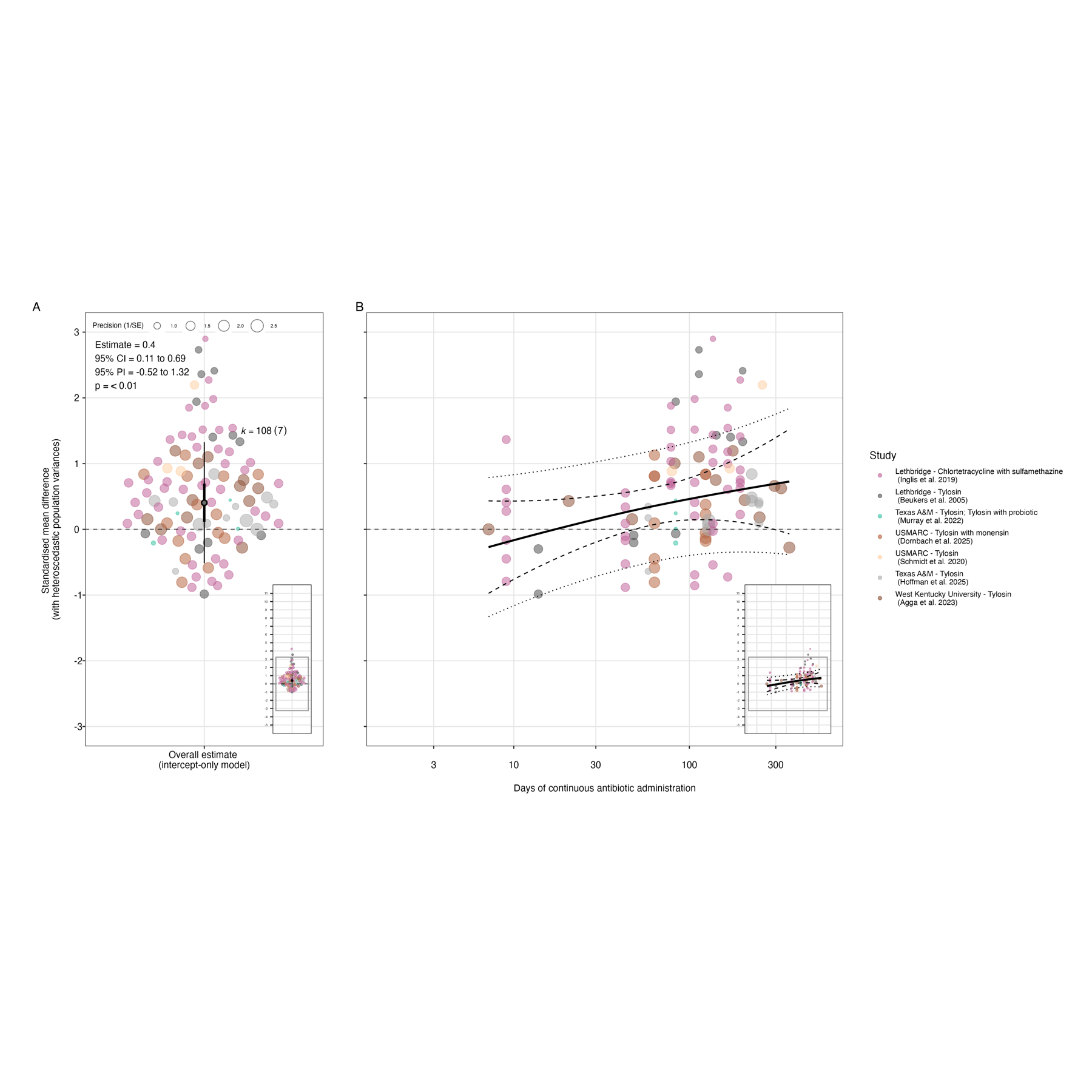

Supplementary Figure 21: Plots representing results of meta-analytical models testing our two hypotheses for effects on the absolute abundance of all determinants measured during the administration of antibiotics. In these plots, colours of the points represent the studies from which they are derived, and their size is proportional to their inverse standard error (1/SE; small points are less precise estimates, larger points are more precise estimates). A) Orchard plot representing the modelled overall effect of antibiotic administration on the absolute abundance of resistant bacteria/resistance genes relating to the intervention antibiotic class in the targeted bacterial population. This overall effect alongside its heterogeneity estimate, are derived from the intercept-only model. In this plot, the overall effect (model intercept after accounting for random effects) is represented by the 'trunk' of the tree (black circle), with its confidence interval represented its 'branch' (shorter, thicker line through it), and its prediction interval represented by its 'twig' (longer, thinner line through it). B) Bubble plot representing the modelled moderating effect of linear timing of sampling (days of continuous antibiotic administration) and residual heterogeneity from the meta-regression model. In this plot, the modelled effect of time is represented by the bold line, with the dashed line and dotted lines representing the 95% confidence and prediction intervals, respectively.

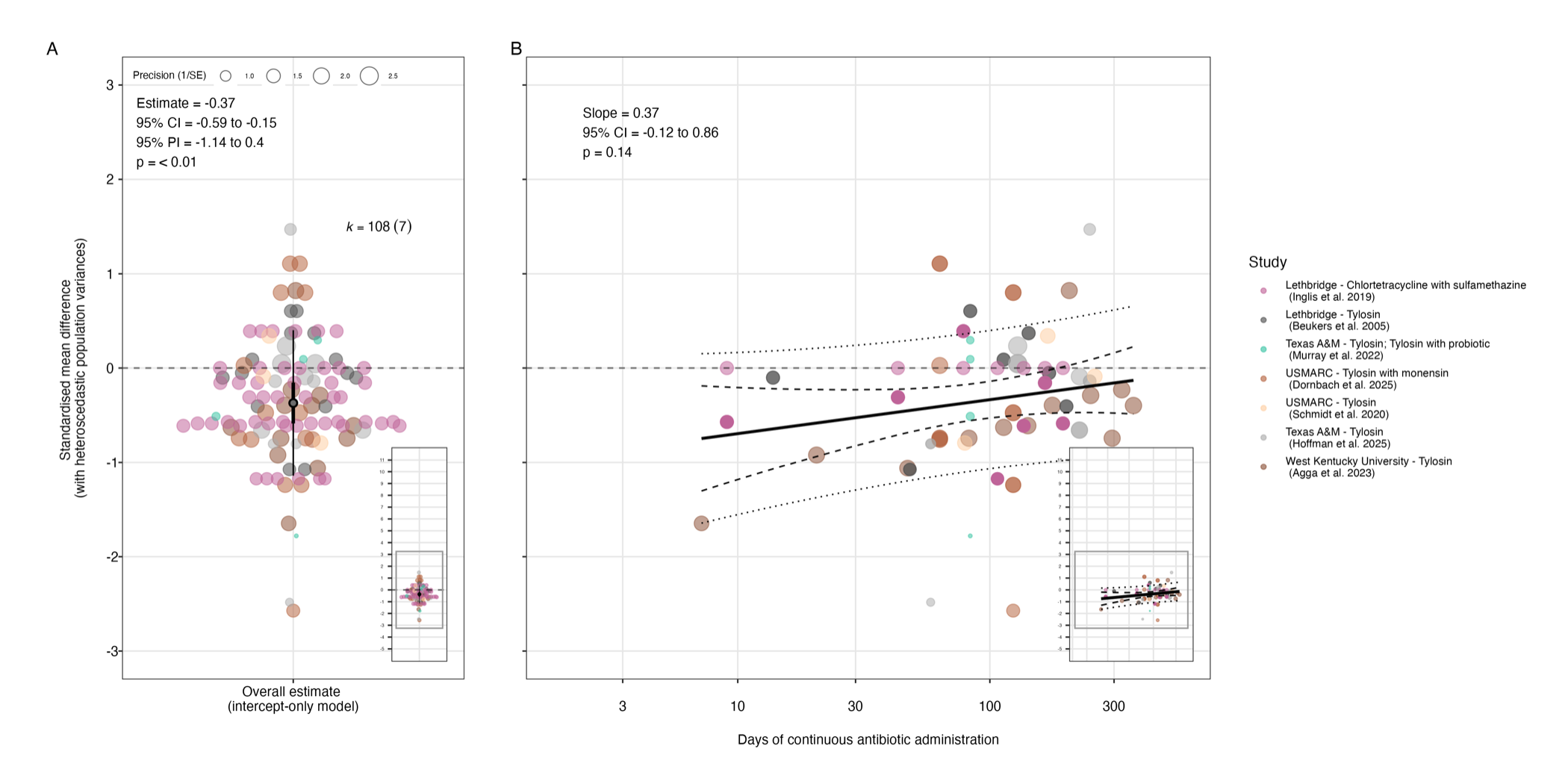

Supplementary Figure 22: Plots representing results of meta-analytical models testing our two hypotheses for effects on the absolute abundance of all determinants measured during the administration of antibiotics. In these plots, colours of the points represent the studies from which they are derived, and their size is proportional to their inverse standard error (1/SE; small points are less precise estimates, larger points are more precise estimates). A) Orchard plot representing the modelled overall effect of antibiotic administration on the absolute abundance of resistant bacteria/resistance genes relating to the intervention antibiotic class in the targeted bacterial population. This overall effect alongside its heterogeneity estimate, are derived from the intercept-only model. In this plot, the overall effect (model intercept after accounting for random effects) is represented by the 'trunk' of the tree (black circle), with its confidence interval represented its 'branch' (shorter, thicker line through it), and its prediction interval represented by its 'twig' (longer, thinner line through it). B) Bubble plot representing the modelled moderating effect of non-linear timing of sampling (days of continuous antibiotic administration) and residual heterogeneity from the meta-regression model. In this plot, the modelled effect of time is represented by the bold line, with the dashed line and dotted lines representing the 95% confidence and prediction intervals, respectively.

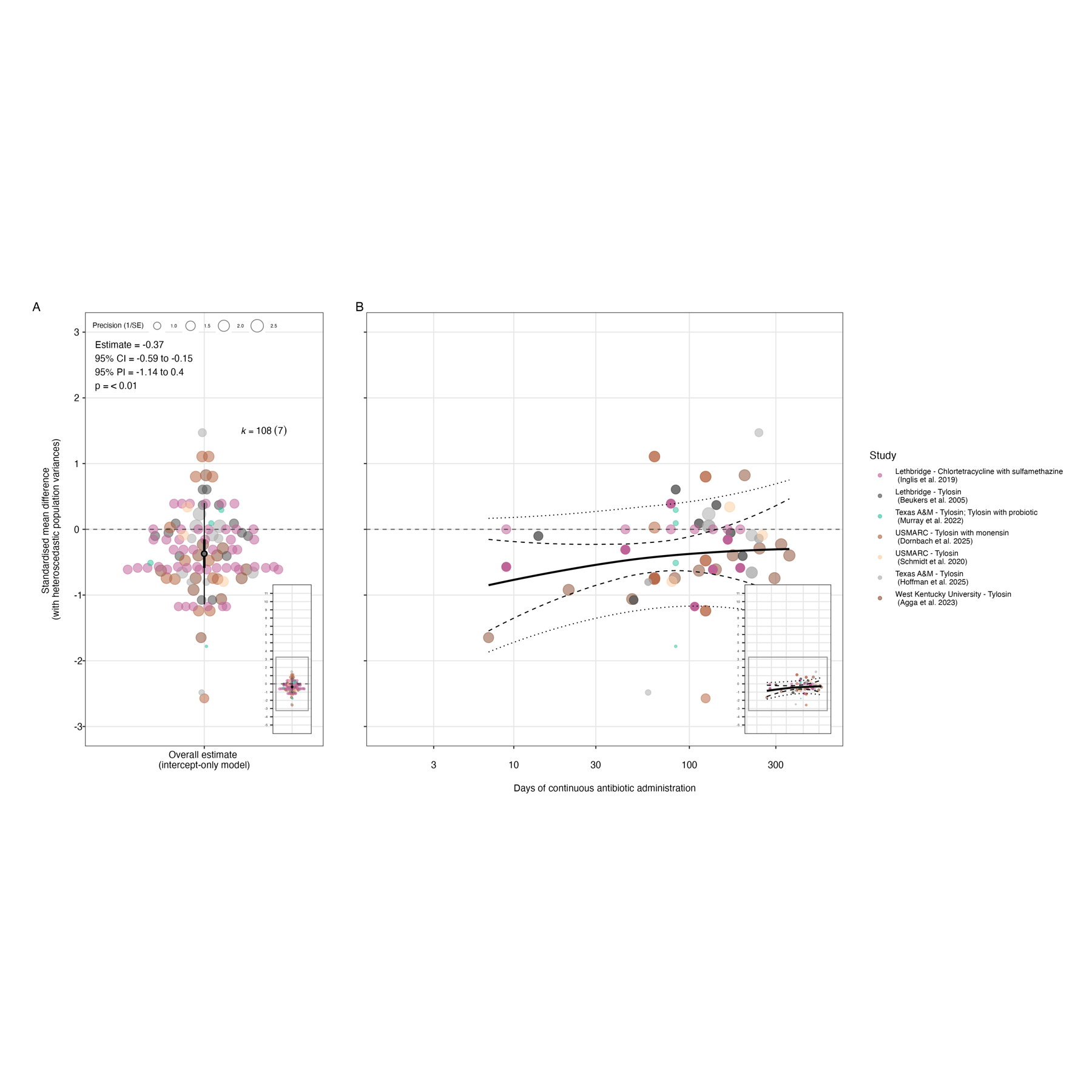

###### 2.4.1.2 Sensitivity testing

Supplementary Figure 23: Orchard plots representing the sensitivity of the modelled overall effect during the administration of antibiotics to leaving one study out. In these plots, the mean overall effect is represented by the 'trunk' of the tree (black circle), with its confidence interval represented its 'branch' (shorter, thicker line through it), and its prediction interval represented by its 'twig' (longer, thinner line through it). The mean and precision (1/standard error) of the individual effect sizes from studies included in the model is represented by the position and size of the coloured points, respectively.

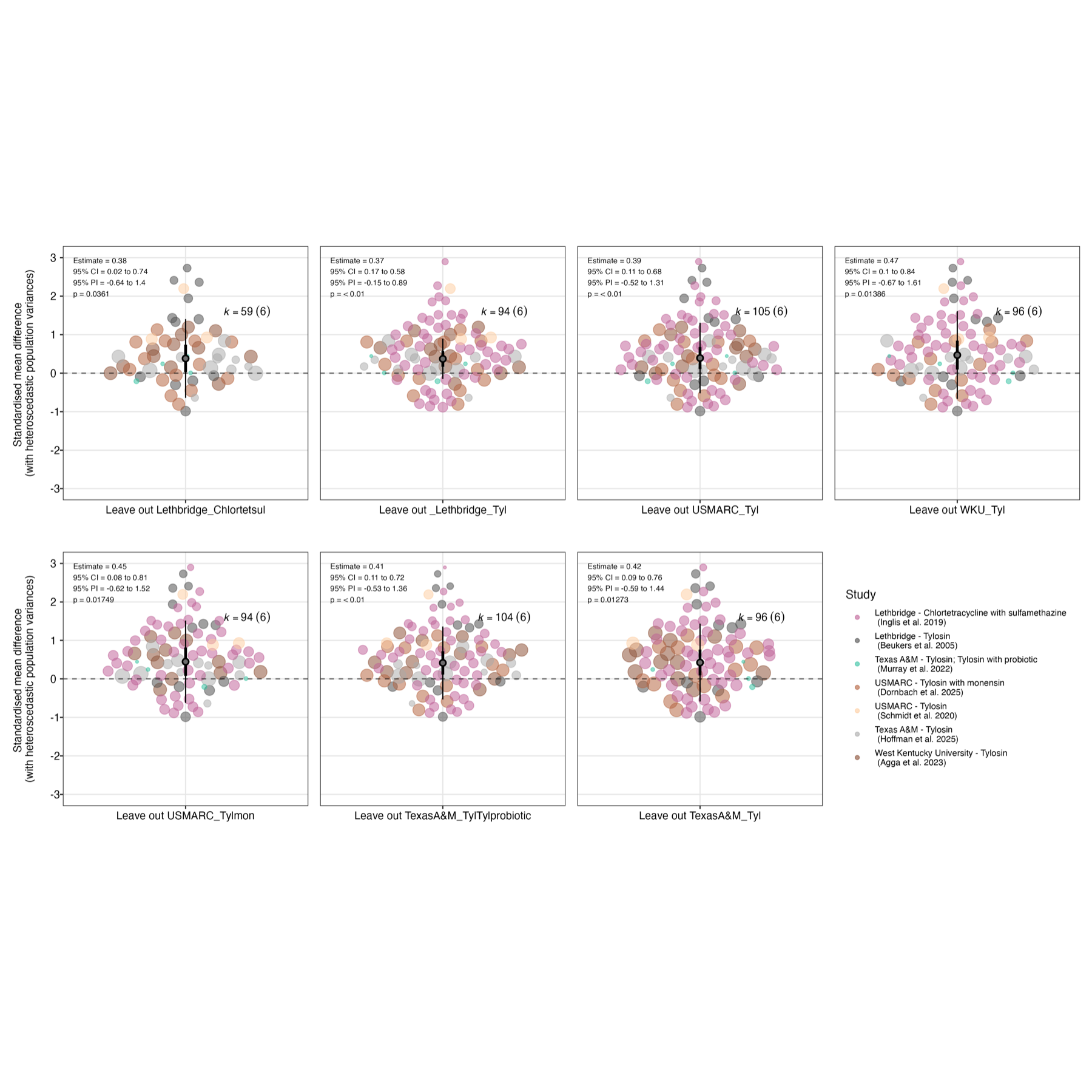

Supplementary Figure 24: Orchard plots representing the sensitivity of the modelled overall effect during the administration of antibiotics to correcting for shared control non-independence and the removal of potentially problematic studies. In these plots, the mean overall effect is represented by the 'trunk' of the tree (black circle), with its confidence interval represented its 'branch' (shorter, thicker line through it), and its prediction interval represented by its 'twig' (longer, thinner line through it). The mean and precision (1/standard error) of the individual effect sizes from studies included in the model is represented by the position and size of the coloured points, respectively.

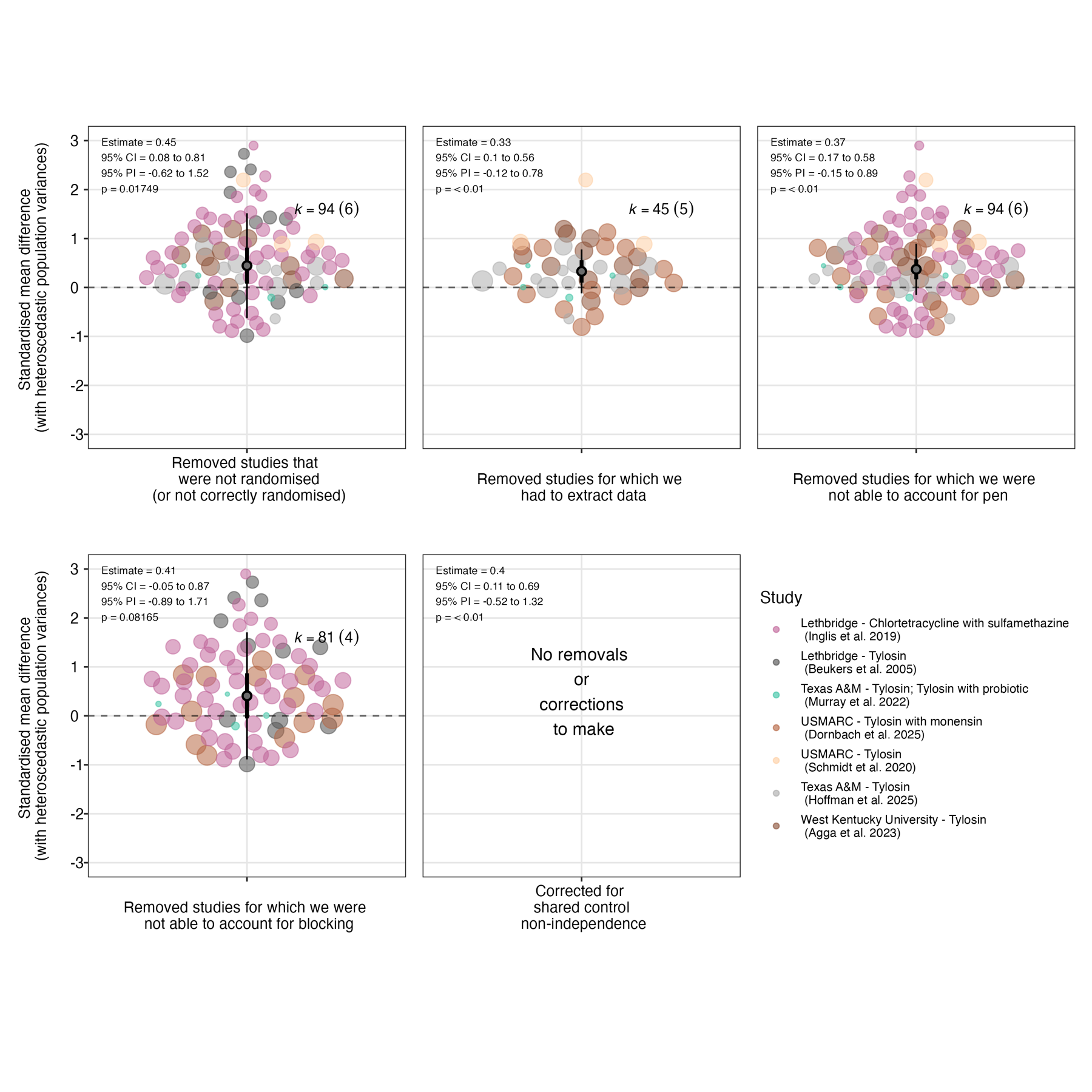

Supplementary Figure 25: Bubble plots representing the sensitivity of the modelled moderating effect of timing of sampling during the administration of antibiotics to leaving one study out. In these plots, the modelled effect of time is represented by the bold line, with the dashed line and dotted lines representing the 95% confidence and prediction intervals, respectively. The mean and precision (1/standard error) of the individual effect sizes from studies included in the model is represented by the position and size of the coloured points, respectively.

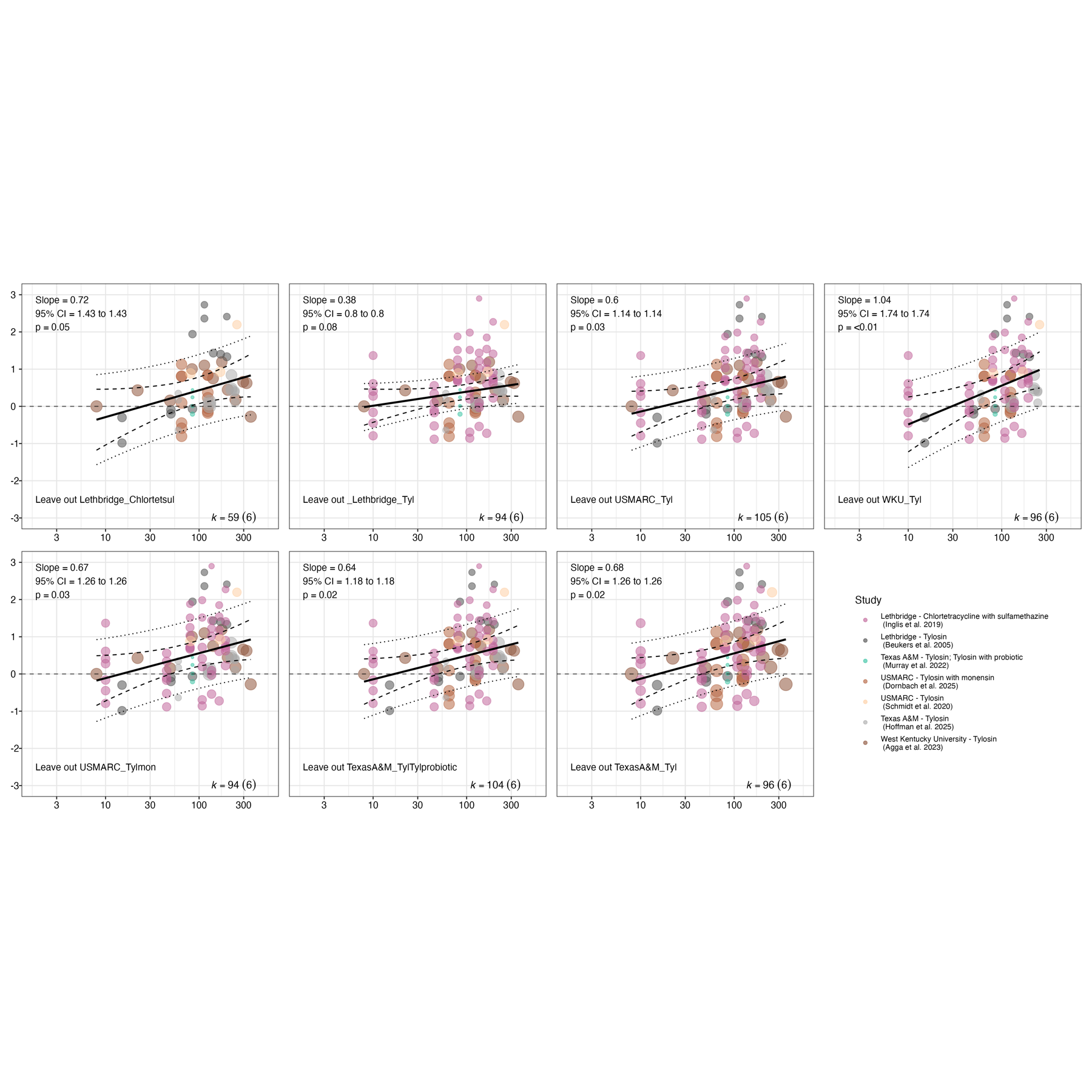

Supplementary Figure 26: Bubble plots representing the sensitivity of the modelled moderating effect of timing of sampling during the administration of antibiotics to correcting for shared control non-independence and the removal of potentially problematic studies. In these plots, the modelled effect of time is represented by the bold line, with the dashed line and dotted lines representing the 95% confidence and prediction intervals, respectively. The mean and precision (1/standard error) of the individual effect sizes from studies included in the model is represented by the position and size of the coloured points, respectively.

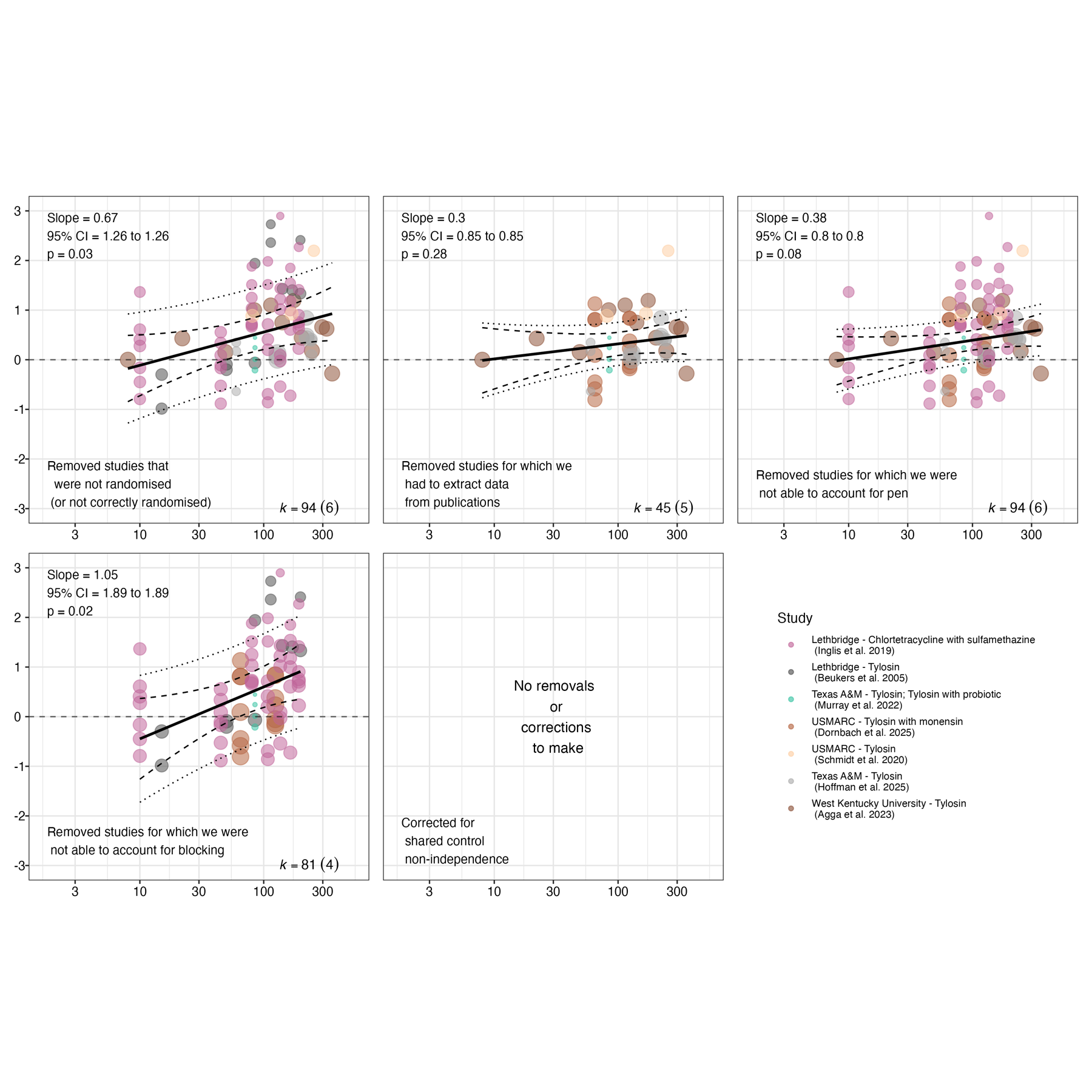

Supplementary Figure 27: Orchard plots representing the sensitivity of the modelled overall effect during the administration of antibiotics to assumptions about the correlation between effect sizes from the same study. Orchard plots of the original model and assuming correlations of 0, 0.2, 0.4, 0.6 and 0.8 are shown. In these plots, the mean overall effect is represented by the 'trunk' of the tree (black circle), with its confidence interval represented its 'branch' (shorter, thicker line through it), and its prediction interval represented by its 'twig' (longer, thinner line through it).

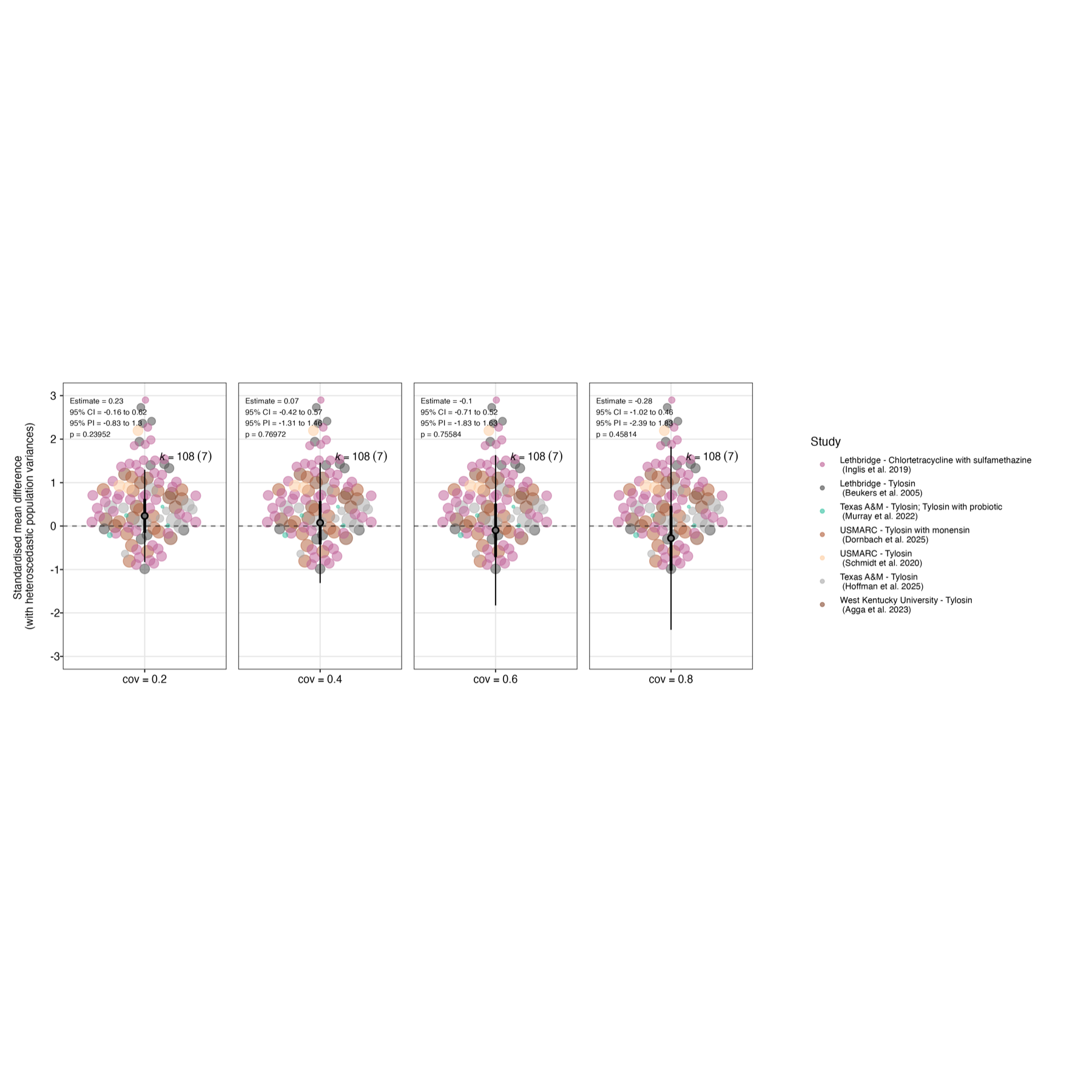

Supplementary Figure 28: Bubble plots representing the sensitivity of the moderating effect of timing of sampling on the effect during the administration of antibiotics to assumptions about the correlation between effect sizes from the same study. Bubble plots of the original model and assuming correlations of 0, 0.2, 0.4, 0.6 and 0.8 are shown. In these plots, the modelled effect of time is represented by the bold line, with the dashed line and dotted lines representing the 95% confidence and prediction intervals, respectively. The mean and precision (1/standard error) of the individual effect sizes from studies included in the model is represented by the position and size of the coloured points, respectively.

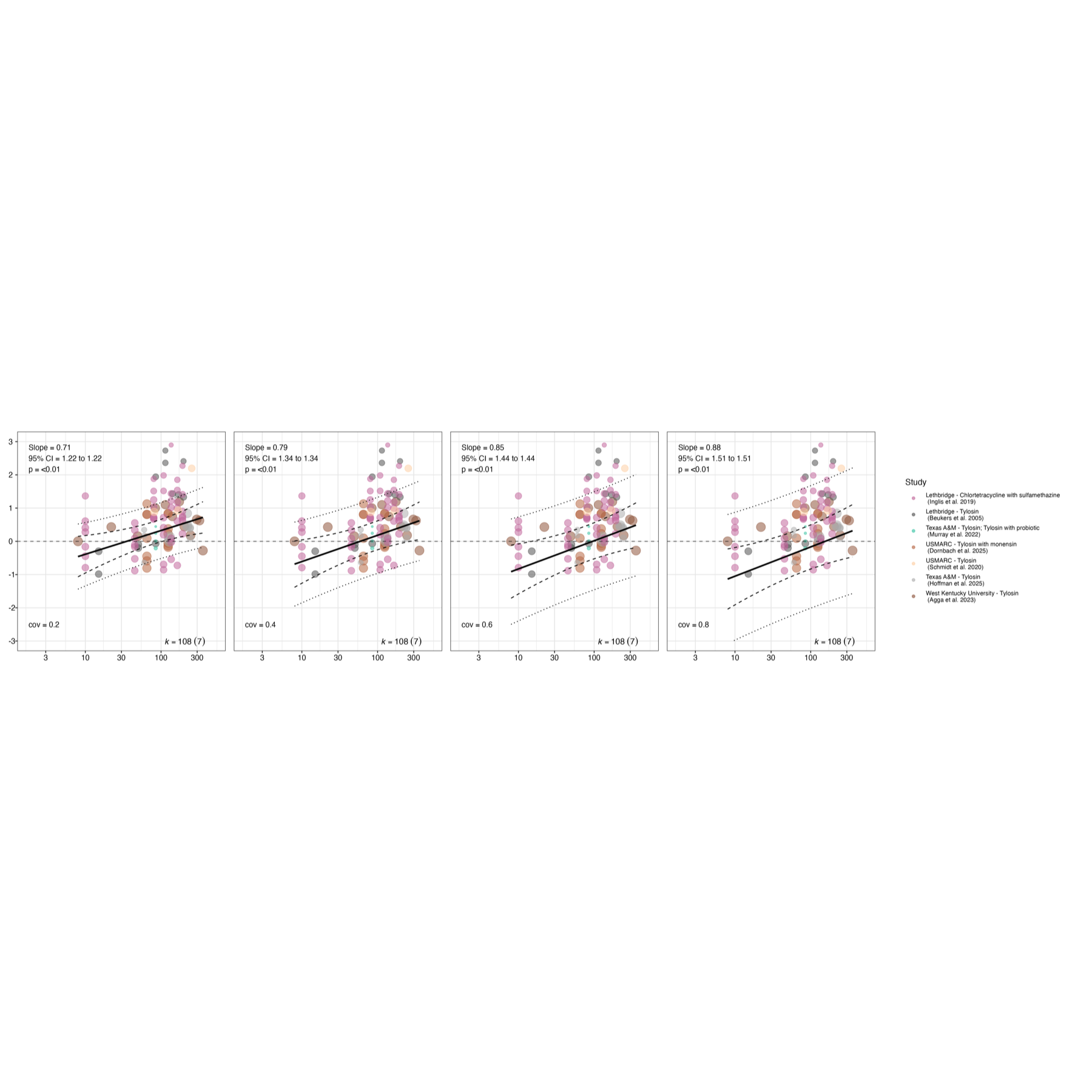

##### 2.4.2 Effects measured after antibiotic administration

###### 2.4.2.1 Primary analysis

Supplementary Information 17: Description of random effect estimates in the meta-regression models of moderating effect of time (hypothesis 2)

Random effect estimates of the model of the overall effect after the administration of antibiotics indicated that the strongest random effect was of temporal autocorrelation between repeated measures/effect estimates from the same study arm, though even this was very small ($\tau^{2}$ = 0; $\rho$ = 0.23). There was no detectable effect of study ($\sigma_{1}^{2}$ = 0) or residual variance ($\sigma_{1}^{2}$ = 0). Profile likelihood plots indicated that these random effect estimates were accurate (Supplementary Figure 29).

Random effect estimates for the meta-regression model with a linear effect of time were similar to those of intercept-only models. There was a very small amount of temporal autocorrelation between repeated measures of antibiotic resistance taken within the same study arm ($\tau^{2}$ = 0; $\rho$ = 0.23), and no detectable effect of study ($\sigma_{1}^{2}$ = 0) or residual variance ($\sigma_{1}^{2}$ = 0). Profile likelihood plots indicated that these random effect estimates were accurate (Supplementary Figure 30).

Supplementary Figure 29: Profile likelihood plots for each of the random effect estimates from the intercept-only model of the overall effect after the administration antibiotics. Where the random effect is accurately estimated, the random effect estimate should occur at the peak log-likelihood value (intersection of the two dotted lines). Profile likelihood plots shown are for the A) σ²₁ parameter, representing the random effect of study; B) σ²₂ parameter, representing the random effect of residual variance due to multiple effect sizes estimates per sampling event; C) τ² parameter, representing the random effect of study arm (outer term); D) ρ parameter, representing the random effect temporal autocorrelation within study arms (inner term)

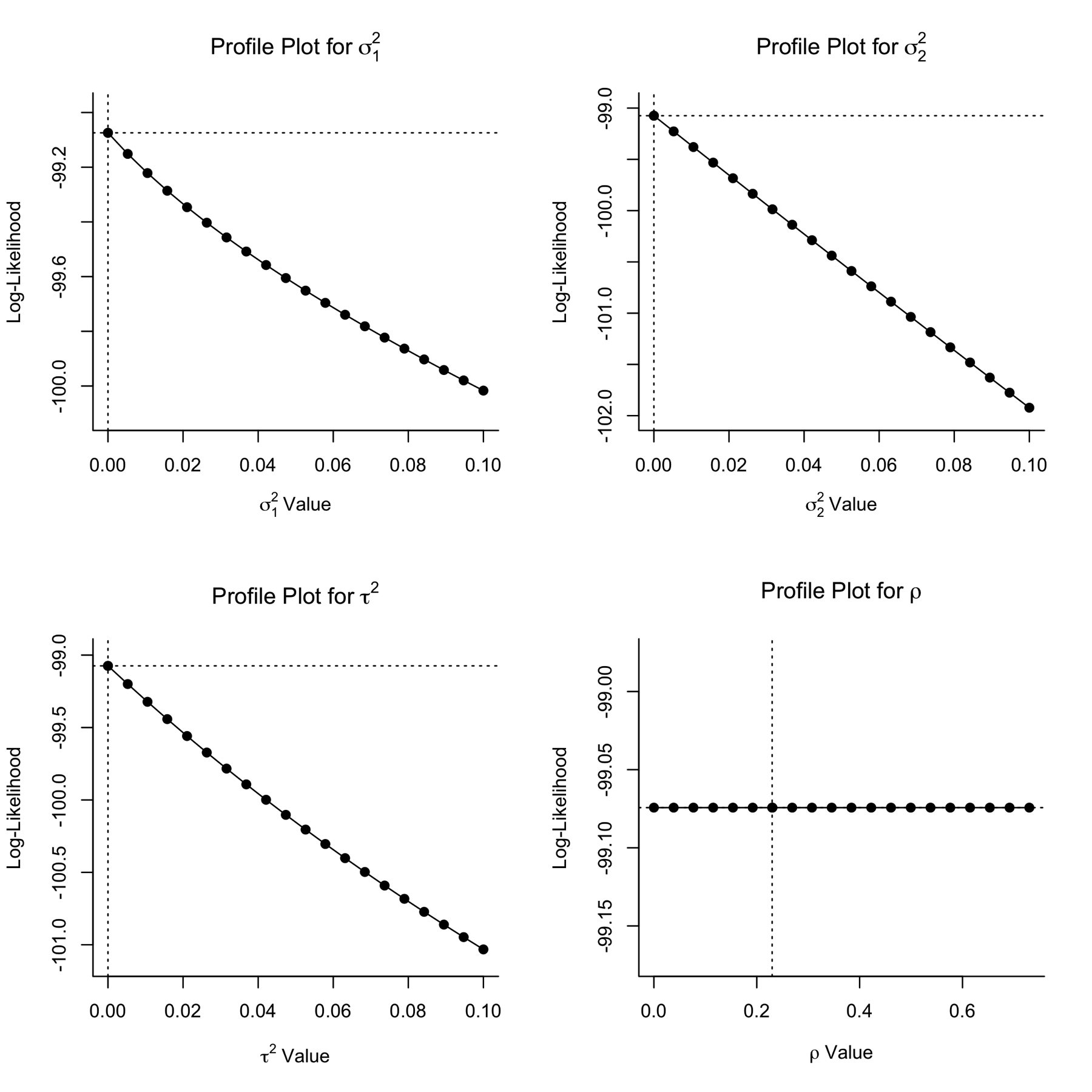

Supplementary Figure 30: Profile likelihood plots for each of the random effect estimates from the intercept-only model of the overall effect after the administration antibiotics. Where the random effect is accurately estimated, the random effect estimate should occur at the peak log-likelihood value (intersection of the two dotted lines). Profile likelihood plots shown are for the A) σ²₁ parameter, representing the random effect of study; B) σ²₂ parameter, representing the random effect of residual variance due to multiple effect sizes estimates per sampling event; C) τ² parameter, representing the random effect of study arm (outer term); D) ρ parameter, representing the random effect temporal autocorrelation within study arms (inner term)

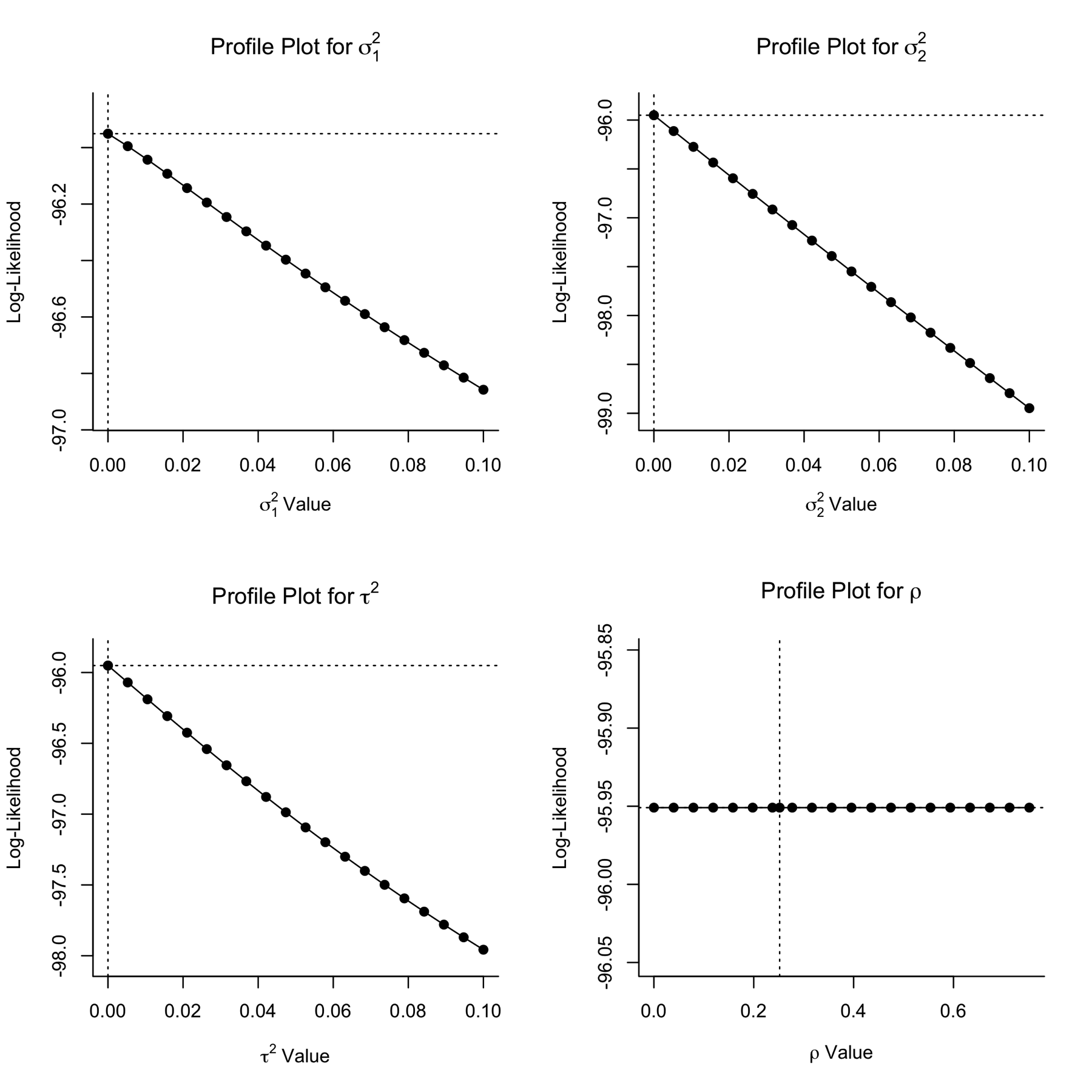

Supplementary Figure 31: Plots representing results of meta-analytical models testing our two hypotheses for effects measured during the administration of antibiotics. In these plots, colours of the points represent the studies from which they are derived, and their size is proportional to their inverse standard error (1/SE; small points are less precise estimates, larger points are more precise estimates). A) Orchard plot representing the modelled overall effect of antibiotic administration on the absolute abundance of resistant bacteria/resistance genes relating to the intervention antibiotic class in the targeted bacterial population. This overall effect alongside its heterogeneity estimate, are derived from the intercept-only model. In this plot, the overall effect (model intercept after accounting for random effects) is represented by the 'trunk' of the tree (black circle), with its confidence interval represented its 'branch' (shorter, thicker line through it), and its prediction interval represented by its 'twig' (longer, thinner line through it).B) Bubble plot representing the modelled relationship between non-linear timing of sampling (days of continuous antibiotic administration) and effect size (SMDH) from the meta-regression model. In this plot, the modelled effect of time is represented by the bold line, with the dashed line and dotted lines representing the 95% confidence and prediction intervals, respectively.

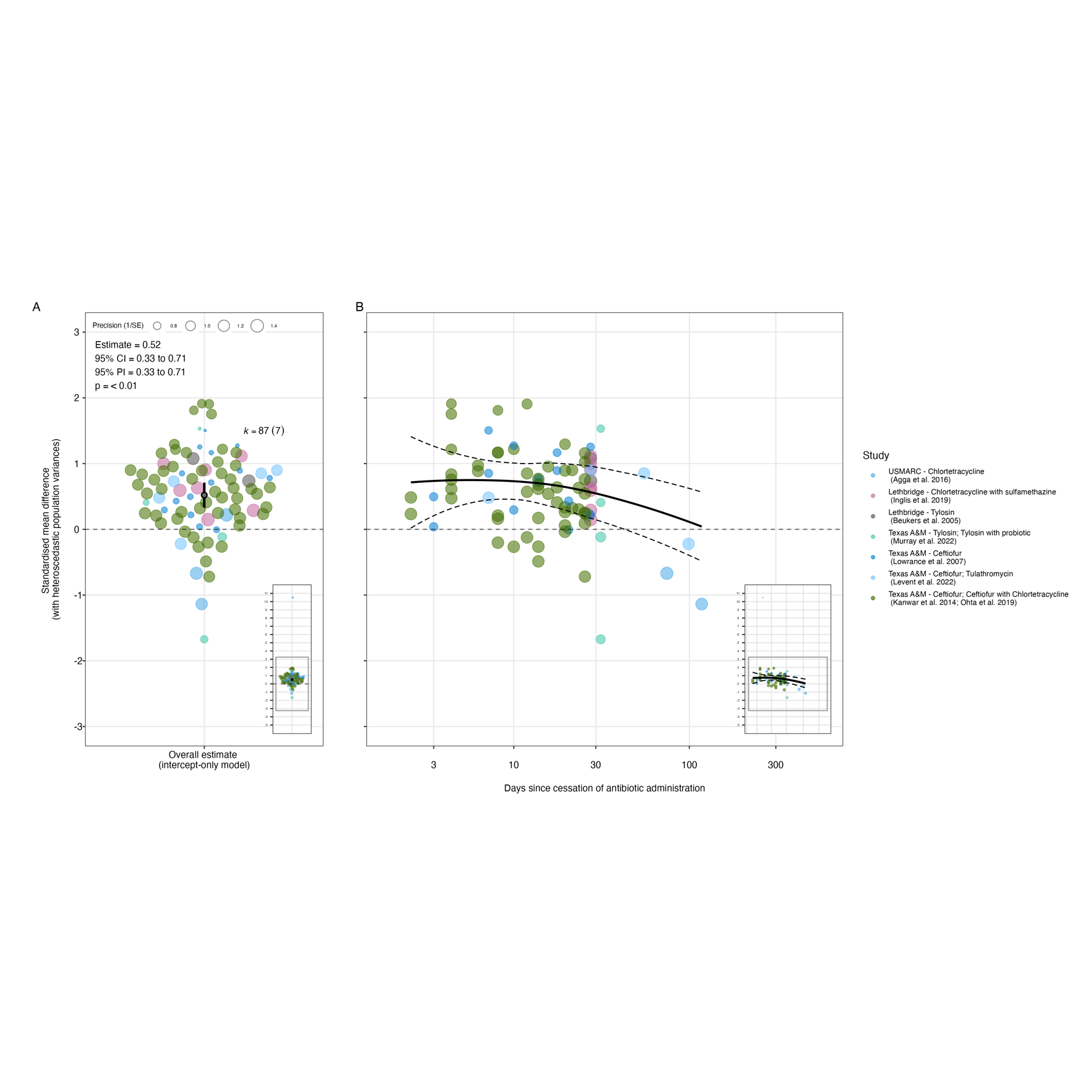

Supplementary Figure 32: Plots representing results of meta-analytical models testing our two hypotheses for effects on the absolute abundance of all determinants measured after the administration of antibiotics. In these plots, colours of the points represent the studies from which they are derived, and their size is proportional to their inverse standard error (1/SE; small points are less precise estimates, larger points are more precise estimates). A) Orchard plot representing the modelled overall effect of antibiotic administration on the absolute abundance of resistant bacteria/resistance genes relating to the intervention antibiotic class in the targeted bacterial population. This overall effect alongside its heterogeneity estimate, are derived from the intercept-only model. In this plot, the overall effect (model intercept after accounting for random effects) is represented by the 'trunk' of the tree (black circle), with its confidence interval represented its 'branch' (shorter, thicker line through it), and its prediction interval represented by its 'twig' (longer, thinner line through it). B) Bubble plot representing the modelled moderating effect of linear timing of sampling (days of continuous antibiotic administration) and residual heterogeneity from the meta-regression model. In this plot, the modelled effect of time is represented by the bold line, with the dashed line and dotted lines representing the 95% confidence and prediction intervals, respectively.

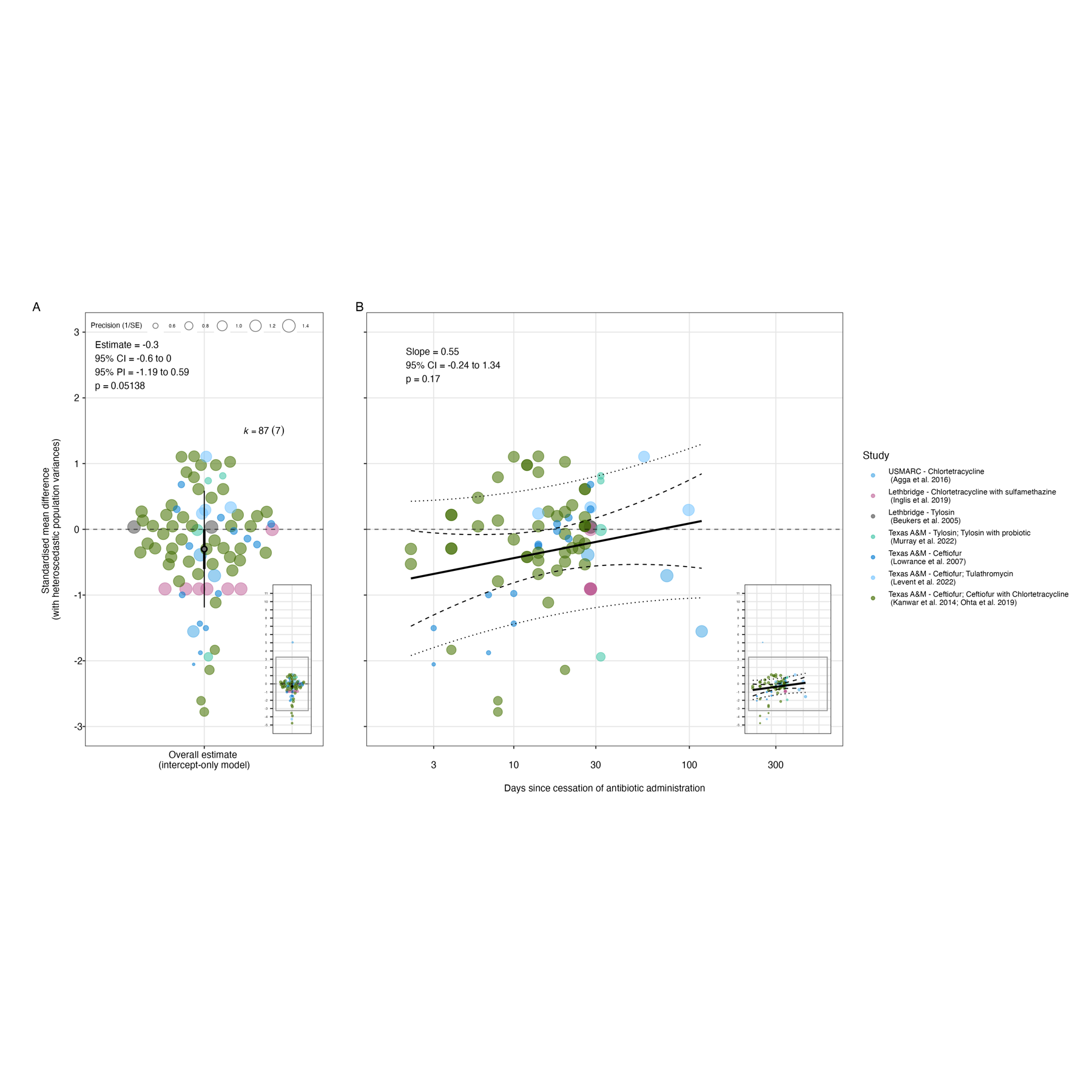

Supplementary Figure 33: Plots representing results of meta-analytical models testing our two hypotheses for effects on the absolute abundance of all determinants measured after the administration of antibiotics. In these plots, colours of the points represent the studies from which they are derived, and their size is proportional to their inverse standard error (1/SE; small points are less precise estimates, larger points are more precise estimates). A) Orchard plot representing the modelled overall effect of antibiotic administration on the absolute abundance of resistant bacteria/resistance genes relating to the intervention antibiotic class in the targeted bacterial population. This overall effect alongside its heterogeneity estimate, are derived from the intercept-only model. In this plot, the overall effect (model intercept after accounting for random effects) is represented by the 'trunk' of the tree (black circle), with its confidence interval represented its 'branch' (shorter, thicker line through it), and its prediction interval represented by its 'twig' (longer, thinner line through it). B) Bubble plot representing the modelled moderating effect of non-linear timing of sampling (days of continuous antibiotic administration) and residual heterogeneity from the meta-regression model. In this plot, the modelled effect of time is represented by the bold line, with the dashed line and dotted lines representing the 95% confidence and prediction intervals, respectively.

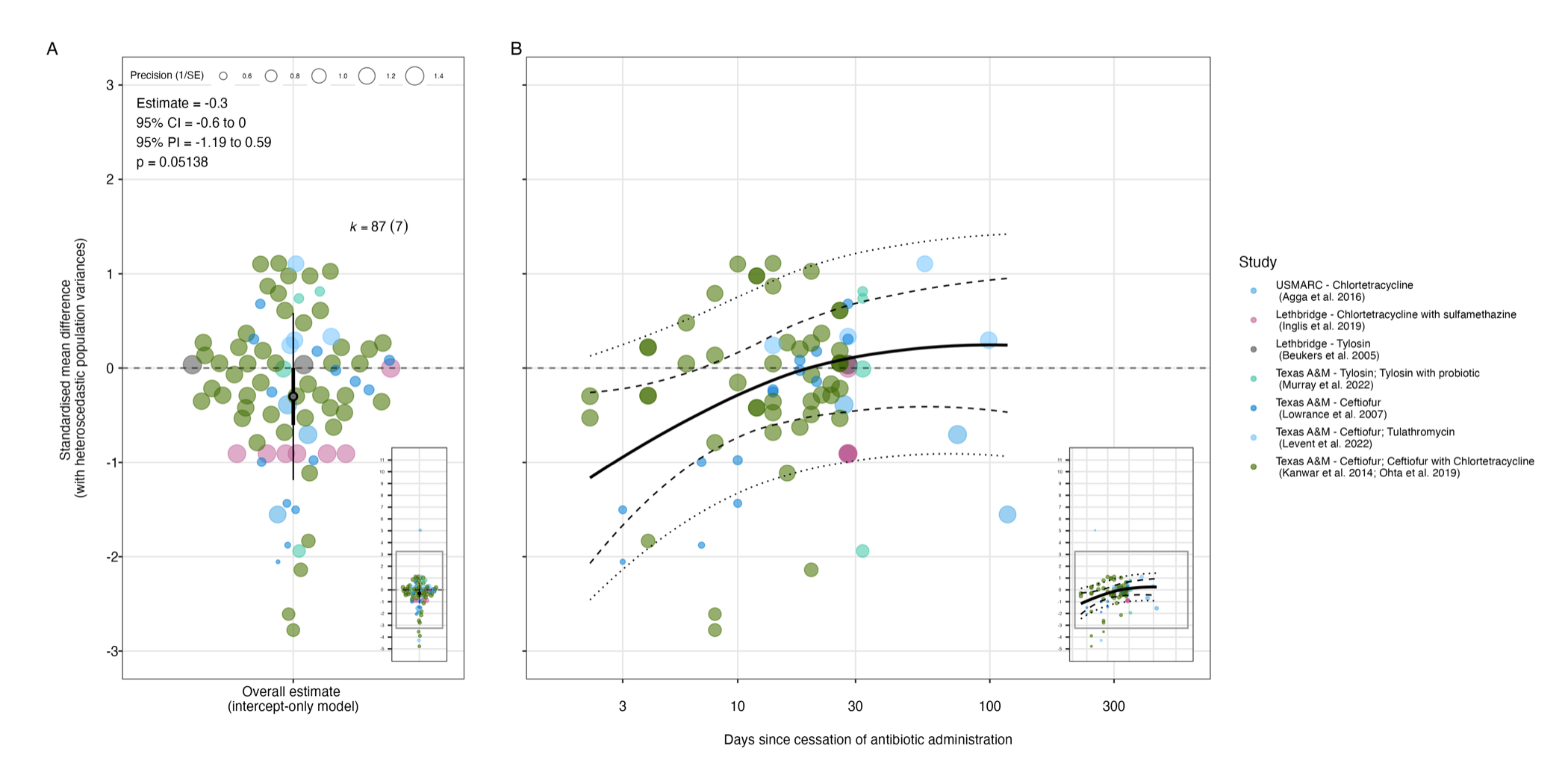

###### 2.4.2.2 Sensitivity testing

Supplementary Figure 34: Orchard plots representing the sensitivity of the modelled overall effect after the cessation of antibiotic administration to leaving one study out. In these plots, the mean overall effect is represented by the 'trunk' of the tree (black circle), with its confidence interval represented its 'branch' (shorter, thicker line through it), and its prediction interval represented by its 'twig' (longer, thinner line through it). The mean and precision (1/standard error) of the individual effect sizes from studies included in the model is represented by the position and size of the coloured points, respectively.

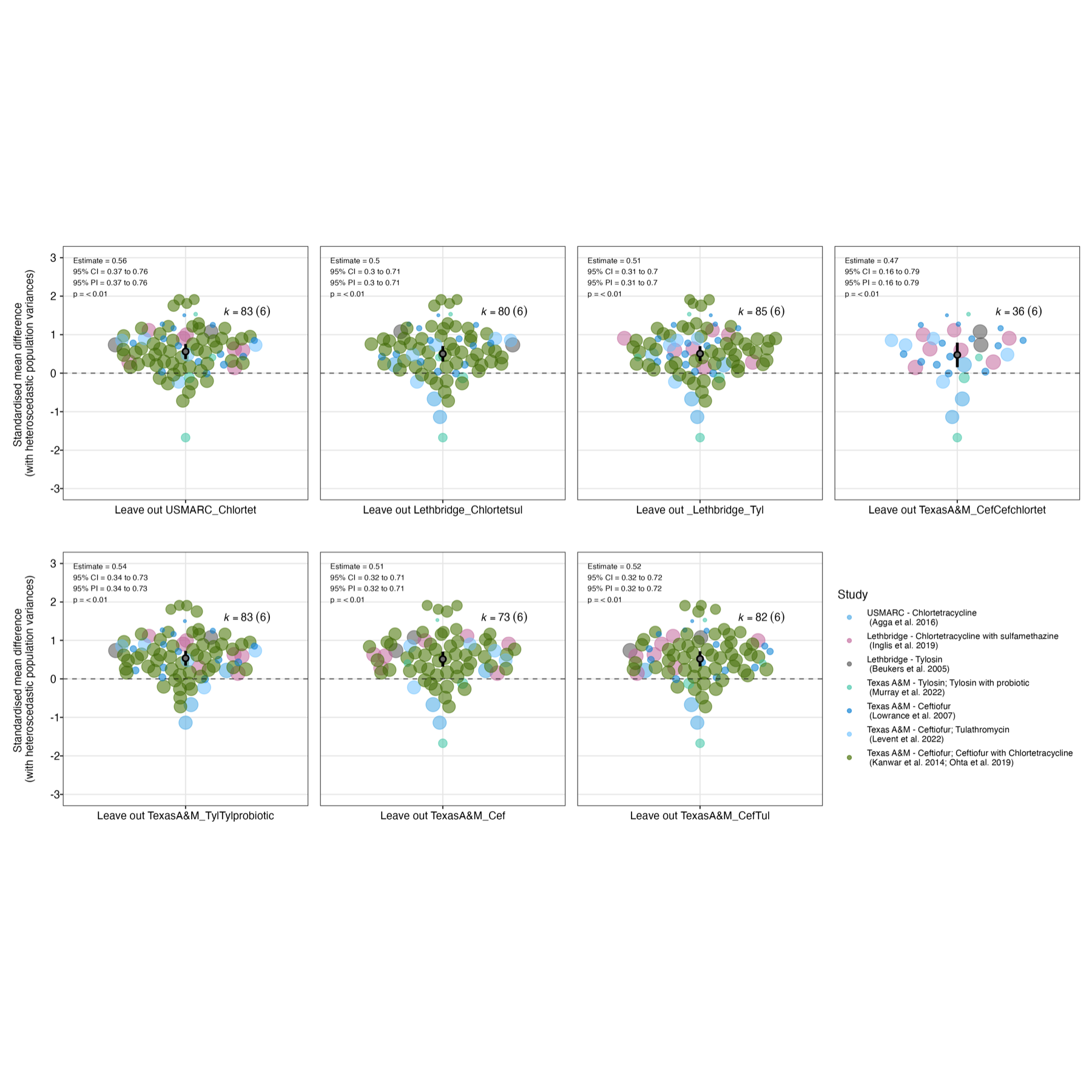

Supplementary Figure 35: Orchard plots representing the sensitivity of the modelled overall effect during the administration of antibiotics to correcting for shared control non-independence and the removal of potentially problematic studies. In these plots, the mean overall effect is represented by the 'trunk' of the tree (black circle), with its confidence interval represented its 'branch' (shorter, thicker line through it), and its prediction interval represented by its 'twig' (longer, thinner line through it). The mean and precision (1/standard error) of the individual effect sizes from studies included in the model is represented by the position and size of the coloured points, respectively.

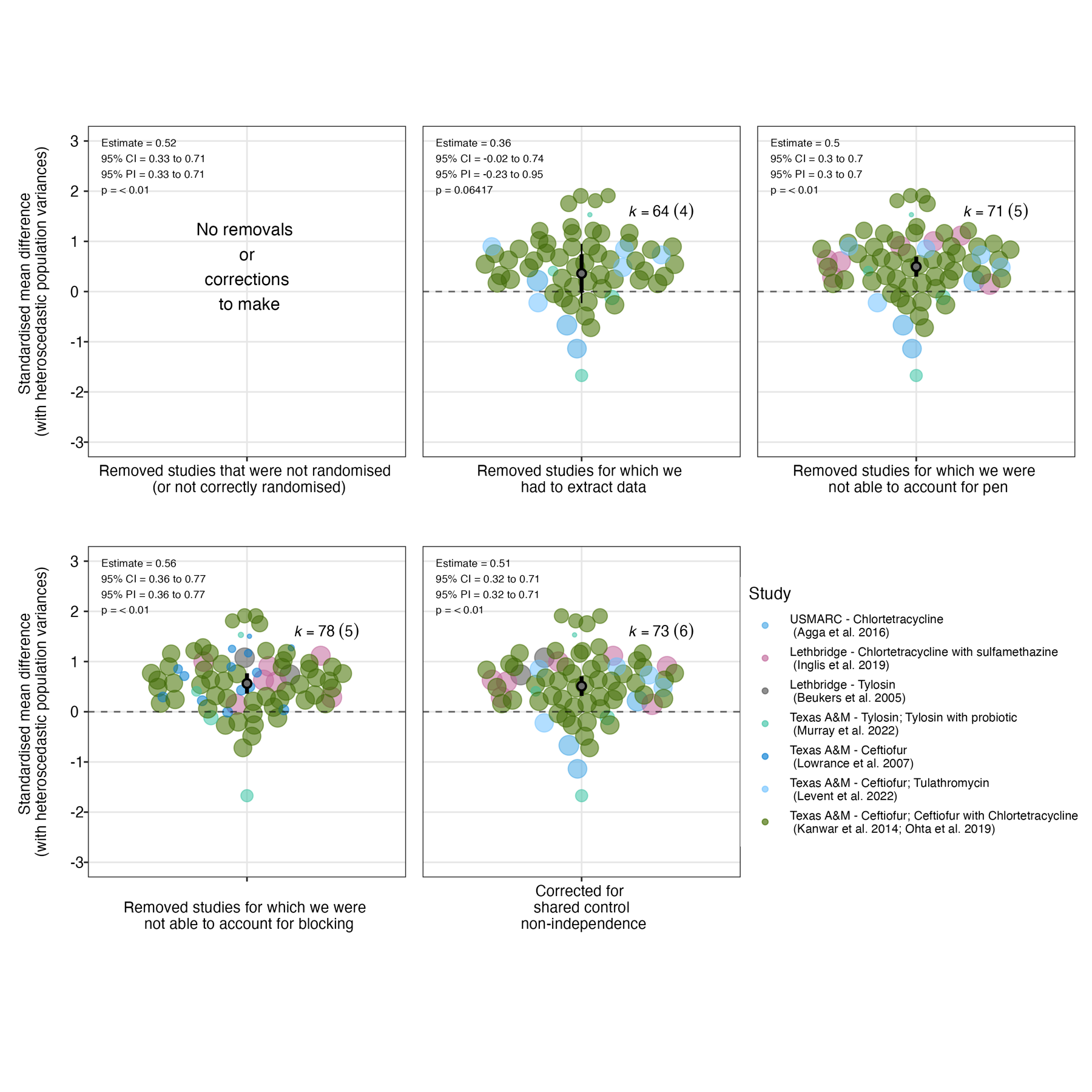

Supplementary Figure 36: Bubble plots representing the sensitivity of the modelled moderating effect of timing of sampling after the cessation of antibiotic administration to leaving one study out. In these plots, the modelled effect of time is represented by the bold line, with the dashed line and dotted lines representing the 95% confidence and prediction intervals, respectively. The mean and precision (1/standard error) of the individual effect sizes from studies included in the model is represented by the position and size of the coloured points, respectively.

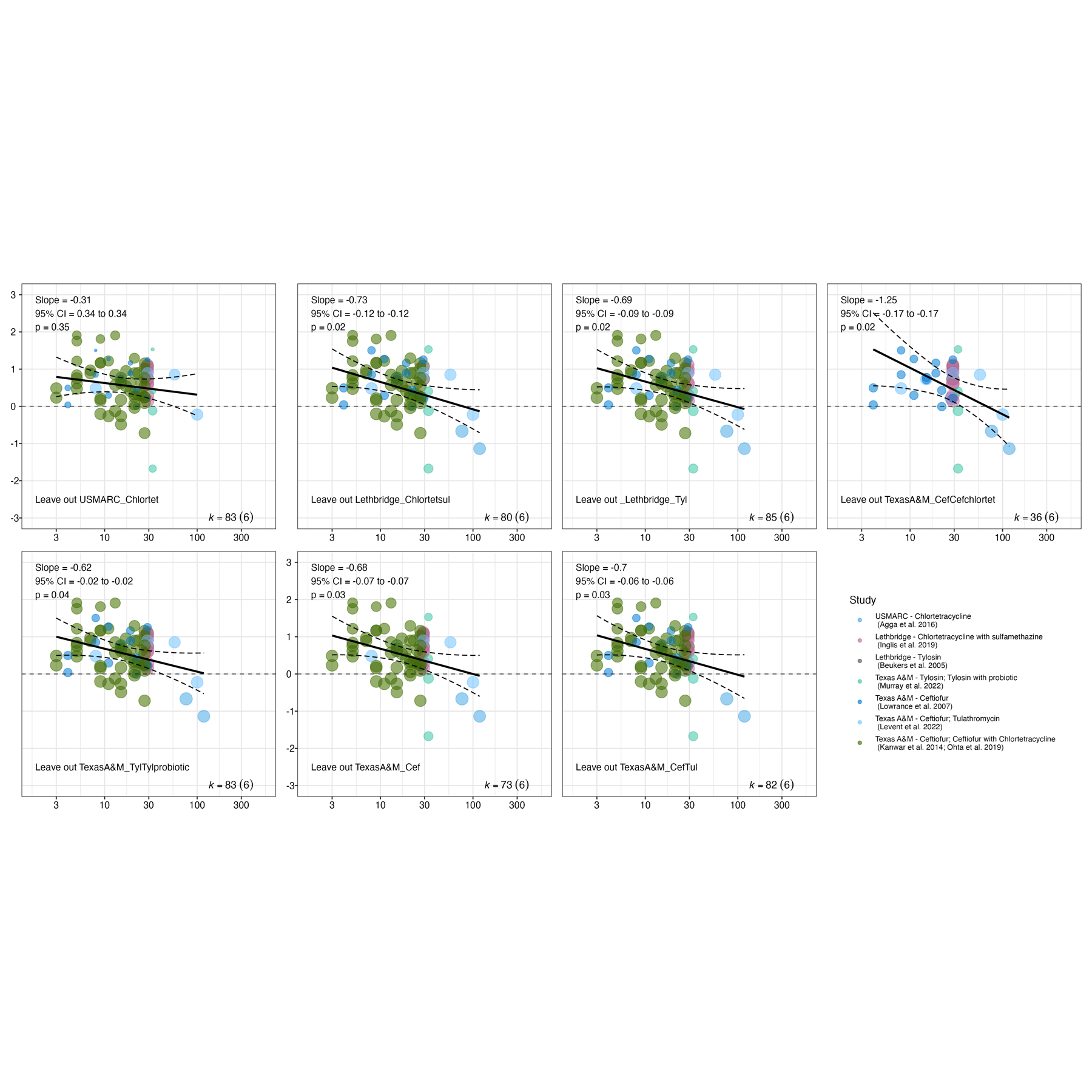

Supplementary Figure 37: Bubble plots representing the sensitivity of the modelled overall effect after the administration of antibiotics to correcting for shared control non-independence and the removal of potentially problematic studies. In these plots, the modelled effect of time is represented by the bold line, with the dashed line and dotted lines representing the 95% confidence and prediction intervals, respectively. The mean and precision (1/standard error) of the individual effect sizes from studies included in the model is represented by the position and size of the coloured points, respectively.

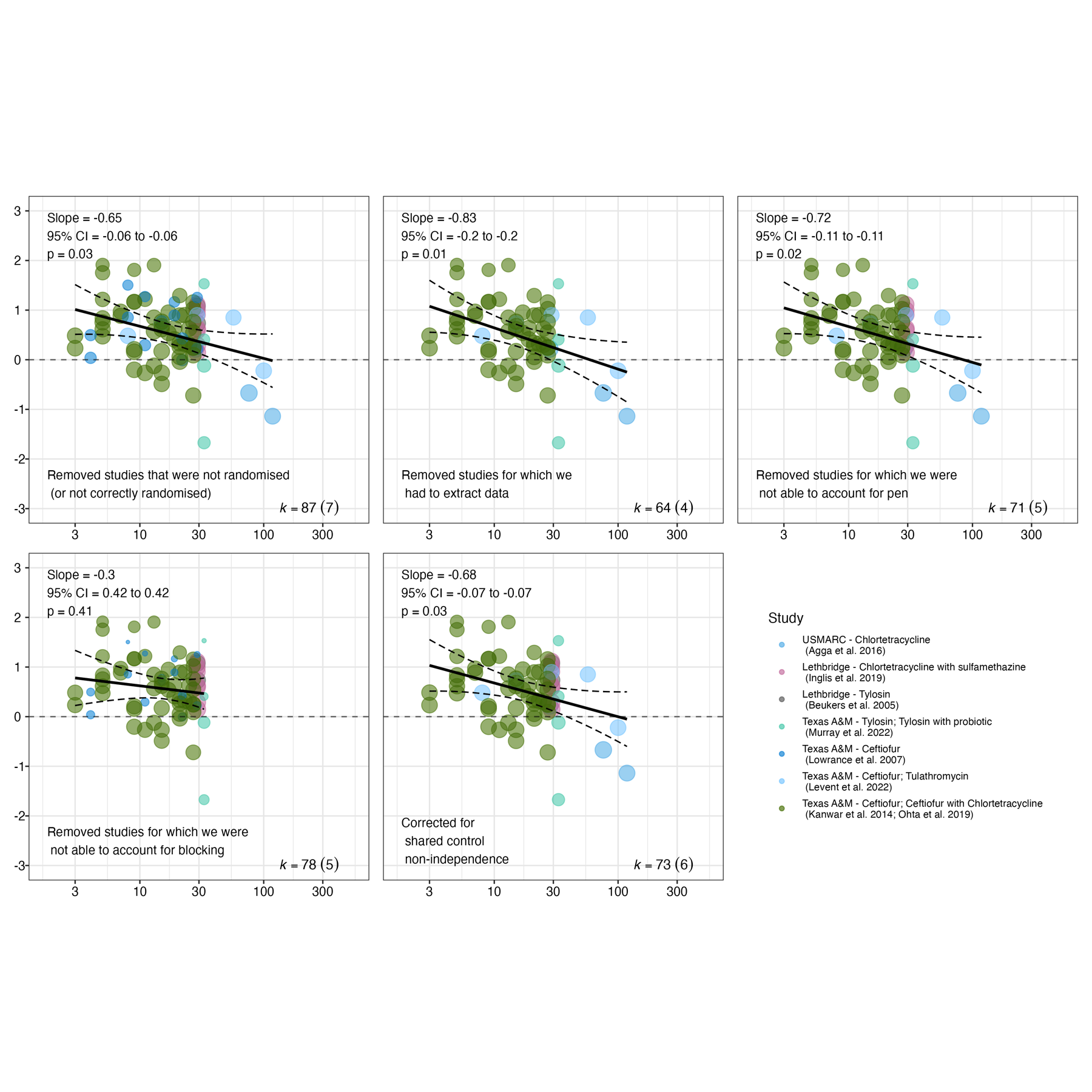

Supplementary Figure 38: Orchard plots representing the sensitivity of the modelled overall effect after the administration of antibiotics to assumptions about the correlation between effect sizes from the same study. Orchard plots of the original model and assuming correlations of 0, 0.2, 0.4, 0.6 and 0.8 are shown. In these plots, the mean overall effect is represented by the 'trunk' of the tree (black circle), with its confidence interval represented its 'branch' (shorter, thicker line through it), and its prediction interval represented by its 'twig' (longer, thinner line through it).

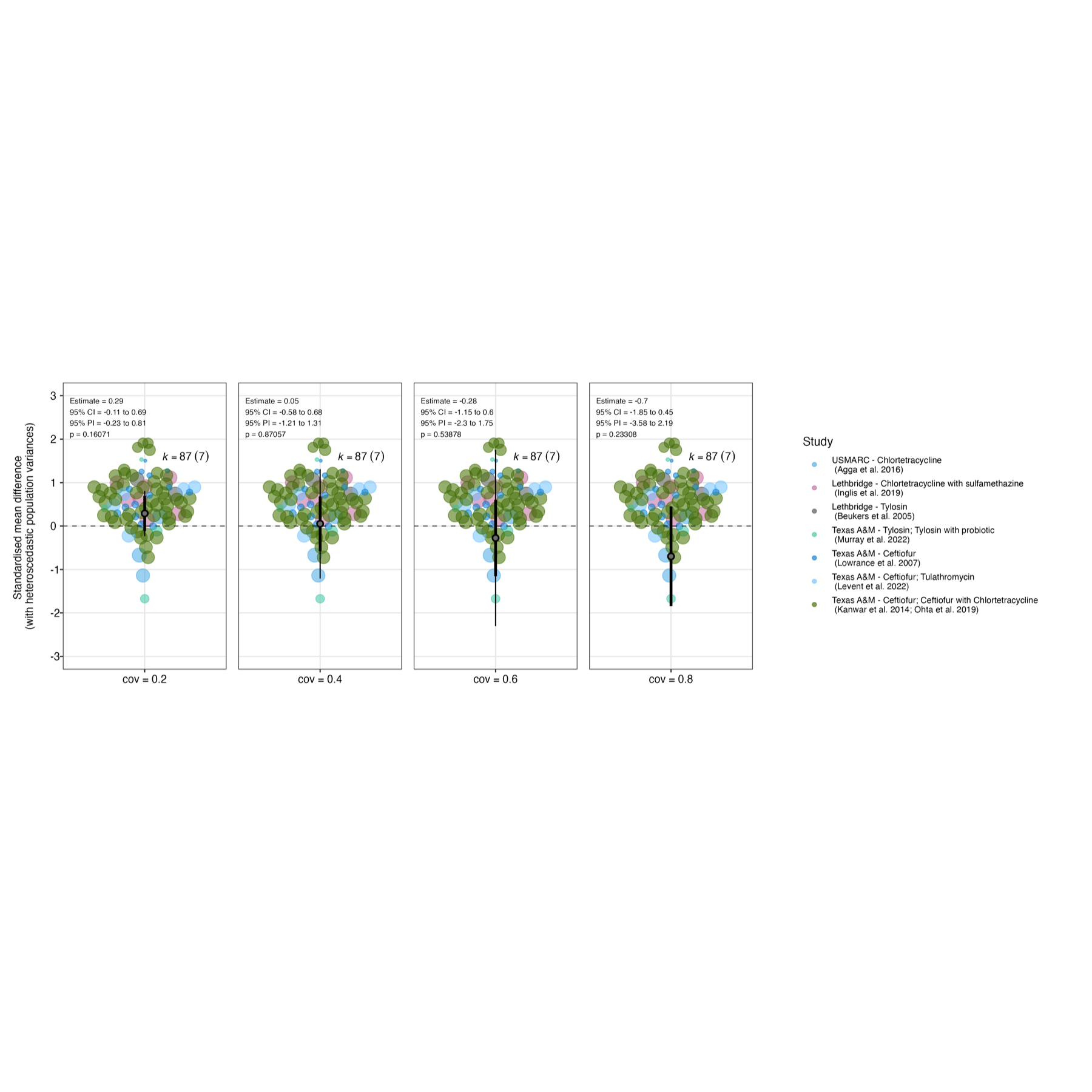

Supplementary Figure 39: Bubble plots representing the sensitivity of the moderating effect of timing of sampling on the effect after the administration of antibiotics to assumptions about the correlation between effect sizes from the same study. Bubble plots of the original model and assuming correlations of 0, 0.2, 0.4, 0.6 and 0.8 are shown. In these plots, the modelled effect of time is represented by the bold line, with the dashed line and dotted lines representing the 95% confidence and prediction intervals, respectively. The mean and precision (1/standard error) of the individual effect sizes from studies included in the model is represented by the position and size of the coloured points, respectively.

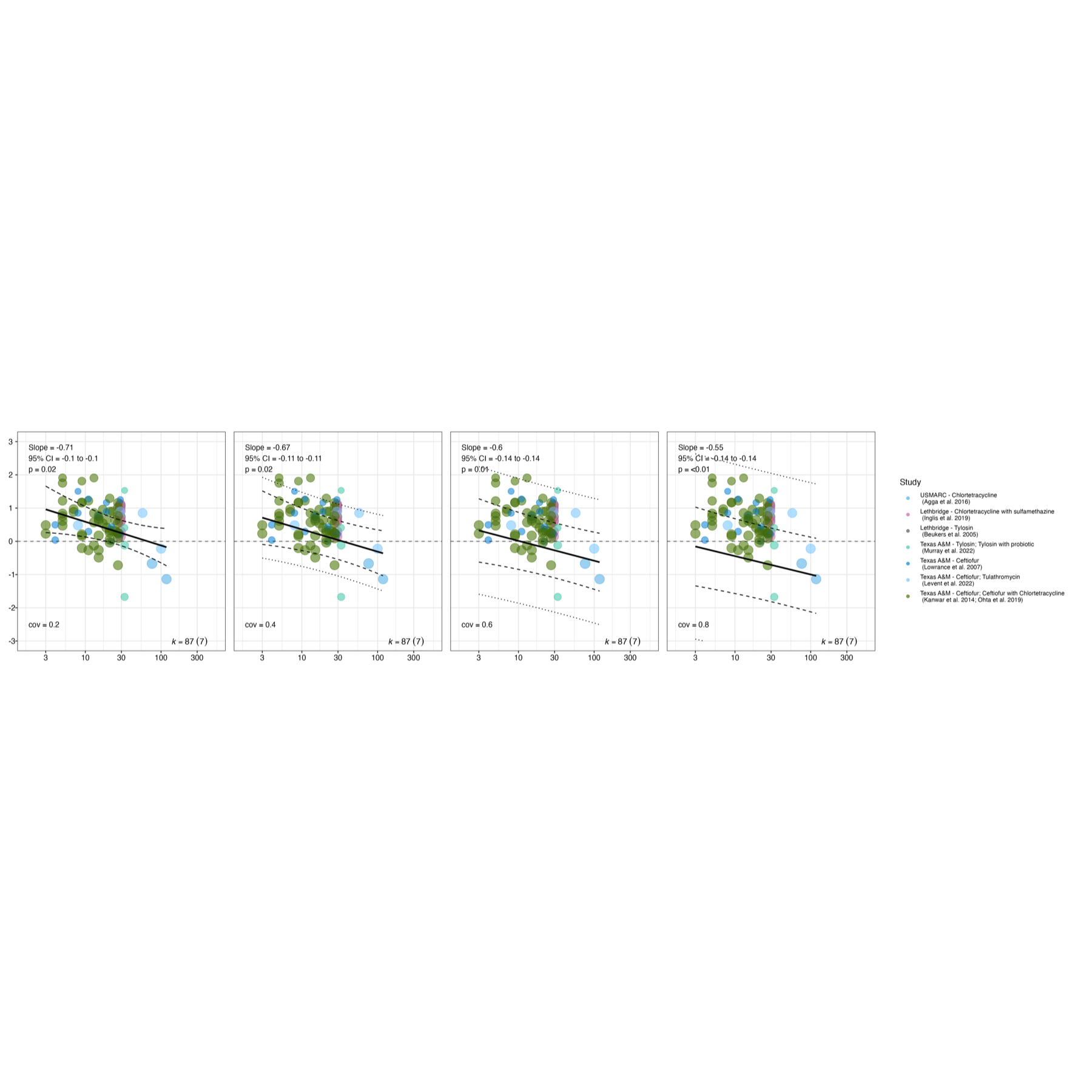

###### 2.4.2.3 Subgroup and other alternative analyses

Supplementary Information 18: Full results of subgroup analyses

**Population**

All study populations included in the quantitative synthesis were feedlot steers, precluding a subgroup analysis by production system. All other population characteristics that we attempted to extract from publications (cattle reproductive class, breed, weight, age, and antibiotic treatment history prior to the study) were poorly reported (resulting in lots of missing metadata) and/or too heterogeneous (e.g. a mix of reproductive classes in each study unit) to permit reasonable alternative meta-regressions based on them.

**Interventions and controls**

Subgroup modelling according to class of the antibiotic administered in the intervention detected a significant positive effect for the two most well-studied classes of antibiotics (Figure 5). In macrolides, lincosamides and streptogramin B/MLSB (which made up the majority of measurements taken during antibiotic administration) the size of the effect sensu Cohen^37^ was small-to-medium (Estimate = 0.38; 95% CI 0.12 to 0.65; 95% PI -0.33 to 1.1; p = <0.01). In extended-spectrum cephalosporins (which made up the majority of measurements taken after antibiotic administration) the size of the effect was medium (Estimate = 0.57; 95% CI 0.27 to 0.87; 95% PI -0.16 to 1.29; p = <0.01). Tetracyclines and sulfonamides were represented by two studies and one study, respectively, and no significant effect was detected in either class (Figure 5).

The subgroup analysis by mode of administration produced similar results, given mode of administration was confounded by class of antibiotic administered (Supplementary Figure 40). The pooled effect of feeding antibiotics (which was typically assessed during tylosin administration) was small-to-medium (0.37; 95% CI 0.16 to 0.59; 95% PI -0.29 to 1.04; p = <0.01), whilst the pooled effect of injecting antibiotics (which was typically assessed after ceftiofur administration) was medium (0.57; 95% CI 0.27 to 0.86; 95% PI -0.13 to 1.26; p = <0.01).

Administering the intervention antibiotic in a pen also containing control cattle, or administering the intervention antibiotic alongside a cointervention antibiotic (defined as another antibiotic given to both intervention and control groups) did not appear to moderate the effect (Supplementary Figures 41 - 42). There was some evidence of a positive relationship between the cumulative dose of antibiotic administered before resistance measurement and effect size (Supplementary Figure 43).

**Outcome**

A subgroup analysis of target microorganisms detected a positive overall effect in all groups (Supplementary Figure 44). The pooled effect increased in strength from Enterobacteriaceae (Estimate = 0.3; 95% CI 0.03 to 0.56; 95% PI -0.36 to 0.95; p = 0.03) to Enterococcaceae (Estimate = 0.5; 95% CI 0.25 to 0.74; 95% PI -0.15 to 1.15, p = <0.01) to qPCR measures of bacteria generally ( Estimate = 0.53; 95% CI 0.22 to 0.84; 95% PI -0.15 to 1.21; p = <0.01).

A subgroup analysis of resistance measurement method indicated a positive small-to-medium size effect in outcomes measured by culture-based direct plating (Estimate = 0.4; 95% CI 0.2 to 0.61; 95% PI -0.26 to 1.07; p = <0.01) and a slightly larger small-to-medium size effect in culture-independent qPCR measures of resistance (Estimate = 0.53; 95% CI 0.21 to 0.85; 95% PI -0.36 to 1.24, p = <0.01), though the latter subgroup was only represented by two studies (Supplementary Figure 45). We also performed sub-subgroup analyses to explore the differences within direct plating and qPCR approaches further (Supplementary Figures 46 – 47. Within direct plating approaches, an effect was detected for the best-represented groups ceftiofur (Estimate = 0.67; 95% CI 0.22 to 1.13; 95% PI -0.18 to 1.52, p = <0.01) and erythromycin (Estimate = 0.36; 95% CI 0.07 to 0.65; 95% PI -0.41 to 1.14, p = 0.01). Analyses of the concentration of the antibiotics onto which samples were plated to assess resistance (i.e. study resistance breakpoint) suggested it may partially moderate the effect, with some evidence that effects might not be detected when tested at a study resistance breakpoint above the CLSI/NARMS breakpoint (Supplementary Figure 48). However, we did not find a linear relationship between the study breakpoint relative to the CLSI/NARMs breakpoint and effect size (Supplementary Figure 49). We also did not find a linear relationship between log-transformed study breakpoint and effect size (Supplementary Figure 50). Within qPCR approaches, there was no clear pattern in effect sizes (e.g. by antibiotic class) though only two studies used qPCR approaches with no overlap in the genes studied between them.

Finally, we explored potential moderation of the effect according to the unit of analysis in the study (Supplementary Figure 51). Animal was regarded as the unit of analysis for one study for which there was only one pen per treatment^29^. In this study, we found a medium effect (Estimate = 0.27; 95% CI -0.18 to 0.73; 95% PI to 1.06, p = 0.2). In the remaining studies, there were multiple pens per treatment and therefore pen was regarded as the correct unit of analysis. In these studies, there was also small-to-medium positive overall effect (Estimate = 0.46; 95% CI 0.27 to 0.65; 95% PI -0.21 to 1.14, p = <0.01). Among the 10 studies where pen was the correct unit of analysis, the potential influence of unit-of-analysis errors which we were not able to correct was tested for separately in the previously-reported sensitivity analyses.

**Study design**

Our subgroup model did not suggest strong moderation of the effect due to overall study design (Supplementary Figure 52), with a similar small-to-medium effect for completely randomised (Estimate = 0.58; 95% CI 0.35 to 0.81; 95% PI -0.06 to 1.21; p = <0.01) and randomised complete block (RCB) designs (Estimate = 0.32; 95% CI 0.03 to 0.62, 95% PI -0.34 to 0.98, p = 0.03). The only non-randomised study^19^ was reported as ‘completely randomised design’ but this only applied to a more secondary intervention; see Risk of Bias assessment), and was not found to be significantly different from randomised designs despite its lower point estimate (Estimate = 0.21; 95% CI -0.24 to 0.66, 95% PI -0.53 to 0.96, p = 0.4).

Supplementary Figure 40: Orchard plot representing results of the meta-analytical model testing for a subgroup effect of mode of administering the intervention groups (via feed or via injection). Points are effect sizes, with their colours representing the studies from which they are derived and their size being proportional to their inverse standard error (1/SE; small points are less precise estimates, larger points are more precise estimates). The pooled effect for each subgroup (coefficient after accounting for random effects) is represented by the 'trunk' of the tree (black circle), with its confidence interval represented its 'branch' (shorter, thicker line through it), and its prediction interval represented by its 'twig' (longer, thinner line through it). For the numbers above each subgroup, 'k' represents the number of effect sizes, whilst the number in brackets represents the number of studies from which they were derived.

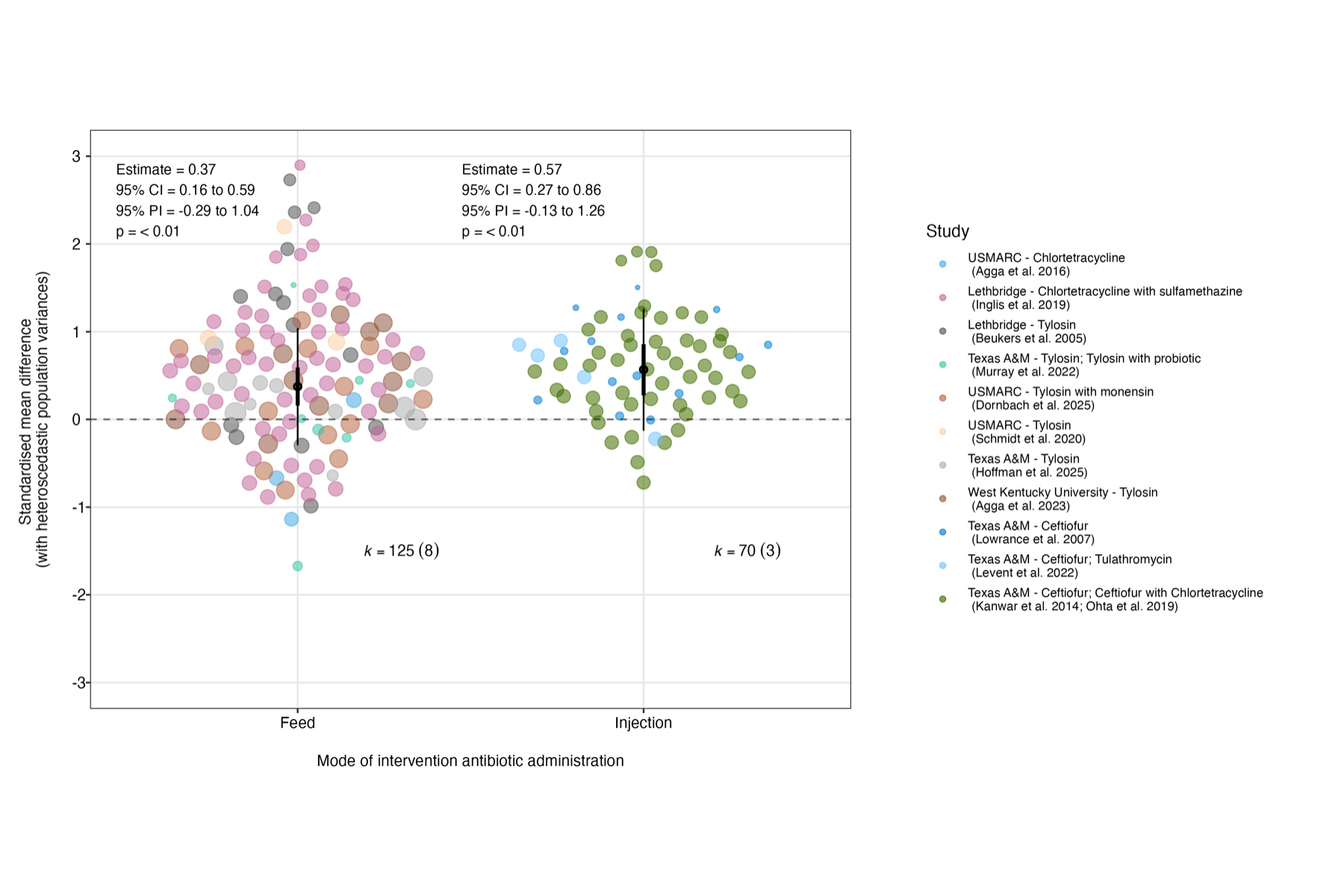

Supplementary Figure 41: Orchard plot representing results of the meta-analytical model testing for a subgroup effect of whether or not intervention cattle were housed alongside control cattle. Points are effect sizes, with their colours representing the studies from which they are derived and their size being proportional to their inverse standard error (1/SE; small points are less precise estimates, larger points are more precise estimates). The pooled effect for each subgroup (coefficient after accounting for random effects) is represented by the 'trunk' of the tree (black circle), with its confidence interval represented its 'branch' (shorter, thicker line through it), and its prediction interval represented by its 'twig' (longer, thinner line through it). For the numbers above each subgroup, 'k' represents the number of effect sizes, whilst the number in brackets represents the number of studies from which they were derived.

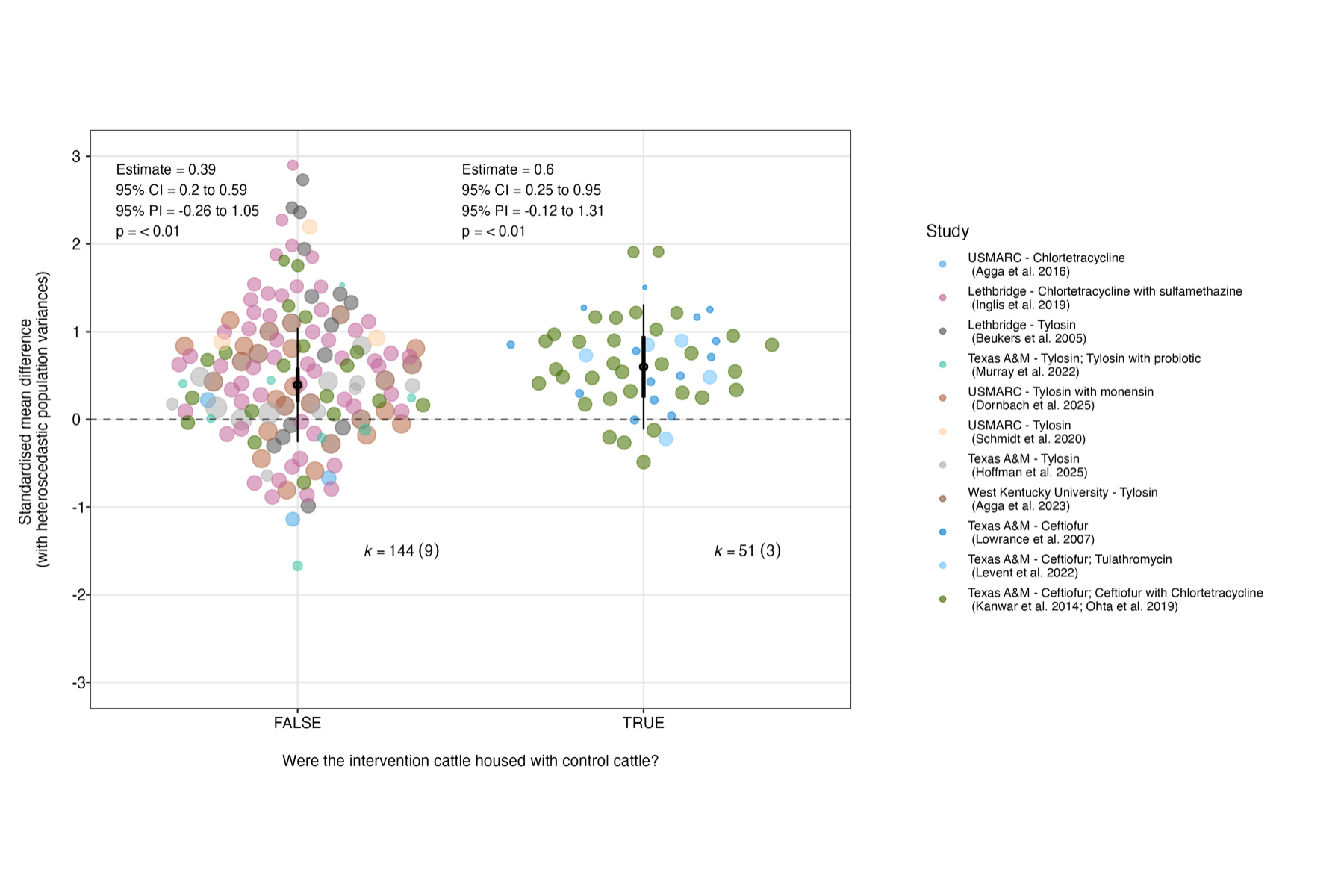

Supplementary Figure 42: Orchard plot representing results of the meta-analytical model testing for a subgroup effect of whether or not another antibiotic was administered to intervention (and control) groups alongside the intervention antibiotic (i.e. presence of a cointervention). Points are effect sizes, with their colours representing the studies from which they are derived and their size being proportional to their inverse standard error (1/SE; small points are less precise estimates, larger points are more precise estimates). The pooled effect for each subgroup (coefficient after accounting for random effects) is represented by the 'trunk' of the tree (black circle), with its confidence interval represented its 'branch' (shorter, thicker line through it), and its prediction interval represented by its 'twig' (longer, thinner line through it). For the numbers above each subgroup, 'k' represents the number of effect sizes, whilst the number in brackets represents the number of studies from which they were derived.

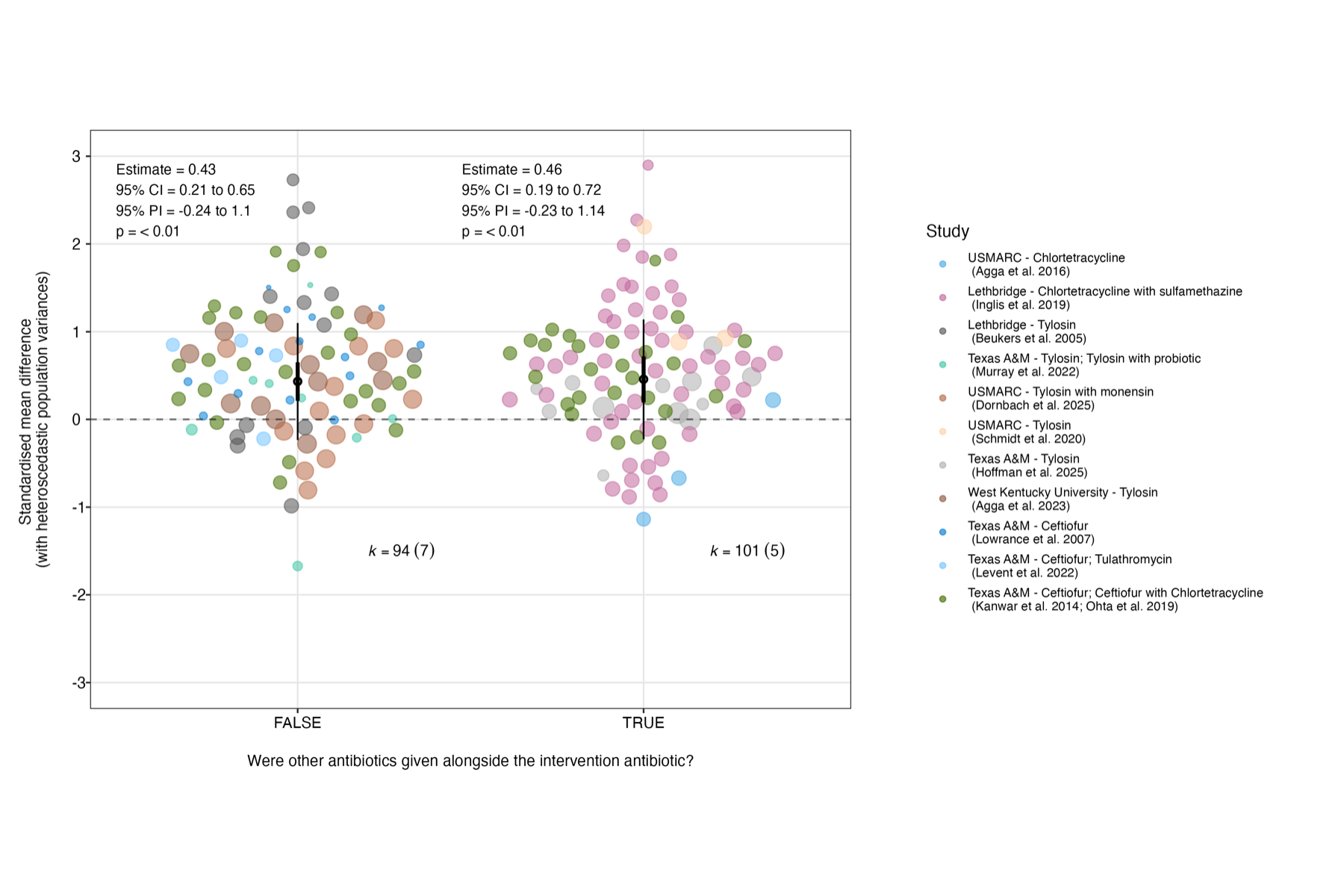

Supplementary Figure 43: Bubble plot representing the results of meta-analytical models testing for an effect of dose. Points are effect sizes, with their colours representing the studies from which they are derived and their size being proportional to their inverse standard error (1/SE; small points are less precise estimates, larger points are more precise estimates). Plots the meta-regression fits of tests of dose (mg/kg) in the following formats: A) Daily dose of antibiotic administered B) Log10-transformed Daily dose of antibiotic administered C) Cumulative daily dose of antibiotic administered before each effect estimate D) Log10-transformed cumulative daily dose of antibiotic administered before each effect estimate.

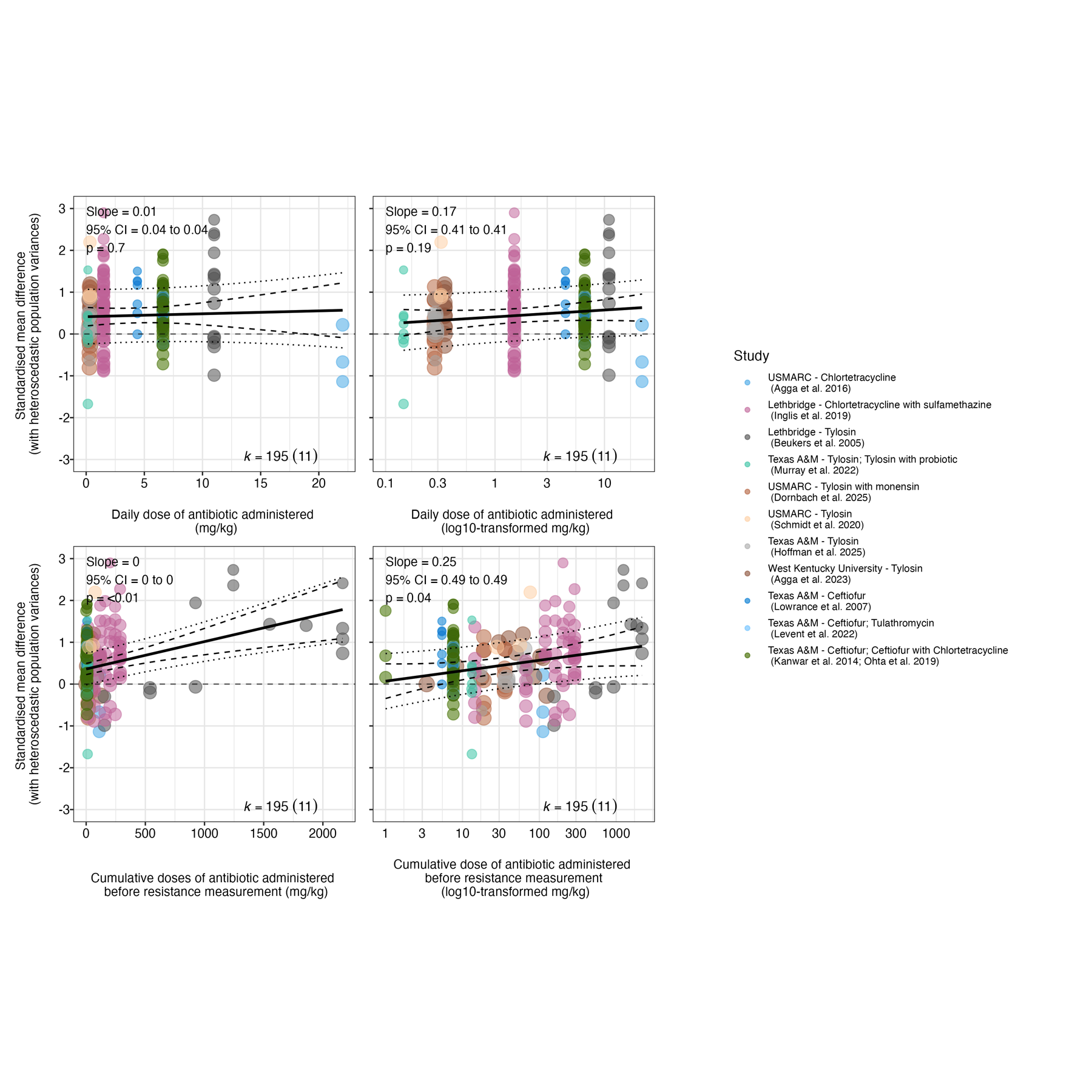

Supplementary Figure 44: Orchard plot representing results of the meta-analytical model testing for a subgroup effect of the microorganism(s) targetted in the outcome assessment. Points are effect sizes, with their colours representing the studies from which they are derived and their size being proportional to their inverse standard error (1/SE; small points are less precise estimates, larger points are more precise estimates). The pooled effect for each subgroup (coefficient after accounting for random effects) is represented by the 'trunk' of the tree (black circle), with its confidence interval represented its 'branch' (shorter, thicker line through it), and its prediction interval represented by its 'twig' (longer, thinner line through it). For the numbers above each subgroup, 'k' represents the number of effect sizes, whilst the number in brackets represents the number of studies from which they were derived.

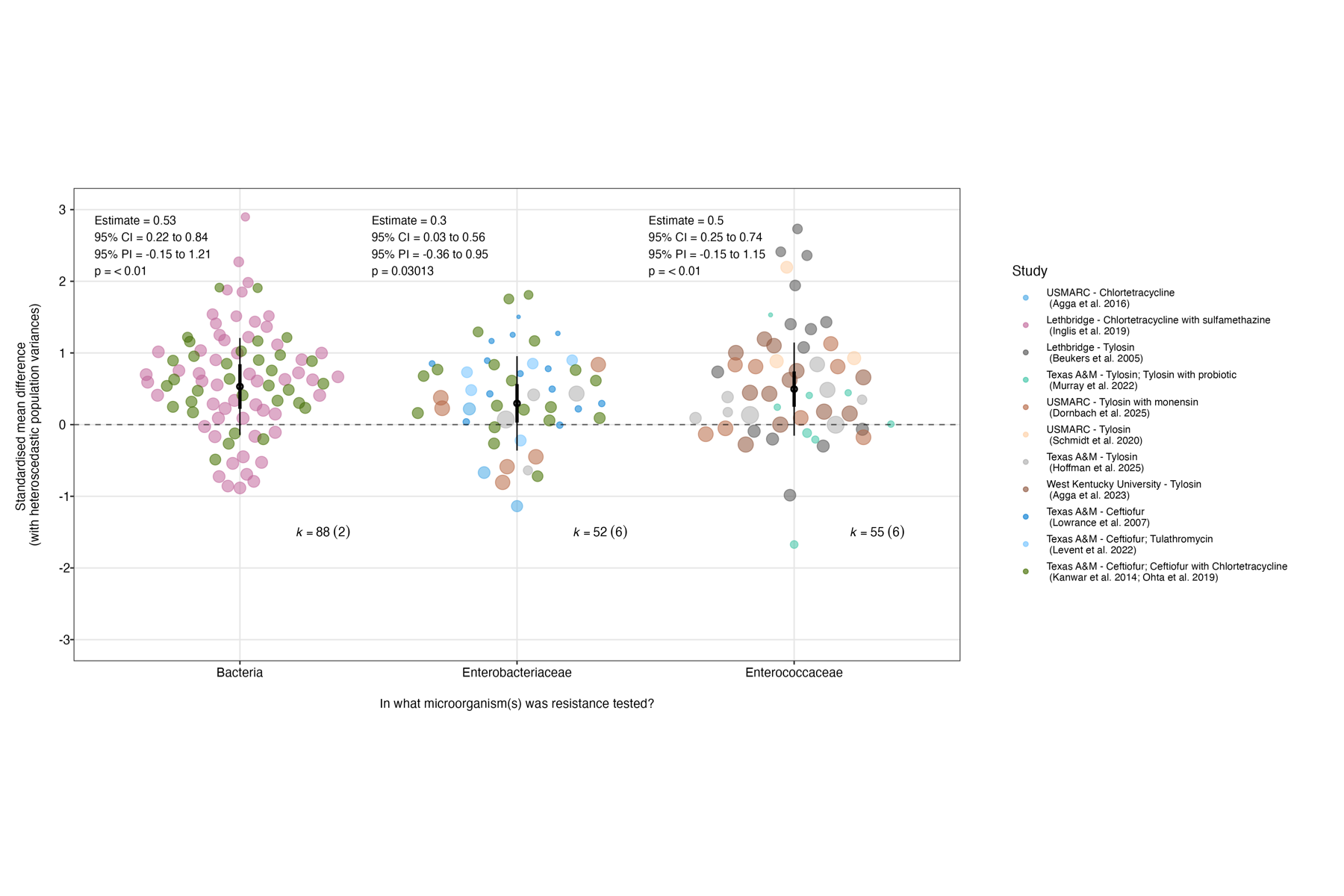

Supplementary Figure 45: Orchard plot representing results of the meta-analytical model testing for a subgroup effect of the method used to estimate the proportion of the targeted bacterial population that was antibiotic resistant in the outcome assessment. Points are effect sizes, with their colours representing the studies from which they are derived and their size being proportional to their inverse standard error (1/SE; small points are less precise estimates, larger points are more precise estimates). The pooled effect for each subgroup (coefficient after accounting for random effects) is represented by the 'trunk' of the tree (black circle), with its confidence interval represented its 'branch' (shorter, thicker line through it), and its prediction interval represented by its 'twig' (longer, thinner line through it). For the numbers above each subgroup, 'k' represents the number of effect sizes, whilst the number in brackets represents the number of studies from which they were derived.

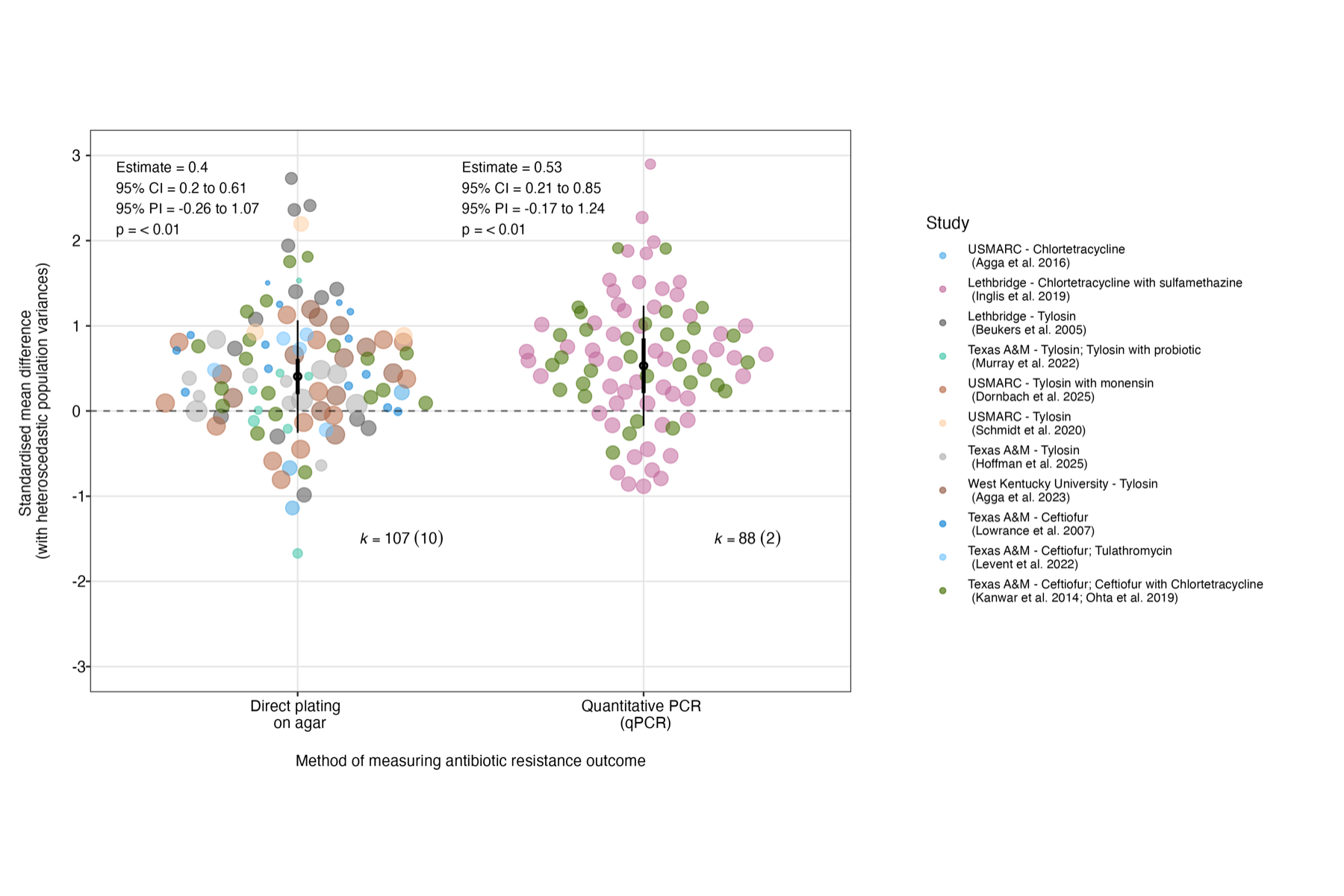

Supplementary Figure 46: Orchard plot representing results of the meta-analytical model testing for a sub-subgroup effect of the antibiotic onto which samples were plated in CFU measures of the proportion of the targeted bacterial population that was antibiotic resistant in the outcome assessment. Points are effect sizes, with their colours representing the studies from which they are derived and their size being proportional to their inverse standard error (1/SE; small points are less precise estimates, larger points are more precise estimates). The pooled effect for each subgroup (coefficient after accounting for random effects) is represented by the 'trunk' of the tree (black circle), with its confidence interval represented its 'branch' (shorter, thicker line through it), and its prediction interval represented by its 'twig' (longer, thinner line through it). For the numbers above each subgroup, 'k' represents the number of effect sizes, whilst the number in brackets represents the number of studies from which they were derived.

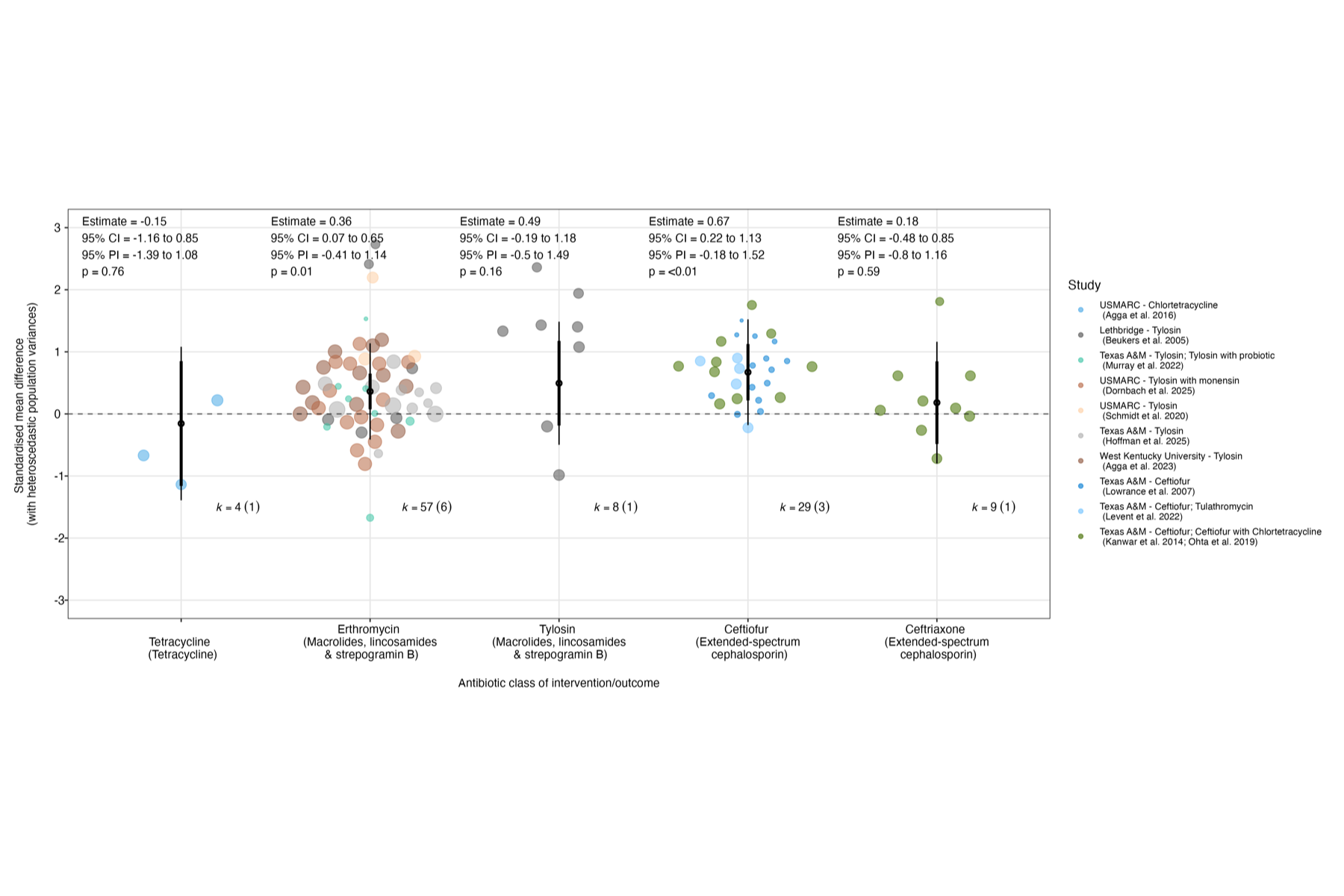

Supplementary Figure 47: Orchard plot representing results of the meta-analytical model testing for a sub-subgroup effect of the antibiotic resistance genes targeted in qPCR measures of the proportion of the targeted bacterial population that was antibiotic resistant in the outcome assessment. Points are effect sizes, with their colours representing the studies from which they are derived and their size being proportional to their inverse standard error (1/SE; small points are less precise estimates, larger points are more precise estimates). The pooled effect for each subgroup (coefficient after accounting for random effects) is represented by the 'trunk' of the tree (black circle), with its confidence interval represented its 'branch' (shorter, thicker line through it), and its prediction interval represented by its 'twig' (longer, thinner line through it). For the numbers above each subgroup, 'k' represents the number of effect sizes, whilst the number in brackets represents the number of studies from which they were derived.

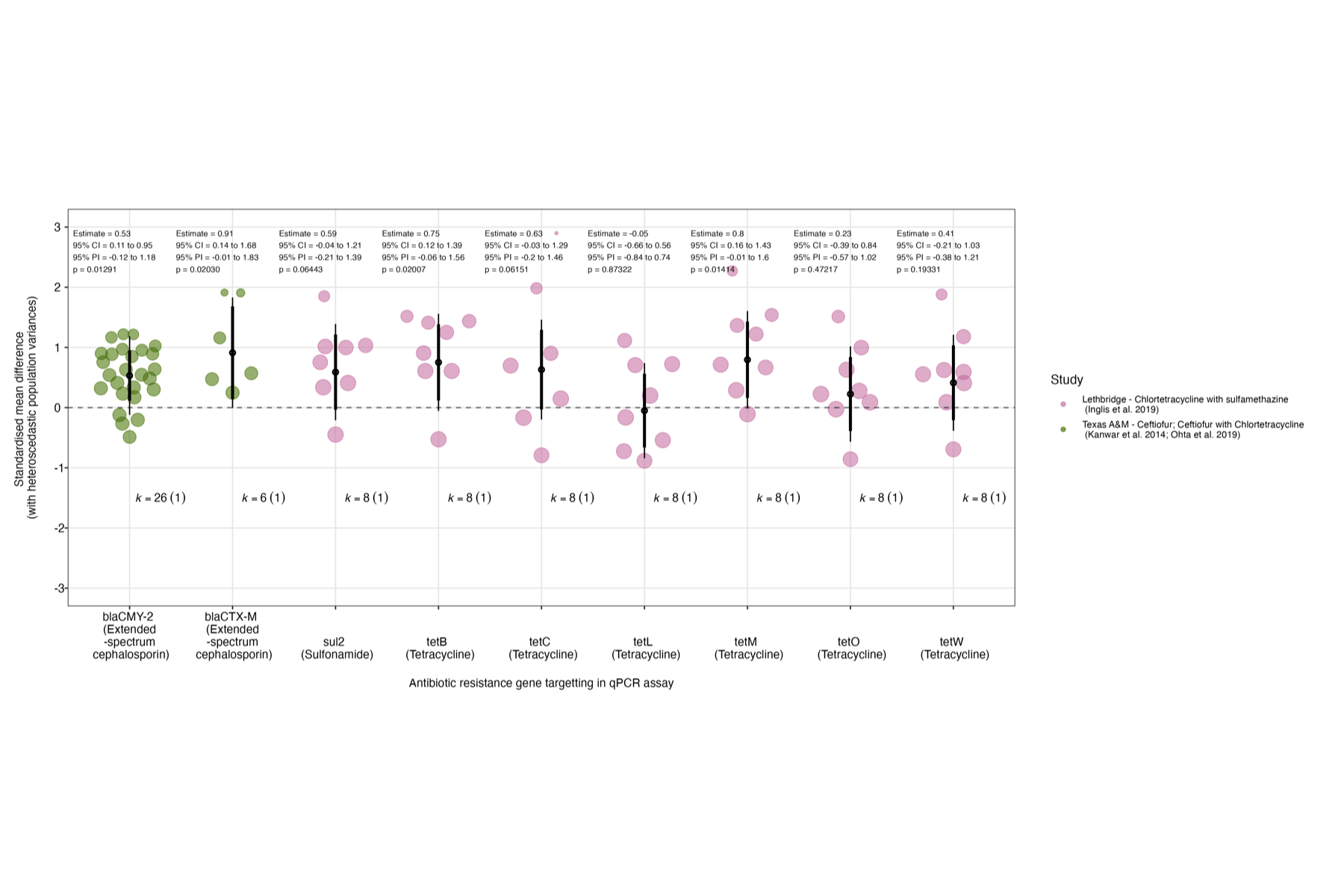

Supplementary Figure 48: Orchard representing results of the second meta-analytical model testing for a sub-subgroup effect of the relative (to the CLSI breakpoint) antibiotic concentration onto which samples were plated in CFU measures of the proportion of the targeted bacterial population that was antibiotic resistant in the outcome assessment(i.e. study breakpoint relative to CLSI breakpoint). Given the uneven spread of concentrations in the previous analysis, for this analysis we categorised antibiotuc concentrations/study breakpoints into four categories based on whether they were below, at, or above the CLSI breakpoint (or there was no CLSI breakpoint for the antibiotic-microorganism combination tested). Points are effect sizes, with their colours representing the studies from which they are derived and their size being proportional to their inverse standard error (1/SE; small points are less precise estimates, larger points are more precise estimates). The pooled effect for each subgroup (coefficient after accounting for random effects) is represented by the 'trunk' of the tree (black circle), with its confidence interval represented its 'branch' (shorter, thicker line through it), and its prediction interval represented by its 'twig' (longer, thinner line through it). For the numbers above each subgroup, 'k' represents the number of effect sizes, whilst the number in brackets represents the number of studies from which they were derived.

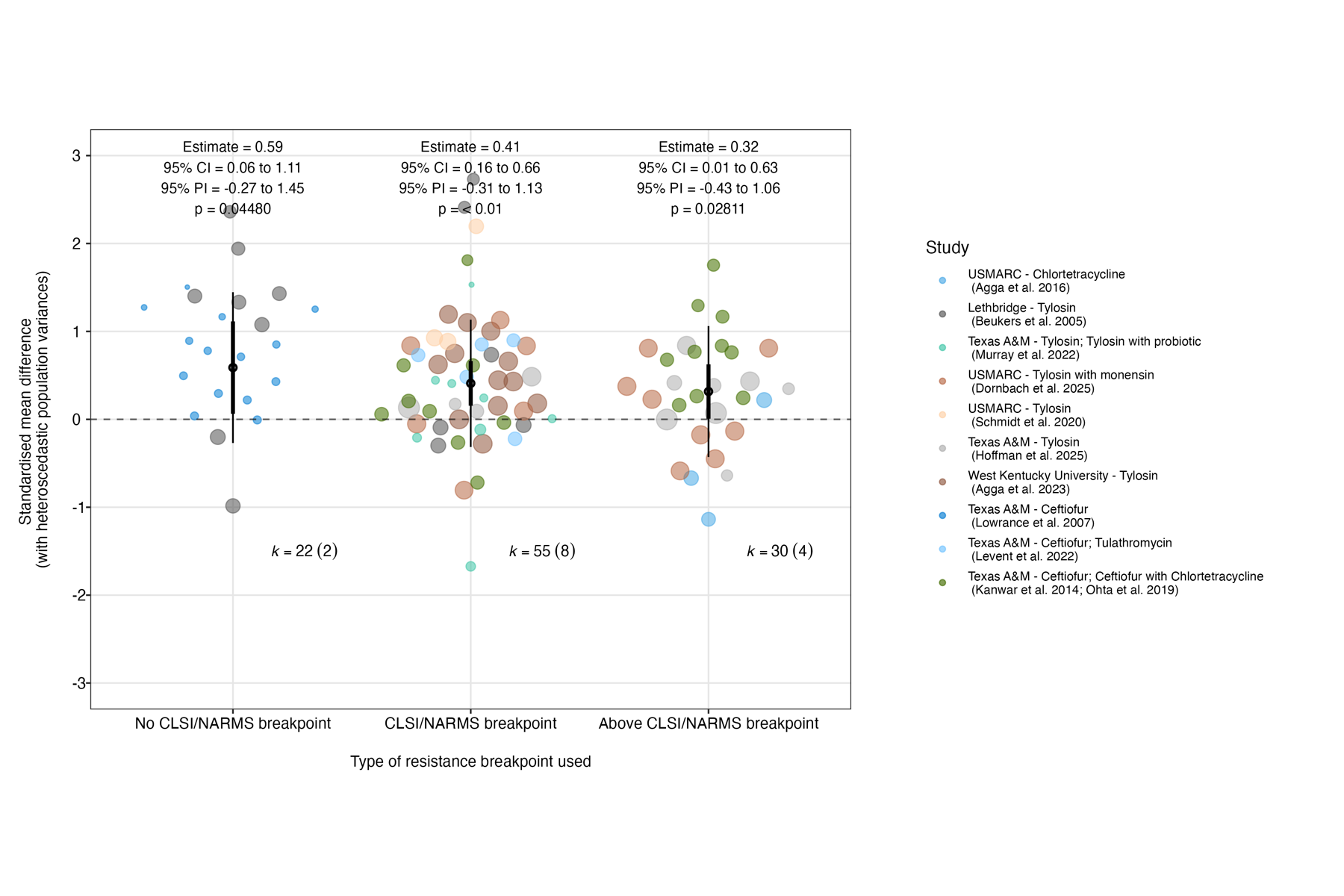

Supplementary Figure 49: Bubble plot representing results of the meta-analytical model testing for a sub-subgroup effect of the relative (to the CLSI breakpoint) antibiotic concentration onto which samples were plated in CFU measures of the proportion of the targeted bacterial population that was antibiotic resistant in the outcome assessment(i.e. study breakpoint relative to CLSI breakpoint). Points are effect sizes, with their colours representing the studies from which they are derived and their size being proportional to their inverse standard error (1/SE; small points are less precise estimates, larger points are more precise estimates).

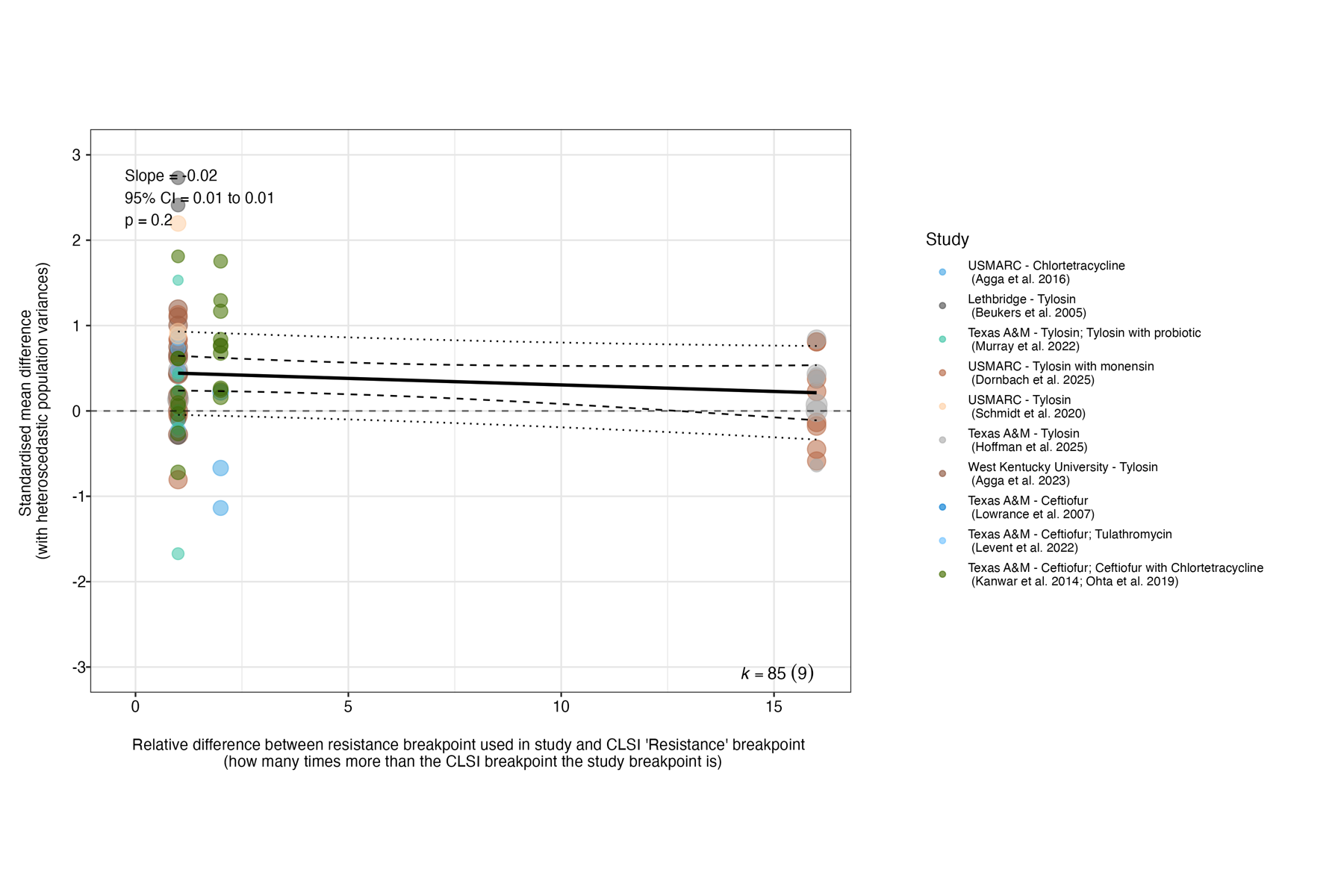

Supplementary Figure 50: Bubble plot representing results of the meta-analytical model testing for a sub-subgroup effect of the absolute log10-transformed antibiotic concentration onto which samples were plated in CFU measures of the proportion of the targeted bacterial population that was antibiotic resistant in the outcome assessment (i.e. study breakpoint). Points are effect sizes, with their colours representing the studies from which they are derived and their size being proportional to their inverse standard error (1/SE; small points are less precise estimates, larger points are more precise estimates).

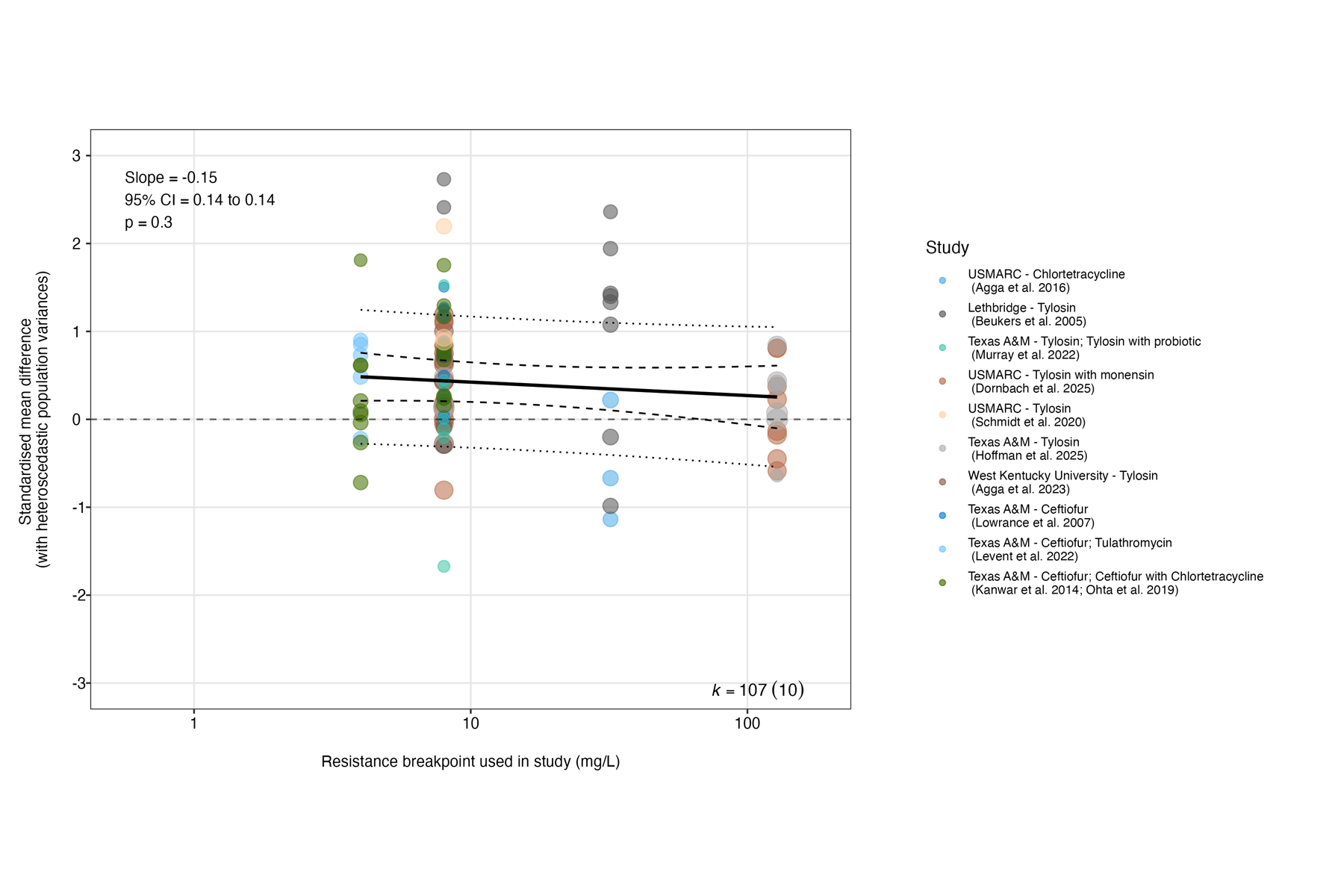

Supplementary Figure 51: Orchard plot representing results of the meta-analytical model testing for a subgroup effect of whether or not the outcomes underpinning effect sizes accounted for potential pen-level effects. Outcomes were classified according to the study design and analysis underpinning any effect sizes used in the meta-analysis. Briefly, 'Individual (pen effect not accounted for)' refers to effect sizes from studies in which there was a single pen per study arm/one pen for all study arms; 'Individual (pen effect not account for)' refers to extracted effect sizes from analyses at the individual level but which did not account for the fact that study arms are spread across different pens; 'Individual (pen as random effect)' refers to extracted effect sizes from analyses at the individual level but which did account for the fact that study arms are spread across different pens by using pen as a random effect in the model; 'Pen (individuals as pseudoreplicates)' refers to effect sizes from analyses at the pen level i.e. which average out individual-level variation by treating individuals as pseudoreplicates (this was the method we used to process raw data where available, hence it is the largest category). Points are effect sizes, with their colours representing the studies from which they are derived and their size being proportional to their inverse standard error (1/SE; small points are less precise estimates, larger points are more precise estimates). The pooled effect for each subgroup (coefficient after accounting for random effects) is represented by the 'trunk' of the tree (black circle), with its confidence interval represented its 'branch' (shorter, thicker line through it), and its prediction interval represented by its 'twig' (longer, thinner line through it). For the numbers above each subgroup, 'k' represents the number of effect sizes, whilst the number in brackets represents the number of studies from which they were derived.

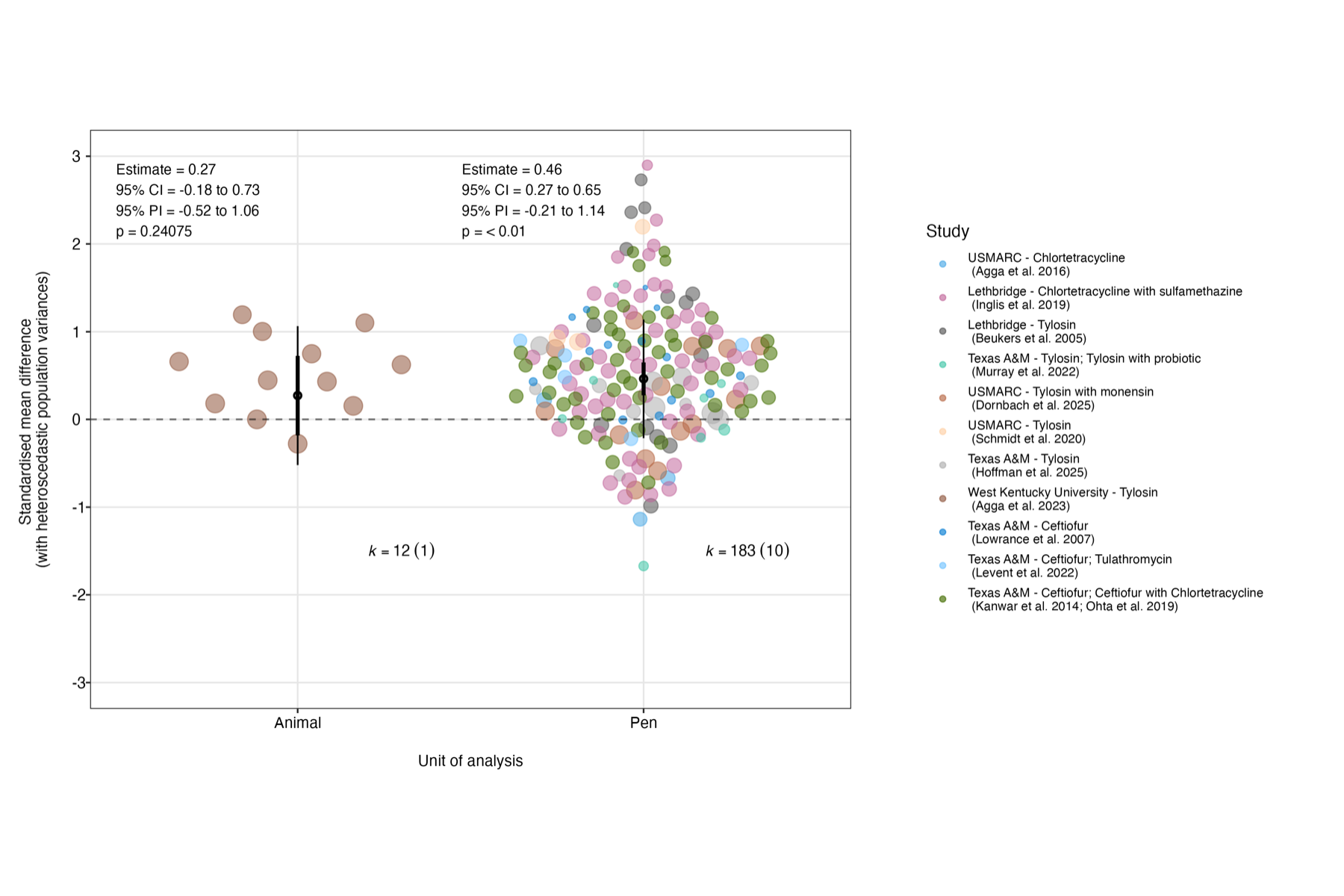

Supplementary Figure 52: Orchard plot representing results of the meta-analytical model testing for a subgroup effect of study design. Points are effect sizes, with their colours representing the studies from which they are derived and their size being proportional to their inverse standard error (1/SE; small points are less precise estimates, larger points are more precise estimates). The pooled effect for each subgroup (coefficient after accounting for random effects) is represented by the 'trunk' of the tree (black circle), with its confidence interval represented its 'branch' (shorter, thicker line through it), and its prediction interval represented by its 'twig' (longer, thinner line through it). For the numbers above each subgroup, 'k' represents the number of effect sizes, whilst the number in brackets represents the number of studies from which they were derived.

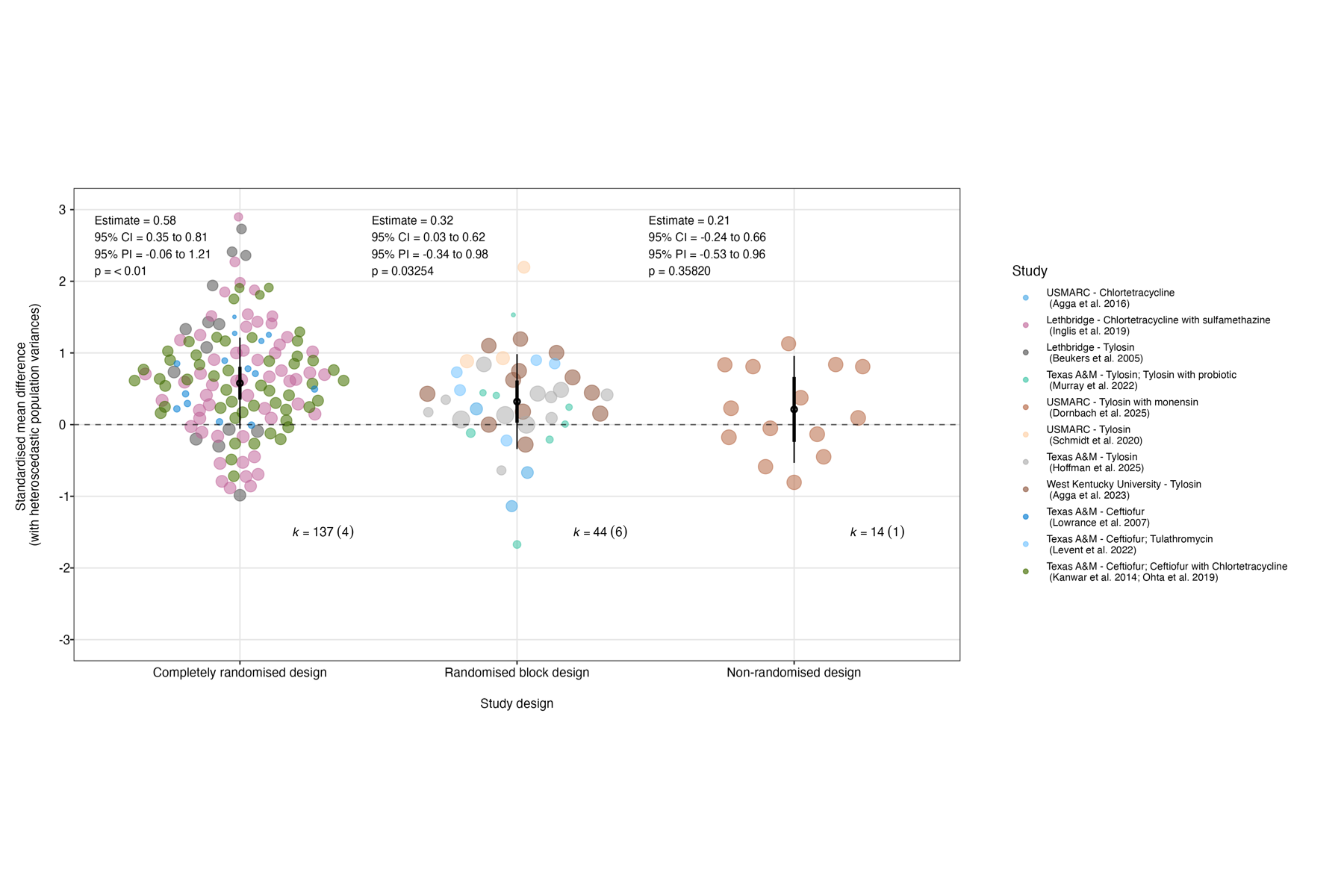

#### 2.5 Reporting Biases

Supplementary Figure 53: Publication bias plots

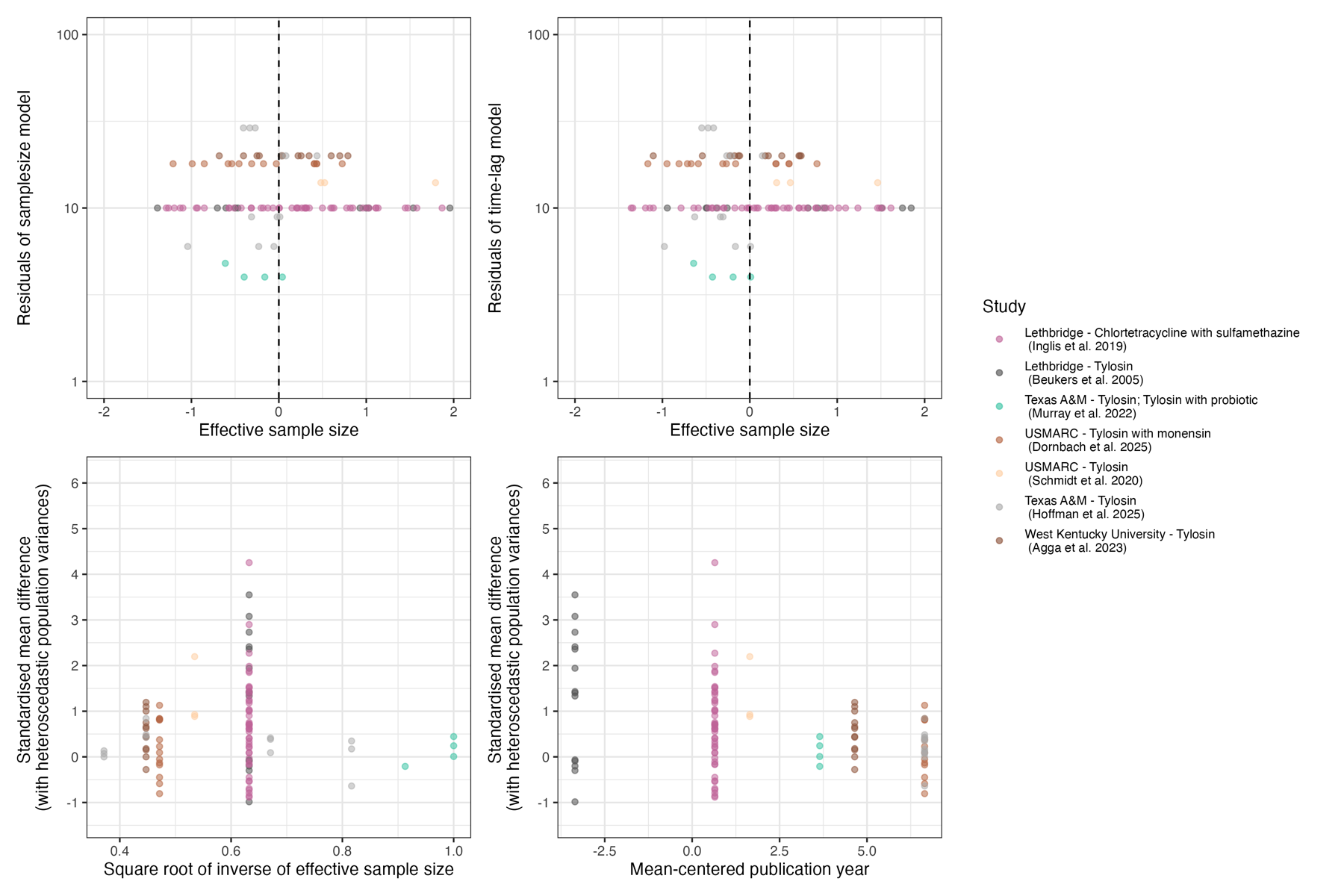

Supplementary Figure 54: Publication bias plots

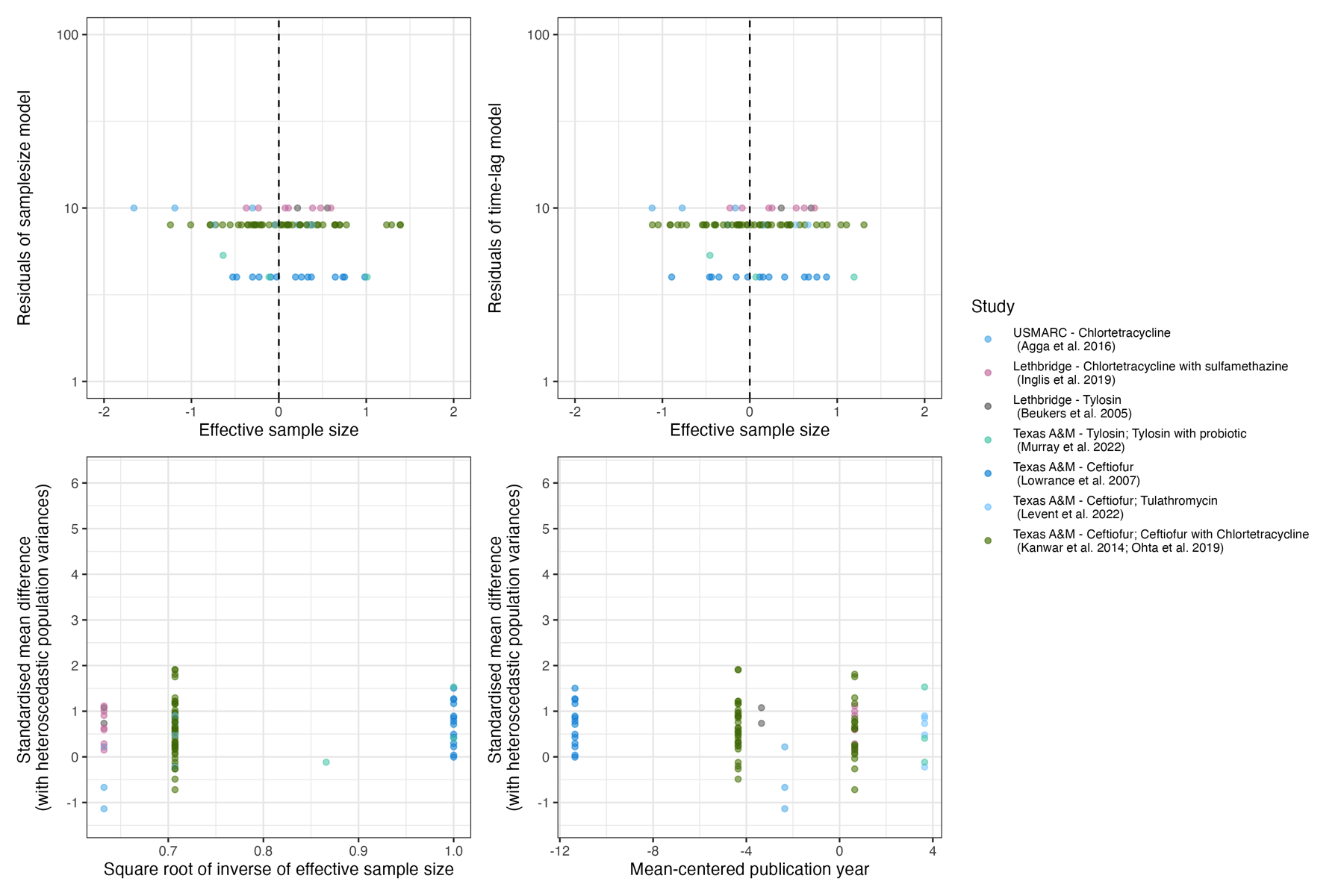

#### 2.6 Certainty assessment

Supplementary Table 4: Certainty assessment table

| **No. of studies** | **Outcome (N.B. Confidence is assessed for size of the effect)** | **Studies/study designs** | **Starting** | **Downgrading domains** | | | | | | **Upgrading domains** | | | **Certainty of the evidence (GRADE)** | **Comments** |
| --- | --- | --- | --- | --- | --- | --- | --- | --- | --- | --- | --- | --- | --- | --- |
|  |  |  |  | **Risk of bias** | **Inconsistency** | **Indirectness** | **Imprecision** | **Publication bias** | **Score after downgrading** | **Large effects** | **Dose-response** | **Plausible confounding** |  |  |
| 7 | Faecal density of resistance determinants for a particular antibiotic class during its administration to beef cattle | 7 (3 completely randomised; 3 randomised complete block; 1 non-randomised) | High certainty (4) | Serious limitations (-1)^a^ | Serious limitations (-1)^b^ | No serious limitations (0)^d^ | No serious limitations (0)^f^ | No serious limitations (0)g | Low (3) | No^i^ | No^j^ | No^k^ | Low (3) | Downgraded for risk of bias (due to universal lack of blinding and pre-registration, amongst other things) and inconsistency (due to results of heterogeneity tests) |
| 7 | Relationship between faecal density of resistance determinants for a particular antibiotic class and days of its continuous administration to beef cattle | 7 (3 completely randomised; 3 randomised complete block; 1 non-randomised) | High certainty (4) | Serious limitations (-1)^a^ | Serious limitations (-1)^b^ | No serious limitations (0)^d^ | No serious limitations (0)^f^ | No serious limitations (0)h | Low (3) | No^i^ | No^j^ | No^k^ | Low (3) | Downgraded for risk of bias (due to universal lack of blinding and pre-registration, amongst other things) and inconsistency (due to results of heterogeneity tests) |
| 8 | Faecal density of resistance determinants for a particular antibiotic class after its administration to beef cattle | 7 (4 completely randomised; 3 randomised complete block) | High certainty (4) | Serious limitations (-1)^a^ | No serious limitations (0)^c^ | Serious limitations (-1)^e^ | No serious limitations (0)^f^ | No serious limitations (0)g | Low (3) | No^i^ | No^j^ | No^k^ | Low (3) | Downgraded for risk of bias (due to universal lack of blinding and pre-registration, amongst other things) and indirectness (due to focus on MLSB and extended spectrum cephalosporin intereventions) |
| 8 | Relationship between faecal density of resistance determinants for a particular antibiotic class and days after its administration to beef cattle | 7 (4 completely randomised; 3 randomised complete block) | High certainty (4) | Serious limitations (-1)^a^ | No serious limitations (0)^c^ | Serious limitations (-1)^e^ | No serious limitations (0)^f^ | No serious limitations (0)g | Low (3) | No^i^ | No^j^ | No^k^ | Low (3) | Downgraded for risk of bias (due to universal lack of blinding and pre-registration, amongst other things) and indirectness (due to focus on MLSB and extended spectrum cephalosporin intereventions) |
